# Metatranscriptomic analysis of common mosquito vector species in the Canadian Prairies

**DOI:** 10.1101/2023.08.15.553461

**Authors:** Cole Baril, Bryan J. Cassone

## Abstract

The microbiome plays vital roles in the life history of mosquitoes, including their development, immunity, longevity, and vector competence. Recent advances in sequencing technologies have allowed for detailed exploration into the diverse microorganisms harboured by these medically important insects. Although these meta-studies have catalogued the microbiomes of mosquitoes on several continents, much of the information currently available for North America is limited to the state of California. In this study, we collected >35,000 mosquitoes throughout Manitoba, Canada over a two-year period, and then harnessed RNA sequencing and targeted RT-PCR to characterize the microbiomes of the eight most pervasive and important vector and pest species. The consensus microbiome of each species was overwhelmingly composed of viruses, but also included fungi, bacteria, protozoa, and parasitic invertebrates. The microbial assemblages were heterogeneous between species, even within the same genus. We detected notable pathogens, including the causal agents of Cache Valley Fever, avian malaria, and canine heartworm. The remaining microbiome consisted largely of putatively insect-specific viruses that are not well characterized, including 17 newly discovered viruses from 10 different families. Future research should focus on evaluating the potential application of these viruses in biocontrol, as biomarkers, and/or in disrupting mosquito vectorial capacity. Interestingly, we also detected viruses that naturally infect honeybees and thrips, which were presumably acquired indirectly through nectar foraging behaviours. Overall, we provide the first comprehensive catalogue of the microorganisms harboured by the most common and important mosquito vectors and pests in the Canadian Prairies.

## Introduction

Microbiome refers to the assemblage of microorganisms harboured by a given organism, including viruses, bacteria, protozoans, and fungi (Huang et al. 2020; Cansado-Utrilla et al. 2021). A range of biotic and abiotic factors can have profound affect on the microbiome, which in turn may impact the life history of the host. For mosquitoes, their microbiome plays critical roles in development, longevity, immunity, and vector competence (Minard et al. 2013; Jupatanakul et al. 2014; Hegde et al. 2015; Guégan et al. 2018; Strand 2018; Caragata et al. 2019; Cansado-Utrilla et al. 2021; Dada 2021). Relatively recent advances in sequencing technologies have greatly expanded our understanding of the highly diverse and dynamic microbial flora carried by mosquitoes (Shi et al. 2019; Ramos-Nino et al. 2020; Wang et al. 2021; Batson et al. 2021; Zhao et al. 2022). These studies indicate that the microbiome can vary among different mosquito species, within populations of the same species, and even among individual mosquitoes within populations. Consequently, cataloguing the microbial communities of notable mosquito vector species within a given geographical region is likely to provide key insights into their life history traits with potential applications for disease and pest control.

A major component of the mosquito microbiome are viruses (i.e., the virome), some of which (i.e., arboviruses) may be transmitted to humans, livestock, and other animals (de Almeida et al. 2021; Hameed et al. 2021; Moonen et al. 2023). Of the ∼3,500 extant mosquito species, only a small proportion vector viruses of public health or veterinary importance, with the majority belonging to the genera *Aedes* and *Culex* (Tolle 2009). In the Canadian Prairies, the most ubiquitous mosquito vector is the inland floodwater mosquito, *Aedes vexans* Meigen (Baril et al. 2023). This species is capable of transmitting West Nile virus (WNV), California serogroup viruses (CSGVs), Zika virus, and Rift Valley fever virus (Drebot 2015; Weissmann 2016; O’Donnell et al. 2017; Parry et al. 2020). The summer saltmarsh mosquito, *Ochlerotatus dorsalis* Meigen, and *Culex tarsalis* Coquillett are also commonly found species in the Prairies. Both are competent vectors of Western equine encephalitis virus (WEEV), WNV, and CSGVs (Wood et al. 1979; Anderson et al. 2015). The cattail mosquito, *Coquillettidia perturbans* Walker, is distributed across the Prairies and typically breeds in permanent swamps where cattails and other aquatic plants are present (Wood et al. 1979). In addition to WNV and CSGVs, this mosquito is a carrier of Eastern equine encephalitis virus (EEEV) (Wood et al. 1979; Andreadis et al. 2008). Other mosquito vector species occurring in the Prairies include *Aedes canadensis* Theobald (CSGVs, WNV, EEEV), *Ochlerotatus triseriatus* Say (La Crosse virus, EEEV, WEEV), *Ochlerotatus flavescens* Müller (canine heartworm, *Dirofilaria immitis*) (Wood et al. 1979; Berry et al. 1986; McMahon et al. 2008; Anderson et al. 2015; Koloski et al. 2021). It should be noted that *Ochlerotatus* was previously ranked as a subgenus of *Aedes*, but has since been reclassified as a distinct genus (Reinert et al. 2004).

In addition to arboviruses, the mosquito virome includes insect-specific viruses (ISVs) (Bolling et al., 2015). As their name suggests, these viruses establish an infection in mosquitoes and/or other insects but are incapable of replicating in vertebrate hosts. These viruses are thought to be vertically transmitted transovarially by mosquitoes from infected females to their offspring (Saiyasombat et al. 20111; Bolling et al. 2011; Haddow et al. 2013), though the mechanisms by which ISVs establish an infection in the mosquito is not fully known (Öhlund et al. 2019). Insect-specific viruses have been recognized for decades (Stoller and Thomas 1975), but were vastly understudied until recent advancements in microbiome research, sequencing technologies, bioinformatics tools, and phylogenetic analyses. Indeed, the genomes or partial genomes of hundreds of ISVs from diverse viral families (e.g., *Bunyaviridae, Flaviviridae, Reoviridae, Rhabdoviridae, Togaviridae, Birnaviridae, Nodaviridae*, *Phenuiviridae* and *Mesoniviridae*) have now been sequenced and characterized from various geographical locations (for review see Atoni et al. 2019; Carvalho and Long 2021). Despite their strict host tropism with no (known) direct implications to the burden of infectious diseases, ISVs may impact/regulate mosquito vector competence (Bolling 2012; Schultz et al. 2018; Öhlund et al. 2019; Olmo et al. 2023) and in some cases may be used in biocontrol (Chen et al. 2023) or serve as effective biomarkers for viruses of public health concern (Nouri et al. 2018; Martin et al. 2019).

Two Canadian Prairie provinces (Manitoba and Saskatchewan) carry out mosquito surveillance activities annually to identify and assess the prevalence of arboviruses carried by vector species. However, these provincial programs are mostly limited to monitoring *Cx. tarsalis* for WNV infection using serological and/or reverse transcriptase-polymerase chain reaction (RT-PCR)-based methodologies. Consequently, there may be other arboviruses contributing to an under-recognized burden of disease within this region. For instance, clinical cases of CSGVs have been reported in Manitoba (Lau et al. 2017; Vosoughi et al. 2018) and mosquitoes harbouring the viruses have been detected in an adjacent U.S. state (Anderson et al. 2015). Further, nothing is known regarding the ISVs or other microorganisms (e.g., protozoa, fungi and bacteria) that may contribute to the microbiome of common vector species in the Canadian Prairies. Metatranscriptomics has filled in these knowledge gaps for mosquito species in other geographical regions, including studies from Asia (Shi et al. 2016), Australia (Shi et al. 2017), United States (Batson et al. 2021; Chandler et al. 2015; Sadeghi et al. 2018), and the Caribbean (Shi et al. 2019). With this in mind, we carried out RNA sequencing on over 35,000 mosquitoes collected throughout Manitoba, Canada, over a three-year period. The microbiomes of eight of the most common vector and pest species were characterized, which included an array of viruses (arboviruses, ISVs and novel viruses) as well as bacteria, fungi, protozoa, and invertebrate parasites. We also carried out targeted RT-PCR-based diagnostics to assess the prevalence of CSGVs in the competent vector species.

## Materials and Methods

### Mosquito collections and identification

Host-seeking females were trapped between June and August in 2020 and 2021, as previously described (Baril et al. 2023a). In brief, CDC Miniature Light Traps (Model 1012, John W. Hock, Gainesville, FL) were placed on tree limbs ∼1.5 m from the ground with the carbon dioxide (CO2) regulators set to 15 psi and the light disabled. Traps were activated from dusk until dawn twice weekly (Monday and Tuesday) in eight West Manitoba communities. In 2020, the City of Winnipeg Insect Control Branch provided us one-time satellite traps from nine additional locations in Central and East Manitoba. In 2019, weekly collections were also done at 4 sites located in Brandon, Manitoba, between July and August.

Mosquito vector species were visually identified using dissecting microscopes and applicable identification keys (Carpenter and LaCasee 1955; Wood et al. 1979; Thielman and Hunter 2007). Eight target species were sorted out with the remaining specimens omitted from our study: *Aedes vexans* Meigen, *Aedes canadensis* Theobald, Ochlerotatus dorsalis Meigen, *Ochlerotatus flavescens* Muller, *Ochlerotatus triseriatus* Say, *Culex tarsalis* Coquillett, *Coquillettidia perturbans* Walker and *Anopheles earlei* Vargus. Mosquitoes were then pooled into 1.5 mL tubes containing up to 50 individuals sorted by species, location, and year. Pools were stored at −80 ℃ until molecular analysis.

### RNA isolation and sequencing

Total RNA was extracted from each pooled mosquito sample using the RNeasy Mini Kit (Qiagen, Hilden, Germany), following the manufacturers recommended protocol. The quantity, quality, and concentration of RNA was evaluated using a Nanophotometer NP80 (Implen Inc., Westlake Village, CA, USA). We then combined RNA samples derived from the same species, location, and year to form larger pools for sequencing. In some cases, nearby locations with relatively small mosquito numbers were combined to form a pool. Approximately 2 μg of RNA per pool was sample was sent to Génome Québec Innovation Centre (McGill University, Montreal, QC, Canada) for mRNA library preparation (New England Biolabs, Ipswich, MA, USA) and paired read sequencing (100 bp) on the NovaSeq platform (Illumina, San Diego, CA). We then used CASAVA 1.8.2 to carry out bcl conversions and demultiplexing. Image deconvolution and quality value calculations were performed using the Illumina GA pipeline (version 1.6). The raw sequence reads can be retrieved from the NCBI short sequence read archive under the SRA accession number PRJNA793247.

### Read processing and *de novo* assembly

The Chan Zuckerberg ID Metagenomic Pipeline v6.8 (Chan Zuckerberg Biohub; CZID), an open-sourced cloud-based bioinformatics platform (https://czid.org/), was used for quality control and host filtration of reads, assembly *de novo*, and taxonomic binning as described by Batson et al., (2021) and Kalantar et al. (2020). Each sample (i.e. mosquito pool) was analyzed individually, as described below. The CZID pipeline employs STAR and Bowtie2 to perform host filtration (human and mosquito), We used Trimmomatic for adapter trimming, Price Seq for removal of low-quality reads, LZW for the removal of low complexity reads, and CZIDdedup for duplicate read identification.

Preprocessed reads were then assembled *de novo* into contig sequences using two algorithms independently: 1) SPADES with default settings in CZID and 2) CLC Genomics workbench version 20 (CLC Bio, Aarhus, Denmark) with the following parameters: mismatch cost = 2, insertion cost = 3, deletion cost = 3, length fraction of 0.7, and similarity fraction = 0.95 (default settings therein). Exploratory functional annotation and pairwise BLASTn between assemblies indicated a high similarity between algorithms, thus, we opted to use CLC for subsequent analyses.

### Microorganism identification

To identify sequences of microbial origin, contigs were subjected to desktop-downloaded BLASTn and tBLASTx searches against the NCBI nucleotide (nt) and non-redundant (nr) databases, respectively. Each contig was also mapped against the local NCBI virus blast (curated via Entrez Query) using CLC and a customized database omitting improbable viruses (e.g., HIV, influenza). We used the following criterion for a positive hit: minimum E-value of ≤1x 10^-100^, nucleotide and amino acid similarities of >90%, and contig length of ≥250 nt. In addition, a minimum coverage of 10X was used as the cut-off value for a confirmed virus. Contigs meeting these criteria were further scrutinized through analysis of protein function using the NCBI ORFfinder and NCBI conserved domains tools to eliminate possible false positives. Contigs of viral origin with percent amino acid (aa) identities <85% were flagged as potentially novel viruses.

### Targeted screening for California Serogroup viruses

In addition to the high throughput analyses, we screened a subset of samples for CSG viruses using RT-PCR. Reverse transcription was first performed on RNA pools using the RevertAid Kit (Thermo Fisher Scientific Inc., Waltham, MA), following the manufacturer recommended protocol. Amplification of a 251 nt fragment of the S segment was then carried out using Platinum SuperFi PCR Master Mix (Thermo Fisher Scientific Inc.) and the following primer set: BCS82C: 5’-ATGACTGAGTTGGAGTTTCATGATGTCGC-3’ and BCS332V: 5’-TGTTCCTGTTGCCA GGAAAAT-3’ (Kuno et al. 1996). Thermocycler (Biometra TOne, Analytics Jena, Germany) conditions consisted of 39 cycles of 94°C for 1 min, 56°C for 1 min, and 72°C for 1 min, with a final extension at 72°C for 5 min. Amplicons were visualized in 1% agarose gels stained with ethidium bromide using a ChemiDoc Imaging System (Bio-Rad Laboratories, Hercules, CA). We then sent amplicons to the Génome Québec Innovation Centre (McGill University, Montreal, QC, Canada) for purification using a Biomek NX robot with a bead solution and Sanger sequencing of the forward strand using the 3730xl DNA Analyzer (Applied Biosystems, Waltham, MA). Sequences were identified to CSG virus using BLASTn and tBLASTx.

## Results

### Mosquito microbiomes are largely composed of viruses

A total of 45 cDNA libraries representing 35,866 mosquitoes collected throughout Manitoba, Canada were subjected to RNA sequencing (Supplementary Table S1). The number of sequencing libraries and specimens sequenced per species varied considerable and largely reflected their prevalence in our sampling region. We generated more than 3.1 billion paired-end reads of 100 bp; Figure 1 displays the number of reads passing quality control and host filtering for each library.

**Figure 1.**
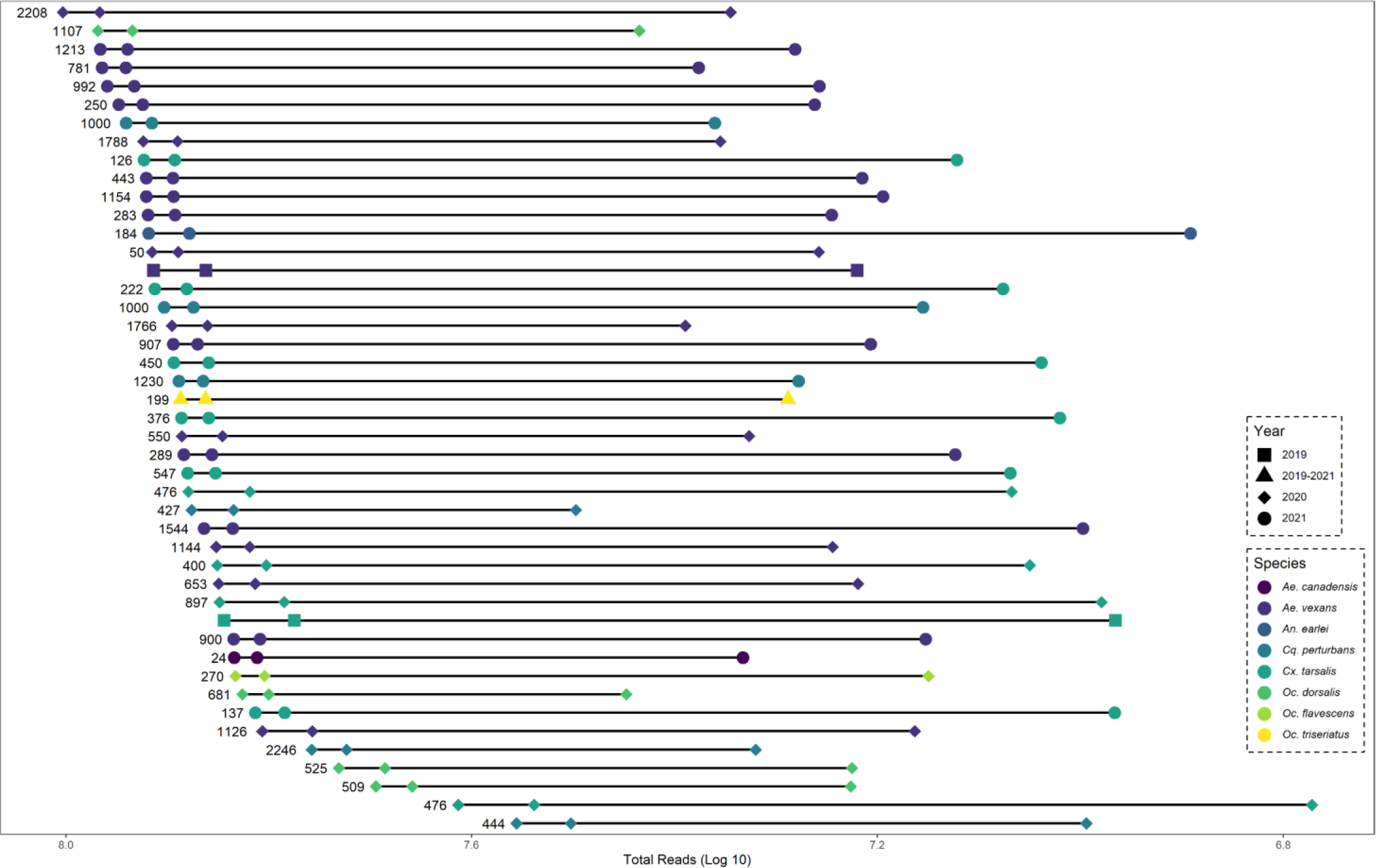
Metadata statistics for each sequencing library (horizontal line). Each library consists of mosquitoes pooled by year, location, and species. Libraries are represented by three icons; total number of reads (left), number of reads after quality filtering (middle), and number reads non-host reads (right). Also labelled for each library is the number of mosquito specimens comprising each RNA pool.

The non-mosquito read subset of each cDNA library was then assembled *de novo* into contigs, functionally annotated via orthologue prediction, and classified to species level (where possible). Our results show an array of different types of organisms associated with the mosquito microbiome, with viruses comprising >99% (5,637,086) of non-host reads mapping to contigs. Our finding of viruses making up the bulk of non-host reads is consistent with other metatranscriptomic studies (Batson et al. 2021). A total of 49 previously reported (i.e. known) viruses were identified in our dataset, which included five types of viral genomes (+ssRNA, - ssRNA, dsDNA, dsRNA, and ssDNA) from 18 families (Table 1). The number of contigs assembled per virus ranged from 1 to 183, with a maximum contig length of 11.7 Kb and a maximum number of reads of nearly 3 million (Supplementary Table S2). Iflaviridae (11) was the most represented viral family, followed by Rhabdoviridae (8), Negevirus (5), and Parvoviridae (4). Nearly half (23) of the viruses were detected in all three years (2019 – 2021), suggesting they commonly infect mosquitoes within the sampling region.

**Table 1.**
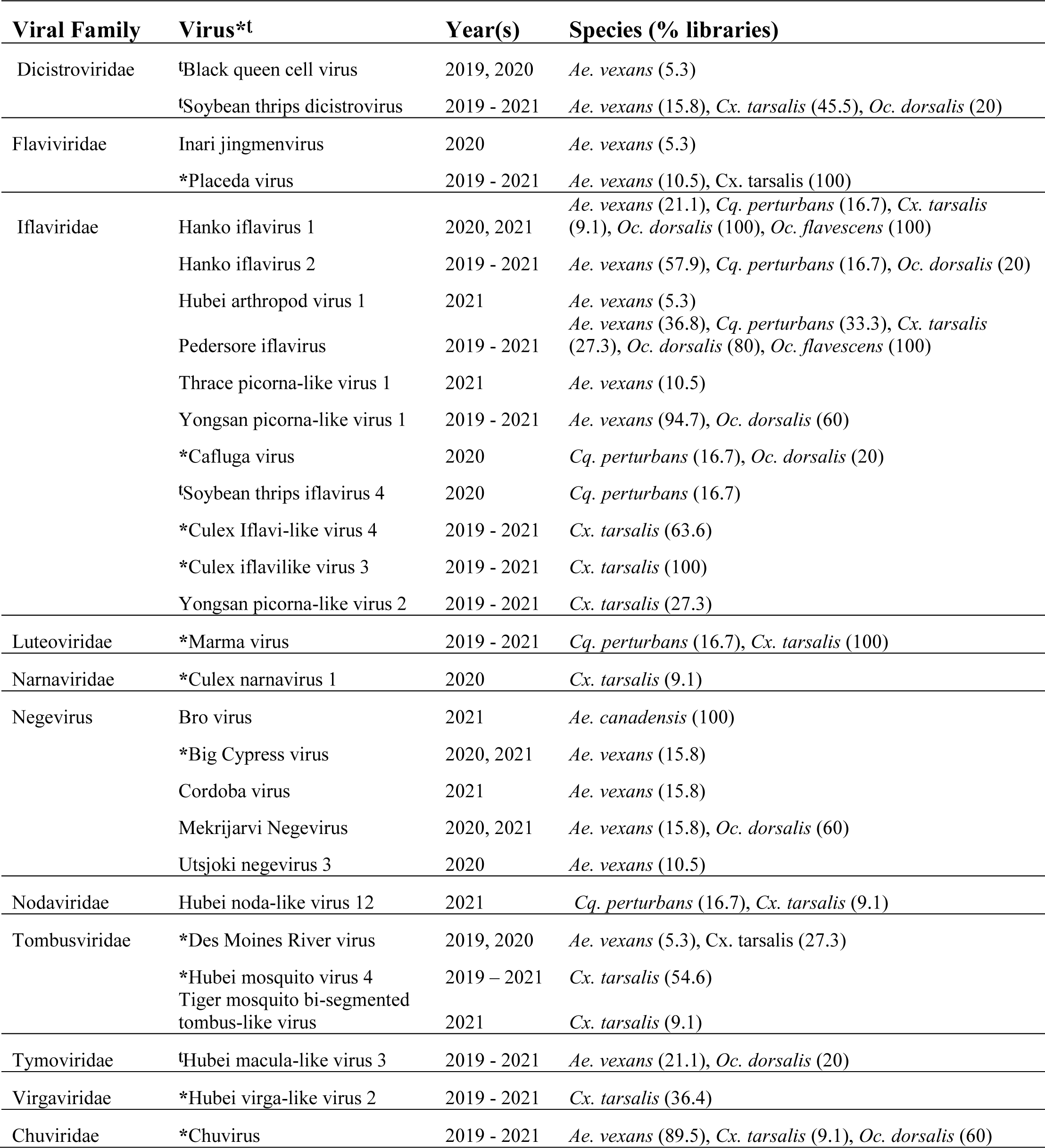
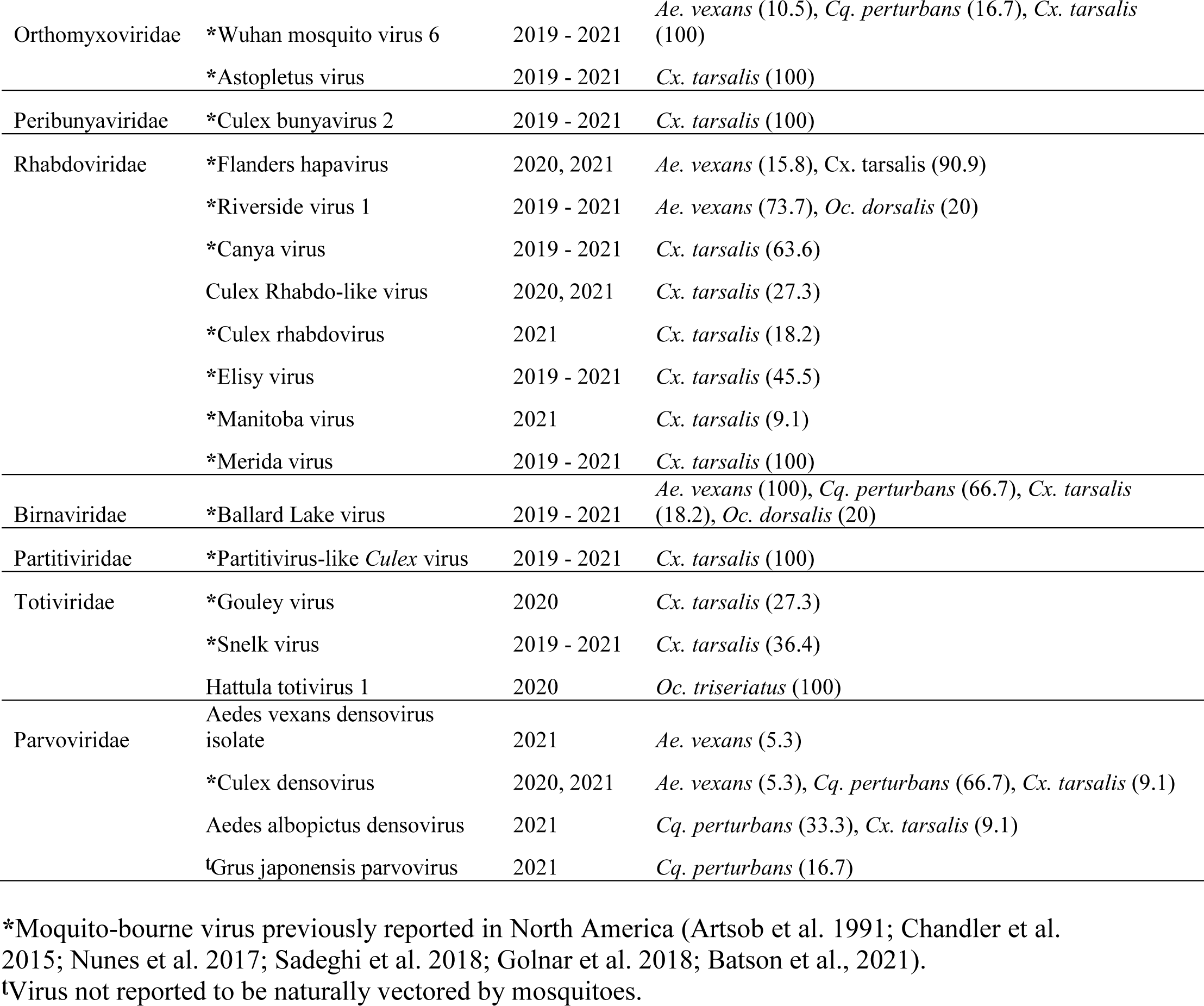
Classification, year(s) reported, and mosquito species each previously reported virus was recovered in.

### Viruses harboured are typically unique to a host species

The proportion of sequencing libraries that each virus was detected in for a given mosquito vector species and year of collection is displayed in Table 1. Some viruses were prevalent in a host species; for instance, Ballard Lake virus was found in all 11 *Ae. vexans* libraries and Merida virus, Astopletus virus, Marma virus, and Placeda virus were identified in each of the 11 *Cx tarsalis* libraries. Others were detected in low frequencies in the mosquito samples, such as Black queen cell virus and Inari jingmenvirus in *Ae. vexans*. The number of viruses and viral families identified in each mosquito species was largely associated with number of libraries sequenced (Figure 2) and to a lesser extent the number of sequencing reads generated (Supplementary Figure S1). Overall, *Cx. tarsalis* harboured the largest number of previously reported viruses (31), followed by *Ae. vexans* (23), *C. perturbans* (12), and *Oc. dorsalis* (11).

**Figure 2.**
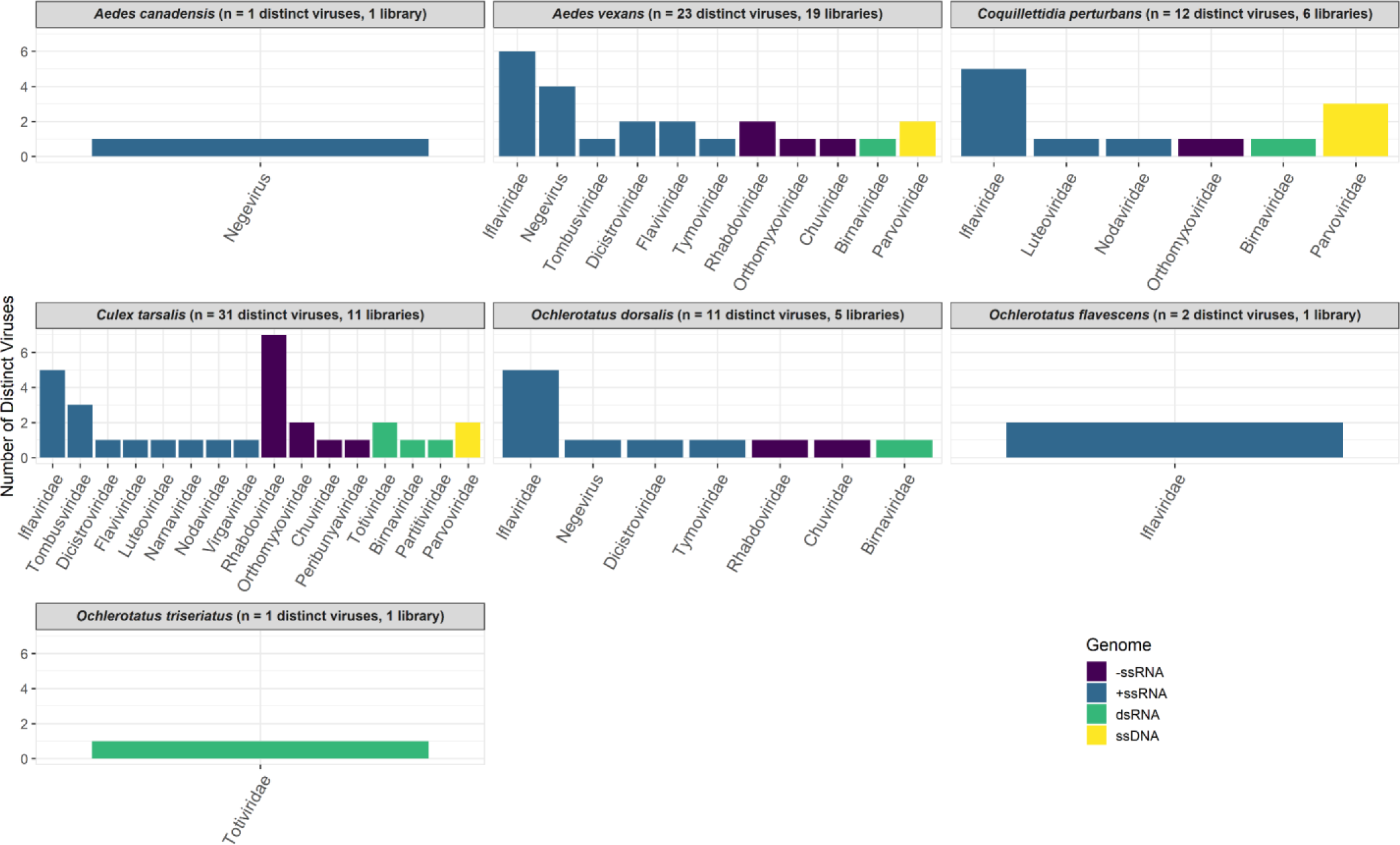
Number of previously reported viruses identified for each mosquito species. Viruses are sorted by family and colour coded based on their genome configuration. Also displayed is the total numbers of viruses detected and sequencing libraries for each species.

The partitioning of viruses by mosquito genera is illustrated diagrammatically in Figure 3. This Venn diagram emphasizes that viruses are largely unique to a given mosquito genus. Indeed, only 39% (19) of viruses were shared among two or more genera, with Hanko iflavirus 1, Pedersore iflavirus, and Ballard Lake virus infecting mosquitoes from all 4 genera. Moreover, there was virus specificity among species within a genus. Both *Aedes* (*Ae. vexans*, *Ae. canadensis*) and *Ochlerotatus* (*Oc. dorsalis*, *Oc. flavescens*) were represented by two mosquito species, and interestingly there were no shared viruses identified among species within each of these genera. No known viruses were detected in *An. earlei*, though this species was represented by the second fewest number of sequenced specimens (184).

**Figure 3.**
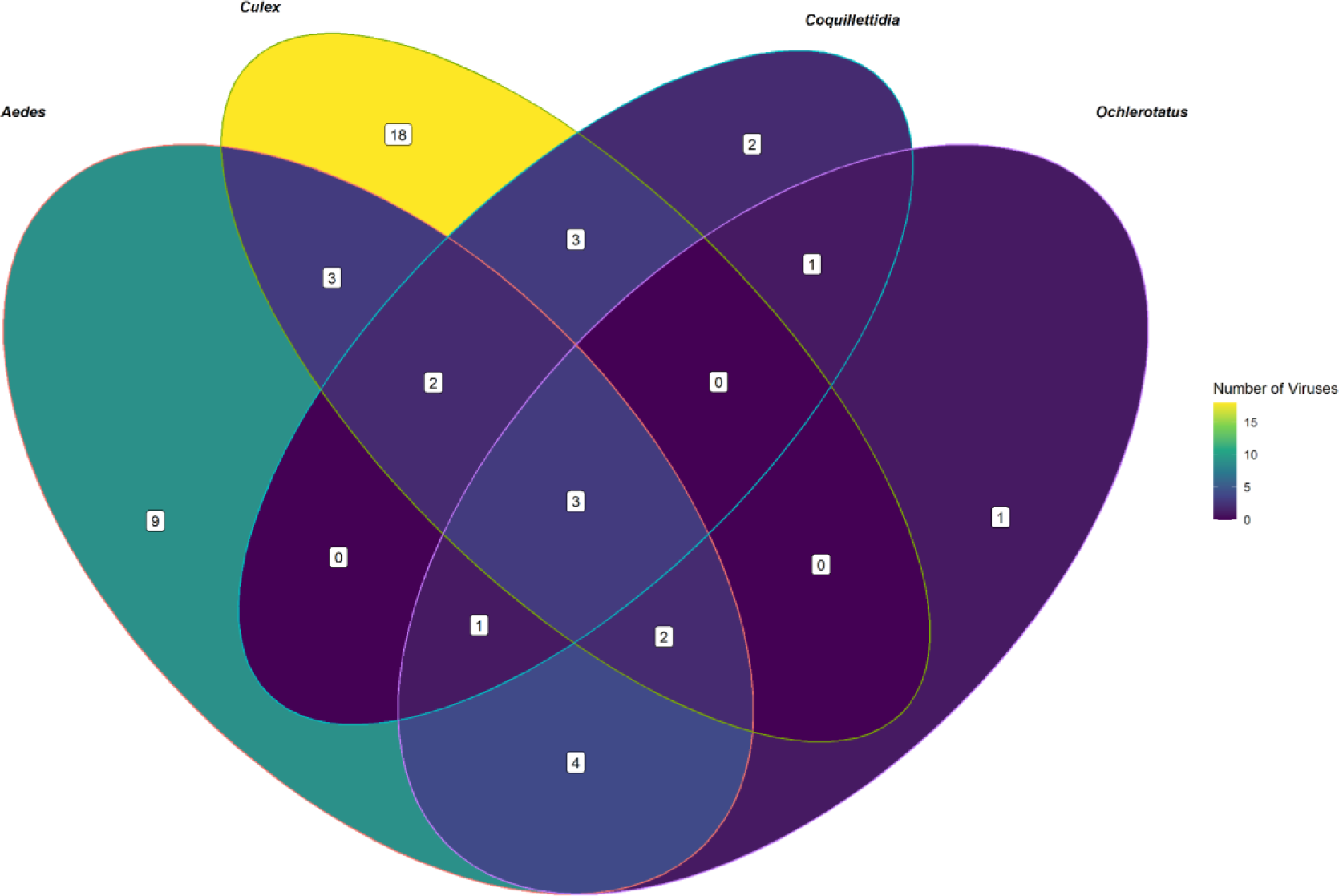
Venn diagram showing the partitioning of previously reported viruses by mosquito genus. The majority of viruses are specific to a given genus.

### Novel viruses were identified in most mosquito vector species

In addition to known viruses, we identified a subset of potentially novel viruses whose contig sequences had amino acid similarities <85% to any organisms currently catalogued in the references databases. We selected this value as the cut-off threshold for the detection of novel viruses, since <90% nucleotide identity is considered a suitable general threshold (Kalantar et al. 2020) and we opted to take a conservative approach in our assignment of previously unreported viruses. We also used ortholog prediction to determine the putative viral genome and family of each novel virus. A total of 17 novel viruses were identified infecting Canadian Prairie mosquitoes, which were represented by 1 to 20 contig sequences per virus (Table 2). While the majority (59%) of these novel viruses were +ssRNA, we also detected putatively –ssRNA, dsRNA, and dsDNA viral genomes. These viruses were classified into 10 different families with Totiviridae, Rhabdoviridae, and Negevirus the most represented at 3 per family. Four of the viruses (Manitoba picorna-like virus 1, Manitoba tombus-like virus 1, Manitoba Rhabdovirus 1, and Manitoba toti-like virus 1) were detected in all three sampling years. Iridoviridae was the only family where we detected a novel virus but did not identify any previously known viruses. Contig sequences for novel viruses have been deposited in the GenBank database (accession numbers OR066146-OR066155), with the complete list of sequences available in Supplementary Table S3.

**Table 2.** Family, year(s) reported, sequencing statistics, and the mosquito species each novel virus was recovered in.

| Viral Family | Virus | Year(s) Reported | No. of Contigs | Longest contig (nt) | Coverage Depth (mean) | Percent aa identity (median)* | Species (% libraries) |
| --- | --- | --- | --- | --- | --- | --- | --- |
| Dicistroviridae | Manitoba dicistro-like virus 1 | 2019 | 1 | 7146 | 149.27 | 82.61 | *Cx. tarsalis* (9.09) |
| Iflaviridae | Manitoba iflavirus 1 | 2020, 2021 | 13 | 2435 | 78.09 | 79.47 | *Ae. vexans* (47.37); *Oc. dorsalis* (20) |
| Iflaviridae | Manitoba picorna-like virus 1 | 2019-2021 | 20 | 2114 | 70.96 | 83.1 | *Ae. vexans* (63.16) |
| Narnaviridae | Manitoba narnavirus 1 | 2021 | 2 | 1760 | 18.77 | 80.59 | *An. earlei* (100) |
| Negevirus | Manitoba mononega-like virus 1 | 2021 | 1 | 11180 | 59.07 | 73.21 | *Ae. canadensis* (100) |
| Negevirus | Manitoba mononega-like virus 2 | 2021 | 1 | 2477 | 20.03 | 75 | *Ae. canadensis* (100) |
| Negevirus | Manitoba mononega-like virus 3 | 2020 | 1 | 5456 | 42.65 | 78.57 | *Oc. triseriatus* (100) |
| Tombusviridae | Manitoba tombus-like virus 1 | 2019-2021 | 5 | 2340 | 157.57 | 76.19 | *Cx. tarsalis* (45.45) |
| Tymoviridae | Manitoba tymo-like virus 1 | 2021 | 1 | 6302 | 23.67 | 66.79 | *Ae. vexans* (5.26) |
| Virgaviridae | Manitoba virgavirus 1 | 2020 | 1 | 1675 | 11.06 | 83.83 | *Oc. dorsalis* (20) |
| Rhabdoviridae | Manitoba Rhabdovirus 1 | 2019-2021 | 20 | 5085 | 26.41 | 77.9 | *Ae. vexans* (73.68); *Oc. dorsalis* (20) |
| Rhabdoviridae | Manitoba rhabdovirus 2 | 2020 | 1 | 4648 | 51.7 | 75 | *Ae. vexans* (5.26) |
| Rhabdoviridae | Manitoba rhabdovirus 3 | 2020, 2021 | 5 | 6071 | 16.51 | 84.62 | *Ae. canadensis* (100); *Cx. tarsalis* (36.36) |
| Iridoviridae | Manitoba iridescent virus 1 | 2021 | 1 | 797 | 16.99 | 78.74 | *Cq. perturbans* (16.67) |
| Totiviridae | Manitoba toti-like virus 1 | 2019-2021 | 5 | 8930 | 39.82 | 68.1 | *Ae. vexans* (10.53); *Cx. tarsalis* (27.27) |
| Totiviridae | Manitoba toti-like virus 2 | 2021 | 1 | 3123 | 13.45 | 60.24 | *Ae. canadensis* (100) |
| Totiviridae | Manitoba toti-like virus 3 | 2020 | 4 | 4626 | 15.11 | 73.5 | *Oc. dorsalis* (80) |

\*All contig sequences for a given virus had amino acid (aa) identities <85% to the NCBI databases.

Supplementary Figure S2 displays the number of novel viruses in each family detected per mosquito species (S2A) and the viruses shared across mosquito genera (S2B). The largest number of novel viruses were identified in *Ae. vexans* (6), followed by *Cx. tarsalis*, Oc. dorsalis, and *Ae. canadensis* (4 per species). Interestingly, both *Ae. canadensis* (4 vs. 1) and *Ae. earlei* (1 vs. 0) were infected with more novel viruses than known viruses. Similar to the known viruses, the novel viruses are largely unique to a given mosquito genus. Only 24% (4) of viruses were shared among two or more genera, with no viruses were found across all 4 genera. Moreover, we did not identify any shared viruses identified among species within a given genus. The only species not infected with a novel virus was *Oc. flavescens*, though this species was represented by a comparatively small number of sequenced specimens (270).

### The non-viral component of the microbiome is significantly smaller

While viruses comprised far and away the largest component of the microbiome for each mosquito species (>99% of non-host sequencing reads), we also detected other sequences of non-host origin (Supplementary Figure S3). The majority of the non-host, non-viral reads generated were fungi (53%), most notably Blastocladiomycota, Microsporidia, and Ascomycota. The second most populous group were invertebrate parasites/protozoa (29%), which included Euglenozoa, Apicomplexa, Nematoda, Acari, and Trematoda. The remaining sequences were derived from bacteria (13%),The Viridiplantae (3%), and Chordata (1%). Notable were sequences from the parasitic roundworm *Dirofilaria immitis* (canine heartworm) isolated from two *Ae. vexans* libraries. Moreover, 25.68% of reads derived from protozoa/parasites were from the Apicomplexan genus *Plasmodium*, specifically *P. gallinaceum* and *P. relictum* (avian malaria) in *Cx. tarsalis*.

### California serogroup virus screening

To supplement our metatranscriptimic analysis, we carried out targeted screening for California serogroup viruses using a primer pair capable of detecting all viruses of the serogroup viruses (Kuno et al. 1996) and leveraging Sanger sequencing. In 2020, a total of 30 mosquito RNA pools representing 17,423 mosquitoes were screened, and in 2021 we screened 68 pools derived from 16,759 mosquitoes. Only two positive pools were identified, one per year and both from *Ae. vexans*. These pools contained mosquitoes captured in West Manitoba during CDC week 32 and 30 for 2020 and 2021, respectively. Due to an RNA integrity issue, we could only resolve the 2021 positive pool to the serogroup, which was unequivocally identified as Cache Valley virus based on a 251 bp fragment.

## Discussion

The primary objective of our study was to characterize the microbiomes of eight commonly found mosquito species in the Canadian Prairies. More than 35,000 individuals were collected in southern Manitoba over a three-year period (2019 to 2021) and subjected to metatranscriptomic analysis. This approach has been harnessed to catalogue the microorganisms harboured by mosquitoes from other geographical regions (Shi et al. 2017; Sadeghi et al. 2018; Batson et al. 2021; Ortiz-Baez et al. 2022; Li et al. 2023); however, to our knowledge this is the first study done in Canada. A distinct advantage of RNA sequencing over 16S rRNA or shotgun metagenomics is the capability to detect RNA viruses (Wensel et al. 2022). Mosquitoes harbour a diverse range RNA viruses, many of which can detrimentally affect human health (Shi et al. 2017; Batson et al. 2021). With the exception of *Ae. earlei*, each mosquitoes species we examined is known to transmit microorganisms of public health concern. Viruses dominated the microbial signature, and included representatives of five types of viral genomes from 19 different families. Similarly, Batson et al. (2021) showed the microbiome of mosquitoes from the state of California was overwhelmingly composed of viruses

In terms of the virome, several clear trends emerged from our study. As expected, there were strong associations between the number of mosquitoes collected/sequenced and virus discovery. *Aedes vexans* and *Cx. tarsalis* comprised 80% of the specimens and in turn they harboured the greatest number of viruses. The greater overall sequencing depth and number of libraries and collection sites for both of these species may also contribute (i.e., less false negatives). Nonetheless these are the most ubiquitous species within the sampling region (Chen et al. 2013; Brust and Ellis 1976; Baril et al. 2023a), and thus it is conceivable that they naturally harbour the greatest viral diversity. There were some notable exceptions; for instance, *Ae. canadensis* was infected with the same number of novel viruses as *Cx. tarsalis*. This may be attributed to lack of microbiome-related research conducted on this species, as it harboured no previously reported viruses. Another noteworthy pattern was the majority of viruses were unique to a given species, with no observable correlations between phylogenetic distance (i.e. same genera) and viral diversity. This is consistent with the literature, as Batson and colleagues (2021) also reported heterogeneity of the virome between species of the same genus. This specificity between virus and host was not discernable at higher taxon ranks; however, as the common viral families (e.g., Iflaviridae, Rhabdoviridae, Negevirus, Parvoviridae) were represented across several mosquito species.

An aspect of our study of considerable interest was the possible detection of medically important pathogenic microorganisms. West Nile virus, the causal agent of WNV encephalitis, is the primary mosquito-bourne pathogen endemic to our sampling region (Chen et al. 2013). The province of Manitoba has undertaken active surveillance of this virus since 2013, which includes the collection and molecular-based detection of WNV in pools of the regional vector, *Cx. tarsalis*. We did not identify WNV in our mosquito collections, which may be due to relatively low natural circulation of the virus during our sampling years. The province recorded a total of 120 infected *Cx. tarsalis* pools and six human cases between 2019 and 2021, which is well below historical maximums (Manitoba Health, 2023). Clinical cases of neuroinvasive disease caused by CSGVs have also been reported in Manitoba (Lau et al. 2017; Vosoughi et al. 2018) and positive mosquito pools have been identified in nearby regions (Anderson et al. 2015). While we did not detect any of these bunyaviruses through RNA sequencing, our more targeted RT-PCR approach identified two positive pools, confirming the presence of Cache Valley virus. This virus was first isolated in 1956 in Cache Valley, Utah, and is considered endemic throughout Canada (Zeller et al. 2000). While NGS is highly sensitive, it is likely that the sequencing depth coupled with the minimum coverage threshold requirements of our study was not sufficient to detect low frequency pathogens. We also detected Chuvirus, specifically Chuvirus Mos8Chu0, in 90% of *Ae. vexans* libraries and to a lesser extent in *Oc. dorsalis* and *Cx. tarsalis*. Chuvirus Mos8Chu0 was previously reported in *Culiseta minnisotae* from USA (GenBank accession: API61887.1), and the family has been associated with febrile illness in China (Quan et al., 2020). If this virus does induce disease, it could be of concern given its broad host range (i.e., detected in three mosquito genera) and ubiquitousness (found in all three sampling years). No other pathogens of known human or veterinary importance were identified in our study.

Of interest was the detection of Flanders hapavirus (FLAV) in >90% of *Cx. tarsalis* libraries. The virus has no known pathology but its transmission cycle shares the same avian hosts and *Culex* spp. vectors as WNV (Poh et al. 2018). Both viruses were shown to co-circulate, with FLAV detectable in *Culex* pools 1-3 weeks prior to peak WNV transmission (Poh et al. 2018). This suggests that FLAV could act as an early warning system for periods of high WNV transmission. Another bird virus was identified in *Cq. perturbans*, *Grus japonensis* parvovirus, though little is known about its transmission cycle (Wang et al. 2019). Additionally, we detected viruses that are not naturally vectored by mosquitoes, likely occurring due to horizontal transmission through nectar foraging behaviours as evidenced by the diverse plant transcripts found in our dataset. Sugar feeding is an important source of nutrients for both sexes, with females ingesting floral and extrafloral nectar throughout their adult life (Barredo and DeGennaro 2020; Foster 2022). We identified a pathogenic honeybee virus, Black queen cell virus, in *Ae. vexans*, which was recently reported (Baril et al. 2023b). We speculated that the virus was indirectly acquired by mosquitoes foraging at the same nectar sources as honeybees harbouring the virus. Three soybean thrip viruses (Thekke-Veetil et al. 2020) were also identified across multiple mosquito species. One of the viruses, Hubei macula-like virus 3, was previously detected in two mosquito genera, with the coauthors speculating their involvement in a horizontal transmission cycle between arthropods and plants (Colmant et al. 2022). Soybean is ubiquitously cultivated in Manitoba and represents an abundant nectar source, with each plant producing 200 to 800 flowers and yielding 0.5 µL of nectar per flower (van Schaik and Probst 1958; Erickson 1984). To this end, the legume is a preferred nectar source of mosquitoes in the Canadian Prairies (Cassone et al. *unpublished*).

The vast majority of the remaining viruses identified in our study are likely ISVs; however, it should be emphasized that little to no research has been done on these viruses to assess their pathogenicity or host tropism. Although ISVs infect diverse arthropods, the majority identified thus far have been isolated from mosquitoes (Öhlund et al. 2019). Despite their host-range restriction (i.e., only replicate in arthropods), many of the viruses we identified have been found on multiple continents, suggesting they (and perhaps most ISVs) encompass a cosmopolitan distribution. More than half were previously reported in North America, primarily in California where mosquito metatranscriptomic studies have been concentrated (Chandler et al. 2015; Sadeghi et al. 2018; Batson et al., 2021). To our knowledge, 17 of the mosquito-bourne viruses we detected are newly described in North America and largely infect different species in the same genera. These viruses were primarily discovered in *Aedes* and/or *Ochlerotatus* species from Finland (Suvanto et al. 2020; Nguyen et al. 2022), Australia (Shi et al. 2017; Ramírez et al. 2020; Ortiz-Baez et al. 2022), and Central Europe (Reuter et al. 2016), but also through meta-analysis (Parry et al. 2021) and unpublished GenBank deposits. Future studies are needed to determine if these viruses are truly ISVs (i.e., do not infect vertebrates) and their potential application as biomarkers, in biocontrol, and/or disrupting mosquito vectorial capacity.

In addition to known viruses, we identified sequences of viral origin that were not previously reported. The International Committee on Taxonomy of Viruses (ICTV) sets specific standards for virus discovery, with pairwise sequence similarity a primary criterion used. However, many of the viruses we detected are unclassified beyond order or family, making it challenging and somewhat arbitrary to determine a minimum identity threshold to define a new virus. Our amino acid sequence similarity cut-off threshold of 85% for all representative contigs is considered conservative (Kalantar et al. 2020), suggesting that we may have underestimated the number of new virus species in our dataset. To this end, a limitation for virus discovery from metagenomic or metatranscriptomic analysis is we currently lack bioinformatic tools that can accurately detect viruses exhibiting minimal to no sequence similarity (Cobbin et al. 2021). Nonetheless, we identified a total of seventeen novel viruses from eleven families, represented by one or multiple contig sequences. In some cases, our assembly algorithms generated a single contig that appears to encompass a nearly complete viral genome. For instance, Manitoba mononega-like virus 1 has one contig of 11.18 Kb and negeviruses typically have genome sizes between 9 and 10 kb (Vasilakis et al. 2013). Similarly, Manitoba tymo-like virus 1 had a contig of 6.3 Kb (typical genome size of Tymoviridae is 6-6.7 Kb; ICTV 2012) and Manitoba dicistro-like virus 1 had a contig of 7.15 Kb (typical genome size of Dicistroviridae is 8-10 Kb; ICTV 2012). As metatranscriptomic studies become increasingly more commonplace, it will be interesting to define the geographical distribution of these viruses, and to determine whether they have pathogenicity or potential applications in research.

In addition to viruses, the mosquito microbiome was made up of various fungi, bacteria, protozoa, and invertebrate parasites. Of interest were *P. gallinaceum* and *P. relictum*, which are causal agents of avian influenza. *Plasmodium* parasites that cause avian malaria have been well documented in Manitoba bird populations, infecting birds at rates of upwards of 50% and is present in both migratory and non-migratory birds (Enslow 2017; Enslow et al. 2020). Given the well-established ornithophilic blood-feeding preferences of *Cx. tarsalis*, this provides a suitable explanation for its mode of transmission from migratory to non-migratory bird. Avian malaria can be detrimental to bird populations that have not yet been exposed to it, which places birds in captivity (e.g., in zoos) and birds in northern regions at elevated risk (Grilo et al. 2016). We also recovered sequences of *D. immitis* in *Ae. vexans*. This roundworm is known to be present in the Manitoba area, and is the causative agent of heartworm disease in domestic dogs, cats, and in rare cases humans (Hendrix et al. 1980). There were also a variety of transcripts belonging to entomopathogenic fungi (e.g., *Coelomomyces stegomyia*) that could have future application in mosquito control.

In conclusion, our work builds on the current body of literature characterizing the microbiomes of mosquito species. Advances in metatranscriptomic analysis have allowed for unparalleled resolution into the suite of microorganisms harboured by these haematophagous pests. These studies have taken place on a global scale, though the vast majority of data collected in North America is from the west coast of USA. Cataloguing the microorganisms infecting mosquitoes provides the baseline information needed for more targeted studies aimed at elucidating how the microbiome influences development, longevity, immunity, and vector competence. We demonstrated that the virome is rich in diversity and represents the largest component of the microbiome, consisting of pathogens, ISVs, and even non-mosquito-bourne viruses. We report on several new viruses, and as metatranscriptomics becomes more pervasive, a nearly exhaustive list of novel viruses should emerge. Future studies should explore their human and veterinary implications, interactions with other arboviruses, temporal relationships, and rates of coinfection.

## Acknowledgements

We thank Ben Pilling, Jessica Sparrow, Milah Mikkelsen, and Carlyn Duncan for assistance with mosquito collections. The authors also thank Manitoba Public Health for use of trapping equipment and City of Winnipeg Insect Control Branch for mosquito collections. We are grateful to Dr. John Anderson and Angela Bransfield for providing positive CSGV samples.

## Funding

This project was funded through a grant from the Public Health Agency of Canada (PHAC) Infectious Disease and Climate Change Fund (IDCCF), awarded to Bryan Cassone.

## Conflict of interest

The authors declare no conflict of interest.

## Data Availability

The raw sequence reads can be retrieved from the NCBI short sequence read archive under the SRA accession number PRJNA793247. The contig sequences for each novel virus has been deposited in the GenBank database (accession numbers OR066146-OR066155).

## Author Contributions

BJC and CB conceived and designed the research project; CB conducted the field work and the laboratory experiments; CB analyzed what data and both authors interpreted the data; BJC and CB wrote the manuscript.

## Notes

### Competing Interest Statement

The authors have declared no competing interest.

