## Supplementary Table S1 for "Metatranscriptomic analysis of common mosquito vector species in the Canadian Prairies"

**Supplementary Table S1.** The year, location, species of mosquito and number of specimens comprising each mosquito pool that was sent for RNA Sequencing. A total of 2 pools, 19 pools and 23 RNA pools were sequenced from mosquitoes caught in 2019, 2020 and 2021, respectively. 1 pool (*Oc. triseriatus*) was comprised of specimens caught in Manitoba in 2019 and 2020.

| **Year** | **Location (s)** | **Species** | **Number of Mosquitoes** | **Year** | **Location (s)** | **Species** | **Number of Mosquitoes** |
| --- | --- | --- | --- | --- | --- | --- | --- |
| 2020 | Brandon | *Ae. vexans* | 1766 | 2021 | Shoal Lake | *An. Earlei* | 184 |
| 2020 | Shoal Lake | *Ae. vexans* | 1126 | 2021 | Shoal Lake | *Ae. vexans* | 1154 |
| 2020 | Virden & Souris | *Ae. vexans* | 1788 | 2021 | Shoal Lake | *Cx. Tarsalis* | 450 |
| 2020 | Boissevain | *Ae. vexans* | 1144 | 2020 | Shoal Lake | *Oc. flavescens* | 270 |
| 2020 | Killarney | *Ae. vexans* | 653 | 2021 | Souris | *Ae. vexans* | 992 |
| 2020 | Carberry & Cypress River | *Ae. vexans* | 550 | 2021 | Souris | *Cx. tarsalis* | 126 |
| 2020 | Cypress River | *Cq. perturbans* | 2246 | 2021 | Virden | *Ae. vexans* | 907 |
| 2020 | Eastern MB | *Ae. vexans* | 2208 | 2021 | Virden | *Cq. Perturbans* | 1230 |
| 2020 | Western MB | *Cq. perturbans* | 427 | 2021 | Brandon | *Ae. vexans* | 1544 |
| 2020 | Brandon | *Cx. tarsalis* | 897 | 2021 | Brandon | *Cx. tarsalis* | 547 |
| 2020 | Shoal Lake, Virden, Souris | *Cx. tarsalis* | 476 | 2021 | Brandon | *Ae. canadensis* | 24 |
| 2020 | Carberry & Cypress River | *Cx. tarsalis* | 476 | 2019-2021 | Various | *Oc. triseriatus* | 199 |
| 2020 | Brandon | *Oc. dorsalis* | 509 | 2021 | Boissevain | *Ae. vexans* | 781 |
| 2020 | Shoal Lake | *Oc. dorsalis* | 681 | 2021 | Boissevain | *Cx. tarsalis* | 376 |
| 2020 | Souris & Virden | *Oc. dorsalis* | 525 | 2021 | Killarney | *Ae. vexans* | 250 |
| 2020 | Boissevain & Killarney | *Oc. dorsalis* | 1107 | 2021 | Shoal Lake | *Ae. vexans* | 283 |
| 2020 | Eastern MB | *Cx. tarsalis* | 400 | 2021 | Souris | *Ae. vexans* | 900 |
| 2020 | Eastern MB | *Cq. perturbans* | 444 | 2021 | Cypress River | *Cx. tarsalis* | 222 |
| 2019 | Brandon | *Cx. tarsalis* | 2089 | 2021 | Cypress River | *Cq. Perturbans* | 1000 |
| 2019 | Brandon | *Ae. vexans* | 1783 | 2021 | Brandon | *Ae. vexans* | 1213 |
| 2021 | Newdale | *Ae. vexans* | 443 | 2021 | Cypress River | *Cq. perturbans* | 1000 |
| 2021 | Carberry | *Ae. vexans* | 289 | 2020 | Shoal Lake | *Ae. vexans* | 50 |
| 2021 | Carberry | *Cx. tarsalis* | 137 |  |  |  |  |
