## Supplementary Table S2 for "Metatranscriptomic analysis of common mosquito vector species in the Canadian Prairies"

**Supplementary Table S2.** Sequencing statistics for the 40 previously reported viruses detected in our mosquito pools.

| **Virus** | **Contigs** | **Longest Contig (nt)** | **Reads** | **Coverage Depth** | | | **aa Percent Identity** | | | |
| --- | --- | --- | --- | --- | --- | --- | --- | --- | --- | --- |
|  |  |  |  | **Mean** | **Min** | **Max** | **Mean** | **Min** | **Max** | **Median** |
| **+ssRNA - Dicistroviridae** | | | | | | | | | | |
| Black queen cell virus | 2 | 6035 | 1,613 | 17 | 11.15 | 22.85 | 100 | 100 | 100 | 100 |
| Soybean thrips dicistrovirus | 11 | 9121 | 11,192 | 17.32 | 10.11 | 29.01 | 100 | 100 | 100 | 100 |
| **+ssRNA - Flaviviridae** | | | | | | | | | | |
| Inari jingmenvirus | 1 | 1267 | 139 | 11.02 | 11.02 | 11.02 | 100 | 100 | 100 | 100 |
| Placeda virus | 169 | 11737 | 106,682 | 90.37 | 10.02 | 1,114.82 | 97.87 | 86.24 | 100 | 98.55 |
| **+ssRNA - Iflaviridae** | | | | | | | | | | |
| Cafluga virus | 2 | 3426 | 976 | 20.82 | 19.41 | 22.24 | 99.77 | 99.55 | 100 | 99.77 |
| Culex Iflavi-like virus 4 | 23 | 9811 | 13,815 | 22.81 | 10.65 | 58.9 | 99.81 | 96.2 | 100 | 100 |
| Culex iflavilike virus 3 | 183 | 1824 | 266,451 | 284.65 | 18.65 | 1,269.06 | 99.49 | 94.19 | 100 | 100 |
| Hanko iflavirus 1 | 30 | 9246 | 2,938,402 | 3,394.01 | 10.99 | 19,795.35 | 97.49 | 93.45 | 100 | 97.3 |
| Hanko iflavirus 2 | 16 | 9206 | 19,456 | 93.72 | 18.5 | 208.48 | 97.05 | 89.58 | 100 | 100 |
| Hubei arthropod virus 1 | 1 | 2967 | 352 | 11.95 | 11.95 | 11.95 | 100 | 100 | 100 | 100 |
| Pedersore iflavirus | 25 | 9899 | 39,865 | 28.41 | 10.35 | 266.71 | 93.47 | 89.14 | 100 | 93.38 |
| Soybean thrips iflavirus 4 | 1 | 3835 | 636 | 16.6 | 16.6 | 16.6 | 100 | 100 | 100 | 100 |
| Thrace picorna-like virus 1 | 2 | 1172 | 2,010 | 82.45 | 34.1 | 130.79 | 91.62 | 89.58 | 93.65 | 91.62 |
| Yongsan picorna-like virus 1 | 64 | 4085 | 55,324 | 62.58 | 10.27 | 180.88 | 89.4 | 85.05 | 100 | 89.11 |
| Yongsan picorna-like virus 2 | 4 | 9573 | 58,132 | 165.26 | 34.98 | 535.91 | 100 | 100 | 100 | 100 |
| **+ssRNA - Luteoviridae** | | | | | | | | | | |
| Marma virus | 23 | 3160 | 39,344 | 70.95 | 11.62 | 129.64 | 100 | 100 | 100 | 100 |
| **+ssRNA - Narnaviridae** | | | | | | | | | | |
| Culex narnavirus 1 | 1 | 513 | 63 | 12.07 | 12.07 | 12.07 | 100 | 100 | 100 | 100 |
| **+ssRNA - Negevirus** | | | | | | | | | | |
| Big Cypress virus | 3 | 9570 | 6,079 | 37.12 | 18.89 | 65.64 | 98.77 | 96.3 | 100 | 100 |
| Bro virus | 1 | 11375 | 3,826 | 33.85 | 33.85 | 33.85 | 90 | 90 | 90 | 90 |
| Cordoba virus | 14 | 5133 | 135,594 | 548.17 | 17.51 | 2,225.85 | 97.1 | 93.75 | 100 | 97.23 |
| Mekrijarvi Negevirus | 6 | 9873 | 167,549 | 294.01 | 30.28 | 1,262.63 | 94.13 | 90.78 | 100 | 93.4 |
| Utsjoki negevirus 3 | 2 | 989 | 416 | 21.12 | 18 | 24.24 | 99.02 | 99.01 | 99.03 | 99.02 |
| **+ssRNA - Nodaviridae** | | | | | | | | | | |
| Hubei noda-like virus 12 | 2 | 4762 | 4,244 | 54.47 | 23.6 | 85.34 | 95.83 | 91.67 | 100 | 95.83 |
| **+ssRNA - Tombusviridae** | | | | | | | | | | |
| Des Moines River virus | 4 | 2361 | 35,117 | 486.06 | 34.94 | 897.17 | 100 | 100 | 100 | 100 |
| Hubei mosquito virus 4 | 6 | 5103 | 2,919 | 16.68 | 10.02 | 39.59 | 96.5 | 92.65 | 100 | 95.7 |
| Tiger mosquito bi-segmented tombus-like virus | 1 | 2351 | 13,049 | 557.41 | 557.41 | 557.41 | 100 | 100 | 100 | 100 |
| **+ssRNA - Tymoviridae** | | | | | | | | | | |
| Hubei macula-like virus 3 | 8 | 6059 | 530,385 | 1,092.52 | 11.15 | 7,749.86 | 96.13 | 92.15 | 100 | 96.63 |
| **+ssRNA - Virgaviridae** | | | | | | | | | | |
| Hubei virga-like virus 2 | 6 | 947 | 505 | 13.26 | 10.94 | 15.5 | 99.04 | 96.27 | 100 | 100 |
| **-ssRNA - Chuviridae** | | | | | | | | | | |
| Chuvirus | 39 | 6799 | 22,130 | 28 | 10.17 | 214.52 | 97.49 | 89.66 | 100 | 97.02 |
| **-ssRNA - Orthomyxoviridae** | | | | | | | | | | |
| Astopletus virus | 29 | 1044 | 5,975 | 32.63 | 10.91 | 107.95 | 99.06 | 95.77 | 100 | 100 |
| Wuhan mosquito virus 6 | 101 | 2467 | 133,667 | 176.35 | 12.58 | 641.58 | 99.76 | 97.69 | 100 | 100 |
| **-ssRNA - Peribunyaviridae** | | | | | | | | | | |
| Culex bunyavirus 2 | 19 | 1896 | 6,255 | 23.92 | 10.28 | 49.84 | 100 | 100 | 100 | 100 |
| **-ssRNA - Rhabdoviridae** | | | | | | | | | | |
| Canya virus | 9 | 2444 | 2,613 | 29.27 | 11.64 | 66.33 | 90.87 | 85.05 | 100 | 85.92 |
| Culex Rhabdo-like virus | 4 | 1973 | 807 | 14.68 | 10.87 | 23.28 | 100 | 100 | 100 | 100 |
| Culex rhabdovirus | 2 | 429 | 997 | 116.26 | 74.47 | 158.06 | 100 | 100 | 100 | 100 |
| Elisy virus | 21 | 1817 | 3,849 | 19.29 | 10.1 | 40.69 | 99.27 | 96.41 | 100 | 100 |
| Flanders hapavirus | 81 | 8773 | 104,235 | 113.05 | 10 | 686.87 | 99.9 | 95.96 | 100 | 100 |
| Manitoba virus | 5 | 1589 | 1,003 | 19.71 | 15.74 | 24.8 | 99.76 | 98.8 | 100 | 100 |
| Merida virus | 55 | 7483 | 76,697 | 91.6 | 11.06 | 276.87 | 99.86 | 97.14 | 100 | 100 |
| Riverside virus 1 | 17 | 8033 | 18,623 | 45.08 | 10.8 | 120.41 | 90.2 | 85.32 | 93.01 | 90 |
| **dsRNA - Birnaviridae** | | | | | | | | | | |
| Ballard Lake virus | 48 | 3472 | 682,541 | 431.81 | 10.84 | 2,260.59 | 99.86 | 93.27 | 100 | 100 |
| **dsRNA - Partitiviridae** | | | | | | | | | | |
| Partitivirus-like Culex mosquito virus | 12 | 1751 | 4,477 | 23.28 | 11.49 | 45.83 | 100 | 100 | 100 | 100 |
| **dsRNA - Totiviridae** | | | | | | | | | | |
| Gouley virus | 3 | 368 | 117 | 12.22 | 10.42 | 14.89 | 95.04 | 92.86 | 97.67 | 94.59 |
| Hattula totivirus 1 | 1 | 4909 | 702 | 14.41 | 14.41 | 14.41 | 89.07 | 89.07 | 89.07 | 89.07 |
| Snelk virus | 4 | 295 | 496 | 41.15 | 23.15 | 67.61 | 100 | 100 | 100 | 100 |
| **ssDNA - Parvoviridae** | | | | | | | | | | |
| Aedes albopictus densovirus | 4 | 3355 | 14,051 | 139.82 | 94.27 | 203.18 | 97.46 | 89.86 | 100 | 100 |
| Aedes vexans densovirus isolate | 1 | 3367 | 594 | 17.69 | 17.69 | 17.69 | 100 | 100 | 100 | 100 |
| Culex densovirus | 8 | 1697 | 6,035 | 127.22 | 11.65 | 490.03 | 99.56 | 97.71 | 100 | 100 |
| Grus japonensis parvoviridae | 1 | 570 | 366 | 60.82 | 60.82 | 60.82 | 100 | 100 | 100 | 100 |
