## Supplementary Table S3 for "Metatranscriptomic analysis of common mosquito vector species in the Canadian Prairies"

**Supplementary Table S3**. Complete list of contig sequences generated assembled from novel (i.e., not previously reported) viruses harboured by Canadian Prairie mosquitoes.

>Manitoba_dicistro-like_virus_1

TTTTTTGCTATGAAAAGGAATCTGCCAGTGAAGCTGACAGGTTCGACCCCATCCTTTACAATTTGGGTCCAAAATCAAATTGCTAAATTTTGAAAATTTTAAAATTTCGGGAGTGGTTAAGCCACATACTTTAAAAGTACTTCACGTGCCATTCGGACACTAATGCAGGGAAACCCGACAAACCTGCATTCACGTGACTCAAAACGGTCGTAATTCCAAACACTAACTCCAGTTGATAGCCTTTCGGCATCTGAAGCACCTAATTAAAGGCATGTTTTTACCCTAACAGCACCTCATAACCTTTCGGTGTATGCTGCACTCAATGAAGAGCAGGTTCAAGCCCTAAAAACACTGAATAGCCTCTCGGCATATGTTTCACTCAATTAAGAGCAGGCTTGTGGAATTATTCTGTTTTGGTTTACCATACATCTATGACCTTGTTATTGAACTTAAGTCGAGTTTGTATGGTTCTTGGGCTGTAGGAACGGGACTTTCACCGCATATTCGCAACGTTAATTGTTCTGGCAAACAGTCCGGAACTCCTAAAAGGAATCCAAGATTGTAGTCGTCACCCGCTGCTTTAGCAATAAATACATCACCGCCCCCATCACGAGACAACAATCCAGTAGTTTGAGGTTGAGTAAAATGAAAATATAAATCTGGTTCTTGAGTTGAAAATTCTTTATAAAATGAATTTATGAATGTGTAAACTGGTGAATAAAAAGGAAATTGAAATTCTGCTCCTCCTTTAACGACGATAGCGTCCATATCACACAATCCAGTAGATCCGCAGTTTTTAGCGCTGGCTATGGTTGTGAAATTTGCAATCTGTGAAGCAGCATGTTCGGGTATGGCTCTGGCTGTTATTAAGTTGGAACTTGAGTCCCAAGCCTTGTAACGAAAGCCGCCTGTCCTAAACGCATACAACGCAGAAATAGAATCCACAAGAGCACATTGCTGTACCTGGTATGGTAAACCTGTTGTGAAGGTATAATACTGAATGTATGGAACGGATGGTAGAGAGATCAACTTGGAAGAAGTACGTATTACCCAATTGAAACGCTTAATGAGGGCGCGCAATGAATAAACTTCCTCTCCCGTAGTGTAATCTGGTATCTTATCATCTGTAGAAGTTGGAGAGAGTCCTGTTATTGATTTTATTTCTTTAGTTGAATTTTGCATACTCGATCTGGTTTTCATGATACCATCTGCCACAAAAGCCTTTGCATTGAACTGGGCTTCTGCTTCAATTATGGTAGGTTCTTGGGGTGTTATTTCGGGAGCATCATTAGCAATCGGAAAATATGACCGTCCGCCCATTGGATAGGCAAATTGAAAATTGTCCGCTGCACAAAACTCCGTTATCATATTCACTGAGTTTGAAACCACGGCTGATGAGGCCTGTAGCTCAGTATCAAGAAACACTGCCACATATCCACAGTAAGTTGAAAGTAATTCCAACTGTTCACCGGGTACATTAGGCGAGACTTGTGTGTCAAAAGGTACATTCTTCCAGGGAGTTGATGAAATAAAAGGTAGTCTTACATAAAAATCTGTTTTCTCTCTGAAATCAATTACTACTGAATAACAATATTCTGAATTGGCATAAGTTATGTTTGCTGGATTCGTGGCCATCGGGTCGTAAACGACCTTCAAACGCAACGAATGGTAATCTGTTTTAATGAGTCCTATGTGAAGCACTGTGTCTCCACGCCAGTATTTAAAATTTGAGCCAATATATGTCTGTAAATTTGGTTGTCTCACTTGGATTGTTCCTCCTGTTGATGTGTTCAAAACTAATTGTGTGTCAAGGTCAAATTTAAATGGTGATACTTGGTAGGACGCTAGCATTGTGCCAACGTTAGTTGATGTTGTTATTGTATGTGTGTCGTAATATTGTAGTGTTTGCATAAGGTATGTTAGGGAGAGTTCATCTGTATCGGATCCTGCGAATCCGGGTAGCATCTTGATCTTGTTATCTCCATGCAACGCTAACATATGGCCATTATCGATTCCTGTGATTGTTGCGAAATTCGAAAAGGCATGCAAACAACGTGGTGCAATTTTGTCGAGGTTGGTGGGTTTGCCTAGCCCAAAGCTAGCATAAATATTGAGTCCAGCCATTGCCGCATTATTGACTGCGTCTACAATTTTGGAAATTGCTGGTATATTTTTGCCTACTCCGTGAAGAATGCCATAAGCTGTTTGACGAATATCGTGACCGATATTGCTTATGGGACCATCGCGTTCTTCTTGCATGGTCACCTTGGTCATTCCTTCAATGCCTAGCTGCGCCTCCGCCTGAACAATCACTGGCCGCAACGTTTCTAACGCTGTAACCAGTGAAGACTGCGTAGGAGCACCCAACCGGACATTGTCATAATATGCTCTGACTGCTATGTTCACAGTCTGAGTTGAGGCTGACGACAGTGGATCATAGACCCATAAGTCGAAGATGCCCCAATCTCCTCCACCGGTCGTGAGATCGTAATGGGTAAAAGAGGATATATATGGCAATGGTATGTTTACTTCAGTCTGTTTTGAGATGTCAATGATTACACTTGGTAGTGATGTTGAACGACACAAGGAGCGTCGTGTTTGTTGAAACCTGTCTACTAAATATTTTGGGACTGGAGTATATGTACAGATCAACCTTCCCGCTTGGAAAGGCTGACTGTTAAACTGTAGCTTCAAAACAATGTCTGCGCGAAACAGACCAAAGCCCTTAAGCTTTTCTCTATACATTGTTGTTAATGTATCTCCAGGTATACTCAGTGAGAGCAAGTTATGGTGTTTAGCATCACTGGTGGACCACGTCACTGCAGCGATCTTTTCTGGCCTTTTCAGGAACGAAATTACGCTGTGTTCTCGCGGTTCATCACCATGTTTAATCAATGATATAGGTAAATTTGTAATTTCTGTGACACTTTCCGTTACAACTCTAGCATCTTCAGTGAAATTAGTAATTTCACGTCTCTCTTCAAAAGAGTTCTCAGAAGTTGTTCCTAAGATTCCTTGTTGCTGATTTGTATTGGAGGTGTCTGTTGTGTTGTCTGTAATTTGGTGTATTGAAGCAAGTCTGGTATTTAAAGGATACTGCTGACTTAGGCGAGTACCTCGACTGACGTTTGCCTGGGTAATAAAGGGCTGCTTCGAGAGACCCTGGCAGGTAACACTAAATAGTGCCTCTCGTTCTTCAGATAGCAATACTAGTCTTTTCTTTTGCGTCAGAGATTTGTATAGATTAGCAAGATCACATCTAAAGCTTCTGGAAGTAGTTATACTCGTCTTCCTCATCACGAGTTTTGTTGAAAACACTGTGCTTCCGCAAATGTTTCAACAGTGTCAAAATCTTGTTCAAGTTGGTGAATATATTCGATCCGCTCGGGGGAAATTTCAGGCACCCAATTAAGGGCATCTTTGCAGACTAACCTGATTTTCTTCGTCCACAGCTTGAATGTTTCTTCATCGTGAAGAGATAGCTCTTGAATAGCCCACTCCACGTTATCTTGGGTTGTCAATTCTTCTTGTATGTTTTTCTTAAACCAAAATGGTGTCTCAAGCACTGTGTCTAGAGTTAGCGGTGCAACATAGCGATTGAGTTTTGGTTCCCACCGAAAGCCTCGTTTTAAAAACGTTACTTGTTCAATGGGTCTATAGTCTGGCATTTGTTTATCTGATTTCTGTTCATCTGTGTATGTCATTCCTATTGTTTCAAAAAGTTTTGGCAGGGTATTTTGGTTAAAAATTGGAAGTAATTCCTTGATTAACGCTACTACATGGTCATCTCCATAGGAGATCATCTTGAATCTTTTGCATAACTTAGCTATTTCTAGATAAGTTTTCAAATTTGCACTTTTGCAAAGTAGATAGCGAAAAACAAGATTTACATAAAGTGAATTTACTATGGCTGTTAAATAATGTCCACTTGGTAGTGAGTGAGTCCAGGAATATAGCTGATCTTTTGCTAAATGGGTTGAGTTCACTAAATCACTGAAAAGAACATTTCTGATTTTCGTGTCTTCGGGCGTATAGTCTTCTAACAATTCTCCAAGGGCATTCAAAATGTCTCTTGCTTTCTCAAGAATCTGGCTTAACAATGAGGCATCAAAGTTAGCAAAGTCTCCGGCGCACATGTAATTACTGCATTGCTGCAGATGTTTTACACATTCGTCCCATTCTCGAGAATATACGTTTATTCCGACACAAATTCCTGTCTGAGTTCTGTGACGAGTTAGTAAACCAACTATTCCTTGGTAATATTGTTTACAGGCGATATAATAATCTTGGGAGCATGCTGAGAAAACTCTGGTCTTAAACCACTTTTCTTTAAGTTTTCTCTCATCCTTAAGAGTATCCACGAATACATGTGGGTATCTTTTATTTTGTCTTGCCAAATCAAATATCTTTTGACATCTTTCTTTAACTGGCTTAGCAAGCTCCAAATCAAATTCATCGCCATCTCCAAACCACTTTTTCTTTCCTGGTTTGCCCTCTTTGCTCCATCCATATCCAGGAGAGGTTCCTCTGTTAATCGAATTAACATATGGATCTCCATCGATTCCTTTTACTGCTTCTTCAAACGAATATTTGCTTTTATAAGTAGGATACTTATGTTTTTGGCAAACATCGATAAGTTCTTGTTTGTAGGCTTCTGCTACCTTGTTGAGTAAACTCTGGTCAAGGCAAGCTGCAGGTAATCCACACTTTTCTAGTCTTGCAATAAAAGGGTCATAGCGAGTTCCTTCCTCTTGGGGAATTCCATCTTCTTTTCTCACTGTGACAGGATGCAACAAAGTTGGACGAGAAACACTTTCTCTCCAAGCTTGAGTATTATCTGGGTGTAATTTTGATTTTTGAATTTTTGATTGAGTTGGTTGTCTAAATGGTTCTGGTGAAATTCCAATCGGGTGAAATTCTCCCTTAAAAGGAAGACTTTGTGTTTCTGCAACAACTGAGTGTTCTGGTGGCTCAATTGTATCAACTTCATCAAATAGTTTTAAAAGATCAACTAAATCTTCTTGATAGAGAGCAACTGAATATCCTTCATCCTTTGATGGTATTCCCATGATATGCATACCCACTAATTTGGCCTTAGCCATAGGATGATTAACTGCTAAGATTGCTCCGCAGTCTCCTTTTGTAGTGGATATAGCATATCTCCAAAAATCTCTTGTATAGAAATGCTCAGTATCGTCTTCAAACTCTACTTCTGAATGAACGCAAGCGCTATGGTCGTGCGACCTGTACATAGCATTGGCGTTACATGCAGAGTATTCTGTGGCAATTTCTTTACCATTCTCTCTTAAATGACCCAAGATTACTCCTACTTTGTCAATTTTATGTTGTTCTTTTCTTGTCATAAAATGTTTGGTTCTGTCAGGGTGATTGTCTGCTGTGATGAATTTAAAAGCAACACAGTCATGCACATATGTTTCATCGGGAACGTCTTCATCATTGTAGACATTGATTGGATCAACATCTGATACAAGCATTTCATAACTAGTATGAAGTCCTCTGATTTTTATCTTACAATTTTCATTCTCGTTTCCCATTGCAACAAGTGGTACAAGAAAGTGTCGGGGCATAATAAAAACTTTTCCGGCCAAAAAGAAACCATTACCAATAAATTTGTCATCTATCGAAACTGTATACATTCCTCTTCTGGCAAAAGAGGCCATCACGTCACGGGAAGCCTGATCGTAAGCTTCTGCTGCTGCTTTCACAACTTTCTTTAAAACTCGATTGTGAGTTTTCAATTTCCCGTTGTACGCGGACTGCCGTACTACATGTGGTTTAGGAGTATACGCTGATTCATTTTGGCTTCGTTTAGGGAAGATTCCCTTAAACAAAGATCCAATCTTGTATATCGAAACTGCCACACCCAGTATCCCAAGGGCTTTAAGGACTGTAGAATGTTCCGCTAGCTTTGCCTTTAGTCTATCTTTGATGACTGTCATTTTCTCTAAATAAACGGGCAATGCTTCCTTTAAGGCTCTAAAATCTGCCAGTTCCCAATACTTGATAGGGAACATTGATGTTTGTCGCTCTGGTAGTTGGTGTTGTTGAAGTGTCAATGGTTGGGGCAAGAAAGAAACAGGTGTTTCACTAACTGGTTGGAAAAATGAAGCCGCAGTTTGGACAATCGATGAGGGTGTCCACATTTGAGATTCTGCTGGGACGAATCCCAACTCTTTTGCGAGAGTAGAATCAAATTTCTGTCTAGATGTTTCAAAACTAAAACGTGCATTATATTTTTGAAAAGCATCTTGGACAACTGCATCAATATCAATTGCTTCGCCTATTTTCTTACTTGAGCTGATGTCAAAAGGAATGAAAGTTTGTCTGAAAGGATCATAATCATGTTGATATTGCTCATCCAAAAAGTAATGAAGTTTTCCGTCAATAGTTACAGGAGTTGTAAACTCTCGTTTAATCTGGATTTCATATGACTTATCAATTCTTCTATAAAAAGCTTCTGGGGCAAAAAGAGATTCAATTTTGGGTTTCTTTGCATTTGTTGTGAGAATGACTATACGAGACTGAAAAGTTGTTGTGTTTTTATCAGCCAAGTCAGCCATGTGCAAAGCATAAGGAAAAGTATTTCCAGCTCGAATCATTTCAAAGAGTTCAGGGTTAGGTTCCTGAGCGAAATCACGATGTTGTGCAAAATCGTCAAATACACAGACGAGTTGTCCGTGATACCCATCCCAAAACTCTTGTTCCACGTTTCGTGCATAGATCAAATCTGAAATATTAATTTTAGTTTTATCAATCTTTTCTTTATCACAAATTTTGGCTAGCAATTTTGTTGCTAGTGTAGTAACTATTGATGATTTTCCTTGTCCTGTTTGTCCATAAAGTTGAATTATGATTGGGGCCAGTCTCTGTCCCTTTGATGATACTCCTCTGAGTCCCAATCTTTCATACAACTTTCTAAGCAATCCATACTGAACTGAAATAGCCTGTCTATATTCTCCTAAACCTGGGGTCTGTAGAATTTTAAGGCCAAGAAGTTCAGCTTCCATGACATGAGCTTGTAGTACCGAAGAATTTTGAATTTGATTCTTTTTATCACCTTCTGCTATTAGTCTCATCAATGTATTAAATCTTTCTATCTGCGGGTGTTCTGCAGATTCCTCCAAT

>Manitoba_iridescent_virus_1

AATTAATACAAGTATTTAAATGTATTTTTTAACCATAACTCTATTATTTCTTTGTCTTAGATTCTTTTTTATTCTTTGCTTCTTTGACTTCTTTTTCTTTTTTTGTTGTTGATTTTTTAACCGATTTAGGCTCTTCAATAACCTTAGTTTCTTTCTCCTTAGTTTCTTTTGGTTTTCGACCACGTTTAATTTCTTTTTCTTCTGGAATAGTTGATGGTTCAACAACTTCGTCAAAACAATTTTTTAAATATTTTTGCATCGTAGGATATTTAAGGACCGTATCATCATCAATTTGTAGTAATTTTTTCAACTTTGCATCAGGAGTAATTATGGTTCTATTGTCTGGGTCACGAAGTTCGTTTTCCTTGATATATGTACATAATATGTTGGTGATATCATACCGAGACTTTTCTGTTTCACCATATATCCAACCCTTACACTCTGGCAAGTCACAGAGACTTGCAAACATAGCCATATTTTCAGAAATCTTACGTAACTTTTCTAAACCTGTATTGTGAGACTCTCTGGTTCTCTTGGGCTTGTATTGAATATTTTTATACTTATATAAAAATTTCCTTGCTTCTTTGACTTCACGCAAGACATTACGTAATTGTTTTCGCATTTCAATAACGGTTGGTTCTGATAATTTAATAATATTATCTATGGATACAACCGATCCTTCTATCTTTTCCAAAGATTTTTCGAAATGCTCGAAAGCAATCTCTTTATTTTTACGTCCCTTTTTGGGGACAATGGTTAGTTCAACTTGTTTTGGGGCAACTGAAAACATATTCTTT

>Manitoba_mononega-like_virus_1

AGGAGGAAAAAGCAGAAGCTGAGCGACAAGCAGACCTCGAGAAGAAAGAGAAGGAGAGGAAAGAGCTGGAAGAGAAGAAAGCTAGAGAGATTGAGGCACAGAGAGAGAAAGAAGCCGCTGAAGCAAAGAGGATGAAGGAGCCTGTCTTATTTGAGGATCTCGAAGATCTTAAGAAGGCAGCTGTTAAGCAAAACCGTAATCTTGATCTTCTCGTTCAGCAAGTTGATCTACTCAGCAAGCTTGTAGCTCTGGAAAAGGAGGATACAACCAGCGAGAAGCCACGTGATTGTGATTACATGGAAGGTATGGCGCCTCAGTCTACTTCTCTATTTAAGAAGATAAATGCCGGTTTCTCTAAATTCTCGTGCAGCTCACTCACTACACCTCAGTGTGTACTTCGGATCTTGGAGCAGGATTATCCGTTTGATCTTGTACGGCTTGACTCAATCGAGGATGCTAAAGACTTCATTAATGAGGTTGAATCAAAATGGAACACCTCTGCCTTACGGGAGATCACACACAGTAATAACTTCGATCGGCAAGTCAGTGATATCCCAAGAATAAACATGGTAGATAGTGTTGAAAATAAGCTGAAAAAAGTAATTGAGAATCACGAAACTCTGACTGCCTTGAATCGGAATGTTATAAATTTGTCCTCCCAGATAAATAGAATTCACAACATGAATGCCTCCAGAATTGTCCCTCTGGCAGCAGGTGCATTTGGTAAATCTCCAGATCCCGTGGAGAATGAAGATTTTCTACTCGGCTATTCTACCCTCGGTTCACAGGTGACCGATATGGCAGAGTGCTCCGTCTCTTTTAAGTCAGGAGCGAATAAGGAACGTGTAGGATCAAAATCGGAACGGGAACAGCTCGGACAAGCACTAACTCAGAGCCTTCAAGCCGCGCATGCAGCGAAGCAGCTCCCAGCACTACACACCCTCACAGAAGAGGAGGAGGTTGCGCTTGCTATCAAGCAAAGCCTAGTTATGTCAGAACAGAGCAAAAATAGAGAACTAATAATATACGAAGACTGGGACAAATCACCTGCTGCTGCAGAAGAAGCCATAAAGAAGAGACAGGAGATAATACTGTCAGAAGCAGCCAAGAAACATGTTCCTACACCTGATTCCAAGAAACAGCCACCATCTCAACAGAGAGCCTTGACTGCCCGCGAGATACTATTACAAAGAGCTCGAGAGAGGTCCAAATCAAAAGACCAAAATTGACAAGGACATCTCTTAGATACTTCTTTAACCATAAGCAATAAAAAATTGTTATTCTCATTGATTAATAAAAAACCCGATAGTACAATTCACATAGATTAAGCCGTTTACTGGAGAATTTGAACAACACAGGAGTAAAACAAAATCAACCCCCTACAAAATAAAAATAAAGGAAAATAAAAATAAAATAAAACGATATTCAGTATACTCTTGTAATTGCTCGGAGGAGCAACGGAACAAAGATGTATAAATTGGTACTACTATTCGCCACCGTATTCTCACTCTATATCACCGTTAATGCCGCAATCATTGCGTATGATTGCAGAACTGAAGAGATCAACAAGACGGCTATCTCCCTCGTTCAATTGCCTGATTGTACTAGCCGGCGAAGCAAGCCCCTCATTCGAACTACACAAGTTGTAGTCTCTCAGACGGAGCAAAGAAGAAAGATCCCGATATCGCGGTGCTTGATTACTGCATCGCACCTAATTTATCGTTGCGGAAAATGGCTAGGGGCAGACACTCCGGTGGACACATACACACAAGTTATTCGTGTCGGTCGAGAAGAGTGCCAACGGCTGATCAAGGATGGAACTTTAGTTCTTCCAGGGACCATGGATCAAGTAATACAATTTTCCGGAACGGGACGATTCAGCCATAGCTACTGGAGTTGGGGCGGATTTAGCGGATCATCCTGCGATCCTGGACCAACTTTGACAGTGAACGGACGAACCTGGGAGCGTGCAGTTAGGTTAACCAATCTTGAGGTGACCTATACAGCAAGTCAAGCGAGCCTAGAGGTGGACACAGGGAGATTAATCTTTGCTAATGGAGAAAGATGTGAGTTCCAATCTGAACAATGTGATACCGCAGATTATGGAAATATTTATTGGCATAGACCTACTCCAATTTGTACGGCAGAGGTTCCTACTGTCACTGTGTACAAAGGACTAGGAGAACTGGCAACTATGATAGACAAGTCCAACAGAACTCTCCAATTCATCCATGTAAGTCATGGCGGGTATGATTTTCAGATCAAGTTGGAAGAGACAAACAGGGCATCGGTCTGCGGTTATCCATCAAGACATACAGAGCATCCAGATTTGTTTGTAACACTCCTAGATTCCTCGAGTCCTCCATTTCCTAATGTTCCAGAACTTGATGCAGATAATGTGAACCTGCTCAGTTATATCAACTCAAAACTGGTTTACTCTATGCGGCACACAAAAGAGCAGGTTGATACTCTTTATGACCTCTTCCATCAGGAGAGGTGCAACATGCAGAACCGAATCACACAAAACCTACAGACATTGGCCCTGCTCTCACCAAGGGAGTTCGCTTATCAATACTTCGGACGACCCGGGTTTACCGCTGTAGCTCGAGGCGAAGCAATATATGCAGCACAGTGCAAGGCTGTTGCTGTAACTCCCCGACCACTGGATCTGGGAACCTGTTACAATGAACTGGTAGTAACCTATAACAACCAAACATGGTTTATGACTCCTAGAACAAGGATACTAATTGCGGCAGGCACCATAATCCCATGTTCTGCAGAGTTCTCTCCAATCTATCTCCTAGGTAACAAATGGGTTAGTCAAAACAGTTTGGGTTTAACAGAGGTACCTAAGCCAACAGTTATTACCACGGAGGACATCGAATATGAATTTGAGACAATGGCCAATCTTGCAAATGGAGGGTTGTATACACCTGAAGTACTGCGGGAATACCAAAGAATCCTTACTTCACCCATGGAGGAGTCAGTTATAGTGGCTAGAATGACAGATTCATTGAGAGGCACATCACAACTGCCGAATGGGTACAGTTTTTCAAGGGGCCTTACGGAAGAGGATCTGACCTTCATCGGTGACAAGGTAGGCTGGACGAGTGTACTTGGATCTTGGATTATGAATGCTGGCATAGGTTTTTCATCTTTCCTATTTCTAGTGTTTGTTGTCAATAATATAGCTAGTGTTGTTAATTGCTGCATTCATTTCCGTTATGCGAAGAGGGAAAAAGGGTTGAAGACTGCTATTCTTGCCGGACTATTTAGCTCAATCGGCAATTTGGTACTATCAGGGAGGTGGAAGAATCGTGAGATGTCTGAAACCAATCAAGAGAGAAAATCACTCCAAACTAATACGAGAGGGTTTGTTTGGAGGCCATCTGTACCTTTGGAAGACGATTGTGAAATGGAAGAACAAGTTTGACAAGATAAATATACATTAATTTTGACTGCAACCCTACCAAGTGCCATTCTTTATTATTTAATGTGTGGTTTTTGGAGCCTATTCTCCAGAGATTAATGTTTAAGATACAGTAAACAGCGAACAATGTTGCTACATTGAAGAGTAATAACTATAAATAAATTTTAGAAGTGCCCAACTATTTTATAAAAACCCGATAGTAACTTTCACATCACATCAGCTAGTATTTATTATTGTGGAGGAATGTTGACTAACATAGTTTGTCCAAGATGTGGAGCTCTAGTTCCAGTAGAATTGAAGACAACAGAGGATATAAGAACAACAAGCTGTTCTTGCCCATTCTGTGGGAGATTGTTCTACTTGAGTAATGTTGACTTGCAAGTGTAATTCAGGGACTACTTAAACACCCAGCTCGAAAGCAATCGTGGATCCATAGAGCTTTAGAAAAACCCGCTAGTAAGATTCACACAATTGCGGACAATGGCACTGAGCATTGACATCGTTGTCTGCCTTCCCCCAAAAGAAGAAGAGTTCAAAGGGAATCCGGGCAGAGCTTTTACAAAAACCATATATCCGTCATCAGGTCCCTCCAATCTTGGGTGATGAGAGACAATTGAGAATTTCCATAGATTTGCTTCATGTCACTTCTCAGGCACTTATAAAAACCCGATAGTACAGATCACTTAAGTTAAGCAATAACATTGGTGTGAGAGAAAACAATTCTAACAAAGAGTCAACAAAGGAAACCAATGGATTGGGCAACAATATTTTCCGCACTTGAGGAAGAAGGGGAGGAGTCGGAGGAGGTGCAGAGGGATAGTGAGTGGGATAAGTACATGAAAAAACGGGAACGATTGACATTCATCGGAGGTCACTTGTGCGCACCAATTGTAGAAGAAGACTGCAAATTCTTGCTGACAACACCATTGGAGACAATGAAAGGATGCAAGAGACATACCGTCCTGAAGAATCGGCTTACAAGTACGTTTCCCAACTGCAAGCAGATTGCCGACAAAGTAAAAGATCCGATCGGCTTTATCCAAGATTGGACAAAACAGTTGGATATTGATGACACTGGTTCGGATTTCCGAACTTGCCTAACTGCTATGGACAATTTTGTACTCCACAAAATGAATTACTTGTTGCCGTTCTGTAATTCTCAGGAAGAAATTGACTTCTGCCTCAAAAAGACTAGAGATATGACAGCAATGTGTCAGAAAGCCGTCCAAGGATTAGCAAGACAATACCACGATATGAGGAGAGCAGAAACAGTAAAGGATTCGATTGGTTTCACTGAACTCGAAGACGGTAAGATGATCCGATACCGAGATCTGTGGGCTGAACAGGATGTTTGTGTCTTGCGTTCAGCTTATTCGGGAGATTTATTATTCTCAACAAATTTATTGCTGTGCTCTTTGGACAAGGTTCAGAGTCGATTTGCTCTGGAAGCCTATTGGCTGATTAGCGACACGATGGAGAAGTATCCAAACAAATCCATATACAAATCTGGAATTGCCCTGAACAAACTCTTTGAAGGTTTCCGTGCGACTATGGGTCAAGACTTCTTTACTCTAATGGCAGGGTACGAATCATTGATTGTTGGTTGGACAGTCGCAGCCAGACATGATTTGGGTTGCAAAGACCTGTTCCGCTCAGAGGCAGAATCTTCACTCAAATTGCTTGAAAAATATGGTCTCGAAAAACAGTTTTATCAAATGCTTCCTCCAGATCGTGACTTCAATACCGTTTTGATGCACTTGGAGTTGACGGGGATAGTTAAGATCTTTGGATACCCGACACTCGAGGCAGATAGAATGCTTGACCAGATATTTGAATATGGAACAAAGAACAAATTTGATATTGACACAAATCTCTTGAGGAATGTGGAAGGTGTTATGAGACGAGAGTTCTGCTTAAATTATTATAAAAAGCACACAAGGTACCCTAAATTGCTTTACTACCCCTCAGCACTGAGAAACCTAGGGAAACAAAAACCAATGCAAGAGTTTACTCACAAACTATATGAGGAATGGTCCACTGTCATCTTTTCAAAAACTATCGAATTCGATTATGTACCTGATCAAGCCGACATGGTAAAAGATTCGGCTTCAGCTGTTAATAGGAAGGCATGGGCTACAATGTATGACGGATGTGCATTCAGGAAAAGGTATGGCAAGCCTCCTCCAAGACGAGTGAAGCGCAAATACTCCGCAAGAACGATAGAGGCATATCTTGAAGGAGGTGAGGATGAGATTCGAAAGCTGATAACTGCTCAAGACAATGGACACTTTGACTATGAAGATTGGATTTGTGTTGAGTGTCAGAAAGAATGCGAACTGAAAGGAGTTTGTGGAAGAGCGTTCACCAAACAAACTCCTCCGAGACGTCTTATCCAAGTTACAATGGAGCACAATATTGCTGAAAGTATTTTTCCCTATGTGCCTGAACAAAGTATGACAGATTCGGAGATAACTAATTCAAGGAAAATATTAACACAGGTGAAGAGCATGCAACGATCATCTCATTTTATGAGTCTAGATCTTAAAAAATGGTGTTTGAACTGGAGACATGAAGTGGTTGAGATGACCGGAAGAATGTACGATGAGCTCTTTGGACTAAAAACATTATATAAGAACAGCCACCTGTTCTTTGTCGGATGTCTTGTTCTGTCGAACAATCGATTGACTCCACCTGATTATGACCAAGCCGGTGAACCTATTGAGGGACCAGTATGCTGTAGAAGTTTCATTGGAGCTATGGAAGGCTTGGCTCAAAAGTTCTGGACCCACCCAACCGGGGCACTGATTCGATTCGTAATGGAATCTTGCTCCCTCAATGGGGATATAATGGGGCAGGGTGACAATCAAATTATAGTCCTACACTTTTCAAAAGATGACGAAAAAGCGAACGACAAGAGAGAAGCTTTCTTGAAAGCTTTGTCTGCTGCTTTCCAGGGGATTAATCATGAGCTCAAGCGACAAGAGACATGGTACTCTCAACATCTTCATGAGTACAGCAAAACGAGGCTCTACCAAGGTGTCAGCGTGAGCTACGGAACAAAGAAGGCTTCAAAGTTAATACCAGATTCCAACGATGGTCTGTTTTCATCATCATCGTGCATGTCCACTATAAACACAATGACTGAGGGTATTGCCAGGGCAGACTACAAGGCCGATGCTGCATTCATGCTTAACCAGTTCTGTCAAACAAGCTTCATGTTGAGGAAAAATATTATCACTCGGGATTTCTCGAAGGATACAATTCAAGGTTTATTAAATTGGCCATTAGACTTTGGTGGTCTACCTATTAGTTCTTATCTCGAGCACACCATCAGGGGAAATGACGATAAGGTGAGTCAGTGGTGGAGTATTCTTAATACAGCGAAGCAGTTAGGTTTTCCTGTTGTCAAAGAGATGGAGCGACTGTGGGAAATCAAGCCTCCAACACCTGCATCTAGCGCACTGGAGAGATCTCGGCTGTACGAGGATGTCTTCTGTCTCAAGGTCAAAGTGCCTCCTTCGTGTGAAGCAAGAATGAAAGATCTTATTGCTAATTACCTCAGAAATCCTAAGTATGTGAAGAACCCTGTTATCCACAAGCTGTACAATACAGATTGTTCGATGAGCTATGACTCCTTAATCCAAGCTATTGACTTGATTCGTCCCAATTTCTTGCCAGCATCCCACGAGCTTCTAGCTTCTAGCAATGCAGGAAGACTTCTCCAGCTGCAAGGAAAACTTACATCATCGAAAACCCTGCAGAAGATTGTTCAACTACATGAGAGAGTCTCCTTAATAGACCTAATAAAAAAGAAGAATGATGAGTTCCGCGTGTATGTTCAAGAACGAGTTCGACGACCCGGCTCCATGTTCAGGAACAGGATAGGAGAATGCCCTAGTAAGATTGCTCGACAGCTTCGGTCAGAGAGTTGGGGTATTGAGATGATTGATATTACAAAAGCACCATTATCGCACCAGATAAAAGTCAAACCGGTAGATCTTTGTACACCAGAAGAACTCGAGTCAGGAATATCAGTTCGTGTGTCCGGTGAGATGGAACAGAGTCCGAGGCAATGCTACTTAACCTATGGGCCTGCAAGGAATTACATAGGATCACGAACTAAATCTAAACTTAAGAAATCTACAATCAACATAGTAGAAAAGTCGGGATTTGTAAAGAACCTTCAGAAAATAGGTCTAATTAGGTCGTGGTTCACGAAGATCGGGGACCAAAATATGGTTGATCTTTGCACAATGCTTATGGAGGAGAAAAGATCACTCATTACCGACATGCCTGAAGGCTTGGATGACTTGGGAGAACTATGCTCAACGGTTACATCAGGCAATATACTCCACCGCTTCAAAAGCAGCGTCGAGCACGAGACAGCTATGCTTAACTGTTTGCCTTCGATATCAGGACACTTTGAGTACAGCTCCAACTCAATGAAAAGACTCACCGCTGGGGGCGTTGATCTTGAGATATTTTATCAATACATCATGGTGGGTGTTACAACAGGACTAGCTATGTACACAAATTTGACCGGGGTATGTCACGGATCTTATATGGTCCTTTTTCCTTGTTCTGAGTGCACGCGAGTGATTCCTGATATCACAATCGGTATGGATCAACTACCTGCTATGTTTAAGCCTATGAATGTTCCTAATGATGTAAAACCAGTCAGTTTGTCTACTGACTTGTCAGCGGAGGAGGCAAGATTTCTGATGGCTGTATCTGTTGGCTACAATATGGCTCGAAATTTTGATGAGAACTTTAGAGTTCATCATACACAAGGGAATAGCAGCCTCACTGCGATGGATTACAAAAAGGATGTAGTATCCCTTAATGACCTCCGACTTTTAGATCAAAAAGTCGTTCTTGATGTTGCTTTGGCTACATCTCAACATGGCAACAGGCTCTATTGGTCAAGAGACAGCCTCCTGGTAGCTCACTCGGAAGATAGATCTTTCATGTACTTTGCGGAATGTATACTCGAAGCAGGATTGGTACCTCGATTGTTGGATATAGTCGGTACGAGAGTTAATGATCATTCAGCAGTAACGACAGCACATGGACTAGCCAGCTTTCTGTCGAGACATGCCGGGCAATACATGGAACGTAATGGACAAACAATAATCGACAACAGTACAAAATTCATTTTTCAAGATTCATCTCCTTATTTAGTGGATAACATTCTTAGATTTTGCTGTTTATGGAAGTCATCCCACAAGAAGATAACTCCGGCACAGAAGGGTGTAGTATTGAATGTATTAGCAACCACAAAGAACCTCGGTTTAACTGCTAAATTACTTCATGTCCCTCTGCAAGTTGTAAAGGTTGATATGAGCGACATGATCACGTTATGGAGAACAATGAATCATGAGTTTTCCAACGATCTATCGTTTACACCGATGCAATTATCTGTCCCTGAGCTCCCTTTAGACCAGTACGACTACTTTAATGTTGGCAGACTTCATTCTTGCAGGGTTCTGAAGAGGGTTGGAGAGATAGGATACACAATTCCGCAGCTCTCTTTCTTTGCCAGGAGTATTGGTGTGATCTCTACAGCTGCCAGCAAGTTTATCGAAGTTCTGCTCATTCTTGATTTAGTGGATTATTTGAAAGAGAGCACAGGTGTGATTTACAGCTTGGCAGAGGGATCAGGAGGGACATTGATAACTCTGCTGTCTCTGTTTCCGCAGCTGAGAGGGCTATACAATACGTGGATGAGATCAGACATTGCCAACAGGGATCGTGTAAACGATTACAGTGTGCCGGCTGCGACTGCACTTAAGATGAGCAATGATAGGCTTATATCTCCTAATTCGCTTATAACAGGCGAGACAAACATTATCACTCCTACTTTTCTCAGGAAGTTGGATGAATGTCTTGAGGAACACCCTCCTGTATTGGTGACTATTGACGCTGAATCACCAGAAGGTTCTAGAGAGGGGGGTTCAAACATTCATTTCCTACAAACAACTGTGCTAACTATCTTAAGGAAACAGTCCACATTGATCATACTCAAGATGTTTCTTCAAAATGAGACGAGAGCAATAATAAGTGACATTCTTGAACCATACCACACGGAGATCCTGTGGTATATTTGCAAGCCTCTGAGCAGTAATCCAGTCGGAAAGGAGGTCTACCTCGTGATCATTCCGAATAACATATCAAGACGAAAGAACGAGCAAGTTCTGCGGGATATGGAGCACAGCATTCTTCAATATGTTCCTCTAGAGACATCAATGACGAAGAGCATGATCAATGATTACGTGATACATTCCCGTCGGTTCTCCAATATGATGATGAACATGTGCCCTCCAGAACTTTTGATTCGCCACAAATATACTGAGTTTCTTCCTGCTATGGGCTGTAGCCTGTACTGTCCCAAATTTTTTGATAATCTTCTCGAGATCTTTGACAAGGTTCATGCAACCAATAGCGACTTTAGTGTCCATCTTGCTATCAGGATGAAGGGAACAAATGGAACCCTATTTTCTCTCGCCCATGAGATAATTTTCTTAATGCTATGGCATTCTAAGGGAAGTGACAAAAGTTTCGGCACCATACTTGCAAGCATGACACAATTAGAGCTACCTGCGGACTTAGCGCATGTCCGATCAAATCTGACTAGAGTAAGTCCATTCAAACACATCAGCGTGTCAGACCTCGGGTTGTGGAAAACATGGACAGATGCGAGAATGTACCTGAGTGAACAAGCACAGGGTAACTGGGGGTGTGACTGTTCCCGACCTATAAGGTACTATGTTAATACCACTGGAAGAACTGTCGACCGCAAGAAATTGTTAAGCACATATATATGGAGTGGTTTGAAACTAAGAGGGATGGTACACCAGGATACTCTCGAAGAGATGAGATTGACACCGGAAAGAAGAATGTTAGGTCAGGTGGACGAGAATGCTATATCTGTCCTGATCGAAAGAGTCACCCTAGAGAAGGTGGATCAACAACCTCTCGAAGGTTTTGATCAGTATGATTGGGTAAATTAGCTTATTAATATAGAAACGCATGAGATGATCCATAAAATTTTATAAAAACCCGAGTCTCGGGTTTTTATTAATATTATTTTGTTTTATTTTATTTTATTTTATTTTATTTTTCAATTTTCTTAATTATTATTAACTTACTGTCATTTAGTTGTTATATATTTAACTTGAATTTTTAATTTTTAATTTTTAATTTGGATTGTGTTTTATCGTTTTTTGCCAACTCACTGTTGTTGTATCTGGCAAATGGAAAGTTAAGAGGAAACTCCATTTGCTGAGCTCCTAAGGAGACACCAGCGCATGGTTCTAACCTAACGTAAAGCTCGATATTTATTTAACTTTATCCCTGTCTGAGATGCTCTGTTCTGTTTATGTTTGGTCAATCCACTTCTTTCCCTCCATCCTCATCCCTCTGTCTAACCGATTCACGGTTGGCGTGTTTGAGTAGGCTACTCTTATCTGATTGTTTATTCATTTTCGTTTTCACAAAAAGCAAAATGCTTCTCTTATTTCTTTATTTTATTTTACTCTATTCTATTTTATTTTCTGTTGATTGCTTTCACTCTGTCTGCCTGTTCATTTATTTATTCATTCATTCAGTAGTCTATATTAAATTGCCCTTAAAATTTTAGTTTTTTTGAGGTTGGTTTTTATTCAATGGACTGGGCGTACAGGCGGGCTGGTCGCACAACAAAGTCTGGTGTATGTTCCATGAAAGAATAGTTTTCTTTGTTTCTTTGTTT

>Manitoba_mononega-like_virus_2

TTTTTTTATTAAATATTAAATATAGACTGTTGACATTATGTTCCAACGGTAGATTTAGCATTTAATAAATAAGTAAATAAACAGACAGCCTGAAAATCCATAAATAATAAATGTGTTTAAAGTATGAGCAGTTGCTTTGCTTTGAGCATATGAGTCTAATTTTCTTAACGACTATAATTTGGAGGGTATCAAAATATAGAGTCTGAAAGAAATGTTAAATCCTCACTTCATTATCATTCACCATACATGGGATATTTGAATTACTATATATTTATAGCACCCCGGGCATTTGAGAGGTAGAACCACAGGGTAGTCAGAGGTGTCAACTGTCACCTTTGATGTGTGCCCGCAGTTTTGACATACTATAATGTTCTCCATTTTCTGATTTTTGACTATCGGGTTTTACTAAAGTTAAAAATGGTACTTAAATTCAAGCATTAAGAAAGAAGAAGAGAAGAATAAGGAGTCGACTCTAACTAATAGAGAGATGCCCAATCAAACGACATCGTCAATTTCTCCGATCTCTCTCATAAGAATGTTGCTCTCTGCATTTGCATGTTTATCCTTGTTATTTGTGTTACTATTTCTATTTTTCAATAGTTGTCCATTTAGAATGAAGTGGGTAATTGTGTCGAAGATGCAGAAAAGAAGCGCAATCAAGCACCCATGAGACTGTTTTAAGAATTTGAAATTGACCAGGCTGTTAATCAAACTTATGAACAGCTTGAGGACAAATACCAAAAATAAAATAAATCCAAACCAACCTCCTATTTCCTTCGATGCTGTACTCACTGCCTTAGACCAGGATGATATTTTGTTTTCGATCATCTGATAGTCATTTGTTGAGAATGCATTTCCCAAGGAAGACCCTTCGGGAAGAGAGTATCGCCCAGCAATTGCATCAGTTATACGTGACGATATTATCTTTTCTTCAAGAGGAGATGTCAATATTTTCTGATACTTCAGTATTGTCTCTGACGTGTAAAGGCCATTCGATGCTAAATCCTCAAGCTGCTCAAACTCATAGTTCAATGGCTCCGGGGTAATAACAAGAGGGCTTTTTACAGTCATCAAACCCATAGACGTCGGAGTTACCCATCTATTATTCAGAAGAAATTGAGGACCTAGATCGGCTGCGCATTCCACAATTGAGCCAACTGCAACCAGTATTCTAGTTCTAGGCATCATATACCAGGTTGCATTATTATACATAACAGCAAGTTCATTGTAACATGTCTGATCAACCTTAATAGGAATCACTGCGACTTCACGGCATTTTGCTACATGAACGACTTCTCCTCTTACAACTGCTGTATACCCAGGCGACTGGAAATATTGGTACGCAAATTCCTTGGGTGAGACCAATGCAAGTGTCATGAGGTTCTCTGTGATGCGATTTTGAAGTTTGCAACGCTCTCTTTCAAATACTCGGAAAAGTCTATCTACTTCTTGTTTAGTATGTCTCATTGCGTATATAAATTTAGAATTTATGTAGTTCAACAAGTTGACATCTTCTGATCCAACCTCCTTCGTTAAAGGAAACACGGGAGAGGATTGAGGTATAATGGTGACAAATAGTCTCGGATGTTCCGTCGAGTAAGAAGAGAAACCGCAGATAGATGCCCCATACTTGGATAGCTTGATTTGGAAATCATAATGACCTTGGACCACGTGAACAAAAAGATTTGCAGGAGAGCCTGTTGGAACATCATTGTCTGTTATGAGTACTCCATTTCCTTTGAATACTAAAGAGTTCTTGGAATTTAATGAGTCGCAGCCAGGGCTGGGATTTTTCCAAAATACTTGACCATACCCAGAATGATCACAACGCTCATCCTCGATCTTACATGATATTCCGTTGGGAAACTTGACTACACCTTCGTCATGGAAAAGTCTTGCCGTCCCTGTCGTATATGTGATATCTAGTCTTGTAGATCGTATGGCGGAGCCCCACGATATTCCTGAAGGGGATACAAGAGTCCCTCCCGACGAGCATTCTCCTCCGTCAATTGATCCAAAGCTTATATACGAAAATGACTTTGTTCCAGTATCTGGTAGAGTTAATCGAATGTTCTTTCCCCATCGAAGATTGATGAATTTCCCCTTGATCATATCATCACACTCTTCTCTAGTTACGTGATAGATTTCAGAGTAGAAACCATCCTTATGCCAGGTATCGATCATTTTTCCACATCTTTGAACTATATGAAACGCCTCTATTGAGCATCGTGTGAAGGGAATTTCTGAGATGGTTGTGGTTTGAGTTACTGCTAGAGTGATAGGCACTGTTGTTATATTGGGTTTACGTTCAAGGCAGGTAGGTGTTTCGACAAGGGAAATGGTTGTTTTGTTTATAGATACACCTCTGCAGTCATAGGCAACAAGGCTGTAAGTTATGTCCAACGACATAAAAGCTACTAACGCGAAAATAAAGTTTTTGCCAATCATCTTTACGACTTTGATTTTTGACTAATCGG

>Manitoba_mononega-like_virus_3

CCCTATCTTAGATGCTGAAAGCACCTTGAGTCAGGTTTTTGACTATGGAATGCAAGATGACCTTGATATCGACATGAACTTAATCGCTGACTTAGAGGGCTTAGCTAGACGTGACCTCTGTGTGAACCATTACCGGAAATATAGGAAATTGCCTAACATATCTGAATGCCCGCCAGAGCTAGACCATCTTTTAAGAAATAAGCTACCCCCCGTGAACATGAACAAGAGATATGACTTGTGGAGACAGGTCAAGTTTACAAAGACTCTTGAATATGACTACAGTCCTGATCAATCCGATTTAATTAAGGATTCAGCTGGTGCAATCAACTCCGAGGCATGGGCACAAATGTTTGACCCGTGTGCCTTTCGTGAGGACTACGGGACAGAACCACCACCAATGGAAGAGAAGATCTACAGCAATCGGATTATCTTAGAGTACTTAAGCTCCACACCAGAGAAGGTTAAACAGCTAATCACAGAAAGGGACAACGGAATACTATCTCCGCAAGACCACATCTGCGTTCTCTGTCAAAAGGAATGTGAGCTGAATATAAAAGGGAGATCATTTACAAAACAAACTCCTCCACAAAGGTTAATTCAGGTCTCTTTAGAACATAACATAGCTAAGTTCATCTTTCCTTTAGTCCCTGAGCAATCCATGATAGATAGCGAGATAAAAAATGTGAGAAGGCATTTACAACAAGTGAAGAATATGCAGAGCAGCTCACAGTTCATCAATCTGGACCTGAAGAAGTGGTGTCTCCATCAAAGACATGCATCAAATGAGTACTTGGGTCGAATATACGATGAACTATTCGGAATGAAAACTTTATTTCAAACTAGTCATCTATTCTTCAATGGATGTCTCATCCTCCCCAATAACCGGCTGATTCCTCCCATGTATAATAGTCGTGGTGGTCCAACAGAAGGGAATCGTTGTACTAGGAGATTTATTGGAGGGATGGAAGGGATGCAGCAGAAAAAGTGGACTCATGGTGCTATAGTATTAATTAAGTATGTCATGGAAAGTTGCGGGTTAAAAGGAGAAGTGATGGGACAGGGAGATAATCAAGTAATCCTATTGCACTTTGCAAAGGGAGACAACAGTGCTGATGAGAAGAGAGATGCTTTCATTACAAGTTTGGAGCTTCACTTTAGAAGAATCGGTCATAGGTTAAAGAAAATGGAGACATGGTACTCGAAATACTTGCATGAATACTCAAAACAAAGAATGTATATGGGATCTGCAGTGAGTTATTCCCTGAAGAAAAGCACAAAAGTTATTCCTGATATCAATGATGGGTTGTTTTCTATTCCTTCGTCTTTATCTACAATAAACACAATAACTGAAGGAATAGCAAGGGCTGATTATGATCCTGATGCAGCTTACATTATGAATTGTGTCAACATAACAAATTACCTGTGCAGAAAAGGTATTATTAATAAGGATTCAGACAAATTGACCTGTCTACTATTCTTGAACTGGCCAATAGATTTTGGAGGAATCAATATAAGCAGTTACCACTCTCACGCGGTCAGAGGGAATGACGACAAAGTGTCCATTTGGCTATCTCTACTAGATACCATAAAAGAGTTTAAGCCTGACCTATATGAGCTGTGTCTATCCAAATGGATCCTTAAACCGAACACACAACCAGAGGATGCCATTGATAGGAAGAATCTCTATGAAGATATCTACAGTTTGAATGTCCCACCACACCCCTCTGCAGAGTCTGACATAAAGGATATGGTCGTAAGTTTCTTGAAAGGACCACTAGTAACAAACCCTGCAATTACGCGGTTGTACGATGAGAACCATTCAGCTAGTTACGAGGAAGTAATCCGGGATGTAGACCAGATCAGACCTGTTTTCCCTCCGTTGTCTCATGAGCTCCTGAAGAACAGCAATGCAGGACTTTTTATTCGACTCCAGGGCAAGCTTACACATTCAAAGACAATAGAAAAGATAGTGAATGAAAATGTGACTAAATCACTCATTGATCTTATAAGAGAGAAGAACGAGGACATGGTGCAATCCATCAACCGTAGACTCAGGAATGATAGAACATGGGAGAATAGAGAGCTGATCACCCAATTGCAGGAAGATTATTGTCCCACACAGCTAGCAAACATTCTAAGAAATGAGAGCTGGAATGTCGACATAATAGGCGTCACAAAGCCCAGTTGGTCTCATCAGTGCATAGTTAAGGATATGGATGACTGTTCAGAGGAGGAGAAGCGAAGAGGAATTACTGTGAGGGTATCAGGAGAAGCCTTGGATAATCCGCGGTTGATGCATAAAACTTATGGTCCTATGAGGCCTTTTGTAGGATCAACAACAAAAGAGAAGATCAATAAAAAAGCTACCCTTGCATTCCAGGATACGTCAACTTTCTCATCTTCTGCAACCAAGCTTAGAAAGGTGAGATCATGGTTGACTCGTCTTGCTGATGAAAAGTGTGTTGATTTCTGTAATAAACTGATCCATGAGAAAAGAGACCTAATAGAGGTTGAGGAAGACATATGGGACGCTTTAGACGACGAACTTCAGACAACAGTAGCAGGGAATCTGTTCCATAGATTTAAGTGTAGTATTGATCGAGATACTGCTATGATAAACTGTTTGCCTTCTGTTACAGGGCAGGTCGAGTATAGTTCAAATTCTATGAAGGATATGACAGCTGGAGGGAAAGATTTCAATATTTTCTACCAATATCTATATACGTCATGCACTGAATCCTTAGCCATGATGGCTCGTACTATGAATAAGATTAGCCCCATGTATATGATACTGTTCCCGGATTGTGACTGTGTTAAAGAGACTCCTAATCCTCAACTGAGGATGTCAGGCAGTAGTGGCTATGATCCCCCCAGCTTGAGAGATCTAGGCATATTAGCTAGGGTAGCAGACCCTCGAGGTACAGAAAATTTAAGATCATATGCCTCGTTCATAATAGGCCGTGAGATTGCCAATAATATTGATGAAAATTTTAGATTATCTCACCAGCAAGGAGACATCACAGTAAGCTCCACTGAGTACAAAAAAGGATTGGTCTCAATAAATGACATGAGACTTGTTGATTTTGAGTCTCTGATTTGCAGTGCCATCTTGTTCTCTAGGCATTGCAGAAGACTGTATCTCGAGAAAAACAGGATGATGTTATGTTTGTCAGATGATCGCTCCTTCCTTTACCTAGCTGAGCTGATACTAGAGTCAGATCTTGTGCCTAAACTCTTAGAACTAGTTCAGCGGAGTACCAACGAGCACTCGGGAGTTTGTTCCATATATGGGATTTCATCATACATCTCTCGAAACATGCCGATCATCATGAGACAGGTGCTTAGCAGCAGCAGAAACACACAGCTCGTGAGGTTTGCAGAAGACGACTTCTCGCAATGGGTTAACTTGTTTAGATTTTTAGTGAAAGCTAATAAGCTATTCCCAGGGTTATTGCAGGGATCAATACACAATAAGCTAGATAGGAACCTCAAGAGAGAATGCAATTTATCGCGCTTAATACGAGATTCAGAATTGAACATCCAGTTCGAGAGATTGAGTCAGGGTGAAATACTGGCGGCATGGAAGTTGTCAGACCGAGGGGATGACATATCGGTCAGGTTCAGACCCCGGATTATAGGATGTAGGCCAATAACAGTAGATAGATATCCAATTGCAGAGGTCGGGCATTATCACTTCTTAAGATCCAGTGAGGATGCTGGTGATTTAAAAGTCCCACAGTTATCCTTCCTAGCTCGGCCTCTAGGATCAATTTCATCAGCAGCCAACAAAGCTCTTGAATGCCTCCTAGCCACGGGTCTCTTTTGGTTTCTACATGAAAGTGATGGATATTTCTACTTTACTGCTGAGGGATCCGGGAGTATAGTAGTGGGAGTGGGCACAGCTTGCCCATATTCGAAAATAATATACAACACATGGATGCGTCCTGATATCGCAAATAGAGACTTAGCCACAGATGCACAGGTGCCAGCATGGACAGATGCAAGGTTAGATCCAAGCAGGTTAGTAACACCAAACCCACTGGCCGTGGGTGAGACTGATATCGTGAGTGAAAGATTCCTTGGAAAAATGTTAACTATGATGGAAAAGTACCCACCTACATTCATCTCCATGGATGCAGAATCAGATGTAGAAACAAATAACCTCCATTTCTTAACCGATTTTATTATTCCTGTTGCTGAGCACTGCAAGGCTATTTTCTTGGTGAAGTTGTTCCATATGTCTCCTCTCCACGAGAAATTAACAAGGTATTTGGCGGGTTTGAATGATTATTCTTGGACGTTATACAAGCCAATATCTAGCAACCCCGTCGGTCCTGAGGTATATTTAATAATGTGCCCTAAGGAGCACGTGGGAAACTCTTTCAGGGACTGTCAAGCCATGCAGCCATCATTTGAAGGATACTTGGGGAATAACATGAACCTTTCTGAACAGTCATGTAATAGATTCGTCGATATTGCATTTAAGGTTAGCTCCTTGCTCAGAGATATATTTCCAACTGGCCAACTCGATATCCCACACAGGTTTCAGGAGTTGCAGCAGACAGGTGCTTTTTGCTCTGTTTATTGCAAGAAGTTTTTAAGCGTCTTTATGAAGACTATCGATGAGCTCCACACAAGTGAAAACGACACTCTATCTCACTTAGCCACCCGAGGGATAGGAATTAACGAGAAGTTGTATGATCTGTCACATAGCCTGATCTTTCTTTTCCTGTACCATGGCTATAATGACACCCTGTTTGGAAAGCTCCGGAATTTGTCTACTTGCAAAATAACACGGAATTATGAAGAAGCAAGAAAGAAAGGGGAAGGTTACCTTAGCAAGTGTCCACCAGGAGAAGGGGACCTATGGGGAACATGGAAAGATGCTAGGATGTTTATGAGAGAAATGGGTAAATTAGAGAACCCTTGCAGTTGCAAACCTGAAGCAGAAATGTGGACTTGGGATGTTCTAAATCATTTTTCACCCAGGCAAGCAATAAGCTGGCAACTTTTTTCATCTATCATTAAAGAGACTATTGCCCCAGGGATGAAGTGGGATTTATACTCAGACTTAAGGCACAAGCCAAAGAAGATTTCAGTCAATCTTCTAGATCACAAGATATGTAATCCTGTGAGCCAATCTGCGATAACGCCGGAGCATGTTCGAATGGAGATTTATTGATGTGCAAAGGGCATAAGTAGTTTGATGTTAAGCGGGTTAGTATAAAATAAATATTTAAACATAATATTACTTTTAAAGAATAAATTAATGGTCAATTAACAATTATAGAAGTAATTCTTTATTACTTTGGAACCATCAACCTTAGTGAGAATAATTTGTATCTTCTTCCCTTGTTCTTAAAAAACCCGCAGACAAGAAGAAAAGGTATGGGGGAACAAACGAACAACGGAGCTGTCGTAATGTGACAGAGTCATCTTTTTCGTTTTGTGTTGTT

>Manitoba_narnavirus_1

AGGTCGTCCTGAATATCACCGAAGATGAATCTTTTATTAAAGATCCATCTGGTGGGTGAGGGTGCAGACAGATATGCCTGGTACTCCTCATCCTCAGATATTGAGAGATCATAGCCCCCCCACTCCTTTGATAAAAGGATGTGGGAGAATGCTTTTGGGTTAAGTTTCTTCGAGGGGAGTAATGCTCCCATTCTATTAATAAATAGATCTCGGATGGAAACTTTATGACTGTGAGGCCACAATCGTGACTTCGGTAGCCATTTTAAAGCGCCGGCGAGTTGGGAAGATTTTCCAATCGCCACGTTTTTATTATCCTTTAGGAGGAGGGTCGATTGACCTTTCTCCATAAGACGAACTTTGATTGAGTCAACTATTAGGGAGTTTCCCTCATAATTGTCCCAATCAATGGCACAGGGTTTCTCGTTTTCCAGATTTGGAAGACAAAGAACCCTTTCGCAGTATGAAACAGCAAATCTGGATATACCATGTTTATCGATAGATATCATGGATCCTACACTTTTGTGTATTTCTGTTATTCTGCGTAAATAGGAGAGAGGACCCTTTACTAAGTGGTCGTCACCTCCTATGTGTATAAACCTCCAGTCGCTCTTTGGAGCGCTGTAATAACAGGTTCTGGGGTTTATTTTGTGGTGTTCTAAAAACGCAAGTTCTTCAATCACTAGATTGAGCAGCGTTAAAACTGGCTTGGTAATAGCCTCGCCCATCATCACACCTCTTTCTGATGTTACAAAAGTACCATCGGGGAGGGTAATGATCCTTGGTCCTATCATTGATAGTACCAGATCAGTATACTCACTAATTGGAAGGCCATAGCCCTCCATAAAGGCCCTAAGTAACTTAGGAGCCAATTGGTGAGATATTGTGTCTGAGGCGGCCGTTAGGTCGGAGCTTAGCACAAAGTGTTCTCTAAGACTTTTGTCTTTGAGTCTACACATATCCTGTGCTGCTTGCCAAGTTTGATCCTGTCGGTGAAAACTTGAGAAGCAACTAGGATGCCATTTTAACATATACTTTAGTGTATGTGCAAGTGGTGCCATTAGAATGTTCACCCAATAGGGTGTTATACTAATGATACGAGCTTTGTCACCCATTTCTGGGATGACCTCAGCCCTTGCCTCTATGACCTCGGAAGAATCCTTCCAAGCTACATAGAGGATTTGTTTTCCAAGGACTTCATCAAGCCCTTGGAGAGCACCAAGGCGATCCTTTGGTATTCCCGATAATCGGGGACCATCGAACTCGCCTTCAGTTATGACTTGGCCTCTAAAGAGGTATCTCCATAACTCTAAGCCACCTATATGATGAACTTCTCCAAAAGGAGTAGATTCAAAGTAAGACCTATCGGTCTTTACTAATAGTCGGGCTTTTACAGCTGCTAGACAGGCATTAGCCTGGCCGCCTTTGGCGACTGAACTAGTTAGCTCACCTGAGTTGGTCAAACTAATGTGTGGCACATTCTCAGGTAGAGTTGGGTGGATGGATTGACAAATCCCTCCAAGTCTTCTTGCACAAGCAACAAGATCACCAAACAACTTTGTAGGAATGTCAACTTTTGTTGACATGACCGCTCTAAACTTTTCAAGGGCAGAGGATATTGTCCTCTCTCCCATAAATGGGAGTTGCCTTGAAGAGACTAAGTGAGATGAGATTTGAAGTAACTTCATAATGCTCATATTCCTAGCTTCCATCTGCTTTATTACTAATGTATTAAGCAATGGTTTGAAGATGTTATTTGTTTT

>Manitoba_picorna-like_virus_1

AGAAATTACGTGAGAATACGGTGCCAAGGTAAGACAATTACGGCTTGGGGTTTAAAGGGATCAACTTTCTTGATGCCAAGCCATTTGAGCAATTTTGTTAATCATGAGGAAGGCTTTGAGATTTTGCTTATGCAGACAACAGATTACATTCATGTCGGTTTGGACAATCGAGAAGTTGTGGTGGTTCCTGGAAAAGATTTGNNTTTGCAGATTGTTCAAAGTTTTTGCAGAAGAGACCTTTGTCACAGATAGTAGGACAAGAAGCAATAATGATGGAAGTTGACAAAGAAGGTGTAGTGGATTACGCAGTTCACATAGACGCTCTTGTGGACACTACTTCGGCCACTGATGCTAAAGGAAGAGTCTATGAGAATACTGGAGGGGTAATGTATTCCTACCAGAAGGCTGGATTGTGTGGCTCCTTATTATGCATGGATGTTATACATCCTATAGTGTCTATGCATATAAGTGGTAGTCAAGCGCAAGATGCTGGTATTGGCGTTGTCTTGTATCAGGATGATTTTCAGGTTCAGGGAGAGTCTTTGGAAGACTTGGAGCTTGAAAACATAAGTCGAGATTATGGGGAAGGTGTTGCTTTGCAGGTACTTGGAAAGGTGGAGACAGAACTTTCCAGTTTTATCCCCGTGAAAACTGAGCTTATAAAGAGTCCTTTGGCTCCATTTTTGAGCCCATGTCTCACTATGCCAGCGTTTTTGTCGAAGACGGAGGATTACCCGCATGAATTTCCTCCATTGTATTATGGAGTTAGAAAGAATGGTGTCTTAGCAGTTTCGTTCCCTGAGAAAATTGTTTCTCGAGCTTATGAAATGGTTCAGTGTTTGTTGCTTAGTGGAAATAAGATACCCGGTGTGGAGTATAAGGTTTACGAGGTTGATGAGTCTATTTGTGGTCTTAACTTGGATATTGAGAAGACTGGTGATGTCTATTTTGGCCATATTCCACTTGATACAAGTGCTGGATGGCCGTATAGTACCAGTGCGTATATGAAGAAGATTGGTACAGGGCTAAAGAATAAAACTCCATGGGTTGACGTGCAGTATGGCGCTGATGGTTTTCCTTTTAACGTTGTCCTTTCAGACATAGTAAAGGAGGATCATAGAAAGAAAATGGAACTACGCTACAAGGGTGAAGCTGCTTTCAATGTTTTTCAGGATTGCTTAAAAGACGAGCGACGACCCATAGAGAAAGCAATGAAACCGGGAGGAACTCGCTTGTTTTCGATGTCAAACTTTGAAGGTACTATTGCGTTACGACGATATACGTTGTGTCTGGTTAATCACATGCGCTTTAACCGCATTCGTAATGGTATAGCTATAGGTATTAACCCTGAATCAATGGAATGGAATTTCTTAGCTACAACGCTTAAAACCCATCCTAAGTTTNNTTTTGGGGCTAGTCTTGAGTATAACTGTGGAATGGCTTTTGCACGGTTGATGCAAGATTTTGCGCAGTTGTATGAGTTTGGTCATGAAATGGAAAATGTTTCAAATGCGCTCATGAAGGAACTTATGGGTTCGTATCATATAGCTGGTAACGTTTTGTATAAAACATTTGGTGGTAGTCCATCAGGCGCGTGTATAACAGGTGAGATAAATTCGTTTGTCCACATGATGTATATGGCAGTTTGTTGGTTGATAGTAGGAGAAATTGTGTATGAATTGCAACATGAAGTTAAAGGTAGATTCTCTAATCGCTATCCTGAGGTTGTGACTCTTATTGAGGGGCTTAACTTGGACAACTTGGAGTTTTCCCCTGAGAGTTACCGCCAAAACTTTGTCATTGTAGTATATGGTGATGATGGTGTGTATTCTGTTGCAGAAGAATACGCTGAGGTGTTTAATGCCAAGATTATTAGCGAAATCTTGGCTCGACATAATTTGGTTGTTACCAATGCAGATAAACGGGCAGAGGTACGACCTTACTTTGGTTTTGAGGATTGTGAATTCTTGAAACGAAAATTTGTGGCCCATCCATTGTTGGACAATTTTTATTGTGCACAAATGAGATGGGAGGTTGTTGAAGAACTAGTGAAGTGGGTGCGACGCAAACCCCTTACTGCAGCAGAAGCTAGTTTTTCAAATGCTGAGAGTAGT

>Manitoba_rhabdovirus_1

CTCACGGTCAAGGCTGGTAAAATTCTGTTTCTTGATTGCTTGAAGTCTTTGTAAACAATCTTATTAACTGATACCTTTTATTCTTAAAACAGTGGAGGAGTTTTGACCAAAAGATTAGAATCAAAACGCCGATTAGAACTGTTACAGCGCATGCTGTTAATGTATATAACATAGGTGTCCAGTGCAAATCTGTCAAAGAGCTATCATCCACGAAATTGTAATCTGTCTCTGTACCGGGGTGTAGCTGTCTTCTCCGGTATTCGGGAGACCGCTTTGCCATCGGTACTCTATAGTGAGAGCTTAGTCGTGCGATTCTTTTATAGTTTTCTATGTCTTGGATGCCAAGGATCAGTGTCTTATTCTTGACATACAACCCATTGGGTCCATCGATCACATCTGTTTTATTTGATGTAACCCAGTACGGCCATGTCCAGAGTCGGTTGTCTGATCTTCTTAGACTAATATACGGAAAACTCAGGCTCGGTTCGATAGTCACTTGTTGGTATAAGCTTAGTCCCATCTCAAATTTTCCTGGACTATAGCGATAGACTGGATGAAATCCGGGATAACGAGGTGTCAATGTTTGCAACTCCGCGCGACTAACCATTTCTCCTGCTATGATTTTATCGACAACTATCTCACACTCATGATCTAAAAATTCATCCAACAAACTGAGGACAGCGTTCTGTATAAATTCTTTGTCATTAATTATGTTAACTTGTGACCCAGTCGGGCAATCCGGAACTTGGTAAAAGTACTCCCCTATCCAGTCCTGGTGCGGGATGTTTTTTCGGTCCAGTGCCATCCAGTCCCCAGAGGGATATCTCAACCCTCGGTGGCCGCAGAAATTCATTGTACATGGTTTGTCATACTCGGATATGTATATATCAGGACTCCAACTAGCTACTGTATGATTGGGAGTGTAATATATCACTAAGGATCCATCCAACATATGAGAATCATCACATGAATCTTTTATCGAGTGATCCGATATCCAAAGACGATTGTAAAAATGTGTGATACAAACAGAGGACGAACAGGTTCCGTCCAAGAAGATCGAGGATTTGAAGGCATCAGAATACGGATTGTACGAGACAGAGTGAGAACTCACTGAGATGAAGATGCGAGAGGTGGTCGAGCTCTTCATCCAGGAGCATGAAGGCGTTGGGTGCTCAGAAGATGAGAATGCTCCCTCTGATACTTTCTGGATCTCCTCTCTACATAAAATCTCCGAAACATCTGATGGTTCAGTGTGTAACTCCAAAGTGATACCTCCAAAGAAACCCTTAGAACAGGTTGTACTCAATTTTATGAAAGTGCATATGGATCCAGCAATATTAATGGTTGTCCCTATCAAAGGTAATCTTATTGGAATTGTTATGCCTGATTCAATTTTTGCATAACTTTCCCCAATGGGACATGTTAGCGCACTCGGTGTCACATCGTGCCACTGAATATTGTCAGATATTGGAAATAGCATTGTATGTTGTTGTCCTCGTAATGAGTCTCCTAATGTACCCTCTGTCATGGCGACTATTTGTACCGCCACATAGATAAGACTCTTGATCTTCTGGCGAGAGCAATCCATTATTATGACGGGTCATAGCATAGTTGACTTGTGTTAGTTTTTTTCTTGAATAGCTCATTCGAACTCTGAGGCTCAGACAAGAGGCATCCCAAACGGACTCAACCGCCACCACAAGAACAGCTTCTAGAGCTCGTTATCTCCTATTACCATATCCTATATAAATACAAAATATAAGCCTATTAGAATTAATCATAAAATTTGATATAGAAAATATCTAGAAATATACGGTGATGATTTTGAAGCACGCTAGTCTGCAACAGTTTTCTGATAAGCTGTTTAGTCCTGAGTCAATTCTTCTTTATAGAAATACACTCTTTATCGCATTCCACCGTTATAGCATACGGTTTAAAGAGTGATAACTTAATAGTATGCCTCTTACGATATTTATTAGGCGAACGAATGCGAATATAATCCAAAAGAGAGGTATGACTCATAGTACTTTCTTCAGACGTTAGATGGAAGGACACCAGTATATTTGGGATTAATCCTATACCCTCACCATCATAATCATATGTAAGAGGGAATATGGGTGAGATGTTGCATAATTTAATTTTGAAAACTCCGGTTACGGTACCACGATAGTGATAAAAGGCTCCAACTTTATGTATGTATTTGCAAGTGAACCCAGCCAATATGTAAAACAAAAAGATATACTCCTTGTCATTGACATAGCCTTTATAATCATCCTTGATAATCTCTAATGACGCCATGAGTTCTTTGTAAGTCCTAGGAGGGTCACGGAATTTTATATCTAAGAGCGCATCGACTCTAACAGTCTTAATAAGAGACATTGTTACTCAAAATTGCAATACCGAGTTTTTTTCTCGATCTGTGTTACTGTTTTGTGATTTTGATCGTTATCTCTCGGTAACGGCATGTCAAAACGAATTGATTGTAGGCCTTATGTGCTCTCAGAATTGCCTTCCCCTGTTCTGGTTCAGTGTCATAGGACTCCAAGATTTCTAAGCAATCCTCATGTGGGTACTCGACAATGATCGGTTTGCCACCCTTCTTACGGGGCATTACCACCTGGACTTTCCCTCCTCTCTGAACCGTAACTGGAGGTTGCGGATCCGTGGGCTTCAACTCGAGTTGAGGATTTCTCTCCGGTTTTGATACCTTCTTAGCCTGAGAAAGGTTTCTTGGCTTCACGGGCGGAATGAAAGGCGGCGGAGTCTCCTGCTGAGATGTTCGTGTCCTCAACTGAGGATTACTCTGGATACGGGAAGCTTTATTCCTGAGTTCACCAATTAAGAATTTACCCTCTGCACGGAACGCAACGATCTCCTTATCTGCCAATTCGATTGCCTGAGATAGCATTAAGTTAACTCCGATCAACAATTCATCAGCAGTGATTATCCCATTTCGGTCTGGGGTCATTTCTACAGCCAGTTCCTTGGACTTTTCCGTGATCTCACCTGCCAGTAGACCTAATTCGTGACATTCTGCCAGTGTCTGGAGATCGATATCAGATGGTTCCGATTCTCTCCCGAGAAGAGTTGCTGTCGAGGAATCCGATCCACCGCTAGCGAGATCGGTTGGATCGAAATCGGTCCCACTTCCGCTTTCCTGAGAGTAGTGCTGAGTCCGTTGCAATTGTAGGTTTCTCTGCGAGCTAGTAATGATACCGCTGGTTTGAACTGATACTGTAGCAGTCTCATTACCTTCTAACTTCGATTCCCGTGACGGCCCGAGCTGGACGATCTTACGATTGTTTTGATCTGGAAACGTCTTTGGAATCACTCCAGTCGCAATTGGTGCCTCTTCATCCAGTCCAGGATCCAACCCGCCATCTCCAAGCGGATCCAACACCTCTCCCAATTTCTCCTCATTTAGAATTTTCTGAGGATCCAGAAGGCGACCCGGACGTGCATTTGGCTTTACTTTCAAGCTTGTACTAGCCGGTTTGTTCTTCTTGCTTTCCGTCTGTTTTCCTTTTCCACGATTTCCCATGGCGAGTTCAGATTTTCCCTCTTTCTTGCCTGAAGTGTCTGGTACACTGGCAAACACATTCACATCTGAAAAGATTGCAGACTTATTCTCCATTTTGATATGTGTTAATGTTTTTTTCACGGACATTACGGATCTAGGAGCAGCGGTCCGGCCTTCTCATACAGCAGTTTTCCGATCGAATCATCCCGACAAGCAGGATTTTGCTCCCACTGCGTAGCCACCTTCTTGATGATGTGATCAGGTACCTCTCCCTGATTTTTCAGAATATGTGCCAACCACAAGGAGGACTCCTTCTCCTGTGGCTCTCCTTCTGGCGGAGGTTGCGGATCATCCTCTTCTTCCAATTCGTCTTTCTCGCCATCCACTCCGAATTGCTGAGAGAGAGTCGCAAACTTGCCCATCACATAATGCATTACGACTGCATTGGAGATGATGTTGTTCACTTCTGGATCCCCAACTACTCGGGCTCGTTTGGATCTTTCGAGATTAGAGCTTACTCCCAACGTGTGATAGAAGAAATGAAGATCCAGGTTCACCGAGGCCGAATAGGGTGATTTAGAAGACAATCCCAAGTCCATGAAGTACATGGCATAAGATCTGGGATTGTCCATTTCCTCCCCTCCTTTCACAATTCTCTCGAATTGGTCGGCGCATCTGTCGGTCCAGATCCACTCCGCGAATGCAGGGAACGTCAATCCCAGACACTTGGTAATCATTCGGGTGGCAATGAGAGCTGCGCAATCCTTGAATCTGGTGACAATAGTTCCGATCCGAGCTTGAGAGTAGGGATGAAATGGAAACTCATTCAGGAACATATCAATTGCGGCCATGATCTGCACATATGGTTGATAGGCAATCCACTGCTTGCAGGCATCAATTATATCTCCCAATTCTGCGGTTTGATTCAACCCCAGCGGTACCAGAAGGTTCTTCACATTCTTCAGGACTTGTTCCTTGTACTCTACCCGTGCAATCTGTCCAACTCGGTAAACGCTGCAGATCAGTATGATGCCCCGAAGAATCTCCTGTTCGGTCAAGAGGACGTTTCCCTCGACCACAGCTGGCACCGATTCCCTTTCGTCCGGAATGATGATCGCACTTAATCCAACGCGGTCTGTTCGCTTTCCGATGTCGCGATTGAATGAAGACCAATCCTGATTGAGGAGTCCCTTGACCAACTCGAACTCGTCGTATAGGAAGCGTACCACAAATCGATGGTCGACTTTGCCTTTGGTTAGTCCTGATCTTATCGCGGCATGAATCTGTTCTCTTGTCGCACCTGTCCCGGGTATCATCATTATAGTCGGCTTAGTATTACCCGGCTTATTAAACCAGGCCGACGGATACTCAATGGATCTCTCAGTTCCTATGTCAAAATGAACGATAGGAGTAGCGGTCGCAGGATTGTCAGTGATACGTTTGATTACACGTGTCTCAGCCATTGAATACAAACAAAATCGATCTGATACTTTCCTCAGAGGTACTTACACAATTCTGACTTGAAGATGTGTTGACGGTACTAGTCGTTTAATTGTTTTGTTTTT

>Manitoba_rhabdovirus_2

CTCGTAAAATTCGTTTTCATCCATAATGATAGATATTAATTGTTTTTGGGTTTTTTATATAAATATATTCAAGTTAATCTTGTTCTAAAGGACCATCTATCCTAGAATAGTAAAATCCGTTCTTATGAGCTTGCAGATCCTTCGATATTTTTCTATTAGTTAAGTGCTTTGATAGAGTGTCGTATAAAGAGGTTATAATTACATTAGATTTAATGTTTTTATAGACCAAATACACCACCATTACTATCAAAAATATCAACAAACACATCTCTATTTCATGCCACATTGTATGAGTTTTTTATATATTCTTCTCTTTTATTTTTCTTGAATCAAGGTTTGAAGTTTTTTTACCAGTTGACAATACCATCTTCTCCTCGAGATAGGAGTGCGATTCTAGGTTTTCTTTTCTTATTTTTGCATAATTTACACCATAATATGTAAATCACAATTAAAATTAAAATCGCTTCTGCAGCCCAAGGAGATAATTTGATCAACAATTTCATTATAGTATTCATACTTTTAGTGTAAATAATGTCTTCACTATGACCCTGTGTAATTTTTATATCTCTCGGGTATGCAATTATACCAATTTCGACTTCATCTAAATGGATTCTCAGATCTTCTTGATTAAATAATTTCCCATAGATTTCAGTTAGGTTAGCTTGTATTATCCCATTATATGAGCATTTTGTTTTCTTTCCGCACTTCTCCCATGATGCGTATTTATCCAGTTCTTCTTTCTTAACTTCTGAATATCTCGCTAGAGCTTTCATCAAAGTATTATTTCCATTCAATTTATATACTGGATAAATCCCTCCTGTGCTAGGTATGAACTTCATTAAAGACTTGTAATTAATTTCGTTCTTACTCATTAAATTGACTTTTTCTTCCAGGCACTCTTTTAACTCTTCCTTGATCTCAGATATCTCTGTATTGGAGTATAACTCGTTGTTAGAATGATTACATTTCTTTACATTGAGAGGTCCTGGCAATTTGAAGTAGACATGATCTTTTGTGACTGCAAGAGTTACCCCACAATGATCATGATAACATATGTTATCCAAATCATAGACTTTCCCATTAACATCTGCTAATATTGTACTGCTTGTTTTGGATTTCCATATTTTTATTGTTAAATCTGAGTAAGAAATTTCTTCAAAGGTTGAGCAAGTGTGTAGTCTTTGTTGATTGTCTCTGATCCAGTAACCTTTCTTATCTTTATATTCATAGATCATCCCTTCTTTGTGAGAGAAGATCTTATCATAATTGGCTAGGTTGTTTAGCTCAGGATCATATATATAACTTTTCTGTCTACTGATAATATGTTCTTCACTCACGTAATGGTATGAATTGTCAAAAAGACCCCCTACGCATTCTTCTTCCGGAAATTTTATCACATCTACAGATGGTTGTTCACAATTTTCGATACTGGTTGTTTTTCTCTCTATTTTTTGCATTATATCATTTGCTGCAAAATAATGGAACTTGCATCCCGTCTGGAATGAAGCCACATAACAATCTTTTCCTTTTATCTCGTAAGGGTTGAGATCTCCATCTTGAATATCATATTCATAAGAATTATAATGATGTATCATATCAAATGATTGCTCATCTATTAAAACTTTATGATATGATGGGCATGATAATTCATTGAATGATATAGCCTTGAAGGCTGTCAATAGTAATACTGGTAGCACAAAATGCATTTTAATAGTTTTTGGGTTTTTTATATGTCTACTTATATCATATATACTAGGATACAATTACATATTATCAAGATGTTATGTGTTCATAAACTTCAGCAGAGTCTTCCCCGAATGCCTTCTTTATAATATCTTTGTCACATCTAGGGATAAGCTTCGTGTTGCATGATGGGTGTAGCTTCTTTACTTTGATGATACAATCCCCTTTGATTTCATGACCATCATAATACACAACAAAGTCCTTATTGAATACAATTGTCTTCTCTGAATCTTCAAACACCCAGTGTTTCTTTACCGGGAAAGTTATATTCTCCACTTGGTAGCTGGTTGCTCTCCCGTATACGTCACTGTAGAGACCCTCTACCAAAGCCTTGTACAATATATAATTGTAAATACCTTCGTATTTTAGAGGACCTTTATAGTGCTCTTGAATCAAAATTGAACTTTTCTTCAGAGGCTTTGCTTTATTCACTTCCAAATTAACCAAGTAAGCGATATGCATCTTTTGATCCTTATAATTCCCCATGCTATTTGATTTTCTTTCTTCTCTGATAGGCTTGCCCATAAAATTCATTGTCTTAAACATTTTTTATTTGTTTTGAGTTTTTTATATACTCTCCTAATTTGAGATTACTTTAAGACAGACAGTACTCCTTTGTAGTCTCTCTCTTTCATTTTTAATATAATCTTTAATCCCTCAACGGAGATCTTAGCATTATATATCATCCTCTCAGGATCCATTTGACCTAGACCTTTATTAGCAGTCAAAATTGACTCTAAGGTCTTCTCTGTGATCTTATAATGTCCTGAAAACGCCTTAGGTACGTTCCAAGTTTTCGTACCTTGATCATAATAACTCTTCTGATCTGACACTTGTATTACCTGTTCCAGTTCAGGTGCTGTAGGTATTTCTGTATAAACACTTGTTTCTTCTACGAATTTATTAGTATAAGGTAGACAAACTTCCTTATTAAGTATATTGTACGTTTTCTCTTTAAAGTCAAATCTGATCCCAAATCCTTCTCTTTCTAATACACCATTGACAAATTTTTCCATCTCCGTAATCGTTGAGGTCTTCTTTACAGTCTTTTTAACCTCAGTATATATAGGTGGGGCTGTGACAATCTCATCTTCAAATGATACCTTCTTGATAGTCACTTCTTCATTCATCTTTTGATCATTTTGCTTGTTCGGAAATGCAATTTCACAGTCCATGTTATCATAATCTAATTTGTTCTCATTTATCAAATACGTATCAGCCCTATATGGATTCAACAATTTCAAGAATTGTTCATCTCCCATCACCTTCTTGGACTTATCAACAATAGTATTTCCACTTGCCATATTTCTGAATCGTTTTAAATTGGTTTTGGGTTTTTTATATATTGTATTAAGTATTGTATAAGAAGTGTGCATTTAACTCATCTTGCTAACGAGATAAGCTCCGATTGTTCCAGGCCTATCATCAGAGATGACATTCTTCTTTCCTATAAACCAATCCCAAATGATTCTTGCTTCTTCTCCATCATCTCTCAAAGCCTTCATGTTCTCTGCAGTATCAGACATCCCCTTATCATCTTCTCCTTCTCTATTAATAATCTCCGCTTCTCTGTTCTTACTTTCTCGATAGATAGTAGCCTTGATATTAGATTTCGACCTGTAAGTATATGCACATGCCAAAGCATTTAGAAACACTGCATTTTGATTGCCACCTGCAAGCCTAGCATTCATAGATCTTTCATTGCTCAATAGAGCGCATATCATATGCATCCAGTTGTGAAGATTCGGATTGGCTGTAACCGAAAATGGTGATTTACCCATTGCTCCCATTTCCTTTGCATAATTTGCAAACTCTCCTGAATATCCATTGCTTAGTATGGTCATTATATTAATGATTTCTTTAGCAATATTATGATCAAGAGCATGTTCCCATGCTGCCATGGGTTTCTCACTCAATGACGTTAAGAAGTGATCCATCTCACCGATCAACGCGCAATCTTTGAATTTTGAAGACTGAGTCGCCATTCTCAAAGATGAATAAATATGATTCTTCTCTTGAAACAAGAGTAAATCTATTCCTGCGCACAATCTAGTAAAATTTGGATTGTTAACCCATGTCAAAGCATTGTAGTAATTCTCCTGTATCTCTTCTTTGATGATTTGATTCTGCACAAATGTCCTATTCATTCTGGATTTTATGTTCTGAATATACTTATTGTCGGCTCCACCTGAGATCCCACAAATCCTGTAATTTGCAACTAAAGTTGCAAGAATCTGCAGTTCTTCAAATGATTCACCTCCGGTTGTATTGTCCCCAGACATCTCCAAAACTTCTCCCTCTCTTATAGCAAATATGTCTGAGATAGACCTTTCTCCCGCTTGTAAGATCACTTTATCTTTTAACTCTAGATCCTTTTTAATATTTACCTTGATATATTTCTTGCTAAGTGCATATAGCAAATATAGAGCAACTTCTGGCTCAATATCTTCAGGCTTTCCTGATTGAGAAGAGAAATATTGGCATACTTCTCCTAGTTTTGCAGTACCTGCCTTAGGCACAGTTAATTCTGGCTTCTTCCCGTCTCTTAACCATTCGGTAGGATCATCGGCTTCTTCAACCAGCTTGATTGTTGGAATTTTCTTAGTAGCCAGCCTTCCATCTTCTCCTTCATATCTGAATGTATATCCCTCCATCTTATGTTGATTGCTTTTTCTTTTTCCTTTTGCTTACGTTTGTCTTATTGGTTTTGAAGTTTTTTTGTTATTTTCTTATTTGACTCTTATTTTTTACTGTGAGATCCTATACTCTTTACTTTATTATTTTCATTTTCATTCGTTCTATATTTTTTATTTTATTCTCATTTTCTCATAATTTCTATTTATTCTTTACCTCTTATACCTTAATAATATCCCTTAATTCTATAGTTTTCATGTGTTTTGATCGG

>Manitoba_rhabdovirus_3

ACCTCGCACACCCATTATGATAAACAAAAAAGAAAAAACGAAAATCAACAAAACCATTTTATTCTTATTTTTGAAAGAAAAAAGACCTATCGATTAACTACGATATGACTTTTTATAAATATATCTAATAATATTTATGCAACGTGTAAAATATACGTAAACAACTAAAATTCCTAATTGTTACTTGCATGTATATGACCCCATACTCTCCATATATATTTAAGGTGTCGAGACTCCCACTTAGGAATATCCCTGATAGTGTAATCATAGATAACTGTCTCATTATAGATTATTTTATTCGGCAAAGAATCAAAACAATTGAGATATATTTGGTACGCTTCCTGTATGGTTGACTCTGCTATGACCGCCTTCAATATGTAACAACATGTGATTGCCTCTGTATCTCGATCCATAGTCTGGCTGGCGCAAGGGTTGACTGCCAGTCGTGATGCTGAGTCCCACGGTAATTTCAGGGATCTATCAGAAGATAATGTGGAGAATATTGCTGCCAACTTTTCCATATTGCTAATCTTGGCCTGAATGCATACTGTTAACCAATCAGATACCTTGCGAGGTGTAATATCTTCTGGAGCAAACATATTTGTTAAACAATACAATGTGTGTCTCAAGTTGCTGTGATAGCCCAATTGCTCTCCAATTAATAACATGGCTCTTCGTATATTATCTTGCCTGCTTATAAAAGGTAGTGTTTGAGAGATTCTGATTTTTTGATAATTTTGTATCTTAATCAAATATTTCCTGAAGTTGTCTTCCACGTATAGGGCATCAGTTATTTTAGGTTGTCGATCTAACTTGAATCTTAATACTTGAGCTACAATAAAGCACTCTCTGTTTTCAAAACTTGAGAAGGCAGGTGTAACAACTTTGACATCCATGTAATATAGCCGTAGGTTACTGATTGTTGCTGCCAGCATCATGGGATTATCTATATACTCCTTCCAAATACATGCATATGGTTGCTTCTCTATTATGATCAGCAGCAATGCATTTATGATTCTGATCTTAGTTACTGGATCATACAGCCCTGACACATCTGCATCACACGTTAATAATGAGGAGTGTTGTGGGATCACCTCAACGAATCTCCTCAGATATCCCATATCAGTAAGATCACCACCGTAGATCAAGCTCAACTGATGCCCTTTAATCTTTTCTCTATCTCTCAAGAAACTACCAGGTACAAAACTTTCATACCTCTGTGTTTGAATGTCTCTACTCTGCAGAGAGTTATAATACACTTCCATATTGCTATAACACGAGATTGCATGAGCTACTCCGGCCTCCCCGTCGGCAGTGCATATGGCAGGGTTCACAAAAATGATATTCTCTTTAATTATCATCTGCATATACTTAAGATAAGCAGTGGTCAATGTTCCCCTAGTGCGCCCTATTTGATCTAACCTTGTACGTCGCACCTTCAATATTGTATCCTCTGGCATAGTCAATGTCTCTAGACTAACCTCTGACCAAGTATCTGAGATAGGTTCAGTTAATACTAGGCTAGGAATTGGATAATTATGAGTGTATATCTCTGTCATCAAGGCCGGATCAATTTTTCGATCTGTGCGAGGTGTCACTTGGATAGGGGGTACTGCAGACAAACTGACAGGATCTCTGAAATAAAACTCCGGTGGTTTCCTCATTAATTTGATTCTTTTAGTTGTGGTTATGAGTCTCCATAGAGTTTCTGCATAATCATGATGTATAGATTCTTTGATAACAGTTCTCATAGTAGAATTCCAATCTTGAGTGGTTCGAGCGACCTCCAACATAGAGTATTTGACACCCTCGTAATATCGCTCATCCATGGTGGTGGTCATCTTGACATAATACCACCACATTAAGAATACACTCACAAATTTCACGTTGTTTAAAAGGAAGAACTTTACTTCTGGATATATGATTGCTTTTGTGGGTATCATATGAATTCTGTCAAGCTCTTTAACTAACATTTCATTGAGCAACTTGGTAATTCTGTTATTATGTTTATAGCAATTATCCGATCCATCTATATCCAAGATTGACATAAGTTGTTCCATTACGCTTGGGACTAAACACAGTTCGGAGATTCCGCACCACAAATCGGACTGCATGTCAGGTAAGAATACACGAGTTGTGACATATTTATCACCTATGTATAAATATAGATATTTAGCAAAAGACTTGACTATGGCTAATAGCCCCACATTCAACACCTCGCTCATGGAGATCCTAGTTATATATGGTGCTTTGACAGCTTGGATGGTTATTAACGCATGAGAAGTTATACCTTTGAATCGAGACATCATGTAATAACCTATGGCATCAGATGATGATATCCCGTCAACATAAGTTATTAAACAACTTTGAGGTGTGACCATAGATTTGAAAGTTGTGATATCAGCATACAATAGTGGGTTGCTTTTTGACAACGCAATGGGCGGTCCTGGTACGAGACATGCTGTAGGTGCATTGTCATATGCGGTGTGGCAGCAGTCATTGGTCCTATAAGCTCTCAAATATCTGAACTGATAGCTAGGATCTGCAATGCACTTCTTAATGATGTCATGTATTGCGTAATGATAAAGTCCACTTACCTGTAGATTGAAATCATCTCCACCGCGTGTCGTTAACCCCAACTTGTCACTGCTTAATGCAAAGAACGTGGTGGCAGAGTGTCTAATTGTATTTTTCGTCTCTTTCTTTGTAGCTGCATCTCCCACGCGATGTATATGCGAGCCAAAGTATGATATACCGCTCATACCGCTGATAATGTCATGTGGTACATTTGTTCTGCAACTTATCAGATCTGAGAGATAATCATAAAGTGGGCTCTCTTTATCTACAGCCCACTCTTGTGCTCTCTTCAATCTACATGCATCATCAAATGGTCTGCTCTTGCCACTGATTGTTACAATTCTACCAGATGTTTTCTCCATGGTTTTAGTGCCAACAAAGGCCGGCTGATTCCCTGCTTCCAATAAGTTAGCAAATGAATACGATTCAGGTGCAAGTACAGATATATACTCATCTCCTGGTCCCACATCATCATACACTATTTTTAATACATGTGATGGATGAGGCATGGTTACACCTATATACCTGCGATCTCCAAGAAGTGATTTATAACTCACCTCTCTTATGTGATCAGCCATGTGTGTGGAGCATATATCAGTTCCGAAGTACGGTTCTAACTTTTGTATTTGCACATAAGTTTTCAGCATATGGATGTAAAAATTGAAGTCCGACTCTTGGATCATGATTGTAGATCCGAATTTAGCACCACCAAATGACATACCCTGCAGAGTCTTTGTGTTGGTGAACTTCGATATCATATACAGTCGAGATCCTGTGGGTGAATGAGTCGATATGACATTCAGCAGTCGTGGATAAAGTGGCTCAGAGGAGATGAGGTACTGATAAAATTCTCTATCAAATTGCTCGGTTGTAGCATTAAATAGTTCCTTGACATCTACATTCCGAGAGTTGTTAGACAGGTAATTGAATACTTCTGACTTGAAGCTCCTTGACAGGGGTGTCGGAGTTTTGATGTTAGTTGCTGTCGGATTGCACACCAAAAGTTCAGGAGAGGCTACTCCCACAGGGAATTTGCTGTCCTTGAACCATGTTAATAACTTGGCAGTGACCTCTTTTATCTCCGGTGTAGATGTCTCTGAGTCTCTCATCTTATATATGAATGACAGGTATGTTGTGAGTGGATCAGGGTGACCTCTAAACTCGTAATCCATCAATGACAATGTAGGTATTCCTGTAAATTCCCCACAGCTTAATAAGAAGAATCGCAACATGGGTACAGTCTCCAAGCTATGAAGGGCCTGTGACATGGTGGCTTTGTCTTTTGACAATACAGAACAAACTTCTGCATCCCTGAGCAAGGTCAGACATGCTTCCATCTGAGCCACGAACCAAGGTATAGCCATTATATACGTTTTTTGGACCGCAGAGTAACCAGCTGTCTGGATGGTACCGAGACGGACACTGCAAGTTGGATAATCATCATTTACATCTGCCATAGTACGCCCTACTCTCTTAGATAGTTGTGATGCATTTGCACCGTTGATATAGATTTCTTTGCTATAGATCATTGTTGAATCTGAGGCCCAAGATTCTAGCGGTTTTAACTCAATACCAATCGCTGCTGATACCTCGTTCAAGACACTTAGGAATGTCTCTTTTGCATTCTTGATGGCAATCTGGTTTGCCAACATTCTGTCTTTATGACTCAACATTGACACATCACCGACTGGGATCTTCAATTTGCAGATCTGATTATCACCTTGTCCTATGATGGTGGACTTAATCCCGGTGCGAGCTTCCACAACCAGCAATATCCCAATGGTAATTAAGGTCCATAGTTTTTGCATAATACCTTCTAAACCTCCCATGTGATTCAACCATCTGTACAGTCCGTCCGGTAAGATGAGGTGAGGATGCTTTTTTGACATGTAGAATTCAGGGACGAGAAATTTGGATGACAATGTGACTTCTGATCCTGCAAAAAACAAATGTCCAAATGAAAATAATCCTGGGCAGTCAAACCAATCATCCAGAAGCTTGGAGACATCTCGTGTGTTGAAAAGACTCCAGTGTATATTCCATGATTTGAAATCAATGCCCAATGTTAACTCCAATGTTAAGGTGGTACCCTTCGATCCCGACATATTGATTAATCTCCGAGACAACTTCTCCTCATCCAACGTCATGCTCTGTTGGGGAAAATACTTGAAGACTCGCTTGGATATGTTGCTCTCCGTTACACAGAAATATAATCTCATCTCTAGAACCATCATAGCAAACAGCCTGGGTTTGATGTTGACCTCTCGCTCTTTTGGTGAAAGTTTTATTATCCACCAAGAGGATGGTATCTCACCTCGTTGTATCAGCTGGCAGATCTCTTTCACAGATAATCGCTCTCGTTCCATGATCTCACTCAACACACGTCTGGTGGTAGTGGGTTTAGATGTGGGCACACCGGTTACATGGTAGTGATATGATGTATTATTCTCGCTTGCAAGTGGTGCTATGGCTTTGTCATTTAAAAGGTCTGTGTAATCTATGTGATAGTCAAATTCTAATTCTCCGATCACTCTGAGTTTTGACCAATCTTGAATGCTTATCTGCGAGTGATAATCACTTAGATTTCGATCGTTGCTTTTAACCAAGTTGTGCAATGGGCTCTCTTTCCTCACCTCCTCAGTGTTGCAGTTGGGCCATCGATTATGTTGTGCTATGAACTCTAACACAAACTGTTTGACAAGCTGGCCTGCAATCTCGGTCATGGTCTTCTGGCATGCAGGCCTACGTGTCTTGATTATGTCTTTAACTTTCTTGCAACCTGCAGCTTCATCTACCACTGGATGTCCCCAATGTCTGTATATCCCGAAAACTTCTGCTAGCTGTGGTACAGAGAACTCTTGCAACAGATCATCCATAACGGCAATCAGCTCATATTGGTCTTCAATACTGGACTTCAACAAATATTCATGGAACTCCTTCTTGATGTCAGTGATATCAGTTTTCTCTATGATTATTGAGTAAACTAGCGATTCAAATACTTTTATTTGGTTATACATCTCATTATGGAACTTGGCCAGACCTTGATCAAACACTTTATATACAGCAGTTAGCTTATCTGCCGATATTTTGAACTTGAACTTGCTGTGTATAAGTCTGTGACTATACATGGTATACAGACGTCCAATCATTGTGTCCAGCATGCACAGGAGATGCTCATAATTACATACATGTACTTTGGTTGCACTGTCCAGCAGATTCTGCATAACTGAGTAGTTCATGTGCATATGGCTTTTAACTCTAATCGGAGACTCTATAATTAGGCACATGCTTTTGTTGACCAGGATATTCGGGTATCGCGCAGTTGGCAACGGTAATAAGGACCACTCATCAATTCTGCTCTCGGCCTTCCTGACTATGATATCCCGCACATCCTCCAGTGACATCTTGATAGCATACATTGATTCTTCTTGCTCATTGAAATATAATTCTGCAGATGGTTCAAATTCGGATGTTAATCTGGCATCATAACCTCG

>Manitoba_tombus-like_virus_1

CCCCCATATTGGCTCCCCTTGCGGGGAATATGGTGCAATTTCTTACGTTGGTGGAGGCCCTCCTTGCGGATCCCCTCACGGGGTTTCCCAACGCGTACCAGCGTCCTGCAGGGACGTTTGATCTGGATTTTTCCTTTTCCGTTAGCCGTGGTCTTATACCGTCCTGGTTAAATCTTCCTTTGCATGTCATGTTGTGTTCCGCGCGGATTTTCTAATTCTAACTTACATAATGTTTTTTTTTTGTATTTGTTGGTTTTGTGATATTTTTTTTTTTTTTTGGGATGGAGCCGTTTTGGCGGGGCAGCCCTCCTTTCCGCTGTTATTTTGTGGCTTAAACATAGGTTATTATTTTAGACATGTGATTAAAAATAAAGAGAGTACGCTGATCAGGCGGTTCTCTATACAACTTAGTTATAGGTGCGTTTATATTGTAATTTGGATATGCTGTAATTAAACTAGGTGTGAGATAGTCCCACGTGTATGGTCTATTTCCTAATTGTTTATTTAGAACGTGTATTCTGTCGCGCTCTAGTATCTTCACTCGAGGTCTATCAAGGGGGGCCCACTGCCTGTGGACCGGGCTGTTCTGATTAATCGCCCAATTAGCATGTTGCACGTTCTGCAGATAGCGCTTGTAATTACTGTCGTTATATCTATGGTATGTGCAGCCATAGATCGCGCGTAGTCTGTCAGGGTCAAATGCCATTGTTGGGTGACCGTCCGGGTGATATACCAGCTTAGTGCGACAAAACTCAACAGTGCTAATGTCAGTTTCAGAAGGTAAAAGCTTACATTCCATATTGTATTTCCTAAGCCTGTCTCCAAATCTTGCGGTGTCGATAGCAGCTGTAGAAAATAGGATGAAATCGTCACCGTTAACGATTGCATCCCCTTTATAGCCGAGATCGCGGAGTACTGAGCGCAGGATATACAAGTTGACGATACAGTTGCCAAAGCCCGTGTCAACATCTCCCGACATCCTTCCGCCGTCGAGGATGTAGCTTTCACCATGTCTGGTAGAGCACTTGTTTCGTAACGTTCTGCGAGATAATTTGTGTAGTGTTTTGTCCTTGTCGTAGCATTTCGAATAAAACCTGTGACATAGCTGTAAGTGTTCACGTAGGTTGTGTGCGTCGAAGCTGCTGTGGTCTCCCGCAGTGTAGTAGCGGTATTTCCTAGCCAGCTTGTGAACCTTTGCAGCGATTTGGTCATATGTTCCTTTTCCAAAATGGATGTTAGATTTAAGCGTTCGTTCCAGTGACTTAATGTAGCAACCGTACTCGATGTTAAATGTTGCATGCCGGGCATGAATGAGTCGAGGAGCCCGGTACTTGTCTGTTTTAACTTTCTCCAGCTTTGTGAAAGGGGTGATTCGTGATATGACTCTTCCGTCCTCACGCACCATCCTAGAAGCTTCAGCGTACATTCTTCTCTTTTCTGGTCGGTCCATAGAATCGATGTACTGCTGATGAGTCCACTTAGCTGGTTTCCTAACCAAGTCACGGAGGATTCTGTCGTTTTTGAATAGTCCTGGGTCGTAGTCGTGTAGTCGTTGTGGTTTGTGTCGATTGAGGTAGGCACCGATGGTGTTGTGCCCACACTGTGCATGGTAGGTGTAACTGTCGGGTTTGTGAATAGAAGGCCGAAAAGGAATGACCCGATGTAAACGGTTGTGCCTAGTGTTAATAATAGCAGGCACGTCAACAGGAGTAACTGTAGGAGAGAGGTAAAATCCGCGCGCAGGGAGTTTAAATGACCGAGTGCCATACGGGTGAATGGGAATTCGTAGATCCCGTTGAGGATGCGAACTGTTTTGTTCTGCACAAAGTTCAAGCAGTTGTTGTACTTTTCGATGTTGTGCAGATCGTAGTCATACTGTGGTCGTATGTTGACTAAGTGCAGATTCATCACCCCATGCATTATGGCTAGTTCCTGTTTGTTCGACACTGGTTTGCGAAGGAGGAAGTATTGTCGTATGATTGATGCACAATAGGCTGTGTATGTTGCATCGACAATCTTCCCTTCCACGCGAACTTTCAGGAAGGCAAAACAGTCATCCATGCTTGGCGGACTGGTTTTGTCGTCAATTGTCTTCCTGTGGTCGAACCCCTCGGCTATCGCCCGGTCCCTGCGTAGGTCCCTCTCCGCCCCTGCTGCTGCGTCTTCGTGGTCTGCATTATTGATTGTTTTCATCAGGTCCACCACGACCTTCAATTCGGCTAAGCTTGCCCCAAAGCTCTTCGCCGTCTCCTGGTTCACTTCGTTCTGACTCTTCCCGATTAGTGGCATGTTGTTGCGTAGATCCCTTTTACCTTATGTTTTGTGTGTCGATGGGTCCCA

>Manitoba_toti-like_virus_1

GTTGGATTCTCTGTCCTTTAACTATATAATATATTTTATAAGCACTTACTCTTCAAAGACGACGCTAGTAGAGCTGTCTAGGTTCAGCTCTTTGCCGATGTCTACACCATGGCCATAGAAGACATCGTCTATATCGGCCCACTGCCTAGTAATGGGTGTTATCTTTGCCTCGACGAGGGCACGATTCACAACGTCCTTGAACCTTTTGTATTTATCAGGCCCGTGCCCATGGGCCTGCAACAGAGCAGCTTCACAGTTGACTCTAGTGGAATCTTTAAGAGATGCCCCTTTCCACACCCACTGGGTTGTGCTAACAATTGATTCCCAAGCAAGGGGGGATAAAAACTCTCCCTTACGCGTTGGGTGCAGCACGAATTTTCTTTTAAGAAACGATGCTTCTGACATATGCACCCATGCTGTTATGGCGGCGGTTTTCTCAGCAGATGTCGAAACTATGTTGTGCGACGCGAAGTATTGGTGTATGGTTATACCATTAAAATCACCTATATAGTCATCGCTAACCGTCATTATGGCATCATCCCCATATACTATGAGGGCGACATTAGCCTTAAATGTCTCCACCCAATCGTTGCCGCCGACTATGGACTTCCATGCTAACATAATGTACATCTGGTTAACCATACTGTTGACTATGGTTGTGAAGAAAGCCCCACTTGGCGAACCACCTAGTTGGCTATAAACAGTATTGTGGCATATATGTTTACTTTGAATGCACTCGTAAACTAGAGACCACATCTCTGTCTCATCAACACCTTGAACATAGTTCTTGGTCCAATCTATCATGAGAGAGTAGGCGGCTTTAGCTACTCCAGCATTGTATCCAGGCCCGAAGTTCGAGTAGTCTATTGTGATGAACTTCGTATTGATCCTAGCCAATCGCTTAATCATATCAGACCACTCTTCACTTAGCGGATTGCATCCAACGCCGTGCATCAAATTGCGGCGGGCGCTCATAAAAGAGGCAACAAAGTGTGAGAACGTCTGGCGACATGCTATAGTGTAATCGGCACTCGGATTACATATTACACGTGTACCCCCCCTCTTCTTAATCTTTTCCAATGACCTTTTCTCATCCTTCAAAGTATCGATGTAGAGGGTCTCAGGTATTACTCCATTTCTACGCAAAAGCTCTTTACGAGCTATCTCGTCAGAAAGAGCGGGATCAATCCATTCCACACCTCTAACACTACCGTCACCATTACGTGCGTATGTAATGTAATCTTCTTTAGCTGTCTTCCGAGTGAGCACATAGGGAAAACCCATGCTCGTGGACAAATCCATTGGCTTGTAGTAATCAATACCCAATCCAGTGATTGCTTTATTCACTGTCAGGCGTTCTGGATTCATCACCAATGGTTTCATTGCGCAATACCACCCATCCCACAGTGCCTCCTTAACACAAGTAAGCTCTTCACTCGTGAAGTCTGTGGTCAATAAGCCGTGTTTTCTCACTCCCTCAAACAACGGAGTGGTGTCAAAGTCGTACCGCGCATCCTTCTTGTGCAAGACAGCAGGTTGCATGTCGGTGCTCATACCACAACTGTTATAAACCAAAGACTTAACTAGCCGTGTTGTCTGGGGTAAGTAGGGTACTTTCTCCTTAGGAAGTGAACCCCCGTAATCTAGACGAACTTCCATATCGTATATTATGGAAGCGTCCATGATAGTACCGTGCTCAACTTCTTCCTTGGTGAGCACAATATCGGACTGTGTAGTGGGCAAGAGCGCCTCTTTTGTAAGGACTATGCCATAACCTGTCCCCTTATGCTCTTGACCAAGCCCAGCAACATGCATAGACATAATGGGGCGATTACTACCCTCCCTAAGTAGCAATGATCCACACGCTCCGGGTCTAGAGTAGGTGTACTCTAAACAATCACGTATCTCAAAGCATTTGTTCTCCTCATCAGCTACAACGTAAGTATCGAGGAAACCTATGAGATCTACCTCGATAAGTACCATGTAGTCTTCACCACGCCTAGGTACTGAGAACAAAGTGGCGTTGGCTGGTATAGATGCCTGCATTTCGTCCTCCGTAGCAAGGAACTTGCGCAAGTCCTTGAATAGGGGTTGCGATGGTTGCAACTTGAGCATAGCTATATCGGTATGCGTGGACTCCGTGATATCTTTCTCACATATCACTAGGAGCGACCTCAACTGGGGTTTGTGGAGGGGGTTACCATATATGTTGTAACCCACCTCCATTGCTTTCCTGATTTCGTGTATGTAGTGCCTAGGTAGAAGCAAATTGTGATTGTACAACCCAACACCGTACAATACGCGTTCCTTGTTACTACCTCTATCTATAGAGATACGGAATGTGTTATTGGCTATGTACGACTTAGCCACGGCGTACGGAGTAGGAGTTTCCTCGCCTCCTTGGAAGTACCTGACTTGTCTCTGCCTATCGTAGCGTCGACGCATTACTCGGTGCGGTTGATTCGAATCATATTCATCAGATTGGGGTAGCGCCACAACGCTCTGTGCTGTGCAACAAATCTTATATATGATTGTCGCTATTGCTGTAGCACCAACAAGACCGGCCAGTACGTAGCTGTACTTATGGAGGGTCCCTGTTAAGTATTCCCACCACGCTTTTGGTAGTGCAACATTACTTGGCTCCACCACCCAAGCAGGTCTAAATAGTGCCGGTACAGCCCTGCACGTTTCTCCCGAAGGATCGTTAATGTAGTTAACTATAGCGGTGCGCAATACACGCACATTAGCCTCCATGTACCTAGTAACAAAAGTGGCATACTGGGTACTGCGCGCGTAACAGCTTTCGGAGTTGCACGACGCCATGGGTACGTCAACCATAAGATTGGTTTGCATATTGGGCACACGCCACATATTGCCGTATATCAAAGGCGACACACGCGGGTTAGACAACAACCCATGAGTACAAACAGGCAAAGGAGCACAAGCTTGCATCAATTCCTCTAGTAACTGCACGTTTACAACAGGCTCCATCAAGTCCTGGGTGGTTTGTGCGTCCACATAGTCAGACTGGAATACAGCTTGCACAGCAAATCCCATAAAATGGCCCGTAGCATAGCTCTCGTAGGCGTCCAACCTCCCTGTAATAAGGGACGCTAGATAGTACAAGACTGCCGTCAGTGTAGGCAACGATGCCCCCTCATATATGGTGGAAAGGTGCTTACCCAACCATGCTATCGGCATGCACCCGCTAGAAACTAGTCGAGCTGCCGCATATATAGAGTAGCCTTGCTCAGGTGTAACGCAAGGTATGATCACACCGTCACGACAAGTGGGGCAGCCAGCAACGCGTTCTGCCATCAAATTATTGTAACATGCAGTACATAAGTAGTGTGGGGATTCCAAGTCCACAGAGTTAGAGCACGCATATGCCACTACTCTGGTGTCTAAGCACACGTTACACCTGCTAACGGTATTGGAGGTCTTATTGACGTAGTAATCCGAGATGGAGCGCAATGCTTTCATCGAGTACTTCGCCACAGTTTTCCACATTCCGCTTCCAATAGCAAGCCCTAATATAGTGTGACCCAAATTCTGCTGCCTAGGCAATTCCCAAGGCATAGCTTCAGGTTGAGATGGATTCTCAGGAGCTGGGGCAACAGAAACCTGGTTGACAATATTGCTAACCGCTATTTCTAGCTCTTCGTATGGAGTAAACGCATTTTGAGCGAGCGACTCATCCTCACGTATAGCAGTATTCAATTGGTAAAACAGGGTAAAGGGGTCATTTAACTGTATCTCCCTAACGTCCTGGGAACTCAAGGACGATAAAAGTCTATCCATCCGCTTTTGAACATTGGCAACTTCCGTAGCATGATAGCGTTGGAAAGTATTCTGGAGGTATGACACAGTCTGTGAAAAGGATCTAATAGCTCCATCTAGCACTGTGCCGTCTTTCACGTTCGGGTACCTCCTAAACATGAGGTGGGGGAAGTCGTTTAGGGATCCTATAGGGAGGTCCCTCAAATTGACATTCCGATACTCCTCAGCAAGCTCAGCCCGCAAAACTGTATCCCGACGGCGATAAATGGCCTCGGGATACCTAGCGTAATTCGCTAAGTTTGGAAAAGCCCCATTACACAATATTATAACGACCAAGGGATTACCTCGGCGCTTCTTCTCCTCCAGGTGTGCCATCTCAGGGATGAACAGTGACGTGGATTTCAATTTATTCAATTCCACTATCATATCGTTACATCTCTGGGAATCTTGGGAATTAAGAAACTCGTCATAGACTATCACTGGTTGATTACTATAACCCGACCAGAATTTCTCCCCAGCAGTCCTGTAATATATACTTGCTGACGAGGGACACTGATAACCGATAGACTTGAGGAGTTGTACTACAATTTCCTCACAGGCTGAACTCTTACCAATTCCAGGAGCCCCTTCTATACACAACACGTATGGTTCGTATCGCACCGGGGACGCTGATAAATCTGCAAACCTCTCATTTGCCACTTTTATAACGTCCGAGCATAACTTGGCTAGCTGTGCGTTTCCCACGCCTGATGGAACAGAACATAAGAGTCGCTGGTATTGGTAAGCACTAAGTACAGTCTTCCAACACCGCAAGCGAAAGTTACCATTCGATAAGAGCGAAGTAGATGCTTCACTGGTGACTATCTGGGCTTCGCGTATGAAATGAGCTATCTGTTCACTCCCATCGGCTAGCATCTTGAGAGCACAAGCCTCTGGTGAGACATAACCAAGAGCATGCATGACATACTCCTTAATAATTTCGAATGTGGATTGCACATAACGCAATATAGACAAAAGATACGATATACCGGAGGTATTTGTGATTCGCTCCAATAGAGCTCCCGGCACACTACGCACACGCTTCGGGTCTATATAAACACCCATTAGCGTACCCACAATGCCGGCCAGGATACCAGTGAGCGTAGACTCATTAGAAGGGGGCCCTTGTGTGGTAGGAGTGTCCTGATTTAATTGTGCGTAAAACGCACCCAACTGTGGGCCAAAAGCCGATAAGGCGAGAGCTTGTGACGTGTCAAAGAAATGGGATATGAAACGCACCACGGAAACGCCTACTACGACCCACGATTTTGAGACCCATGCACTGACAATGTCTAAAAATAAGTCGAAAAATATCCTCGCGTACGAGGCCACGGAAGACGTAGTTGAAATAACTTTGTCTACGGCTGCTTTTATAATGCCCTTAAATTCGTCAACAGACGAAGACATACCGGACGCTAATCTCTCTAAGCTGGTAGCAGCTTGATCGATTTTCGGTACGCTCTGGCCGAGGGCGTTACTTAAATTGCTACACGAGCGCATGGCGTTCCTCACTTCAGGTGCCAGCTCGTTCACAGCACTCGCGGCGACTAAGCCACCACCAACGATTGGTATAGCGCCTAGCGCCATACGTGCCACCGAAGTTGGCGAAGTAGATTGTATAACATTATTTATAGCCCCTGTGATGGTATCCATCTGAGTTTTAGGGATACTGCGAACCGAAGTTCCAGGATCTTTATCCAGAAAAACCGCCGGTGACGCAGGTTTCTTAGTAACTCCATTTTCATCAGTATACACTGTGTACTGACCATAATCGATACAGTAGGGTATACCATGGAAATTGGCTACTTGGAAGTCATCCCCTGCTGACCACCACACTGTAACCTTAGCCTCAACTTCGGATGACAGTATTAAATGGCCAGAGTTGTAGTTACCCTTGTCACGCCAGGAATAAGACAAGCCAGGGTTGTCCTCCTGCATAAGAGTCCAGACGTTTTCGGTGTCATACGGGACCTCAAAGGTTAACATGGGATTAACAGCCGGAACTATTACCTCAGTGACCATACCTACGCCCTCCAAAGGTGACCAGGAGATGTGGTTGTTATCACCGCTAGTCGCCTTCTGTATATAGGCGTGATTACCTATAATGCGAGCGCCTGTGTGTGGCATGTATGACACATAAACTGTGGCTCCGTTCCGCACTAAGATTGTATAACGCATACTACCTCTCCACATACGGAACATATTGCATATATTATACTGGGCAGTGTGTTGTACCCCAGGGTTAAATATAGCACTGGCTATGTGCGTCATAGGGTCATTAATCATGGACCTATTCGGAGGCATCAACGGTATAAAGAAAGCCGTGGCAGTATCACTATCGGGTCTCAAATCGATAGTGGCCCTGGTTATAAGGCAGGGCATCCTCAATCTGTCCTTGAAATCCATATGACAGTCCAATGTTTGGAGCTGCGTATTTACGCGATTGACGCGGAAATTAGGCGTATCGTCTTGCGCTACAGCATCCATCTGTGTCCTAGGCATTTCATTCCACGCAGATCCTGTGGCCCTTGCAATAGCAACGAGCTCTCTCGAGGTCATCTCTATAGCAGGAGGGTTGCGCTTGACAATCCTATCTCTGCTATAACCACCCCCGGGTAGCAGCGTGGGGTAATCACCCGGGAAATTATCGATGGGCTTAATAGTATTATATTCCCAGAAGTGCCGCATGGACGCTGGCTTTATGCCGTGCATAGCAAAATCCTCACCAGCTCTGATATATATAATACACTTAACCTCCTTGGCTACGACAGGGCTACAACGCAAAGGTGTGCTAACTCGTATTTTAAGCATGGCTTTGGTTTGTGCTGCAATTCCGATAGCAGACGCACAGACCCTTGGATCTGGCTTAAGCCCATAAAAGAGCGGTAGAGGATCAACCGTTGACCTACGCATGATGGTATCATAAACATAGGGTATGGTGTACTCGCACACGTTACCGCTACTGAGGGGGTAGTACTTGGAGTAAGAACATGACGATGCCGACATATTAGACTCCTCACTAACACGGTTGAACTCCGTGCCTATCATGACTTCTCCACTGTGGCCCTTGTTAGCAATAAAGCGGAGTTTGACCTCTATGGTACCAGACCAGAATTGAAAGTTGGTCATAGCGTACTCCAGAGGGGTAGGCTCGCCACGGTATGTTGTAGGCCGTAAATTTCCACCATTATTGCGGCATGATGGATCTACAAAGAATTCGGCCAAAACAGCACTTACGGCATCGGTGGTTCTCCACTCAAACTGCTTATAGTAACCCCATTTCCGGGCTAGCTCCCAATAATTGCTTGGGTCGTTCTGCGGTATCTCGATCTTGTCGTAGTTAGTTTGGGTATAAGGGTTCAAACGCAGTGAAACTACGTTGATAGGTCCTTTCCCTGTTGCGAAATTCATAGCAGGGTGGGGGATAGCTATGGTAGACTGGGTGTCACTAGGCTTATCCCGGTTACGGGCTCTCCCTCGCACGTCGTAAGCATTTTCATACTTGCCAAGTATGCTCCTTAAGGCACCATTAGCAGCACCTGCAATAACATCAGTTATCCCCAACATCTGGACCACAGCTCTGGCAGTAAGACCAGCAAAGTGAGCCCTTGGCATTCGTATGTATAGAGTGCCAGTGATGGAGGATGGACCATTCTCGCCAACGGCAAGTGGCGCAACACACATGACGGTTATAGAGCAAGACTCTGCTGGTCTAACACCTTTTGACACGCCAGGTGCATTTAGTAGCCTAATGAAGGGTGTGTGGTATATGTACGGGCAGTACAGAACTGCATCGTTGTCCTCCGATAGATCAACCTCGACGTGACGTCTAGCTTGTTCAGATTGATGTCCCGCCATGGCACGTTTACACTGGTACGAATCATGCTTCACTCCCATGGTGTAAAATCCAGCAAACGCATCATTACCAGGCACTACTAGGCGTACTTCGAAATCCATGTTAGCGTAAGCATGGGTCTGGAAGGGAAGCAGAGCAGGGGAATTTGGCATCTGCGTGTACAAAAATTCCGGTAGCACACACGTAAAGATAACTTTACCTATGGCATCACTGGTAGAGAAACTAACCTGTTCCAGGTAAACCCACCTATTAACAAGATCGTCGTAAGTGGATATCTTCTCCGATGTTACAAATTCATCCAACGAAATATCGGGTTGCTCGGGTTGGGCAGTAATATCGGGCCCGGTGTGCACCATTATGGTGTTGCCAGAAACATCTGCCTCCGAGTCCATCTGAACGCGAGGGATACACGCGCACTTCCTAATAGCAGGACAACCATATCGACCTTTGTACGTGTTACGAAAATTGTACATCAAACGCCAGGATGCCTCATTGGTGGCTGCCTTGACCGTAGTAGTAATGATATCAGCTTCGATTACTACAGGTCCCTGCAACGTTGACCAAGTGAGGGATCTTCTACATCTTGCTGTAGCGGGGTCTCTTCCACGTCGCGTGGTGTTGGTGATTTGGCAACTTGCCGACAAATAAAAAGTAGCATTATACCAACTATTAAGGACAAGGGTATAGCTAGTGTTGCTATGATACGCCTCGTCAAAGGAGGCAGCGTATCTATCCAACGATTTAGTGAGCGCATCCATGCACTCACTCTTACTAGAGTAGACTCTAGCCTCAGCTCTCTCGGAACTGCTCTTCTCTCCGGATACAAATGGTCCCATGATGTAAAAGTTGAAACGTCGTTAGACGGGTAAACCAAGTTGTTATATGGCGTACCTTGGACTAAAGCTGAAGAACAATCTTCACTATTAGCTGGGACAGTGATGGTGCAAGTATTTGTCATACAATGACACAATATAGCACTGAAAATCCGGTTGTTCGCAAGAATAAAATAGCAAATTACACACGCAAATAACAAATGCGCCCACGGGTGTAGGACACAAAGGTAAATGAGAGGTGCAAAGGCTAAAATAATTGCTGCACGTTTTGAATGTTTATCATTCGGGGTTTTGTTGGTTGTTTGGCGTTAATAAACGCGTTGTTGTGTGGTTCTTTCTCTGAAGAATCAAATCACGGTAATAAGGCACTTTCTCTGAAGTGTGGGCGGCGTACAATATGAAGAGTTGTTCTCTGACAAATC

>Manitoba_toti-like_virus_2

CTCACTCACACTCTCACTCATAAACACTCTCACTCAATATCTCGATATATCACTCATCAAACCAAATTTATTTAAAGCTTTAACAGCTTCGTCAAACTCACGCTGAACCACACGTTTGGCCATAGAAACGTTTGTCACCCCCCTGTGCACTAATATTTGTGCAGCCCGTTCCCTGGCCAAGTCACTCACCCAAGAAAGGATGATTGGATGCCATTTAAACTCAAAAATCGTCACTCCTACATTCTGGCCGTCCAATAAAACTTTCGTCACATTCACACCCACTCCACACACTAGATAATTAAACGCGATGATGGCTCTCCCTTGCACTGATGGTCTACTACACATCCCAATGCAGATGCGCACTATGTCAACCCACTCCCCCTTCAAAGATACAGCAGCAGCAGATATGCTTTTGAACGGACACATCGATATGGCCAGATTTACTGTTCGTAGAGGCCCACTCGTTTTCTCATTCCTCTTCAGGATGTGCCCAGCTTCAAAGTTCTCTATTGTGTAGTCCAGATCCAAGAATTCACCTGGTTTGCTCAGAAACTCTCGCATGAGCTCATAGTTCACCGTAACCTTAGGGAGGACCAAATCCTCCCTCCATGACCTCAAATCACGCTCCAGAGCCCTCAGACCTTTTAACTTGTCTTCAGGCCTAAGTGAATCGGTCACATTACTGGCGTGAACTAGATCAGCCAAAGCTTCAGCATCGAAACTGGAACCTACTTTCTTACTGACATGCTTGATCCAGTCAGTGCTCATATTTCTAGGGACAGCATCAGCCATATCTTCACTGTACGCACGATAGGTGGGTAGGGGACTAATAGCTTGGGTGCTGAAACTCATCTTGCCAGGTGGTAGTAAACCTAAACCTCCCAAGCTTGGGTGCAAGAATAAAACGCACTCTGGGACGCTAAAACCACCACCAGGCAAAGAAACATGGGCCGCAAAAGGTACCATTGCGTGCCAAAGTTTATTAGCTCCCTGCTCAGTCAAACCTCTCCTACGTAGTATACACATCTGATTGTTGATGGCCAATGCTCTCTCTGCTGGCCCGCTTATGTCCGCACTCTGAATGGGTTTCACTATTAAGGTAGACACAGCCCTGGCTAAATAACCCATGCATCCTTCAGCACTATATAAAACACGTAAAAACTCAGCACGACCAATATCGAACATTTGCTTCTTGGGTTGAAAATCAAAACCTGTCTCTTGCATAACTTTAAAGATGGAAATTGCCCACAGTCGTGAGTCGTTTGATATCCAAACATCATCTCCCTGATGTATGTTAAACAAGTCACGCGGAGCCAGGTCTAGATTATCCTTAACCCAATCCACAGCCACTAGGAAATAGGCCAAATTCAAAGCTGTATTTATAAAATTTGTTCCTCTACAGCCTGAAAACATTCCTTGCAATATCCGTTCCGCTTTTCGAAACGTGGGAAACCTACACCACTGGTTCAACAAGGATTCTGCCGTCCAATTGGCAGCTTTTACTTTATCAGGGTGAGCATTTGCCATCTTGTACAGATCAGCTATGGCATAAAATACCACAGCTTGAGCCTCTAAGGTGTGCTGATAGTTGAAGTCGGCATAATCTATCATAGAGCACTCAACCCCTGGCTGCTTGGCTATATTCATCCTGTTAACTAAGCTTCCAATCACGTCTATTCCTGTTAATCCAGCCTCTATCCCTTCTATTGCATTCATGTTAGGCTCTATTCCCTCTAAGACATATGCTGATATGGCATAGTCTATCGGCTTCGTTCCATATATCGCTCGAGCTTTACCCATTTCATATTTCTCGCTTGCTACTGCTTCGGTCTTCGGTGGGCTATCCAGCCAAGCTATCATCTCCTTTGATGTTAATTTCTCAAAAAGAACGTGTTTATTGATTTTGATGACCTCACCGTTGATCACCATTTTCTCACCGCCCGTACTTCCCGATGAGACCCAGGACTGCCTGTTGATTATGAAGTCTTCATAACTCTCAACGAAATCCTTCCCGATAAACAGTGGTTTCAAAATTTCAAACACCTTCTTCCTAAGTATCTCGCAGTACTTCCTATTGCTCGCTTCATCGCGTTTAATTCCTAGTGGGTCGCCCAAATACATTCTCACAGTCGTTCTCTTTCTTCGTTCCTCTTCCCAGTCAGACACATTCTTAGACCTACCAATCGCCAGCTCCCATCCGGCACAGGTCCCTATCTCCTCTAGCGTAGCCTTACTCCCATCAGGCCATCGTGCAGACCTTCTCAAGGCTACGCTTATCAATTTGAAGATATCGCTGTACATTGCTGAATCTACGCAGCAAATCCCCTTGCTATGGACAACAGAAATTATTTTCTTAACGCAAGGGTTCCCGTAGCAATGTATAGCGATAGCGGCGGCAGTAACGTTGCTAGCGCTTAATCCTAATAATGGTAAAAAGTAGTCAAAATTCTCCCTAAAACCGGCGTCATCCTGGAAGTGCCAGACCAATTGACTTACACTGACATTTGTTTTATGAGCATCCGCAGTACTACTATAGCCCCAAATTTCATCCAGAACGGCCATCTCTCCTTTCCTCGGCATTCTGTCCCATTTAAATTGTATAAAAAAGCACTCCTTGTGCTTAGGACATGGCTTCAACTTTAGCCAAGAATAGTCCTCCAGGCGCTCTAATAGCACCTGGTCAACGTGGAAGTCTCTATCAACACTACCTTTACCCAATAGATAGCGGTCGAGTTCTTCCCATAGCTTGAAACAGGCCATGCTATCGCCCAGTTGGTAAGCATGCCCGTTAACCTCCATCCACTTATCGTACAATGAAGCCGGTATCCACTTCATCTTGTAGCTCTTCTCTACCCGCTCCAACCTACTTAGCAGGCACACATGTGGTACAGCCATCCATAATGGCACATCTAACTCCAGATCAAACTCATTTTTAACCGCCTTGCCGGACCGTGGTTTGTTCGACAATTTCCAGCACTGTTTACATTCCAATGTGCCAAAGTTAACCCTGAAGTACTTCTTCCCCTCTCTGGTTAATGCTAGCTCTGGCCCATTGTGAGCATAACCACTTACTCGGGCCCAACTCAGTTTTCCGAGCGGCTAGCGCCTGCATCAGCCTCATCA

>Manitoba_toti-like_virus_3

TTTTCAAATTTTATAATTTTTATTTGATTTTTTATTTTTTATTATTTACATTTTTACGGGAAATTCACGTCAATTATTGAATTTCCCAGCCCATCATGAGCCTTGATTTACAGGATCCTTAGATCCTTAACACGATAGTTGTCATAATTCGGTATACATTGTTTGTCAAATCCGGTTTACGTTTGGACACCCAATCATCGGGGGTTCCATGGTCATCATTTGTGTTGTAATCAACTGTTGTTGATTGTCCGATAAACCAGCTGTTAATAATGCGATATGTCTATCAACACTGGTCTGTTCCTAGCTGATCGAATGTATCGATCAATTGTCAATCGTATGAACTTGAGAACTTCATTGACATCTCTAACTTGCTTCAGCAAGATCGTTCTCACTGCATAGTCCATAGCATAATCTTTTATCCAAGCCAATATAGTGGGGTGCAGTTCCCCCTGGTACGTGGTACTTCCCGTTCTAATCTCGTCGATCAAAACCGATGCGACGCCTTTCCCACAGTTATCGTTGAGCTGCTGCAAGGCCAAACTAGCACTCCTCCTCAAATTTGAGTTGAAATGGGTAGAGATAGCAACCTCCGCAGCCTCAAACCTGTCCAAACCTGTAGCAACCATAGCGTTCGTCAAGCCTTTAAAAGGACAAGAATTCAAAGCCCTAAAAATAGTAGAGAGAGGCCCTACTACTTTTCTACTCATCTTAGCCAAACTACCATTCTCCAACGCGTCAAGCACTTTATTGAAATCATCAGGGGCTGGGTCGCCAGTGAGAGACCTAGCATATAATTCGCGATTCCGAGTCACCTTAGGCAGATCAAGGCTGGCCTTCCATTTGACACAAGACCTTTCCAGGCTCTTCAAACCAGCCATTCTGTCCTCCATTCGTAATGAATCCATCACGTTGGCTCTATGTAGCACATCAACCCAATTGTCAGCAAGAATAGGAGCCCTAATCTGAGTACTGGCATAGTTGACCCAGTCTTGAGACATTTTAGATTCTATGCTCCTTTCCAATTCAAGGCTCCTAAATTCCATAGACGGTATTGGCGCTACTGAGCTTGTGTGGGAGGCCATTGTCCCTGGTGGACCTAGGTCTAGTCCATTACAATCAAACCGCATTTTTAGAAACGTGACTGGAATACTGAAAGTTCCTGAAGGAAAAGTTGATGTGGCAGCAAAAGGTACTACTGCTGACCACAACAAGTCTACCCCTTTCTCAGTCATCCCACGCCGATACAGTATTTGTATTTGATCATTTAACGCAGTGGCCCTCTCAGCAGGGCTAACCACCTCCACTGATTGTATCGGTTTGATTATTAGCGTGTTAACCGCTCGTGCCAGGTACCCGTTACACCCTTGACTGGTGTAGACAACTCGCAGGAATTCACCCACCTGCGTCCCAAACAATTGCTTCTCTTCCTGAAAAATCAATCCTGTCGCTTGCATAACGTTGAATAGTGCCATAGACCATAACCTCGAATCGTTACTGATCCACACATCATCTCCTTGGTGTACATTTAATAGATTGTCAGGTTTTAAACGTAGATATCTGAAGACCCAATCTCTAGCATGTCTATAATATCCAACATTCAACGTTGTATTTAAAAAGTGAGTAGCCCTACAACCTGAGAACATCCCTTGTGTGATCTTTAGTTCCCTGTTATTATTCGTGGGAAACTTGCACCATTGATTGAGTAGACCAGCTGCTACCCAGTCTGTTGCCCTGACCTTATCTGAGTGATAGCCTAGTTGTGATAGCCTTTTGGAAAGTGCTTGAAAAACAGCCGACTGTGCAGGTAGTGTGTGTTGGTAGTTAAAATCAGCATAATCTACCATAGTACACTCCTTTCTATCCTCCTTTACAGCCGAATATTTTTTGAAGACGGCATTGGTTACATCTAGCCCCAAAAGACCACTTTCAATTCCTTCAACTCTAAACATATGCTCTTCCACCTCAGATATGCTGTAGCTGGTGATGACGTAGTCGGGCGGCTTGGTCCCGTAGATGGCTCTACCTTTGCCCATTTCGAACTTCTCACTTCCTACGGCGCATATTTTCGGTTCTTCATCTAGCCAGCCAAGCATCTCCTTGGTGCTGACTCCTTCGAAATAAGCATGTTTATTAATTCTGATCGTACTACCATCACTCAAATTGGCCTTAGCACCTCCGGTACTCCCTGAAGACATCCACGACTGTCTGGTCTTACAAAAATCAAGGAACGACTTGTTTAACGGAAACGGCTTTACCATTTCCACCATGATAGTGTCAAGAACTTCATCTAGTTGTTTCAAATACAGAGCATTAGTATATTCAGATTTGTCCACCAACTCAGGCGGGCTCAAAGGCACAAAGCATTCTGTACGTTTATGCTGTTCCTCAACCCAATCAGAGACGTTATTAGAACGTCCAATGGCTAATTCCCAAGCCGCACACCCTGTCACCTCGTGCAGGGTAGCTGGCTTACCATCAGGCCATCGATTCGTCCTCCTCAGTGCAACACTCCATTTCTTAAACAACGAGGCATAGTCTTCCATTCCTAGACAGCACACGTTATACCTTTTGAGGTATTGCGACATGACGTCAAAATTGGGATTGCCCACCAAGTGGATTAACATAGCGGTGGCTGTTATGTTATAATAGGCGGTACTAAACATCCATGAATACTTGTCATACATCTTTTTAGCCCATTCAACTCTACTAATGTGCCAAAGCATGTTCTTAGCAGACAAAGTAGTCTTCACACGGTCTGCTTCACTAGAATAAACCCAAATCTTGTTTACGCAAGAAAGGTCATCTGGTTTATTGAGTTTAGTCCACTTTATTTTAACATAATAGCCACGATGTTTTCTACACTGCTCCAAAACAACCGCAGGATGAAGTTCTATATCAACAAGTCTAGCATCTGGAATCTGTTTCTTCTTAGGAATGCTCCTTTCCCATGACTTCCATAATTTCATACATCGTTCGCTAGACCCATGTTGATAATCACCACCATTGATAGAAAGCCAGTTTTCGTACAACTCTTCCGTGATATACCCACTTAAATAAGCTCTCTCTACTTCTTTCAATCTATCAAAAACACATAAGTGTGCTGTAGTCAAAAAACGAGGAATTGGTATTATACATCCAGTTGCGTCGGACCAAACATGAGTAGATAACAATCCGTTCTTTTCGCCCAATTTGCTACAAGTTAGACAACGTAAAACCCCGGCTCGCAGTTCAAAGTTATTCTTGAATTTGGCCCCAACTTTCAGTAGATACTCCTCACCACCAATCAAAACCCAATCTTCACCCAAATATGCTTTTTCTCGCTTCGATGTGAGCAAGTGTCTACGTACTAGGGCCCACCTTAGTTTCCCTCCTCTTCAGCAGCATCACGCCTACTTGCTGCGCGACTGGCACCACCAACTATTAAGTCAGACGCCATCCCGATAGCATAGCCAGTGGCTAGTCTCTCGGGGCGAGGTCCTCCCATAAAGCCGGCGTGAAGTGCCTTCCAATGCTCCTGACGGATAAGGAACTCCCACGTGTGGTCGGCAATTTGAGCTTTCAACCACACAGGTCCTAATACGCAACCTCGAGAATAATCGTATGTTGGCGTTCCAATTTTGATACTGAAGATTTCAGCATAAAAGTCGGGCTGTAGAGTGAGATCATTGATTAGCTGTGGTCCTATAGCCGCACGATTCCCTCCCGATTTATAGATCGGTGTATAGGTAAACGCCGGCATGACTGCTAGTCGTTGGTCCTGTATAATGATGTTGGCCAAAAACATAGCATACGGGATGAACAAATAGGGCCGTGGTCCCGCAGCGATTTCAAGATACTTCGATTCACCTTGCCAAAAGCTGAATTGGTTGCCCTCGTTTGCATTGGACATTATCAATTCAGCACCGAGGTTGACTTCGGGGGCCAATCCACTATAACCATAGATGGATGGCCACTTTTCAAGGATCCATTGTAATGACAAAGGCTCAAAGATGTAGGGGATCTTAATGCCAAAGGCATTGCGCCAGCCGGTGTTGTCCGTCAAGGCTGGCTGATCTACCCAGTCACCACTGAAACTCGTTACGACAAATGGATCGGCTGACAATATACACTTCGTCGCTTGTCGAGCAAAACCTACAGCACCTAGTATTAGCGGGATGGTGGGCATATCCATCTGTTTCTGAAACAGATCAGTTAACATATTAGACGATACTGATGCAGTTCCGTTAGCCCAACCATTTAGCTCCCGGCCCGTAATGTTGATGTTATGCAGTACTGCAGATACTCCCAAACTGTATATGCCAGCTGCTACCACCCCAACTCGAACGTAATCACCAAAATCAAACGAAGACATAGCTTCGAGATCGCGAGCATACTCAGCACTAACCTTGCCCGGTGGGCCCAGTTTTAATAGTTGCCAAAGTGGATTATATCCACCCGGTGTTGGTAGACGAGTAACCCCCGTTTCAAGGCCTGCTGTGAAGTAACGCTTTTCCTGAAGTCCTTGTTCTCCTTCAGTGGCCATTTTCCCGAAACCAATGGACACTGCCCTCAGATAACCACGTACATACCCATCTTGATCTTGTAAAAATTCAGCCATCCTACGTAGGGATGTGTACAGGGCTGAAA

>Manitoba_tymo-like_virus_1

GTTAGAGCAACCGTTGACCTTTGGCGCGTGAGCGGAACAGGAGTTAATTGAAGGCCGGGTAGGCGAGAATGAGTTCACCGCGGAGGAGGGTGGTGACTATTGTGTGGGTGTTTTTGGCGGTGGTGGCTTCCATGTTGAGCCAATGATTGACAGCAACACGGATGCGATCGGTTGGGAGGAAGGGGCTTTTGATCACGGGGGAAAAGCGGTTGAGAGGGCAGGAGACGGCAGAATTGGCGATGAAACCAATGGGGCCACCAGTGGAGATGGAGACTGCGCCAGGATGGTCGATCATGTCAGAAGCGGAGGGAGTGACATCTGACGGCAAGAAGCGGATTTCGACGGTGGAGGAGTACTTGGAGGCGGATTGGCGAGGAAGAATGACAACCTCGAGGGATTTCAGTTTGGCGAAGCGGTAAGGGGCAGTGAGAAGCTTGAGTTTGTCGTCGGAGGCGAGGTCGATGTTGACGAGCTTTTGGTTGCCAACATAATCGTGGAGAGTAAACTGGAAGGGGAGAGTCATACAAGGTCCAAGGTCTGAAGGAGTGTCATAGAAGCTGGGGCGGATCGGTATCGCCGGATTAGATAGTACGTCGTGAATATCCATGTTGCGAGAAGGAGCGGAGTCACGGTTCGGCGAAGAGCGAGCAGGGGCCATAGGAGCAGACGGAGGAATCGAAGCAGAAGGTTGCGGATCGGGACTGGAAGAAAGGGCATTGGAGGCGGCGTTGATGGCGGTGTCGGCGACGGCTGAAGCGAGAACTGGGGCGAGAGAAGCTAAGAAAGACATTTGTTCGTGTTGAGTCGATTGTTGAAAGAGACGATTGAACAAATGGGAATTATGAGCGCAAGCCAAGGTTATACGGGAGATCAGAGGGGGTGAAAGAGAGGTTGTCCTCGGAGTCGAGTTGGAAAGGAATGAGTTCAGGACGTTGGGTGCGGAGACGGAGTCGGACGAAGCTTGGAATCATGTGCCAGAGCGGGCGGGAGAGGTACTTGATTGATTCGGAGAGAAAGGTTAGGATCGAGGATGAGGGTTCTCCAATTTTGAGGGCGTCTTTCAAATGTTTTGGTGCGAAGCGGCAGAAGTAATCGAAGCAGGCGCTTTGGTAAATGGTTTGGTCTGCGGGGAGGAGGTTCCAGAAGCGATCACCGAGAGAGTGTCCAATTGCGAATTCGGTGATGTAGGAGGCGAGCTTGTCACGGATGGAAAGGTCATTGTGGGCGACGTACAATTTTGCGAAAAGGGCACGGGGGGCTCGAATTGCTCCTTCTGGTCCGATATAATATCCGCAGAACAGGGCGTGTGGAGTGAACTCGATTTTGAAGGTCAGGGAGAGGAGTGGTTGGACGGCAGCCCAGCGTTTGTTCGGAGGCGGGACGCGAGAGATGGCGGAATCATCACCGCTGACGCAAAAGGGAATTCCGTGGAGACTGAAGCGGGAGTTGAGGACCGCAATATTGTAGTCGGTGTTGTCGTCATAAGTGCCGGGTTCGCCGGTGAGTCGCATGCTGGTGAGAGGTCCGAATTGGCACTCGATTTCAGTTTTGAGCTTGAGGTGGAGGTCTTGGAACTCGATGGGGATGCCGACACTGCGCATTTTGAGAACCTCAAAAACGACTGCTTCACCACCTTGGGACTGGTCAAAGGCGGAATAGTCGTTGCATACTCGCAGTTCATTGGGTCGGAAGTGTTCAGAAGCCCAATCTGCGAGCTGAGAGGGAGTGTGAGAGGCGTAGATGAAGATGTTGGGATGGGACTTGCGGTCACGGATGATTCGTTGGTACTTCTTGACGGGGCCAAGGAGCAGAACGACGGCGTCGTGAGCGAGAGCGAGGGTTTGGCATGCTTTGGGGCCGGTGAATAGCGTGGACTCATTGATTTTGTGTTGGGACTTGGCGAAAATGCGGACGACGGTGTGTCGCCATGTGGGGTCGGAGCGGGCGGCGTTGGCGACGATGGTGGCTTTCGTTTTGCTGGTGAGTTGGTGGAATTCATTGTCATTGATGCAGTCGGCGAAGAGGATGGGATCGAATGGAAGCTGCTGGTTGGGATTGAGGTCGAAGGCGTCACACCAGGATTCGAAGAGGAGGTGGCCGAGGAGGATGTCGGAATCGGTGAATTGGTATGGGGAATAGGAGCTGCGAAATCGAAGGCGTTTTGGTATGGACATGGGGAGAAGAGTGGGGTCGCGGGCGGGCTGATGCCAAGGGGCGATTTGATTGGGAGCCTGGGAACCGTGTTCGAAGATTTCGGAGGTCGGGAACTGGTAGGAGAATTCGTTTTCATGCATGGTTTCGAGGTCGAACTGGTCTTTGGGTTCGAGTTGGAAGAGGGACATAAGGAGGAGGGAGTCGATGCCAGGGTAAACGGGTTCGATGTGGTCGAATGAGTGGATGCATGGAGTGAAAGAATCGGCGGAAGAAACGGCGGCTGGAAGGTCTTGATGAAGTGGGCGTCGAGTCTCAGGGAGATGGTCGGTCGGGGGTCGAGGAATCAAATCTGGGCCGGAAAGGGTGATGATTGGGGAGTCCCGGAAAACGTCGGAGTTGTCGTCAAGCATACGTTTAGCGAGGCTGGGAGCGCGGCGGGGTGGCAGTGGGGAGGAAGGTGTTTTTGGGCTGCTGAGCGGAGAATTCCCAACTTGACCACCGATGAGTTTCGGGGCGCGGGTGGTGAGTGGAGCGCGGATGATTTCGATGTTGCGAAGTTCGGGCCATGAGGAGAGAGGAATCGATGAGCTGGTGATGAGTGCATTGAAAATGGGATTAAGGGCGTTCTTGTTTCTGGCGGCTTGGAAGTCGCCGATGAGTGTGAGTGAGTCTTTGTGACGGGTGAGGGCGACGAGACAGAGGTTTTGGGAGATTTGGAAGATGGAGCGATCGATGAAAAGGCCAGTATGGTTGAAGGTGAGGCCTTGAGCGGATGCGACGGTTTGGGCGGGAATGCCACCTTGGACGAGACTTTGGGCGGCGGAAAGCGAGGCGGTGAGGGTGCGGAGTCGAGGGTTGGTGACGCGGGAGTAGCGAAGTGTTCCGGGGAGTTCGTTGTGACAGGCGACGTCAAAGAGCTTGGCGATGAGAGAAGGGCAGCGGTAAGTCCAACCGCAATAGACGTCGATGTATTTGGAGAGGACTTTCGTTTCGGGGATCAGGGTTTTCAGGGAGCTGTCGGGGTTGATTGAGTGGTATGAGCCTTGGAGAGGGTCTCCTAGGAGAATGACGAGGGAAATGGTCTGATCGAGGAGGCAGATGAGGTCGAGATATCCGGGGGGCATTTTGTAGATCTCGTCGATGATGAGGATGCGGGATCGGCGGGTGAAGGAGGTTTCCCAGGTGTTAATTCTCCAAGCAAGGTTTTTGGGGAGTCGAAGGGTGGTTTTCCAGTGCTCACGAAGGTCGGTGTTGGGGACGGCGACGCGCAGTTCATTGATGTAAGGTGAGCCGCGGAAGAAGGTTGCGATGGGGTGAGACTTGCCAGAGCCAGCGAAACCAGCGATGTGACAGAGGCGGACGGAGCGTGGGGCAGCGAAGTCAACTAGGGAGTCAAATTTATGGAACAATTTTTGGTCGCCCTCGGAGAGCAGGACTTGGCGTAGTATGCCATCGAATTGGTTTTTGAGGTTAGAGGAGAGGCATTTGGCGCGGTTGATGTTGATCTGGTGAGTGTGGAAGCGGATGAAAGGCAGCAGGTGTCCCTCGAATGTTCGGTGTGACAGGAGATGCTTTTCGAGAGAAGGGTAAGATGAGGGGCTCGGAGTGGAGTGTGAGCCTTGGATGCTTGGGGTGTGTGTCCAGTGTCCGGGTTCGTGAAAAATCGTATGGGTTGCAGGTGCGGCGTACGGACCAATCGGGAAGGTGGGGTACTGGGTGGAATTGACGAGGAAGCGTAAATTGAACAAGTAAGCGAGGACGGCCAGATGATCGATGGAAAAACCTTGGGTAACGGTGGTTTCGTCGAAGAGTTGGGAGTCGGGGAGGTGCGTAGAGAGGGCATTCCACAGCTCGGGGATTGGAATTTTGGTCAAGTTGGAAACCGCGTCGAAGAGGCAAGTGTTGCGGCTTGGGTAGGGAGGGACCTCGGTGGATGAACGCAGGCGGCTTGGGAAAGAAATGGAGGGGTCTTCGGGGTAAACATAAATGACGTTCCATGGGAGAACGGGGCCGGAGGCGGAGTTGTCGACAGAAAGGGGTGAGGCTTGGGGAGTCGTGGGTTCTTTGTAACTCTCGAAAAGAGCTGAGATTTCCTCGGACGGGCAGGGGCTGGGGGGAGGGAGGGGAATCTCGGCGGGCGCGGTGTTTTGGAGGAGAAGTGGGGCTTGCTCGGTTGGGACGTGCTGCTCAGTGACATCCTCACTGGGTTGGGTGGTGGTGTTATGGTCGTCAGGCGAAATTTCGGGAGGTTGGGGAGTGGGGTCGACAGGAACTTCGATGGTGAGCTCGGCGGTTGCAGCGAGTGGTGATGGAGAAGGTTGTCGGATTTGGACGGGAGTTGTAGGAGTCGGAGCAGTTGACGGCGTGCGAAGAGGAGAAGAGGTAGGGGTGGTGCCGAAGGGTTGAGAACGGGCGAGACCGCGGGCGGCTGACCGTTTGAAGCAGAGGCTGAAATGACGGTCGCAGAAGTATTGTTCGAACTGTTTTTGAGTGATCGATGCGTGCGGGGGGATGATGGCTTTGACGGCTAAGTATATGGCGGCGGAGGTGAGGCTGGCTGCGACGAACATTCCACAGTGGGCAGCGATTGAAATGTGTCGGGGGGCCAAGGAGGGAACTTTGATCATGAATGAGTGTTGGAAACTTGGGATGGTGAAGTCTGAGAGGAGCGTTAAAGAGAAGGGGAAACGGAAAATGACTTTTGGAGCGGGAAGAAGAAATGAAAACCACGAGGCGGGAGTTTCATGGCAGAGAGAGCAATTGGTGAACGGGATTGACGCAATCGGACGATAACGTGAGAAGTTGGAAAGCACGAATTGGGGAAGGTTGAAGGTGAGGATTCGGGTTGCGAGTTCGAGTTTCGGGATGGCTTGGATGATGGCTTGAGTGAAGTAAGTGACGGCGAAGGGGGCGGCAGCGAGGAGAAGATGGAAATTTTGACGGTGATTGGATAGGCTGCGGACGAAACGATTCCACCAGGTGAGGTGGGGGTGGTATTCGGCGGCGTTGTCGAGGGTGGCGGTTTTGAGGATAAAGCTGGTGAGGTTGTTCCAGGCGGCAGCGGTGACCCAAGCGTGTTCTTCTTTGTTCATCTGGGTACGGACAAAACCGGCGGGGTCGGTCACGCGGAGGGTGCGGACGGCACGGACGTACTGGAAGGCGTTGTTGTACACGGAGGTCGGCACGAGGCGGTTACGGATGTCGATGGAAGCGTTTGAGGGATTGGGAAGTTCAACGCATTGAGGGATTTGGAAAGTGATGTCTGTTTGGGGAGCGACTGGGCAGGTTCTGGAGATGAGGAGAAGGTGAGTTGATCCAAGGCTTTTAAGAACTTCGACGTAGAAGGTTGTTTCGCCGTTGAAGGTGTGGGTGGTGAGCCATGAACGTGAATGCGAGGGTTGGGTATAATGGTCGGGTGAGCCTTCGAGTTGGTACTTCAGATTGGAAGCATCGAGGCTGTACTTGTACATGAAAGGGAAAAAGCTGGCGTGGGTAGATCCGGGCAGGAGTGATTCAGGTGGGATGACGAGAGAGGCAAAAACTTTGTTGACGAGCGGGTTGAGGCGGAAAAAGTTTGCCATTTCGGAGAGGGTCCAGTAATGCAGTGAGTCGTGGAAGAAGGCGGAAGGAGTGGTGCAAGGTGAGAGGTCGACTGAAGGATATCGGGTGGAGTCGCGAATTGTGTAATGGGCGTTTTTGAGGGAGGAAAAGTGTTGGTTTTTCTCGTGGAGGTTGTTGAATTTCTCGGGTTTCATAAAGTAGACGGTACAGGCTTCACGGATGAAGGAAGGAAGGACGTTGTAAAGGAGGTCGATTTCGATTGTCTTATGTAGGGGATGAGGGTGGTTAAGGTGGCCTTGTGAAGGAATGGTGAGGCCGTGGGAAAGGAGGAAGGTGGAAGCTTGAGGACTGAGACAGTAGGGGAAGAGATTTTGGGCGGTCGTGACTGATCGGATGGCTGCGGAGGCGATTGGCTGGTAAGCGGAGTCACGATGAATGGTGCCACTGAGGCTTTCAAGGATGGTGCGGAGTGACATGTTGATTCGGTTAGACCAAGTTTTAAGTTGGTGTAGAGAGAGAGAAAATCAAGTATTACGTTGACTTATTTAAGTTGAGATATCAAGTATTAAGTTGATACGTAGTTTGCC

>Manitoba_virgavirus_1

ATTTACTTATCTCATGTAATTGTGATCGTGATTGATCTATCAAGCTGATAAGCTTGATTAATACACAGCATTTCTAACTGTAGTATTCTAATTCTAAGAAATCTGCTTCATCGACGTAAGCATCATTTTCTCTTTGTACCACGGAGCGCAATTTCTCTCGCATATTATTAGGTATCTTGCGAGTCCTTAAATTGTACTTCCCTTGGAACAAAGCGACAAATGATTTCTTATTTACCCGTAATAATGCTATAAAATTTACTAAAGTTATAGCTATTTCAAAGGAAATTCCAGGTAATTTCGCAGAATTTCTAATTACTACGGCATTTGCTAATTCATTTCTTACATGTTCATTACCGTAAACTGATGTATTGTCAGCGAAGGATATGTGATATAAGCCCGAATGTTCATCGCTGAACATGTCATCTCTCCCTAATTTAACTATTAATTTTAGTGGATCCGCTACGAATAGATATCTATCATTGACATGTATTAAATATTTTGACGAAAAGTACGGTGCACGTTCAAAATATTCTTGCTTTGCATGCAAATTAAACATCTCAGCTATTTTTTGGGATCTGTCTTCATATTCAAATTCGGCATTTAACACTACTAAGCTATCATCTCCACCAAAGACTCCACCGTAAGCGCAATTTCCGTAATCATATACATAAGCCAATACAAACATCGTTACTATAGTATTTCCAAGGAAAGTTAATGCATCACCACTTTTTCTCTGAAATTTCGTTAATAGCTTAATACCTTCGTTTCTAAACATTAATTCGGTACTACAATGAGTTTCCATCCATTCTTCTAGAAACCAATCCGGTGCACCAAACCTGCGGAATATTTCTAATTGCACACGTAACACTAATTCTACTTGACTTTTATCGAACTTACTGAAATCAACTTCATGGTATTTTACGGTTTTCAATCCATGTAATACATAATTTATATGTCCGTTTAATTCGTTTGGACTTAATCCTTCATTGATACGATATTTTCTTTTTAAGCTATGCTTAATTCTATGCATTAATACTAACATTATTCCTGACACTTTAGCGGTTAGAGCTGGGTTTTGAGCACCTATTACTTGACCTGCCATTAATTCATCATTATGCGTATTATCATATTTTGCTTTTAGATCTGCTTTTAAACTACCCGAGTAATTTACACCAGAGTAATTTAATTTCGAAAATTGTTTAAGTACATCTAATTGTGTGGTTGTTCTATTGCTTAACCAACCTTCAAAGAAAACACCGTTGTCGTATGAAAATATACACTCTGCATCCGTAAGTCTGTAATCATCTATATAAGTTGACATAAACACATCTGCGATTTCAATTTCTATATTTGGTATTTCCATGAGATGCAATTTCGGAGGTCTTGAATTACGTTTATCTATACTACTGATTAAATGCTTCAAATTACCCCTATATTTTTCAGGTTGAGCTGTTATCAGTATGGGTGATAGTCTATTAGATATCTTTTCGCGATATAATGCATTAAAGAATTTACTATGGTTAAATATAGTATATTCTTCAGACAATGGGAATACAGGTTCTAATTTCCTTTCTTCATATAGCATTTTATCAGGTCTATCATAAATATCATCTAATGCGCTTTGTACCGCAACTACATTCTTCGTTAAATCCAGACCTCCGTGAGTCTTTAAAGT

>Manitoba_iflavirus_1_36-2505

AATTGTGTAATCACCAGCAAAACACAAATGTGGTGCCATTAGTAAGGAATGCTGGCAAATATACCTACATTCTGAGGGTATCTCACTCATCTCCTGCATCTCAATGAATTGGCTCTTCAAGTCTTGATACCCTACTTGCGCATCAATTGGATAATAGGAAGTGGAAACGTAAGCATCCTTCAACCAATTATCCACGAAACCACCAATAGTGGGACCTTGTACACAAAATTTTGAACACAAAACAAGAAAACTGTACACTTGACGAATGTACCTTAAAAAGAACACGAATTGTTCTGTTGTCCCATCACTACTGTTCCATCCTGCTGCTAATAAAGAAGAAATCCATTTACTACCCTGCAAAGCATCATCCATGTAATTAGCCACAAAAGATCTAAAGACCCCAACTGTTTTAATCTGCAATTTACTGCCCATATGCATGGGACACATAAACTTCTCCACTGCTTCGGTCCTACACATCAATGTGGCACATGCCAAACACAGATAGTGGTTGCCATTTTCACACCATAAAGAAACACCACTGTGATGAGATAATGTTGAGCACCTATGGCACCGTACCGTCTCTCTCAAACCTTTGTTGAACCATGACTTAACTATCTGCACTATGTTGGTTACTGCTTCTTTCACATCTGTCAAGCCAAAGCCTTGCACCTGCATTTCATCGTCTTCGAGTGGTAACTTTGATAAAATCATCTGCCTATGCTTCATTTCATGGACGAGCATCGCCACTTCATCTTCCAACAATGTGCGGGAATCTCGCATCTCTACATTTGCCGTTTCGTTAGCTTTACGCAACAAATCAAAAGGATTGCCAATGTTCAACATATCTATAGCATTAGTTGTCATACATTGCATCAAAAATTCATACTTCAACTGTTGATTGGCTGCTTCTTTTTCCATGTGCGCACAGAATCGTACGGCCATTTCCTCAGCAAACTGTTTGAAGGAAAT

>Manitoba_iflavirus_1_45-15921

CTGGAAACATTTTTCTTGCAAACCTCATTCACTCTATATTCATTCTTCAAACAAACATACCGTAAACACACATCATTCAATCATACTACTCTAGAAACACACATTAATATCTAAAATCTATGAATTATAATTTCTGGAAACATCTTTCTTGCAATGTCAACGAAATCAATTTGAGGCACTACAACCTCACCTCTAAAAGCCTCTTTGAAAACTTTTTCCAGTTCTTTTTTCCAAGCCAAAAATTTACGTTCGCCATGACCAAATGCAAACATTAAACTACTCTCAGCATTTGAAAAACTAGCTTCTGCTGCAGTAAGGGGTTTGCGTCGCACCCACTTCACTAGTTCTTCAACAACCTCCCATCTCATTTGTGCACAATAAAAATTGTCCAACAATGGATGGGCCACAAATTTTCGTTTCAAGAATTCACAATCCTCAAAACCAAAGTAAGGTCGTACCTCTGCCCGTTTGTCTGCATTGGTAACAACCAAATTATGTCGAGCCAAGATTTCGCTAATAATCTTGGCATTAAACACCTCAGCGTATTCTTCTGCAACAGAATACACACCATCATCACCATATACTACAATGACAAAGTTTTGGCGGTAACTCTCAGGGGAAAACTCCAAGTTGTCCAAGTTTTGCCCCTCAATAAGAGTCACAACCTCAGGATAGCGATTAGAGAATCTACCTTTAACTTCATGTTGCAATTCATACACAATTTCTCCTACTATCAACCAACAAACTGCCATATACATCATGTGGACAAACGAATTTATCTCACCTGTTATACACGCGCCTGATGGACTACCACCAAATGTTTTATACAAAACGTTACCAGCTATATGATACGAACCCATAAGTTCCTTCATGAGCGCATTTGAAACATTTTCCATTTCATGACCAAACTCATACAACTGCGCAAAATCTTGCATCAACCGTGCAAAAGCCATTCCACAGTTATACTCAAGACTAGCCCCAAAATTGGAAAAATCAAGTGTAAAAAACTTAGGATGGGTTTTAAGCGTCGTAGCTAAGAAATTCCATTCCATTGATTCAGGGTTAATACCTATAGCTATACCATTACGAATGCGGTTAAAGCGCATGTGATTAACCAGACACAACGTATATCGTCGTAACGCAATAGTACCTTCAAAGTTTGACATCGAAAACAAGCGAGTTCCTCCCGGTTTCATTGCTTTCTCTATGGGTCGTCGCTCATCTTTTAAGCAATCCTGAAAAACATTGAAAGCAGCTTCACCCTTGTAGCGTAGTTCCATTTTCTTTCTATGA

>Manitoba_iflavirus_1_45-15922

TTCAACAACATCCCATCTCATCTGTGCACAATAAAAATTGTCCAACAATGGATGAGCCACAAACTTACGTTTCAAAAACTCGCAATCCTCAAAACCAAAGTAAGGTCGTACCTCTGCCCTTTTATCCGCATTGGTCACAACCAAATTATGTCGAGCCAAAATTTCACTAATAATCTTGGCATTAAACACTTCAGCATATTCTTCTGCAACAGAATACACACCATCATCACCATATACCACAATAACAAAATTCTGCCGGTAACTCTCAGGGGAAAACTCCAAGTTATCAAAATTGCGCCCCTCAATAAGAGTTACAACCTCAGGATAACGATTAGAAAATTTACCTTTAACTTCATGTTGCAGTTCATACACAATTTCTCCTACTATCAACCAACAAACTGCCATATACATCATGTGAACAAACGAGTTTATTTCACCTGTAATACATGCACCTGATGGACTACCACCAAACGTTTTATACAAAACATTGCCGGCGATATGATATGAACCCATAAGTTCCTTCATAAGCGCATTTGAAACATTTTCCATTTCATTACCAAACTCATACAGCTGCGCAAAGTCTTGCATCAAGCGTGCGAAAGCCATTCCACAGTTGTACTCAAGACTAGCCCCAAAATTAGAAAAATCAAGTGTAAAAAACTTAGGATGGGTTTTGAGTGTTGTAGCTAA

>Manitoba_iflavirus_1_5-8678

AGAAGAAATCCATTTGCTACCTTGCAAAGCATCATCCATATAGTTTGCTACAAAAGATCTAAAAACTCCAACAGTTTTAATCTGTAACTTGCTGCCCATATGCATAGGACACATGAACTTTTCNNNNNNNNNNNNNNNTTTCACACCATATCGAAACTCCACTGTGATGAGACAGCGTTGAACACCTATGGCACCGTACCGTCTCTCTCAAACCTTTGTTAAACCATGACTTAACAATTTGCACTATATTGGTTACTGCCTCTTTCACATCCGTCAAGCCAAAGCCTTGCGCCTGCATTTCATCTTCTTCAAGTGGCAACTTTGACAAGATCATTTGCCTATGTTTCATTTCATGCACGAGCATTGCCACTTCATCTTCTAACAGCGTGCGAGAATCGCGCATCTCTACATTTGCAGTTTCATTGGCTTTGCGCAACAAATCAAAGGGATTACCAATATTCAACATATCTATAGCATTTGTTGTCATGCATTGCATCAAGAACTCATATTTCAATTGCTGATTGGCTGCTTCTTTTTCCATATGTGCGCAGAACCGCACGGCCATTTCCTCAGCAAATTGTTTAAAGGAAATCGCATCGTGAGAAATAGACCCCTCATTCGTTACATCAGAATACATAAAAACTTCCAAGTGTTCAAAATTCTTTAAATCTGTTAAATCCGACTGACTTATTTTAACACCCTCCTTCAATACCAGTTTTGCCAATACGTGCCGACGCCTCCAAATTGCTTTGGGACATGGAATATCTGGATGGGTTACAAATGGATTATTAGTCGTCATTCCCATAACAATTGGTTGTCCCAATTGCTCT

>Manitoba_iflavirus_1_25-9622

CTCAAAAAGAAAACAAACTGTTCTGTTGTCCCATCGCTACTGTTCCATCCTGCTGCTAATAAAGAAGAAATCCATTTGCTACCTTGCAAAGCATCATCCATATAGTTTGCTACAAAAGATCTAAAAACTCCAACAGTTTTNNGCCAAACACAAGTAGTGATTGCCATTTTCACACCATATCGAAACTCCACTGTGATGAGACAGCGTTGAACACCTATGGCACCGTACCGTCTCTCTCAAACCTTTGTTAAACCATGACTTAACAATTTGCACTATATTGGTTACTGCCTCTTTCACATCCGTCAAGCCAAAGCCTTGCACCTGCATTTCATCTTCTTCAAGTGGCAACTTTGACAAGATCATTTGCCTATGTTTCATTTCATGCACGAGCATTGCCACTTCATCTTCTAACAGCGTGCGAGAATCGCGCATCTCTACATTTGCAGTTTCATTGGCTTTGCGCAACAAATCAAAGGGATTACCAATATTCAACATATCTATAGCATTTGTTGTCATGCATTGCATCAAGAACTCGTACTTCAATTGCTGGTTAGCTGCTTCTTTTTCCATATGTGCGCAAAACCGCACAGCCATTTCTTCAGCAAATTGTTTAAAGGAAATTGCATCGTGAGAAATAGACCCCTCATTCGTTACATCAGAATACATAAAAACTTCCAAGTGTTCAAAATTCTTTAAATCTGTTAAATCCGACTGACTTATTTTAACACCCTCCTTCAATACCAGTTTTGCCAATACGTGCCGACGCCTCCAAATTGCTTTGGGACATGGAATATCTGGATGGGTTACAAATGGATTATTAGTCGTCATTCCCATAACAATTGGTTGTCCCAATTGCTCTTTGTTTTCAAGTTCAGCACGTGGCACATTGTAAGGTGCTGACGTCATCACTCCATACAGAATACTGCACATACTACGTATAGTTTCAATGTCCTTAAGATTCCATATGTCATCAAACCACCAAGCAAT

>Manitoba_iflavirus_1_25-10201

AAGGAGTATGGCATCAAAGTTTGATACACGGGAACGTGTATTACAATTTTTGTCGACTGATTGGTGGTTGGAGTACAAATCGAAACATCCCATAGTGTCTCAATTGCTGTCCAAAGTTTTTCCAATTTTAGCAGTGTGTGGAGCTTTGTGGGTTGCAGGTAAGGCAGCTAATGCATTATGGGATTGGATAATGCAATATATCGGTTGGACTCCTAATGGCAAGAAGTATGATGAAGAAGAACAAAACAAAAGAGGAAGGAGAAAATTCTACCGAACACAAGGGTCTAATTTTTGCAATAAGCTTGACAAGATTTGTAGAAATTACGTGAGAATACGATGCCAAGGTAAGACAATTACGGCTTGGGGTTTAAAGGGATCAACTTTCTTGATGCCAAGCCATTTGAGCAATTTTGTTAATCATGAGGAAGGCTTTGAGATTTTGCTTATGCAGACAACAGATTACATTCATGTCGGTTTGGACAATCGAGAAGTTGTGGTGGTTCCTGGAAAAGATTTGATAATGGTAACAGTCAAGAAAGGGCGAATTTTGTTTGCAGATTGTTCAAAGTTTTT

>Manitoba_iflavirus_1_41-1425

GCCTCATTCACTCTATATTCATTCTTCAAACAAACATACCGTAAACACACATCATTCAATCATACTACTCTAGAAACACACATTAATATCTAAAATCTATGAATTATAATTTCTGGAAACATCTTTCTTGCAATGTCAACGAAATCAATTTGAGGCACTACAACCTCACCTCTAAAAGCTTCTTTAAAAACTTTTTCCAGTTCTTTTTTCCAAGCCAAGAATTTATGTTCACCATGACCAAATGCGAACATTAGACTACTCTCAGCATTTGAAAAACTAGCTTCTGCTGCAGTAAGGGGTCACAAACTTACGTTTCAAAAACTCGCAATCCTCAAAACCAAAGTAAGGTCGTACCTCTGCCCTTTTATCCGCATTGGTCACAACCAAATTATGTCGAGCCAAAATTTCACTAATAATCTTGGCATTAAACACCTCAGCATATTCTTCTGCAACAGAATACACACCATCATCACCATATACCACAATAACAAAATTCTGCCGGTAACTCTCAGGGGAAAACTCCAAGTTGTCAAAATTGCGCCCCTCAATANNNNNAATTTACCTTTAACTTCATGTTGCAGTTCATACACAATTTCTCCTACTATCAACCAACAAACTGCCATATACATCATGTGAACAAACGAATTTATTTCACCTGTAATACATGCACCTGATGGACTACCACCAAACGTTTTGTACAGAACATTGCCGGCGATATGATATGAACCCATAAGTTCCTTCATAAGCGCATTTGAAACATTTTCCATTTCATTACCAAACTCATACAGCTGCGCAAANNAAAGCCATTCCACAGTTGTACTCAAGACTAGCCCCAAAATTAGAAAAATCAAGTGTAAAAAACTTAGGATGGGTTTTGAGTGTTGTAGCTAAGAAATTCCATTCCATTGATTCAGGGTTTATACCTATAGCTATACCATTGCGAATGCGATTAAAGCGCATGTGATTAACCAAACACAACGTATATCGCCGTAGCGCAATAGTACCTTCAAAATTGGACATCGAAAACAAGCGAGTTCCTCCGGGTTTCATTGCTTTTTCTATGGGACGTCGCTCATCTTTCAAACAATCTTGAAAAACATTAAAGGCGGCTTCACCCTTGTAGCGCAGTTCCATTTTCTTTCTATGATCCTCCTTTACTATGTCTGAAAGGACAACGTTAAAAGGAAAACCATCAGCGCCATACTGCACATCAACCCATGGTGTTTTATTCTTTAGCCCTGTACCAATTTTCTTCATATACGCACTAGTACTATATGGCCATCCAGCACTTGTATC

>Manitoba_iflavirus_1_4-2392

CCTCATTCACTCTATATTCATTCTTCAAACAAACATACCGTAAACACACATCATTCAATCATACTACTCTAGAAACACACATTAATATCTAAAATCTATGAATTATAATTTCTGGAAACATCTTTCTTGCAATGTCAACGAAATCAATTTGAGGCACTACAACCTCACCTCTAAAAGCTTCTTTAAAAACTTTTTCCAGTTCTTTTTTCCAAGCCAAGAATTTATGTTCACCATGACCAAATGCGAACATTAGACTACTCTCAGCATTTGAAAAACTAGCTTCTGCTGCAGTAAGGGGCTTACGACGCACCCACTTAACTAGTTCTTCAACAACATCCCATCTCATCTGTGCACAATAAAAATTGTCCAACAATGGATGAGCAACAAACTTACGTTTCAAAAACTCGCAATCCTCAAAACCAAAGTAAGGTCGTACCTCTGCCCTTTTATCCGCATTGGTCACAACCAAATTATGTCGAGCCAAAATTTCACTAATAATCTTGGCATTAAACACCTCAGCATATTCTTCTGCAACAGAATACACACCATCATCACCATATACCACAATAACAAAATTCTGCCGGTAACTCTCAGGGGAAAACTCCAAGTTGTCAAAATTGCGCCCCTCAATAAGAGTTACAACCTCAGGATAACGATTAGAGAATTTACCTTTAACTTCATGTTGCAGTTCATACACAATTTCTCCTACTATCAACCAACAAACTGCCATATACATCATGTGAACAAACGAATTTATTTCACCTGTAATACATGCACCTGATGGACTACCACCAAACGTTTTGTACAGAACATTGCCGGCGATATGATATGAACCCATAAGTTCCTTCATAAGCGCATTTGAAACATTTTCCATTTCATTACCAAACTCATACAGCTGCGCAAAATCTTGCATCAACCGTGCAAAAGCCATTCCACAGTTGTACTCAAGACTAGCCCCAAAATTAGAAAAATCAAGTGTAAAAAACTTAGGATGGGTTTTGAGTGTTGTAGCTAAGAAATTCCATTCCATTGATTCAGGGTTTATACCTATAGCTATACCATTGCGAATGCGATTAAAGCGCATGTGATTAACCAAACACAACGTATATCGCCGTA

>Manitoba_iflavirus_1_40-6734

TCTTGATACCCTACTTGCGCATCAATTGGATAATAGGAAGTAGAGACGTAAGCATCCTTCAACCAATTGTCCACAAAACCACCAATAGTAGGACCTTGCACACAAAATTTGGAACACAAGACAAGAAAACTGTACACTTGACGAATGTATCTCAAAAAGAAAACAAACTGTTCTGTTGTCCCATCACTACTGTTCCATCCTGCTGCTAATAAAGAAGAAATCCATTTACTACCCTGCAAAGCATCATCCATATAGTTTGCTACAAAAGATCTAAAAACTCCAACAGTTTTAATCTGTAACTTACTGCCCATATGCATAGGACACATGAACTTTTCCACAGCTTCAGTCCTACACATCAATGTGGCACATGCCAAACACAAGTAGTGATTGCCACTTTCACACCATATCGAAACTCCACTGTGATGAGACAGCGTTGAACACCTATGGCACCGTACCGTTTCTCTCAAACCTTTGTTAAACCATGACCTAACAATTTGCACTATATTGGTTACTGCCTCTTTCACATCCGTCAAGCCAAAGCCTTGCGCCTGCATTTCATCTTCTTCAAGTGGCAACTTTGACAAGATCATTTGCCTATGTTTCATTTCATGCACAAGCATTGCCACTTCATCTTCTAACAGCGTGCGAGAATCGCGCATCTCTACATTTGCAGTTTCATTGGCTTTGCGCAACAAATCAAAGGGATTACCAATATTCAACATATCTATAGCATTTGTTGTCATGCATTGCATCAAGAACTCATATTTCAATTGCTGATTGGCTGCTTCTTTTTCCATATGTGCGCAGAACCGCACGGCCATTTCCTCAGCAAATTGTTTAAAGGAAATTGCATCGTGAGAAATAGACCCCTCATTCGTTACATCAGAATACATAAAAACTTCCAAGTGTTCAAAATTCTTTAAATCTGTTAAATCCGACTGACTTATTTTAACACCCTCCTTCAATACCAGTTTTGCCAAGACGTGCCGACGCCTCCAAATTGCTTTGGGACATGGAATATCTGGATGGGTTACAAATGGATTATTAGTCGTCATTCCCATAACAATTGGTTGTCCCAATTGCTCTTTGTTTTCAAGTTCAGCACGTGGCACATTGTAAGGTGCTGACGTCATCACTCCATACAGAATACTGCACATACTACGTATAGTTTCAATGTCCTTAAGATTCCATATGTCATCAAACCACCAAGCAATATGATCTGATCTCAAACCATTGAAGTAATCAAGAGCACAATTAATCGTAAAAAT

>Manitoba_iflavirus_1_1-1950

CCTCTAAAAGCCTCTTTGAAAACTTTTTCCAGTTCTTTTTTCCAAGCCAAAAATTTACGTTCACCATGACCAAATGCGAACATTAGACTACTCTCAGCATTTGAAAAACTAGCTTCTGCTGCAGTAAGGGGCTTACGACGCACCCACTTAACTAGTTCTTCAACAACATCCCATCTCATCTGTGCACAATAAAAATTGTCCAACAATGGATGAGCCACAAACTTACGTTTCAAAAACTCGCAATCCTCAAAACCAAAGTAAGGTCGTACCTCTGCCCTTTTATCCGCATTGGTCACAACCAAATTATGTCGAGCCAAAATTTCACTAATAATCTTGGCATTAAACACCTCAGCGTATTCTTCTGCAACAGAATACACACCATCATCACCATATACCACAATAACAAAATTCTGCCGGTAACTCTCAGGGGAAAACTCCAAGTTGTCCAAATTGCGCCCCTCAATAAGAGTTACAACCTCAGGATAACGATTAGAGAATTTACCTTTAACTTCATGTTGCAGTTCATACACAATTTCTCCTACTATCAACCAACAAACTGCCATATACATCATGTGAACAAACGAATTTATTTCACCTGTAATACATGCACCTGATGGACTACCACCAAATGTTTTATACAAAACGTTACCGGCTATATGATACGAACCCATAAGTTCCTTCATAAGCGCATTTGAAACATTTTCCATTTCATTACCAAACTCATACAGCTGCGCAAAATCTTGCATCAACCGTGCAAAAGCCATTCCACAGTTGTACTCAAGACTAGCCCCAAAATTAGAAAAATCAAGTGTAAAAAACT

>Manitoba_iflavirus_1_1-8332

CCTACTATCTGTGACAAAGGTCTCTTCTGCAAAAACTTTGAACAATCTGCAAACAAAATTCGCCCTTTCTTGACTGTTACCATTATCAAATCTTTTCCAGGAACCACCACAACTTCTCGATTGTCCAAACCGACATGAATGTAATCTGTTGTCTGCATAAGCAAAATCTCAAAGCCTTCCTCATGATTAACAAAATTGCTCAAATGGCTTGGCATCAAGAAAGTTGATCCCTTTAAACCCCAAGCCGTAATTGTCTTACCTTGGCACCGTATTCTCACGTAATTTCTACAAATCTTGTCAAGCTTATTGCAAAAATTAGACCCTTGTGTTCGGTAGAATTTTCTCCTTCCTCTTTTGTTTTGTTCTTCCTCATCATACTTCTTGCCATTAGGAGTCCAACCAATGTATTGCATTATCCAATCCCATAAAGCATTGGCTGCCTTACCTGCAACCCACAAAGCTCCACACACTGCTAAAATTGGAAAAACTTTGGACAGCAATTGAGACACTATGGGATGTTTCGATTTGTACTCCAACCACCAATCAGTCGACAAAAATTGTAATACACGTTCCCGTGTATCAAACTTTGATGCCATACTCCTTG

>Manitoba_iflavirus_1_27-788

TTCTGCTGCAGTAAGGGGTTTGCGTCGCACCCACTTCACTAGTTCTTCAACAACATCCCATCTCATTTGTGCACAATAAAAATTGTCCAACAATGGATGAGCCACAAATTTACGTTTCAAGAATTCACAATCCTCAAAACCAAAGTAAGGTCGTACCTCTGCCCGTTTATCTGCATTGGTAACAACCAAATTATGTCGAGCCAAGATTTCGCTAATAATCTTGGCATTAAACACCTCAGCGTATTCTTCCGCAACAGAATACACACCATCATCACCATATACTACAATGACAAAGTTTTGGCGGTAACTCTCAGGGGAAAACTCCAAGTTGTCCAAGTTAAGCCCCTCAATAAGAGTCACAACCTCAGGATAGCGATTAGAGAATCTACCTTTAACTTCATGTTGCAATTCATACACAATTTCTCCTACTATCAACCAACAAACTGCCATATACATCATGTGGACAAACGAATTTATCTCACCTGTTATACACGCGCCTGATGGACTACCACCAAATGTTTTATACAAAACGTTACCAGCTATATGATACGAACCCATAAGTTCCTTCATGAGCGCATTTGAAACGTTTTCCATTTCATGACCAAACTCATACAACTGCGCAAAATCTTGCATCAA

>Manitoba_narnavirus_1_21-2268

TTGCAGGACGTTCGGCACGTAATGTGATCGGTCCTTCGGGACGATTATCCGTTATAAACCGATGAAACCTCTTCCTATATTTAGGGAGGGTTTTCCAGATGTGAAAACTCTATCGAGAGATCCCATTTGGTAGTCTTCTGTCTCAGGGTGACTATCGTCACCCCTGTGACCGAAATCTGCAAAAGTCTCTAAATTAATATTTATAAACTTTTGTTCAACAGGGAAACTTCTTATTTTTAAGAAGTCTTCTTTGCTTAGGGGCGAGTCGCCATAGTTGTCGACACCCCGATTCTCCAATTCGTCCCATACATATTTGTATGTCTTGACAAATGGATTTGTATTAAATGAGCCTGTCTTTTCTGGTCCTTCAAGCAATAGCTTGGAGAACAGATCACCTCTTGTGCAGTATTCTGCAAAGTCGCGGTAGCTCATCCATCCCTTTTCTTTAGCTATTGCTAAAGTTAAGTGATTATTGTCCTTTTCGATCTGAAGCTCTTGCTTTAGATCTTTAAAGGATATACAACCAAAGTTCTCTCCCATTAGGGTTAGATCTTCTACGATTTGGTCTTCCATGGTTTTTAAGCATGAAATTCCTCGTCGACTCGTATTTGAGTTCAGGCGCCTAACAATCGTTAGGTCGTCCTGAATATCACC

>Manitoba_picorna-like_virus_1_20-2054

TCTATGATCCTCCTTTACTATGTCTGAAAGGACAACGTTAAAAGGAAAACCATCAGCGCCATACTGCACATCAACCCATGGTGTTTTATTCTTTAGCCCTGTACCAATTTTCTTCATATACGCACTAGTACTATATGGCCATCCAGCACTTGTATCTAGTGGAATATGGCCAAAGTAAACATCGCCAGTTTTCTCAATATCCAAGTTAAGACCACAGATGGACTCATCAACTCCATAAACCTTATATTCCACGCCAGGTATCTTATTACCACTGAGCAATAAACACTGGACCATTTCATAAGCTCGAGAAACAATTTTCTCAGGGAATGAAACGGCTAAGACACCATTCTTTCTAACACCATAATACAGTGGAGGAAACTCATGTGGGTAATCCTCTGTTTTCGACAAAAACGCTGGCATAGTAAGACAAGGACTCAAAAATGGAGCCAAAGGACTCTTGATAAGTTCAGTTTTAACAGGTATGAAACTAGCAAGTTCAGTCTCCACTTTTCCAAGTACTTGCAAAGCAACTCCTTCCCCATAATCTCGACTTATATTTTCCAACTCCAAATCTTCCAAGGACTCACCCTGGACCTGAAAATCATCCTGATACAAGACAACACCAATACCAGCATCTTGTGCCTGACTGCCACTTATATGCATCGACACTATAGGATGTATAACATCCATGCATAATAGAGAACCACACAACCCGGCCTTTTGGTACGAATACATTACTCCTCCAGTGTTCTCGTAAACTCTTCCCTTAGAGTCAGTGGCAGAAGTAGTATCCACAAGAGCGTCTATGTTAACTGCATAATCTACTACTCCCTCTTTGTCAACCTCCATCATTATTGCTTCATGTCCCACTATCTGCGACAAAGGTCTCTTTTGCAAAAACTTAGAACAATCCGCAAACAGAATTCGTCCCTTCTTAACCGTCACCATTATCAAATCCTTTCCAGGAACCACTACGACTTCACGATTATCCAAACCAACATGAATGTAGTCTGTTGTCTGCATAAGCAAAATCTCAAAGCCTTCCTCATGGTTAACAAAATTGCTCAAATGGCTTGGCATCAAGAAAGTTGATCCCTTTAAACCCCAAGCCGTAATTGTCTTACCCTGGCACCGTATCCTCACATAATTTTTACAAATTTTGTCAAGCTTGTTGCAAAAGTTTGACCCTTGAGTGCGATAGAATTTTCTTCTTCCTCTCTTGTTTTGTTCTTCTTCATCATACTTCTTGCCATTAGGAGTCCAACCGATATATTGCATTATCCAATCCCATAATGCATTAGCTGCCTTACCTGCAACCCACAACGCCCCACACACTGCCAAAATTGGNNAATTGAGACACTATGGGATGTTTTGACTTATACTCCAACCACCAATCAGTTGACAAAAACTGTAGCACACGTTCTCGGGTATCGAACTTTGATGCCATACTCCTTGGCCACAAGTATCTAGGCCAGTTTTGCCTAATACTTTCCTTGTTTCCACTTTCCAGCTCTTCTAACATCCCCTGGGAAGTTTGGAAACGCATTTTCAACACTTGCAGGATTTTGACATCTGCTAACACACACACTCCCTCACATAATGCATCCTTAATCTTCACTTCAGGGACACATGGAATTGTGTAATCACCAGCAAAACACAAATGTGGTGCCATTAGTAAGGAATGCTGGCAAATATACCTACATTCTGAGGGTATCTCACTCATCTC

>Manitoba_picorna-like_virus_1_29-1013

AATTAACACTTCTCGGATTTTGATAGCTCTCAATATAACAGCTAAATACTTAATACGTTTATCCCTTTTGTTAATCTCTGCTTCTGCAAAGAAAGTTAAGAATTCATTAGCATCCTTAGCAAATTCTTCCAAATCAGTACCTTGATTCTGTAATGCAGTTATTAAAGCGATATTGGGATCAGCTTCTTTAAAAGCCCAATCAACAATAGCTGAAACTCCTGCAAAAACAGTCCTAGTCAAGCTCAATATTGAATTAATGCCTTGTATCGATCGGTAATCGTGACCCCAGAAGATCTTCTTCAAATGTCCTTGTAGTCCTCCAGAGCTACGCACTTGTAATTTTCCACAAATAACAGCTGTAAGCATTTGCATAAGAGAGCTCCATGCACTGTCATTTGGCCCCTGGGTCACAAACTGTTGAGCTTGTGCATAAAACCCTTCAATTTTTGGACCTAAATCTTTCCAACGAGACATTGCAACGAGATCCAGCTCAACAAGTATGCCAAAAACTGCAAAACCAGCATTGGCCCAAGTACGAGCGATCGCTGCGTGTATCAGATGTTGAGCTACGCCCCACAAAGTCGTTAACATGCTTGAAGTGGCCTTCAATTGTGGGAAAAAGCCATTGACAATCTTACGTAATTCCTCCATCGCCATATTAGAGGATTCTTGAACAGACGCAGCAATTTTACCAGCTGCACTTCGAACCATAGTGTCTGAAACACCCATCAAATCACCAGCCACATTAGTTATAGCGTCTCTGACTACATGTGAAAGCGTACCTACTGCATGTTGACTGGCAGTATCAACTGCTTCAACAATATCCTGCGATATTTCAGTAAACGATGGTCCATCATATTGCATGCACCGCCGTACCTGTATGGCGTTGGCAACACCACCTGTTAGTGTATAAATTGCCACCGATCCTGCCAGACAAGCAACACTTGCTTTCGGTTCATCAGGAACTAAACTAGTTGCAATTGCTGTGAAAACACCAGCAGCTGCGAAATTGACAAAGGACTTCCACCCTGCAGAGGCTAGGTCCTTAAGACGAAAATCCAACTTCCCTTGACATTTCCAATTTTCAACAGGTTTGTTAAGTTCGTACATTAGCATATTATTTTCCCAACTACCAACAACTGTCTTATGTTGCTTAATTTCGTTGGAACACTCTGGACCTTGAGCCACGAAGGCAAGTTTGGTATCTTTGGCTTGAAGCTGCTTGACCCAACCTTGACGTCTATCATCGTCCACTGCAAAATAGGGATACGGATTGAAAAAACCAGGATGACCCAAAAATGCTCCTAACTCGAAACAATCAGGATTATTCTGGTAGATGGTAATTTTAAAACTTGAATCAGACCAAATGTATAATCTTCCATT

>Manitoba_picorna-like_virus_1_36-8618

TCAAATCCAGAACAATCTTGTACATTAATGATTTTCACATAAGAATTTTCCTTAGCGTCTCCACCAAGAGTCCTTGCATATGTTATTGAAATTTCCACTGAACCCTTATGAAGAAAAGTCTTAACAAATTGACCATATATAACCATAGTACCACGATAGTTAGTATATCCTGATGCTACCATTTGCAAGGGGGTAGTTATATCATTAATGGCCAAAGACCTATTGAAGACGTTTTCCTTGCCAGAATCCGTTGGATGTGCAGTCCAGGACCACAAGGTAGTACCAGCTGCATTAGTTGAACTCCAACTAAAAGTAGCCAATAACCCATTCATATTCATCATGTCCTTGTAACTCTTGGGTTCATCTACATACTGCTCCATGACTTCAGACAACTCAACCCAGTTTAATCCCATAACGCATGCAGAATCAATTCCTTCTCCCGTTGTAAAATTTAATCTAGGACGGGGAACAATTGCCATTTGGTGAACTGCATCATTAGGCTTGTCCTGATTTTTAACTTTTCCATTTCTCACTCCACCAATATTTTCAACTGTCTCTATTATAGATGACGCAGTATTGGCAGCTGTCTTGACACCATGTAGCACTGACAATGCAGGACCTTGAACTCGCCAAGTGGCTCTGTATGTAGCGCGCGCCTTGTCATTCTTGTTTTGTTCTGTAACAGTAGTGTAATCTTTCTTGTAATCCTTCTCTGGAACTACATACAACGCAGTGACAGTTCTGTCCGACTTCTTCGTAACAGGTTGCGTCATTGCAGTTACTTGCACATTCGAAAATCGATAAAAGATGCGTGCATTAAACGACGACGCAGTTCCAGTACCCACTTGAATTTTGGATAATGCATGTAATGTCAACATACCAGTAAAAGCTCCACGAACTCCAGGATTACCTCCTTCAACCGTGGTAGCATCAAAGGTACGTGCAAATGTGCGCTGATATTTCTGTGGAACGCGCAAAATTGCTTCACCACCCTGAGTAATGTCCATTATAACGTGCGGGCGTTGTATACTAGCCTCC

>Manitoba_picorna-like_virus_1_36-10843

GTCATGTGGAGCATACGCATAATAAACCGGTGCAGTTCCATCAACTACCAGAGTGTACATCTGACCACCTGCATACTGCCCGAAGCAAGAATTTATGAAAAACTGATGCGTTTGGGCCAAAGCACCAAGAAACTGGCCACTTGGAGCTAATTGCCGACTTAACCATGCAACAAACGTACGACTCAAAGGAGCAACAGGCATTTTCAAATAATTCTTAACTTTGACTTCAACCGCAGGTGTCACTCCAACTTTTGCTGTATACTTCACCGTTGTCGTAGGTTCATAACTAAAATTAGACACCACTTTAACAGGTTGCTTCAAAATAGATTTAAAATTTAAATGCTCAACTGTAGTTAAGATGTTCCCTGTATACTCCGGTTGTTCAAAATCCTCTCCTTGTGTTGAAAAAGCTACATCCTTCACCCATGTGCCAGAATCTGAGTATGGAACCATTCTGTCAGTCGCATTGGCTACATAATGATATTGCCTTGGGAATGCCCCTCCGCCAGTTACAGAATTAAGATGTGGCACTGGTAACATGTCCAAGCCATAATTGTCAACAGCCATAACACTATTAAACGGTCGAGGGTAATTACAACAAAAATCACTTCCACCTTTCAAATACAAAATCACTGGAACTGTATTAGATACAGCATCTGGAGCCAAAAGTTGA

>Manitoba_picorna-like_virus_1_45-1248

ACTTCACGATTATCCAAACCAACATGAATGTAGTCTGTTGTCTGCATAAGCAAAATCTCAAAGCCTTCCTCATGATTAACAAAATTGCTCAAATGGCTTGGCATCAAGAAAGTTGATCCCTTTAAACCCCAAGCCGTAATTGTCTTACCCTGGCACCGTATCCTCACATAATTTTTACAAATTTTGTCAAGCTTGTTGCAAAAGTTTGACCCTTGAGTGCGATAGAATTTTCTTCTTCCTCTCTTGTTTTGTTCTTCTTCGTCATACTTTTTGCCATTAGGAGTCCAACCGATATATTGCATTATCCAATCCCATAATGCATTAGCTGCCTTACCTGCAACCCACAACGCCCCACACACTGCCAAAATTGGAAACACCTTAGACAATAATTGAGACACTATGGGATGTTTTGACTTATACTCCAACCACCAATCAGTTGACAAAAACTGTAGCACACGTTCTCGGGTATCGAACTTTGATGCCATACTCCTTGGCCACAAGTATCTAGGCCAGTTTTGCCTAATACTTTCCTTGTTTCCACTTTCCAGCTCTTCTAACATCCCCTGGGAAGTTTGGAAACGCATTTTCAATACTTGCAGGATTTTGACATCTGCTAACACACACACTCCCTCACATAATGCATCCTTAATCTTCACTTCAGGGACACATGGAATTGTGTAATCACCAGCAAAACACAAATGTGGTGCCATTAGTAAGGAATGCTGGCAAATATACCTACATTCTGAGGGTATCTCACTCATCTCCTGCATCTCAATAAATTGACTCTTCAAGTCTTGATATCCTACTTGCGCATCAATTGGATAATAGGAAGTGGAAACGTAAGCATCCTTCAACCAATTATCCACGAAACCACCAATAGTGGGACCTTGTACACAAAATTTCGAGCACAAAACAAGAAAACTGTACACTTGACGAATGTACCTTAAAAAGAAAACGAATTGTTCTGTTGTCCCATCACTACTGTTCCATCCTGCTGCTAATAAAGAAGAAATCCATTTACTACCCTGCAGAGCATCATCCATGTAATTAGCCACAAAAGATCTAAA

>Manitoba_picorna-like_virus_1_45-1333

GATATATGGAATTTGAAAGATATTGAAACTATACGTAGCATGTGTAGTATTTTGTACGGAGTAATGACATCAGCACCTTACAACGTTCCGCGTGCTGAGCTTGAAAATAAAGAGCAATTGGGACAGCCAATAGTCATGGGGATGACAACCAATAATCCATTTGTGACTCACCCAGACATACCATGTCCCAAAGCAATTTGGAGGCGCCGGCATGTATTGGCGAAATTGGTGTTGAAGGAAGGTGTTAAGATAAGTCAGTCGGATTTGGCAGATTTGAAGAACTTTGAACACTTGGAGGTGTTCATGTATTCTGATGTTACAAATGAGTCTACTATTTCTCATGATGCGATTTCCTTCAAACAGTTTGCTGAGGAAATGGCCGTGCGATTCTGTGCTCACATGGAAAAAGAAGCAGCCAATCAACAGTTGAAGTATGAATTTTTGATGCAATGTATGACAACTAATGCTATAGATATGTTGAACATTGGCAATCCTTTTGATTTGTTGCGTAAAGCTAACGAAACGGCAAATGTAGAGATGCGAGATTCCCGCACATTGTTGGAAGATGAAGTGGCGATGCTCGTCCATGAAATGAAGCATAGGCAGATGATTTTATCAAAGTTACCACTCGAAGACGATGAAATGCAGGTGCAAGGCTTTGGCTTGACAGATGTGAAAGAAGCAGTAACCAACATAGTGCAGATAGTTAAGTCATGGTT

>Manitoba_picorna-like_virus_1_5-1019

CCTTGTGTTCGGTAGAATTTTCTCCTTCCTCTTTTGTTTTGTTCTTCCTCATCATACTTCTTGCCATTAGGAGTCCAACCAATGTATTGCATTATCCAATCCCATAAAGCATTGGCTGCCTTACCTGCAACCCACAAAGCTCCACACACTGCTAAAATTGGAAAAACTTTGGACAGCAATTGAGACACTATGGGATGTTTCGATTTGTACTCCAACCACCAATCAGTCGACAAAAATTGTAATACACGTTCCCGTGTATCAAACTTTGATGCCATACTCCTTGGCCACAAGTATCTAGGCCAGTTTTGCCTAACACTTTCCTTGTTTCCACTTTCCAACTCTTCTAGCATTCCCTGGGAAGTTTGAAAACGCATCTTCAACACTTGAAGGATTTTAACATCCGCTAGCACACACACTCCTTCACACAACGCATCCTTAATCTTCACCTCAGGAACACATGGAATTGTGTAGTCACCAGCAAAACACAAATGTGGCGCCATTAACAAAGAGTGTTGGCAAATATACCTACATTCTGAAGGTATCTCACCCATTTCCTGCATCTCAATGAATTGGCTCTTCAAATCTTGATACCCTACTTGCGCATCAATTGGATAATAGGAAGTAGAAACGTAAGC

>Manitoba_picorna-like_virus_1_5-1020

GTAATTGTCTTACCCTGGCACCGTATCCTCACATAATTTTTACAAATTTTGTCAAGCTTGTTGCAAAAGTTTGACCCTTGAGTGCGATAGAATTTTCTTCTTCCTCTCTTGTTTTGTTCTTCTTCATCATACTTCTTGCCATTAGGAGTCCAACCGATATATTGCATTATCCAGTCCCATAATGCATTAGCTGCCTTACCTGCAACCCACAATGCCCCACACACTGCCAAGATTGGGAACACCTTAGACAACAACTGAGACACTATGGGATGTTTTGACTTATACTCCAACCACCAATCAGTTGACAAAAACTGTAGCACACGTTCTCGGGTATCGAACTTTGAAGCCATACTCCTTGGCCACAAGTATCTAGGCCAGTTTTGCCTAATACTTTCCTTGTTTCCACTTTCCAACTCTTCTAACATCCCCTGGGAAGTTTGGAAACGCATTTTCAATACTTGCAGGATTTTGACATCTGCTAACACACACACTCCCTCACATAATGCATCCTTAATCTTCACTTCAGGGACACATGGAATTGTGTAATCACCAGCAAAACACAAATGTGGTGCCATTAGTAAGGAATGCTGGCAAATATACCTACATTCTGAGGGTATCTCACTCATCTCCTGCATCTCAATGAATTGACTCTTCAAGTCTTGA

>Manitoba_picorna-like_virus_1_5-2690

ATGTCCTTGTAGTCCTCCAGAGCTACGCACTTGTAATTTTCCACAAATAACAGCTGTAAGCATTTGCATAAGAGAGCTCCATGCACTGTCATTTGGCCCCTGGGTCACAAACTGTTGAGCTTGTGCATAAAACCCTTCAATTTTTGGACCTAAATCTTTCCAACGAGACATTGCAACGAGATCCAACTCAACAAGTATGCCAAAAACTGCAAAACCAGCATTGGCCCAAGTACGAGCTATCGCTGCGTGTATCAGATGTTGTGCAACGCCCCACAAAGTCGTTAGCATGCTAGAGGTAGCTTTCAATTGTGGGAAAAAGCCATTGACAATCTTACGTAATTCCTCCATCGCCATATTAGAGGATTCTTGAACAGACGCAGCAATTTTGCCAGCTGCACTTCGAACCATGGTGTCTGAAACACCCATCAAATCACCAGCTACATTAGTTATAGCGTCTCTGACCACATGTGAAAGCGAACCTACTGCACGCTGACTGGCAGTATCCACTGCTTCAACAATATCCTGCGATATTTCAGTAAACGATGGTCCATCATATTGCATGCACCGCCGTACCTGTATGGCATTAGCAACACCACCTGTTAGTGTATAAATTGCCACTGATCCTGCCAGACAAGCAACACTTGCTTTCGGTTCATCAGGAACTAAACTAGTTGCAATTGCTGTAAAAACACCAGCAGCTGCGAAGTTGACAAAGGACTTCCACCCTGCAGAGGCTAGGTCCTTAAGACGAAAATCCAACTTCCCTTGGCATTTCCAATTTTCTACAGGTTTATTAAGTTCGTACATTAGCATA

>Manitoba_picorna-like_virus_1_5-2972

AACTGACCGCTTGGAGCCAACTGGCGACTTAACCATGCAACAAACGTACGACTCAAAGGAGCAACAGGCATTTTCAAATAATTTTTAACTTTAACCTCAACTGTAGGTGTGACTCCAACTTTAGCAGTGTATTTCACTGTTGTTGTAGGCTCATAACTAAAATTAGACACTACTTTAACAGGCTGTTTCAAAATAGACTTAAAATTCAAATGCTCAACTGTAGTTAGGATATTTCCTGTGTACTCTGGTTGTTCAAAGTCTTCTCCTTGCGTTGAAAAAGCTACGTCCTTCACCCATGTTCCAACATCTGAGTATGGAACCATTCTATCAGTTGAATTAGCAACGTAATGATATTGCCTTGGGAATGCTCCTCCTCCAGTTACAGAATTAAGATGCGGTACTGGCAACATATCCAAACCATAATTGTCAACTGCCATTACACTATTAAATGGTCGAGGATAATTGCAACAAAAATCACTACCACCTTTCAAGTACAGAATAACTGGAACCGTATTAGATACCGCATCCGGAGCCAGAAGTTGATTCAAAACTCTAATTGTAACAGCTGTACTA

>Manitoba_picorna-like_virus_1_41-2716

TCGTTGGAACACTCTGGACCTTGAGCCACGAAGGCAAGTTTGGTATCTTTGGCTTGAAGCTGCTTGACCCAACCTTGACGTCTATCATCGTCCACTGCAAAATAGGGATACGGATTGAAAAAACCAGGATGACCCAAAAATGCTCCTAACTCGAAACAATCAGGATTATTCTGGTAGATGGTAATTTTAAAACTTGAATCAGACCAAATGTATAATCTTCCATTGTACCATTCACTATTCTCTCGTGAGGAAGCCAGTGTAAAGTCGCCTGTGCTTGTTGCGACATTACGATTCATCAAAATCCATGGCAATGGAATTGACTGATTAATCTCAATCTTTTCAGTTGGATTTACACTAGGCACAATTATACCGTTAAACAAACCACACGAAGCTAAATCCTCAGAGCCATTGATAGCACCTCCTAATGTTGGTGGTACAATGTTAGCAGTTTTGTCTGGAAAGCCTGATGACAAGATACCACCAGCTCCTCGATACTGCCATTCAGCTAAGGGGCCGTAAAATTTATTAACTCCAGACTTGTCATGTGGAGCATACGCATAGTAAACCGGTGCAGTTCCATCCACTACTAGAGTGTACATCTGACCACCTGCATACTGCCCGAAGCAAGAATTGATGAAAAACTGATGCGTCTGGGCCAAAGCACCAAGAAACTGGCCACTTG

>Manitoba_picorna-like_virus_1_35-4471

AATTATAATTTCTGGAAACATTTTTCTTGCAATGTCAATAAAATCAATTTGAGGCACTACAACCTCTCCTCTAAAAGCCTCTTTGAAAACTTTTTCCAGTTCTTTTTTCCAAGCCAAAAATTTACGTTCGCCATGACCAAATGCAAACATTAAACTACTCTCAGCATTTGAAAAACTAGCTTCTGCTGCAGTAAGGGGTTTGCGTCGCACCCACTTAACTAGTTCTTCAACAACTTCCCATCTCATTTGTGCACAATAAAAATTGTCCAACAATGGATGGGCCACAAATTTACGTTTCAAGAATTCACAATCCTCAAAACCAAAGTAAGGTCGTACCTCTGCCCGTTTATCTGCATTGGTAACAACCAAATTATGTCGAGCCAAGATTTCACTAATAATCTTGGCATTAAACACCTCAGCGTATTCTTCTGCAACAGAATACACACCATCATCACCATATACTACAATGACAAAGTTTTGGCGGTAACTCTCAGGGGAAAACTCCAAGTTGTCCAAGTTAAGCCCCTCAATAAGAGTCACAACCTCAGGATAGCGATTAGAGAATCTACCTTTAACTTCATGTTGCAATTCATACACAATTTCTCCTACTATCAACCAACAAACTGCCATATACATCATGTGGACAAACGAATTTATCTCACCTGTTATACACGCGCCTGATGGACTACCACCAAATGTTTTATACAAAACGTTACCAGCTATATGATACGAACCCATAAGTTCCTTCATGAGCGCATTTGAAACATTTTCCATTTCATGACCAAACTCATACAACTGCGCAAAATCTTGCATCAACCGTGCAAAAGCCATTCCACAGTTATACTCAAGACTAGCCCCAAAATTGGAAAAATCAAGTGTAAAAAACTTAGGATGGGTTTTAAGCGTCGTAGCTAAGAAATTCCATTCCATTGATTCAGGGTTAATACCTATAGCTATACCATTACGAATGCGGTTAAAGCGCATGTGATTAACCAGACACAACGTATATCGTCGTAACGCAATAGTACCTTCAAAGTTTGACATCGAAAACAAGCGAGTTCCTCCCGGTTTCATTGCTTTCTCTATGGGTCGTCGCTCGTCTTTTAAGCAATCCTGAAAAACATTGAAAGCAGCTTCACCCTTGTAGCGTAGTTCCATTTTCTTTCTATGATCCTCCTTTACTATATCTGAGAGGACTACGTTAAAAGGAAAACCATCAGCGCCATACTGCACGTCAACCCATGGAGTTTTATTCTTTAGCCCTGTACCAATCTTCTTCATATACGCACTGGTACTATACGGCCATCCAGCACTTGTATCAAGTGGAATATGGCCAAAATAGACATCACCAGTCTTCTCAATATCCAAGTTAAGACCACAAATAGACTCATCAACCTCGTAAACCTTATACTCCACACCGGGTATCTTATTTCCACTAAGCAACAAACACTGAACCATTTCATAAGCTCGAGAAACAATTTTCTCAGGGAACGAAACTGCTAAGACACCATTCTTTCTAACTCCATAATACAATGGAGGAAATTCATGCGGGTAATCCTCCGTCTTCGACAAAAACGCTGGCATAGTGAGACATGGGCTCAAAAATGGAGCCAAAGGACTCTTTATAAGCTCAGTTTTCACGGGGATAAAACTGGAAAGTTCTGTCTCCACCTTTCCAAGTACTTGCAAAGCAACACCTTCCCCATAATCTCGACTTATGTTTTCAAGCTCCAAGTCTTCCAAAGACTCTCCCTGAACCTGAAAATCATCCTGATACAAGACAACACCAATACCAGCATCTTGCGCTTGACTACCACTTATATGCATAGACACTATAGGATGTATAACATCCATGCATAATAAGGAGCCACACAATCCAGCCTTCTGGTAGGAATACATTAC

>Manitoba_picorna-like_virus_1_4-1091

TTCCCAAGGCAATATCATTATGTAGCCAATGCAACTGACAGAATGGTTCCATACTCAGATTCTGGTACATGGGTGAAGGATGTAGCTTTTTCAACACAAGGAGAGGACTTTGAACAACCAGAATATACAGGGAACATCTTAACCACAGTTGAGCATTTAAATTTTAAATCTATTTTGAAGCAACCTGTTAAAGTGGTGTCTAATTTTAGTTATGAACCCACAACAACAGTGAAGTATACTGCAAAAGTTGGAGTGACACCTGCAGTTGAAGTCAAAGTTAAGAATTATTTGAAAATGCCTGTTGCTCCTTTGAGTCGTACGTTTGTTGCATGGTTAAGTCGGCAGTTAGCTCCAAGTGGCCAGTTTCTTGGTGCTTTGGCCCAGACGCATCAGTTTTTCATCAATTCTTGCTTCGGGCAGTATGCAGGTGGTCAGATGTACACTCTAGTAGTGGATGGAACTGCACCGGTTTACTATGCGTATGCTCCACATGACAAGTCTGGAGTTAATAAATTTTACGGCCCCTTAGCTGAATGGCAGTATCGAGGAGCTGGTGGTATCTTGTCATCAGGCTTTCCAGACAAAACTGCTAACATTGTACCACCAACATTAGGAGGTGCTATCAATGGCTCTGAGGATTTAGCTTCGTGTGGTTTGTTTAACGGTATAATTGTGCCTAGTGTAAATCCAACTGAAAAGATTGAGATTAATCAGTCAATTCCATTGCCATGGATTTTGATGAATCGTAATGTCGCAACAAGCACAGGCGACTTTACACTGGCTTCCTCACGAGAGAATAGTGAATGGTACAATGGAAGATTATACATTTGGTCTGATTCAAGTTTTAAAATTACCATCTACCAGAATAATCCTGATTGTTTCGAGTTAGGAGCATTTTTGGGTCATCCTGGTTTTTTCAATCCGTATCCCTATTTTGCAGTGGACGATGATAGACGTCAAGGTTGGGTCAAGCAGCTTCAAGCCAAAGATACCAAACTTGCCTTCGTGGCTCAAGGTCCAGAGTGTTCCAACGAAATTAAGCAACATAAGACAGTTGTTGGTAGTTGGGAAAATAATATGCTAATGTACGAACTTAACAAACCTGTTGAAAATTGGAAATGTCAAGGGAAGTTGGATTTTCGTCTTAAGGACCTAGCCTCTGCAGGGTGGAAGTCCTTTGTCAATTTCGCAGCTGCTGGTGTTTTCACAGCAATTGCAACTAGTTTAGTTCCTGATGAACCGAAAGCAAGTGTTGCTTGTCTGGCAGGATCGGTGGCAATTTA

>Manitoba_picorna-like_virus_1_4-8285

TCTCTCAAACCTTTGTTAAACCATGACTTAACAATTTGCACTATATTGGTTACTGCCTCTTTCACATCCGTCAAGCCAAAGCCTTGCGCCTGCATTTCATCTTCTTCAAGTGGCAACTTTGACAAGATCATTTGCCTATGTTTCATTTCATGCACGAGCATTGCCACTTCATCTTCCAACAGCGTGCGAGAATCGCGCATCTCCACATTTGCAGTTTCATTGGCTTTGCGCAACAAATCAAAGGGATTACCAATATTCAACATATCTATAGCATTTGTTGTCATGCATTGCATCAAGAACTCGTACTTCAATTGCTGGTTAGCTGCTTCCTTTTCCATATGTGCGCAAAACCGCACAGCCATTTCCTCAGCAAATTGTTTAAAGGAAATTGCATCGTGAGAAATAGACCCCTCATTCGTTACATCAGAATACATAAAAACTTCCAAGTGTTCAAAATTCTTTAAATCTGTTAAATCCGACTGGCTTATTTTAACACCCTCCTTCAACACCAGTTTTGCCAACACGTGCCGACGCCTCCAAATTGCTTTAGGACATGGAATGTCTGGATGGGTTACAAATGGATTATTAGTCGTCATTCCCATAACAATTGGTTGTCCCAATTGCTCTTTGTTTTCAAGTTCAGCACGTGGCACATTGTAAGGTGCTGACGTCAT

>Manitoba_picorna-like_virus_1_4-8382

CATAAACCTTATACTCCACACCAGGTATCTTATTTCCACTAAGCAACAAACACTGGACCATTTCATAAGCTCGAGAAACAATTTTCTCAGGGAATGAAACGGCTAAGACACCATTCTTTCTAACACCATAATACAGTGGAGGAAACTCATGTGGGTAATCCTCTGTTTTCGACAAAAACGCTGGCATAGTAAGACAAGGACTCAAAAATGGAGCCAAAGGACTCTTTATAAGCTCAGTTTTCACGGGGATAAAACTGGAAAGTTCTGTCTCCACTTTTCCAAGTACTTGCAAAGCAACTCCTTCCCCATAATCTCGACTTATGTTTTCAAGCTCCAAGTCTTCCAAAGACTCTCCCTGAACCTGAAAATCGTCCTGATACAAGACTACACCAATACCAGCATCTTGCGCTTGACTACCACTTATATGCATAGACACTATAGGATGTATAACATCCATGCATAATAAGGAGCCACACAATCCAGCCTTCTGGTAGGAATACATTACCCCTCCAGTATTCTCATAGACTCTTCCTTTAGCATCAGTAGCCGAAGTAGTGTCCACAAGAGCATCTATGTGAACTGCGTAATCCACTACACCTTCTTTGTCAACTTCCATCATTATTGCTTCTTGTCCTACTATCTGAGACAAGGGTCTCTTTTGCAAAAACTTTGAACAATCTGCAAACAAAATTCGCCCTTTCTTGACTGTTACCATTATCAAATCTTTTCCAGGAACCACCACAACTTCTCGATTGTCCAAACCGACATGAATGTAATCTGTTGTCTGCATGAGCAATATCTCAAAGCCATCCTCATGATTAACAAAATTGCT

>Manitoba_picorna-like_virus_1_33-6083

GTATCCACAAGAGCGTCTATGTTAACTGCATAATCTACTACTCCCTCTTTGTCAACCTCCATCATTATTGCTTCATGTCCCACTATCTGCGACAAAGGTCTCTTTTGCAAAAACTTAGAACAATCCGCAAACAGAATTCGTCCCTTCTTAACCGTCACCATTATCAAATCCTTTCCAGGAACCACTACGACTTCACGATTATCCAAACCAACATGAATGTAGTCTGTTGTCTGCATAAGCAAAATCTCAAAGCCTTCCTCATGATTAACAAAATTGCTCAAATGGCTTGGCATCAAGAAAGTTGATCCCTTTAAACCCCAAGCCGTAATTGTCTTACCCTGGCACCGTATCCTCACATAATTTTTACAAATTTTGTCAAGCTTGTTGCAAAAGTTTGACCCTTGAGTGCGATAGAATTTTCTTCTTCCTCTCTTGTTTTGTTCTTCTTCATCATACTTCTTGCCATTAGGAGTCCAACCGATATATTGCATTATCCAATCCCATAATGCATTAGCTGCCTTACCTGCAACCCACAACGCCCCACACACTGCCAAAATTGGGAACACCTTAGACAGTAATTGAGACACTATGGGATGTTTTGACTTATACTCCAACCACCAATCAGTTGACAAAAACTGTAGCACACGTTCTCGGGTATCGAACTTTGATGCCATACTCCTTGGCCACAAGTATCTAGGCCAGTTTTGCCTAATACTTTCCTTGTTTCCACTTTCCAGCTCTTCTAACATCCCCTGGGAAGTTTGGAAACGCATTTTCAACACTTGCAGGATTTTGACATCTGCTAACACACACACTCCCTCACATAATGCATCCTTAATCTTCACTTCAGGGACACATGGAATTGTGTAATCACCAGCAAAACACAAATGTGGTGCCATTAGTAAGGA

>Manitoba_picorna-like_virus_1_3-11786

TCTTCTCCTTGCGTTGAAAAAGCTACGTCCTTCACCCATGTTCCAACATCTGAGTATGGAACCATTCTATCAGTTGAATTGGCAACGTAATGATATTGCCTCGGGAACGCTCCTCCTCCAGTTACAGAATTAAGATGTGGTACTGGCAACATGTCCAAACCATAATTGTCAACTGCCATTACACTATTAAATGGTCGAGGATAATTGCAACAAAAATCACTACCACCTTTCAAGTACAGAATAACTGGGACCGTATTAGATACCGCATCTGGAGCCAGAAGTTGATTCAAAACTCTAATTGTAACAGCTGTACTATGAAACATAGTAGCGGTAGACGGCCATGTAACATTAGGCAACAAGAATGGTGTCGATGTACCATCCAAAGTGCGCATTGGTGTGTCAAATATGTACGGAATAGTTATCTCAAATCCGGAACAATCTTGTACATTAATGATTTTCACGTAAGAATTTTCCTTAGCATCTCCACCAAGAGTTCTTGCATATGTTATTGAAATTTCCACTGAACCTTTGTGAAGGAAAGTTTTAACAAATTGACCATATATAACCATAGTACCACGATAATTAGTATATCCTGATGCCACCATTTGCAAAGGCGTTGTTATATCATTAATGGCCAACGATCTATTGAAGACATTTTCTTTACCAGAATCCGTTGGATGTGCAGTCCAGGACCATAGAGTGGTACCAGCCGCATTGGTGGAGCTCCAACTAAAGGTAGCCAATAAACCATTCATATTCATCATGTCCTTGTAACTCTTGGGTTCATCTACATACTGCTCCATGACTTCAGACAACTCAACCCAGTTTAATCCCAT

>Manitoba_picorna-like_virus_1_6-7570

GTCATGTGGAGCATACGCATAGTAAACCGGTGCAGTTCCATCCACTACTAGAGTGTACATCTGACCACCTGCATACTGCCCGAAGCAAGAATTGATGAAAAACTGATGCGTCTGGGCCAAAGCACCAAGAAACTGGCCACTTGGAGCTAACTGCCGACTTAACCATGCAACAAACGTACGACTCAAAGGAGCAACAGGCATTTTCAAATAATTCTTAACTTTGACTTCAACTGCAGGTGTCACTCCAACTTTTGCAGTATACTTCACTGTTGTTGTGGGTTCATAACTAAAATTAGACACCACTTTAACAGGTTGCTTCAAAATAGATTTAAAATTTAAATGCTCAACTGTGGTTAAGATGTTCCCTGTATATTCTGGTTGTTCAAAGTCCTCTCCTTGTGTTGAAAAAGCTACATCCTTCACCCATGTACCAGAATCTGAGTATGGAACCATTCTGTCAGTTGCATTGGCTACATAATGATATTGCCTTGGGAATGCCCCTCCGCCAGTTACAGAATTAAGATGTGGCACTGGTAACATGTCCAAGCCATAATTGTCAACAGCCATAACACTATTAAACGGTCGAGGGTAATTGCAACAAAAATCACTTCCACCTTTCAAATACAAAATCACTGGAACTGTATTAGATACAGCATCTGGAGCCAAAAGTTGA

>Manitoba_picorna-like_virus_1_2-5166

CAAGCCAAAGCCTTGCACCTGCATTTCATCGTCTTCGAGTGGTAACTTTGATAAAATCATCTGCCTATGCTTCATTTCATGGACGAGCATCGCCACTTCATCTTCCAACAATGTGCGGGAATCTCGCATCTCTACATTTGCCGTTTCGTTAGCTTTACGCAACAAATCAAAAGGATTGCCAATGTTCAACATATCTATAGCATTAGTTGTCATACATTGCATCAAAAATTCATACTTCAACTGTTGATTGGCTGCTTCTTTTTCCATGTGCGCACAGAATCGTACGGCCATTTCCTCAGCAAACTGTTTGAAGGAAATCGCATCATGAGAAATAGTAGACTCATTTGTAACATCAGAATACATGAACACCTCCAAATGTTCAAAGTTCTTCAAATCTGCCAAATCCGACTGACTTATCTTAACACCTTCCTTCAATACCAATTTCGCCAATACATGCCGGCGCCTCCAAATTGCTTTGGGACATGGTATGTCTGGGTGAGTCACAAATGGATTATTGGTTGTCATTCCCATGACTATTGGCTGTCCCAATTGCTCTTTATTTTCAAGCTCAGCACGTGGAACGTTGTAAGGTGCTGATGTCATTACTCCGTACAAAATACTACACATGCTACGTATAGTTTCAATATCTTTCAAATTCCATATATCATCAAACCACCAAGCAATGTGATCTGATCTCAAACCATTGAAGTAATCAAGAGCACAATTAATCGTAAAAATATAATCCGGCGTATTAAGCGTAAACTTGGCTTTTTTAAG

>Manitoba_rhabdovirus_1_20-4126

GTGTTAATGTTTTTTTCACGGTCAAGGCTGGTAAAATTCTGTTTCTTGATTGCTTGAAGTCTTTGTAAACAATCTTATTAACTGATACCTTTTATTCTTAAAACAGTGGAGGAGTTTTGACCAAAAGATTAGAATCAAAACGCCGATTAGAACTGTTACAGCGCATGCTGTTAATGTATATAACATAGGTGTCCAGTGCAAATCTGTCAAAGAGCTATCATCCACGAAATTATAATCTGTCTCTGTACCGGGGTGTAGCTGTCTTCTCCGGTATTCGGGAGACCGCTTTGCCATCGGTACTCTGTAGTGAGAGCTTAGTCGTGCGATTCTTTTATAGTTTTCTATGTCTTGGATGCCAAGGATCAGTGTCTTGTTCTTGACATACAACCCATTGGGTCCATCGATCACATCTGTTTTATTTGATGTAACCCAGTACGGCCATGTCCAAAGTTGGTTGTCTGATCTTCTTATACTAATATACGGAAAACTCAGGCTCGGTTCGATAGTCACTTGTTGGTATAAGCTTAGTCCCATCTCAAATTTCCCTGGACTATAGCGATAGACTGGATGAAATCCGGGATAACGAGGTGTCAATGTTTGCAACTCCGCTCGACTAACCATTTCTCCTGCTATGATTTTATCGACAACTATCTCACACTCATGATCTAAAAATTCATCCAACAAACTGAGGACAGCGTTCTGTATAAATTCTTTGTCATTAATTATGTTGACTTGTGACCCAATCGGGCAGTCCGGAACTTGGTAAAAGTACTCCCCTATCCAGTCCTGGTGCGGGATGTTTTTCCGGTCCAGTGCCATCCAGTCCCCAGAGGGATATCTCAACCCTCGGTGGCCGCAGAAATTCATTGTACATGGTTTGTCATACTCGGATATGTATATATCAGGACTCCAACTAGCTACTGTATGATTGGGAGTGTAATATATCACTAAGGATCCATCCAACATATGAGAATCATCACATGAATCTTTTATCGAGTGATCCGATATCCAAAGACGATTGTAAAAATGTGTGATACAAACAGAGGACGAACAGGTTCCGTCCAAGAAGATCGAGGATTTGAAGGCATCAGAATACGGATTGTACGAGACAGAGTGAGAACTCACTGAGATGAAGATGCGAGAGGTGGTCGAGCTCTTCATCCAGGAGCATGAAGGCGTTGGGTGCTCAGAAGATGAGAATGCTCCCTCTGATACTTTCTGGATCTCCTCTCTACATAAAATCTCCGAAACCTCCGATGGTTCAGTGTGTAACTCCAAAGTGATACCTCCAAAGAAACCCTTAGAACAGGTTGTACTCAATTTTGTGAAGGTGCATATGGATCCAGCAATATTAATGGTTGTCCCTATCAAGGGTAATCTTATTGGAATTGTTATGCCTGACTCAATTTTTGCATAACTTTCCCCAATGGGACATGTTAACGCACTCGGTGTCACATCGTGCCACTGAACATTGTCAGATATTGGAAATAGCATTGTATGTTGTTGTCCTCGTAATGAGTCTCCTAATGTACCCTCCGTCATGGCGACTATTTGTACAGCCACATAGATAAGACTCTTGATCTTCTGGCGAGAGCAATCCATTATTATGACGGGTCATAGCATAGTTGACTTGTGTTAGTTTTTTTCTTGAATAGCTCATTCGAACTCTGAGACTCAGACAAGAGGCATCCCAAACGGACTCAACCGCCACCACAAGAACAGCTTCTAGAGCTCGTTATCTCCTATTACCATATCCTATATAAATACAAAATATAAGCCTATTAGAATTAATCATAAAATTTGATATAGAAAATATCTAGAAATATACGGTGATGATTTTGAAGCACGCTAGTCTGCAATA

>Manitoba_rhabdovirus_1_36-4251

GGAGGAGTTTTGACCAAAAGATTAGAATCAAAACGCCGATTAGAACTGTTACAGCGCATGCTGTTAATGTATATAACATAGGTGTCCAGTGCAAATCTGTCAAAGAGCTATCATCCACGAAATTATAATCTGTCTCTGTACCGGGGTGTAGCTGTCTTCTCCGGTATTCGGGAGACCGCTTCGCCATCGGTACTCTGTAGTGAGAGCTTAGCCGTGCGATTCTTTTATAGTTTTCTATGTCTTGGATGCCAAGGATCAGTGTCTTGTTCTTGACATACAACCCATTGGGTCCGTCGATCACATCTGTTTTATTTGATGTAACCCAGTACGGCCATGTCCAAAGTTGGTTGTCTGATCTTCTTATACTAATATACGGAAAACTCAGGCTCGGTTCGATAGTCACTTGTTGGTATAAGCTTAGTCCCATCTCAAATTTTCCTGGACTATAGCGATAGACTGGATGAAATCCGGGATAACGAGGTGTCAATGTTTGCAACTCCGCTCGACTAACCATTTCTCCTGCTATGATTTTATCGACAACTATCTCACACTCATGATCTAAAAATTCATCCAACAAGCTGAGGACAGCGTTCTGTATAAATTCTTTGTCATTAATTATGTTGACTTGTGACCCAATCGGGCAGTCCGGAACTTGGTAAAAGTACTCCCCTATCCAGTCCTGGTGCGGGATGTTTTTTCGATCCAGTGCCATCCAGTCCCCAGAGGGATATCTCAACCCTTTGTGGCCGCAGAAATTCATTGTACATGGTTTGTCATACTCGGATATGTATATATCAGGACTCCAACTAGCCACTGTATGATTGGGAGTGTAATATATCACTAAGGATCCATCCAACATATGAGAATCATCACATGAATCTTTTATCGAGTGATCCGATATCCAAAGACGATTGTAAAAATGTGTGATACAAACAGAGGACGAACAGGTTCCGTCCAAGAAGATCGAGGATTTGAAGGCATCAGAATACGGATTGTACGAGACAGAGTGAGAACTCACTGAGATGAAGATGCGAGAGGTGGTCGAGCTCTTCATCCAGGAGCATGAAGGCGTTGGGTGCTCAGAAGATGAGAATGCTCCCTCTGATACTTTCTGGATCTCCTCTCTACATAAGATCTCCGAAACCTCTGATGGTTCAGTGTGTAACTCCAAAGTGATACCTCCAAAGAAACCCTTAGAACAGGTTGTACTCAATTTTATGAAGGTGCATATGGATCCAGCAATATTAATGGTTGTCCCTATCAAGGGTAATCTTATTGGAATTGTTATGCCTGACTCAATTTTTGCATAACTTTCCCCAATGGGACATGTTAACGCACTCGGTGTCACATCGTGCCACTGAACATTGTCAGATATTGGAAATAGCATTGTATGTTGTTGTCCTCGTAATGAGTCTCCTACTGTACCCTCTGTCATGGCGACTATTTGTACAGCCACATAGATAAGACTCTTGATCTTCTGGCGAGAGCAATCCATTATTATGACGGGTCATAGCATAGTTGACTTGTGTTAGTTTTTTTCTTGAATAGCTCATTCGAACTCTGAGACTCAGACAAGAGGCATCCCAAACGGATTCAACCGCCACCACAAGAACAGCTTCTNNTCACAAAATTTGATATAGAAAATATCTAGAAATATACGGTGATGATTTTGAAGCACGCTAGTCTGCAATAGTTTTATGATAAGCTGTTTTAGTCCTGAGTCAATTCTTTTTTATAGAAATACACTCTTTATCGCATTCCACCGTTATAGCATACGGTTTAAAGAGTGATAACTTAATAGTATGCCTCTTACGATATTTATTAGGCGAACGAATGCGAATATAATCCAAAAGAGAGGCATGACTCATAGTACTTTCTTCAGACGTTAGATGGAAGGACACCAGTATATTTGGGATTAATCCTATACCCTCACCATCATAATCATATGTAAGAGGGAATATGGGTGAGATGTTGCACAATTTAATTTTGAAAACTCCGGTTACGGTACCACGATAGTGATAAAAGGCTCCAACTTTATGTATGTATTTGCAAGTGAACCCGGCCAATATGTAAAACAAAAAGATATACTCCTTGTCATTGACATAGCCTTTATAATCATCCTTGATAATCTCTAATGACGCCATGAGTTCTTTGTAAGTCCTAGGAGGGTCACGGAATTTTATATCTAAGAGCGCATCGACTCTAACAGTCTTAATAAGAGACATTGTTACTCAGGATTGCAATACCGAGTTTTTTTCTCGATCTGTGTTACTGTTTTGTGATTTTGATCGTTATCTCTCGGTAACGGCACGTCAAAACGAATTGATTGTAGGCCTTATGTGCTCTCAGAATTGCCTTCCCCTGTTCTGGTTCAGTGTCGTAGGACTCCAAGATTTCTAAGCAATCCTCATGTGGGTACTCGACAATGATCGGTTTGCCACCCTTCTTACGGGGCATTACCACCTGGACTTTCCCTCCTCTCTGAACCGTAACTGGAGGTTGCGGATCCGTGGGCTTCAACTCGAGTTGAGGATTTCTCTCCGGTTTTGATACCTTCTTAGCCTGAGAAAGGTTTCTTGGCTTCACGGGCGGAATG

>Manitoba_rhabdovirus_1_8-2884

GTTGTCCTCGTAATGAGTCTCCTAATGTACCCTCCGTCATGGCGACTATTTGTACAGCCACATAGATAAGACTCTTGATCTTCTGGCGAGAGCAATCCATTATTATGACGGGTCATAGCATAGTTGACTTGTGTTAGTTTTTTTCTTGAATAGCTCATTTGAACTCTGAGACTCAGACAAGAGGCATCCCAAACGGACTCAACCGCCACCACAAGAACAGCTTCTAGAGCTCGTTATCTCCTATTACCATATCCTATATAAATACAAAATATAAGCCTATTAGAATTAATCATAAAATTTGATATAGAAAATATCTAGAAATATACTGTGATGATTTTGAAGCACGCTAGTCTGCAATAGTTTTATGATAAGCTGTTTTAGTCCTGAGTCAATTCTTCTTTATAGAAATACACTCTTTATCGCATTCCACCGTTATAGCATACGGTTTAAAGAGTGATAACTTAATAGTATGCCTCTTACGATATTTATTAGGCGAACGAATGCGAATATAATCCAAAAGAGAGGCATGACTCATAGTACTTTCTTCAGACGTTAGATGGAAGGACACCAGTATATTTGGGATTAATCCTATACCCTCACCATCATAATCATATGTAAGAGGGAATATGGGTGAGATGTTGCACAATTTAATTTTGAAAACTCCGGTTACGGTACCACGATAGTGATAAAAGGCTCCAACTTTATGTATGTATTTGCAAGTGAACCCAGCCAATATGTAAAACAAAAAGATATACTCCTTGTCATTGACATAGCCTTTATAATCATCCTTGATAATCTCTAATGACGCCATGAGTTCTTTGTAAGTCCTAGGAGGGTCACGGAATTTTATATCTAAGAGCGCATCGACTCTAACAGTCTTAATAAGAGACATTGTTACTCAGGATTGCAATACCGAGTTTTTTTCTCGATCTGTGTTACTGTTTTGTGATTTTGATCGTTATCTCTCGGTAACGGCATGTCAAAACGAATTGATTGTAGGCCTTATGTGCTCTCAGAATTGCCTTCCCCTGTTCTGGTTCAGTGTCGTAGGACTCCAAGATTTCTAAGCAATCCTCATGTGGGTACTCGACAATGATCGGTTTGCCACCCTTCTTACGGGGCATTACCACCTGGACTTTCCCTCCTCTCTGAACCGTAACTGGAGGTTGCGGATCCGTGGGCTTCAACTCGAGTTGAGGATTTCTCTCCGGTTTTGATACCTTCTTAGCCTGAGAAAGGTTTCTTGGCTTCACGGGCGGAATGAAAGGCGGCGGAGTCTCCTGCTGAGACGTTTGTGTCCTCCACTGAGGATTACTCTGGATACGGGAAGCTTTATTCCTGAGTTCACCAATTAAGAATTTACCCTCTGCACGGAACGCAACGATCTCCTTATCTGCCAATTCGATTGCCTGAGATAGCATTAAGTTAACTCCGATCAACAATTCATCAGCAGTGATTATTCCGTTTCGGTCTGGGGTCATTTCTACAGCCAGTTCCTTGGACTTTTCCGTGATCTCACCTGCCAG

>Manitoba_rhabdovirus_1_8-6304

TCAGAGACTCCGACTCCGTCCACAAATCTGAGGTCCAGGTCTCATGCTCCATCTGATGCACCTCTACTTCAAATACATCCAACTCGTTTCTCTCAACATCCGCGAACCGATGACCGTACATGTTCGACATTTTGAATGACAACTATTCGACCGTGATCTTTCGACACTAGCAGTGGAATTGTGTTACTGTTTTTTTCACGGTCAAGGCTGGTAAAATTCTGTTTCTTGATTGCTTGAAGTCTTTGTAAACAATCTTATTAACTGATACCTTTTATTCTTAAAACAGTGGAGGAGTTTTGACCAAAAGATTAGAATCAAAACGCCGATTAGAACTGTTACAGCGCATGCTGTTAATGTATATAACATAGGTGTCCAGTGCAAATCTGTCAAAGAGCTATCATCCACGAAATTATAATCTGTCTCTGTACCGGGGTGTAGCTGTCTTCTCCGGTATTCGGGAGACCGCTTTGCCATCGGTACTCTGTAGTGAGAGCTTAGCCGTGCGATTCTTTTATAGTTTTCTATGTCTTGGATGCCAAGGATCAGTGTCTTGTTCTTGACATACAACCCATTGGGTCCATCGATCACATCTGTTTTATTTGATGTAACCCAGTACGGCCATGTCCAAAGTTGGTTGTCTGATCTTCTTATACTAATATACGGAAAACTCAGGCTCGGTTCGATAGTCACTTGTTGGTATAAGCTTAGTCCCATCTCAAATTTTCCTGGACTATAGCGATAGACTGGATGAAATCCGGGATAACGAGGTGTCAATGTTTGCAACTCCGCTCGACTAACCATTTCTCCTGCTATGATTTTATCGACAACTATCTCACACTCATGATCTAAAAATTCATCCAACAAGCTGAGGACAGCGTTCTGTATAAATTCTTTGTCATTAATTATGTTGACTTGTGACCCAATCGGGCAGTCCGGAACTTGGTAAAAGTACTCCCCTATCCAGTCCTGGTGCGGGATGTTTTTTCGGTCCAGTGCCATCCAGTCCCCAGAGGGATATCTCAACCCTCGGTGGCCGCAGAAATTCATTGTACATGGTTTGTCATACTCGGATATGTATATATCAGGACTCCAACTAGCTACTGTATGATTGGGAGTGTAATATATCACTAAGGATCCATCCAACATATGAGAATCATCACATGAATCTTTTATCGAGTGATCCGATATCCAAAGACGATTGTAAAAATGTGTGATACAAACAGAGGACGAACAGGTTCCGTCCAAGAAGATCGAGGATTTGAAGGCATCAGAATACGGATTGTACGAGACAGAGTGAGAACTCACTGAGATGAAGATGCGAGAGGTGGTCGAGCTCTTCATCCAGGAGCATGAAGGCGTTGGGTGCTCAGAAGATGAGAATGCTCCCTCTGATACTTTCTGGATCTCCTCTCTACATAAAATCTCCGAAACCTCTGATGGTTCAGTGTGTAACTCCAAAGTGATACCTCCAAAGAAACCCTTAGAACAGGTTGTACTCAATTTTATGAAGGTGCATATGGATCCAGCAATATTAATGGTTGTCCCTATCAAGG

>Manitoba_rhabdovirus_1_45-557

AAGTACTCCCCTATCCAGTCCTGGTGCGGGATGTTTTTCCGGTCCAGTGCCATCCAGTCCCCAGAGGGATATCTCAACCCTCGGTGGCCGCAGAAATTCATTGTACATGGTTTGTCATACTCGGATATGTATATATCAGGACTCCAACTAGCTACTGTATGATTGGGAGTGTAATATATCACTAAGGATCCATCCAACATATGAGAATCATCACATGAATCTTTTATCGAGTGATCCGATATCCAAAGACGATTGTAAAAATAAGTGACACAAACAGAGGACGAACATGTTCCGTCCAAGAAGATCGAGGATTTGAAAGCATCAGAATACGGATTGTACGAGACAGAGTGAGAACTAACTGAGATGAAGATGCGAGAGGTGGTCGAGCTCTTCATCCAGGAGCATGAAGGCGTTGGGTGCTCAGAAGATGAGAATGCTCCCTCTGATACTTTCTGGATCTCCTCTCTACATAAAATCTCCGAAACCTCCGATGGTTCAGTGTGTAACTCCAGAGTGATACCTCCAAAGAAACCCTTAGAACAGGTTGTACTCAATTTTGTGAAGGTGCATATGGATCCGGCAATATTAATGGTTGTCCCTATCAAGGGTAATCTTATTGGAATTGTTATGCCCGACTCAATTTTTGCATAACTTTCCCCAATGGGACATGTTAACGCACTCGGTGTCACATCGTGCCACTGAACATTGTCAGATATTGGAAATAGCATTGTATGTTGTTGTCCTCGTAATGAGTCTCCTAATGTACCCTCCGTCATGGCGACTATTTGTACAGCCACATAGATAAGACTCTTGATCTTCTGGCGAGAGCAATCCATTATTATGACGGGTCATAGCATAGTTGACTTGTGTTAGTTTTTTTCTTGAATAGCTCATTCGAACTCTGAGACTCAGACAAGAGGCATCTCACACGGACTCAACCGCCACCACAAGAACAGCTTCTAGATCACGCTGTCTCCTATTACCATATCCTATATAAATACAAAATATAAGCCTATTAGAATTAATCATAAAATTTGATATAGAAAATATTTAGAAATATACGGTGATGATTTTGAAGCACGCTAGTCTGCAATAGTTTTATGATAAGCTGTTTAAGTCCTGAGTCAATTCTTTTTTATAGAAATACACTCCTTATCGCATTCCACCGTTATAGCATACGGTTTAAAGAGTGATAACTTAATAGTATGCCTCTTACGATATTTATTAGGCGAACGAATGCGAATATAATCCAAAAGAGAGGCATGACTCATAGTACTTTCTTCAGACGTTAGATGGAAGGACACCAGTATATTTGGGATTAATCCTATACCCTCACCATCATAATCATAAGTAAGAGGGAATATGGGTGATATGTTGCACAATTTAATTTTGAAAACTCCGGTTATGGTACCACGATAGTGATAAAAGGCTCCAACTTTATGTATATATTTGCAAGTGAACCCAGCCAATATGTAAAACAAAAAGATATACTCCTTGTCATTGACATAGCCTTTATAATCATCTTTGATAATCTCTAATGACGCCATGAGTTCTTTGTAAGTCCTAGGAGGGTCACGGAATTTTATATCTAAGAGCGCATCGACTCTAACAGTCTTAATAAGAGACATTGTTACTCAAGATTGCAATACCGAGTTTTTTTCTCGATCTGTGTTACTGTTTTGTGATTTTGATCGTTATCTCTCGGTAACGGCATGTCAAAACGAATTGATTGTAGGCCTTATGTGCTCTCAGAATTGCCTTCCCCTGTTCTGGTTCAGTGTCGTAGGACTCCAAGATTTCTAAGCAATCCTCATGTGGGTACTCGACAATGATCGGTTTGCCACCCTTCTTACGGGGCATTACCACCTGGACTTTCCCTCCTCTCTGAACCGTAACTGGAGGTTGTGGATCCGCGGGCTTCAACTCGAGTTGAGGATTTCTCTCCGGTTTTGATACCTTCTTAGCCTGAGAAAGGTTTCTTGGCTTCACGGGCGGAGTGCAAGGCGGCGGAGTCTCCTGCTGAGACGTTCGTGTCCTCCACTGAGGATCACTCTGGATACGGGAAGCTTTATTCCTGAGTTCACCAATTAAGAATTTACCCTCTGCACGGAACGCAACGATCTCCTTATCTGCCAATTCGATTGCCTGAGATAGCATTAAGTTAACTCCGATCAACAATTCATCAGCAGTGATTATTCCGTTTCGATCTGGGGTCATTTCTACAGCCAGTTCCTTGGACTTTTCCGTGATCTCACCTGCAAGTAGACCTAATTCATGACATTCTGCCAGCGTCTGGAGATCGATATCAGATGGTTCCGATTCTCTCCCGAGAAGGGTTGCTGTTGAGGAATCTGATCCACCGCTAGCGAGATCGGTTGGATCGAAATCGGTCCCACTTCCGCTTTCCTGAGAGTAGTGCTGAGTCCGTTGCAACTGTAGGTTTCTCTGCGAGCTAGTAATGATACCGCTGGTTTGAACTGATACTGTAGCAGTCTCATTACCTTCTAACTTCGATTCCCGTGACGGCCCGAGCTGGACGATCTTACGATTGTTTTGATCTGGAAACGTCTTTGGAATCACTCCAGTCGCAATTGGTGCCTCTTCATCTAGTCCAGGATCCAACCCACCATCTCCGAGTGGATCCAACACCTCTCCCAATTTCTCCTCATTTAGAATTTTCTGAGGATCCAGGAGGCGACCCGGACGTGCATTTGGCTTTACTTTCAAGCTNNNNNNNNNNNNNNNNNNNNNNNNNNNNNNNNNNNNNNNNNNNNNNNNNNNNNNNNNNNNNNNNNNNNNNNNNNNNNNNNNNNNNNNNGAAGTGTCTGGTACACTGGCAAACACATTCACATCTGAAAAGATCGCAGACTTATTCTCCATTTTGATATGTGTTAATGTTTTTTTCACGGACATTACGGATCTAGGAGCAGCGGTCCGGCCTTCTCATACAGCAGTTTTCCGATCGAATCATCTCGACAAGCAGGATTCTGCTCCCACTGCGTAGCCACCTTCTTGATGATGTGATCAGGTACCTCTCCCTGATTTTTCAGAATGTGTGCCAACCACAAGGAGGACTCCTTCTCTTGTGGCTCTCCTTCTGGCGGAGGTTGCGGATCGTCCTCTTCTTCCAATTCGTCTTTCTCGCCATCCACTCCGAATTGCTGAGAGAGAGTCGCAAACTTGCCCATCACATAATGCATTACGACTGCATTGGAGATGATGTTGTTCACTTCGGGATCCCCAACTACCCTAGCTCGTTTGGATCTTTCGAGATTGGAGCTCACTCCCAACGTGTGATAGAAGAAATGAAGATCCAGGTTCACCGAGGCCGAATAGGGGGATTTAGAAGACAATCCCAAATCCATGAAGTACATGGCATAAGATCTGGGATTGTCCATTTCCTCCCCTCCTTTGACAATTCTCTCGAATTGGTCGGCACATCTGTCGGTCCAGATCCACTCCGCGAATGCAGGGAACGTCAATCCCAGACACTTGGTAATCATTCGGGTGGCAATGAGGGCTGCGCAATCCTTGAATCTGGTGACAATCGTTCCGATCCGAGCTTGAGAGTAGGGATGAAATGGAAACTCATTCAGGAACATATCAATTGCGGCCATGATCTGCACATATGGTTGATAGGCAATCCACTGTTTGCAGGCATCAATTATATCTCCCAATTCCGCGGTTTGATTCAACCCCAGCGGTACCAGAAGGTTCTTCACATTCTTCAAGACTTGTTCCTTGTACTCTACCCGTGAAATCTGTCCAACTCGGTAAACGCTGCAGATCAGTATGATGCCCCGCAGAATCTCCTGCTCGGTCAAGAGGACGTTTCCCTCGACTACAGCTGGCACTGATTCCCTTTCGTCCGGAATGATGATCGCACTTAATCCAACACGGTCTGTTCGCTTTCCGATGTCGCGATTGAATGAAGACCAATCCTGATTGAGGAGTCCCTTGACCAACTCGAACTCGTC

>Manitoba_rhabdovirus_1_5-3063

CACGGTCAAGGCTGGTAAAATTCTGTTTCTTGATTGCTTGAAGTCTTTGTAAACAATCTTATTAACTGATACCTTTTATTCTTAAAACAGTGGAGGAGTTTTGACCAAAAGATTAGAATCAAAACGCCGATTAGAACTGTTACAGCGCATGCTGTTAATGTATATAACATAGGTGTCCAGTGCAAATCTGTCAAAGAGCTATCATCCACGAAATTATAATCTGTCTCTGTACCGGGGTGTAGCTGTCTTCTCCGGTATTCGGGAGACCGCTTTGCCATCGGTACTCTGTAGTGAGAGCTTAGTCGTGCGATTCTTTTATAGTTTTCTATGTCTTGGATGCCAAGGATCAGTGTCTTGTTCTTGACATACAACCCATTGGGTCCATCGATCACATCTGTTTTATTTGATGTAACCCAGTACGGCCATGTCCAAAGTTGGTTGTCTGATCTTCTTATACTAATATACGGAAAACTCAGGCTCGGTTCGATAGTCACTTGTTGGTATAAGCTTAGTCCCATCTCAAATTTCCCTGGACTATAGCGATAGACTGGATGAAATCCGGGATAACGAGGTGTCAATGTTTGCAACTCCGCTCGACTAACCATTTCTCCTGCTATGATTTTATCGACAACTATCTCACACTCATGATCTAAAAATTCATCCAACAAACTGAGGACAGCGTTCTGTATAAATTCTTTGTCATTAATTATGTTGACTTGTGACCCAATCGGGCAGTCCGGAACTTGGTAAAAGTACTCCCCTATCCAGTCCTGGTGCGGGATGTTTTTCCGGTCCAGTGCCATCCAGTCCCCAGAGGGATATCTCAACCCTCGGTGGCCGCAGAAATTCATTGTACATGGTTTGTCATACTCAGATATGTATATATCAGGACTCCAACTAGCTACTGTATGATTGGGAGTGTAATATATCACTAAGGATCCATCCAACATATGAGAATCATCACATGAATCTTTTATCGAGTGATCCGATATCCAAAGACGATTGTAAAAATGTGTGATACAAACAGAGGACGAACATGTTCCGTCCAAGAAGATCGAGGATTTGAAGGCATCAGAATACGGATTGTACGAGACAGAGTGAGAACTCACTGAGATGAAGATGCGAGAGGTGGTCGAGCTCTTCATCCAGGAGCATGAAGGCGTTGGGTGCTCAGAAGATGAGAATGCTCCCTCTGATACTTTCTGGATCTCCTCTCTACATAAAATCTCCGAAACCTCCGATGGTTCAGTGTGTAACTCCAAAGTGATACCTCCAAAGAAACCCTTAGAACAGGTTGTACTCAATTTTATGAAGGTGCATATGGATCCAGCAATATTAATGGTTGTCCCTATCAAGGGTAATCTTACTGGAATTGTTATGCCTGATTCAATTTTTGCATAACTTTCCCCAATGGGACATGTTAATGCACTTGGTGTCACATCGTGCCACTGAACATTGTCAGATATTGGAAATAGCATTGTATGTTGTTGTCCTCGTAATGAGTCTCCTAATGTACCCTCCGTCATGGCGACTATTTGTACAGCCACATAGATAAGACTCTTGATCTTCTGGCGAGAGCAATCCATTATTATGACGGGTCATAGCATAGTTGACTTGTGTTAGTTTTTTTCTTGAATAGCTCATTCGAACTCTGAGACTCAGACAAGAGGCATCCAAAACGGACTCAACCGCCACCACAAGAACAGCTTCTAGAGCACGTTATCTCCTATTACCATATCCTATATAAATACAAAATATAAGCCTATTAGAATTAATCATAAAATTTGATATAGAAAATATCTAGAAATATACGGTGATGATTTTGAAGCACGCTAGTCTGCAATAGTTTTATGATAAGCTGTTTTAGTCCTGAGTCAATTCTTCTTTATAGAAATACACTCTTTATCGCATTCCACCGTCATAGCATACGGTTTAAAGAGTGATAACTTAATAGTATGCCTCTTACGATATTTATTAGGCGAACGAGTGCGAATATAATCCAAAAGAGAGGCATGACTCATAGTACTTTCTTCAGACGTTAGATGGAAGGACACCAGTATATTTGGGATTAATCCTATACCCTCACCATCATAATCATATGTAAGAGGGAATATGGGTGAGATGTTGCACAATTTAATTTTGAAAACTCCGGTTATGGTACCACGATAGTGATAAAAGGCTCCAACTTTATGTATGTATTTGCAAGTGAACCCAGCCAATATGTAAAACAAAAAGATATACTCCTTGTCATTGACATAGCCTTTATAATCATCCTTGATAATCTCTAATGACGCCATGAGTTCTTTGTAAGTCCTAGGAGGGTCACGGAATTTTATATCTAAGAGCGCATCGACTCTAACAGTCTTAATAAGAGACATTGTTACTCAAGATTGCAATACCGAGTTTTTTTCTCGATCTGTGTTACTGTTTTGTGATTTTGATCGTTATCTCTCGGTAACGGCACGTCAAAACGAATTGATTGTAGGCCTTATGTGCTCTCAGAATTGCCTTCCCCTGTTCTGGTTCAGTGTCGTAGGACTCCAAGATTTCTAAGCAATCCTCATGTGGGTACTCGACAATGATCGGTTTGCCACCCTTCTTACGGGGCATTACCACCTGGACTTTCCCTCCTCTCTGAACCGTAACTGGAGGTTGCGGATCCGTGGGCTTCAACTCGAGTTGAGGATTTCTCTCCGGTTTTGATACCTTCTTAGCCTGAGAAAGGTTTCTTGGCTTCACGGGCGGAATGCAAGGCGGCGGAGTCTCCTGCTGAGACGTTCGTGTCCTCCACTGAGGATCACTCTGGATACGGGAAGCTTTATTCCTGAGTTCACCAATTAAGAATTTACCCTCNNNNNNNNNACTCCGATCAACAATTCATCCGCAGTGATTATTCCGTTTCGATCTGGGGTCATTTCTACAGCCAGTTCCTTGGACTTTTCCGTGATCTCACCTGCAAGTAGACCTAATTCGTGACATTCTGCCAGCGTCTGGAGATCGATATCAGATGGTTCCGATTCCCTCCCGAGAAGGGTTGCTGTCGAGGAATCTGATCCACCGCTAGCGAGATCGGTTGGATCGAAATCGGTTCCACTTCCGCTTTCCTGAGAGTAGTGCTGAGTCCGTTGCAACTGTAGGTTTCTCTGCGAGCTAGTAATGATACCGCTGGTTTGAACTGATACTGTAGCAGTCTCATTACCTTCTAACTTCGATTCCCGTGACGGCCCGAGCTGGACGATCTTACGATTGTTTTGATCTGGAAACGTCTTTGGAATCACTCCAGTCGCAATTGGTGCCTCTTCATCTAGTCCAGGATCCAACCCGCCATCTCCGAGTGGATCCAACACCTCTCCCAATTTCTCCTCATTTAGAATTTTCTGAGGATCCAGGAGGCGACCCGGACGTGCATTTGGCTTTACTTTCAAGCTTGTGCTAGCCGGTTTGTTCTTCTTGCTTTCCGTCTGTTTTCCTTTTCCACGATTTCCCATGGCGAGTTCAGATTTTCCCTCTTTCTTGCCTGAAGTGTCTGGTACACTGGCAAACACATTCACATCTGAAAAGATCGCAGACTTATTCTCCATTTTGATATGTGTTAATGTTTTTTTCACGGACATTACGGATCTAGGAGCAGCGGTCCGGCCTTCTCATACAGCAGTTTTCCGATCGAATCATCCCGACAAGCAGGATTCTGCTCCCACTGCGTAGCCACCTTCTTGATGATGTGATCAGGTACCTCTCCCTGATTTTTCAGAATGTGTGCCAACCACAAGGAGGACTCCTTCTCCTGTGGCTCTCCTTCTGGCGGAGGTTGCGGATCATCCTCTTCTTCCAATTCGTCTTTCTCGCCATCTACTCCGAATTGCTGAGAGAGAGTCGCAAACTTGCCCATCACATAATGCATTACGACTGCATTGGAGATGATGTTGTTCACTTCTGGATCCCCAACTACTCGGGCTCGTTTGGATCTTTCGAGATTGGAGCTTACTCCCAACGTGTGATAGAAGAAATGAAGATCCAGGTTCACCGAGGCCGAATAGGGGGATTTAGAAGACAATCCCAAGTCCATGAAGTACATGGCATAAGATCTGGGATTGTCCATTTCCTCCCCTCCTTTGACAATTCTCTCGAATTGGTCGGCACATCTGTCGGTCCAGATCCACTCCGCGAATGCAGGGAACGTCAATCCCAGACACTTGGTAATCATTCGGGTGGCAATGAGGGCTGCGCAATCCTTGAATCTGGTGACAATAGTTCCGATCCGAGCTTGAGAGTAGGGATGAAATGGAAACTCATTCAGGAACATATCAATTGCGGCCATGATCTGCACATATGGTTGATAGGCAATCCACTGTTTGCAGGCATCAATGATATCTCCCAATTCTGCGGTTTGATTCAACCCCAGCGGTACCAGAAGGTTCTTCACATTCTTCAGGACTTGTTCCTTGTACTCTACCCGTGCAATCTGTCCAACTCGGTAAACGCTGCAGATCAGTATGATGCCCCGAAGAATCTCCTGCTCGGTCAAGAGGACGTTTCCCTCGACTACAGCTGGCACCGATTCCCTTTCGTCCGGAATGATGATCGCACTTAATCCAACGCGGTCTGTTCGCTTTCCGATGTCGCGATTGAATGAAGACCAATCCTGATTGAGGAGTCCCTTGACCAACTCGAACTCGTCGTATAGGAAGCGTACCACGAATCGATGGTCGACTTTGCCTTTGGTTAGTCCTGATCTTATCGCGGCATGGATCTGTTCTCTCGTCGCACCTGTCCCGGGTATCATCATTATAGTCGGCTTAGTATTACCCGGCTTATTAAACCAGGCCGACGGATACTCAATGGATCTCTCAGTTCCTATGTCAAAATGAACGATAGGAGTAGCGGTCGCGGGATTGTCAGTGATACGTTTGATTACACGTGTCTCAGCCATTGAATACAAACAAAATCGATCTGATACTTTCCTCAGAGGTACTTACACAATTCTGACTTGAAGATGTGTTGACGGTACTAGTCGTTTAATTGTTTTG

>Manitoba_rhabdovirus_1_25-10086

GGGACTCAGACAAGAGGCATCCCAAACGGACTCAACCGCCACCACAAGAACAGCTTCTAGAGCACGTTATCTCCTATTACCATATCCTATATAAATACAAAATATAAGCCTATTAGAATTAATCATAAAATTTGATATAGAAAATATCTAGAAATATACGGTGATGATTTTGAAGCACGCTAGTCTGCAATAGTTTTATGATAAGCTGTTTAAGTCCTGAGTCAATTCTTTTTTATAGAAATACACTCCTTATCGCATTCCACCGTTATAGCATACGGTTTAAAGAGTGATAACTTAATAGTATGCCTCTTACGATATTTATTAGGCGAACGAATGCGAATATAATCCAAAAGAGAGGCATGACTCATAGTACTTTCTTCAGACGTTAGATGGAAGGACACCAGTATATTTGGGATTAATCCTATACCCTCACCATCATAATCATATGTAAGAGGGAATATGGGTGAGATGTTGCACAATTTAATTTTGAAAACTCCGGTTATGGTACCACGATAGTGATAAAAGGCTCCAACTTTATGTATGTATTTGCAAGTGAACCCAGCCAATATGTAAAACAAAAAGATATACTCCTTGTCATTGACATAGCCTTTATAATCATCTTTGATAATCTCTAATGACGCCATGAGTTCTTTGTAAGTCCTAGGAGGGTCACGGAATTTTATATCTAAGAGCGCATCGACTCTAACAGTCTTAATAAGAGACATTGTTACTCAAGATTGCAATACCGAGTTTTTTTCTCGATCTGTGTTACTGTTTTGTGATTTTGATCGTTATCTCTCGGTAACGACA

>Manitoba_rhabdovirus_1_25-11113

CTGTGTTACTGTTTCGTGATTTTGATCGTTATCTCTCGGTAACGGCATGTCAAAACGAATTGATTGTAGGCCTTATGTGCTCTCAGAATTGCCTTCCCCTGTTCTGGTTCAGTGTCGTAGGACTCCAAGATTTCTAAGCAATCCTCATGTGGGTACTCGACAATGATCGGTTTGCCACCCTTCTTACGGGGCATTACCACCTGGACTTTCCCTCCTCTCTGAACCGTAACTGGAGGTTGCGGATCCGCGGGCTTCAACTCTAGTTGAGGATTTCTCTCCGGTTTTGATACCTTCTTAGCCTGAGAAAGGTTTCTTGGCTTCACGGGCGGAGTGCAAGGCGGCGGAGTCTCCTGCTGAGACGTTCGTGTCCTCCACTGAGGATCACTCTGGATACGGGAAGCTTTATTCCTGAGTTCACCAATTAAGAATTTACCCTCTGCACGGAACGCAACGATCTCCTTATCTGCCAATTCGATTGCCTGAGATAGCATTAAGTTAACTCCGATCAACAATTCATCAGCAGTGATAATTCCGTTTCGATCTGGGGTCATTTCTACAGCCAGTTCCTTGGACTTTTCCGTGATCTCACCTGCAAGTAGACCTAATTCATGACATTCTGCCAGCGTCTGGAGATCGATATCAGATGGTTCCGATTCTCTCCCGAGAAGGGTTGCTGTTGAGGAATCTGATCCACCGCTAGCGAGATCGGTTGGATCGAAATCGGTCCCACTTCCGCTTTCCTGAGAGTAGTGCTGAGTCCGTTGCAACTGTAGGTTTCTCTGCGAGCTAGTAATGATACCGCTGGTTTGAACTGATACTGTAGCAGTCTCATTACCTTCTAACTTCGATTCCCGTGACGGCCCGAGCTGGACGATCTTACGATTGTTTTGATCTGGAAACGTCTTTGGAATCACTCCAGTCGCAATTGGTGCCTCTTCATCTAGTCCAGGATCCAACCCACCATCTCCAAGTGGATCCAACACCTCTCCCAATTTCTCCTCATTTAGAATTTTCTGAGGATCCAGGAGGCGACCCGGACGTGCATTTGGCTTTACTTTCAAGCTCGTGCTAGCCGGTTTGTTCTTCTTGCTTTCCGTCTGTTTTCCTTTTCCACGGTTTCCCATAGCGAGTTCAGGTTTTCCTTCTTTCTTGCCCGAAGTGTCTGGTACACTGGCA

>Manitoba_rhabdovirus_1_41-5959

CCTCGATCTGTGTTACTGTTTTGTGATTTTGATCGTTATCTCTCGGTAACGGCATGTCAAAACGAATTGATTGTAGGCCTTATGTGCTCTCAGAATTGCCTTCCCCTGTTCTGGTTCAGTGTCGTAGGACTCCAAGATTTCTAAGCAATCCTCATGTGGGTACTCGACAATGATCGGTTTGCCACCCTTCTTACGGGGCATTACCACCTGGACTTTCCCTCCTCTCTGAACCGTAACTGGAGGTTGCGGATCCGTGGGCTTCAACTCGAGTTGAGGATTTCTCTCCGGTTTTGATACCTTCTTAGCCTGAGAAAGGTTTCTTGGCTTCACGGGCGGAATGAAAGGCGGCGGAGTCTCCTGCTGAGACGTTCGTGTCCTCCACTGAGGATTACTCTGGATACGGGAAGCTTTATTCCTGAGTTCACCAATTAAGAATTTACCCTCTGCACGGAACGCAACGATCTCCTTATCTGCCAATTCGATTGCCTGAGATAGCATTAAGTTAACTCCGATCAACAATTCATCAGCAGTGATTATTCCGTTTCGGTCTGGGGTCATTTCTACAGCCAGTTCCTTGGACTTTTCCGTGATCTCACCTGCCAGTAGACCTAATTCGTGACATTCTGCCAGCGTCTGGAGATCGATATCAGATGGTTCCGATTCTCTCCCGAGAAGGGTTGCTGTCGAGGAATCTGATCCGCCGCTAGCGAGATCGGTTGGATCGAAATCGGTCCCACTTCCGCTTTCCTGAGAGTAGTGCTGAGTCCGTTGCAACTGTAGGTTTCTCTGCGAGCTAGTAATGATACCGCTGGTTTGAACTGATACTGTAGCAGTCTCATTACCTTCTAACTTCGATTCCCGTGACGGCCCGAGCTGGACGATCTTACGATTGTTTTGATCTGGAAACGTCTTTGGAATCACTCCAGTCGCAATTGGTGCCTCTTCATCTAGTCCAGGATCCAACCCACCATCTCCAAGTGGATCCAACACCTCTCCCAATTTCTCCTCATTTAGAATTTTCTGAGGATCCAGGAGGCGACCCGGACGTGCATTTGGCTTTACTTTCAAGCTTGTACTAGCCGGTTTGTTCTTCTTGCTTTCCGTCTGTTTTCCTTTACCACGATTTCCCATGGCGAGTTCAGATTTTCCCTCTTTCTTGCCTGAAGTGTCTGGTACACTGGCAAACACATTCACATCTGAAAAGATCGCAGACTTATTCTCCATTTTG

>Manitoba_rhabdovirus_1_35-18212

GCTCATTCGAACTCCGATACTCAGACAAGGGGCATCTCACACGGACTCGACCGCCACCACAAGAACAGCTTCTAGATCACGCTGTCTCCTATTACCATATCCTATATAAATACAAAATATAAGCCTATTAGAATTAATCATAAAATTTGATATAGAAAATATTTAGAAATATACGGTGATGATTTTGAAGCACGCTAGTCTGCAATAGTTTTATGATAAGCTGTTTAAGTCCTGAGTCAATTCTTTTTTATAGAAATATACTCCTTATCGCATTCCACCGTTATAGCATACGGTTTAAAGAGTGATAACTTAATAGTATGCCTCTTACGATATTTATTAGGCGAACGAATACGAATATAATCCAAAAGAGAGGCATGACTCATAGTACTTTCTTCAGACGTTAGATGGAAGGACACCAGTATATTTGGGATTAATCCTATACCCTCACCATCATAATCATATGTAAGAGGGAATATGGGTGATATGTTGCACAATTTAATTTTGAAAACTCCGGTTATGGTACCACGATAGTGATAAAAGGCTCCAACTTTATGTATATATTTGCAAGTGAACCCAGCCAATATGTAAAACAAAAAGATATACTCCTTGTCATTGACATAGCCTTTATAATCATCTTTGATAATCTCTAATGACGCCATGAGTTCTTTGTAAGTCCTAGGAGGGTCACGGAATTTTATATCTAAGAGCGCATCGACTCTAACAGTCTTAATAAGAGACATTGTTACTCAAGATTGCAATACCGAGTTTTTTTCTCGATCTGTGT

>Manitoba_rhabdovirus_1_4-4817

CACGGACATTACGGATCTAGGAGCAGCGGTCCGGCCTTCTCATACAGCAGTTTTCCGATCGAATCATCCCGACAAGCAGGATTTTGCTCCCACTGCGTAGCCACCTTCTTGATGATGTGATCAGGTACCTCTCCCTGATTTTTCAGAATGTGTGCCAACCACAAGGAGGACTCCTTCTCCTGTGGCTCTCCTTCTGGCGGAGGTTGCGGATCATCCTCTTCTTCCAATTCGTCTTTCTCGCCATCCACTCCGAATTGCTGAGAGAGAGTCGCAAACTTGCCCATCACATAATGCATTACGACTGCATTGGAGATGATGTTGTTCACTTCTGGATCCCCAACTACTCGAGCTCGTTTGGATCTTTCGAGATTGGAGCTTACTCCCAACGTGTGATAGAAGAAATGAAGATCCAGGTTCACCGAGGCCGAATAGGGTGATTTAGAAGACAATCCCAAGTCCATGAAGTACATGGCATAAGATCTGGGATTGTCCATCTCCTCCCCTCCTTTGACAATTCTCTCGAATTGGTCGGCACATCTGTCGGTCCAGATCCACTCCGCGAATGCAGGGAACGTCAATCCCAGACACTTGGTAATCATTCGGGTGGCAATGAGAGCTGCGCAATCCTTGAATCTGGTGACAATAGTTCCGATCCGAGCTTGAGAGTAGGGATGAAATGGAAACTCATTCAGGAACATATCAATTGCGGCCATGATCTGCACATATGGTTGATAGGCAATCCACTGCTTGCAGGCATCAATTATGTCTCCCAATTCTGCGGTTTGATTCAACCCCAGCGGTACCAGAAGGTTCTTCACATTCTTCAGGACTTGTTCCTTGTACTCTACCCGTGCAATCTGTCCAACTCGGTAAACGCTGCAGATCAGTATGATGCCCCGAAGAATCTCCTGCTCGGTCAAGAGGACGTTTCCCTCGACTACAGCTGGCACCGATTCCCTTTCGTCCGGAATGATGATCGCACTTAATCCAACGCGGTCTGTTCGCTTTCCGATGTCGCGATTGAATGAAGACCAATCCTGATTGAGGAGTCCCTTGACCAACTCGAACTCGTCGTATAGGAAGCGTACCACGAATCGATGGTCGACTTTGCCTTTGGTTAGTCCTGATCTTATCGCGGCATGAATCTGTTCTCTTGTCGCACCTGTCCCGGGTATCATCATTATAGTCGGCTTAGTATTACCCGGCTTATTAAACCAGGCCGACGGATACTCAATGGATCTCTCAGTTCCTATGTCAAAATGAACGATAGGAGTAGCGGTCGCAGGATTGTCAGTGATGCGTTTGATTACACGTGTCTCAGCCATTGAATACAAACAAAATCGATCTGATACTTTCCTCAGAGGTACTTACACAATTCTGACTTGAAGATGTGTTGACGGTACTAGTCGTTTAATTGTTT

>Manitoba_rhabdovirus_1_40-7619

ATCACTGCTGATGAATTGTTGATCGGAGTTAACTTAATGCTATCTCAGGCAATCGAATTGGCAGATAAGGAGATCGTTGCGTTCCGTGCAGAGGGTAAATTCTTAATTGGTGAACTCAGGAATAAAGCTTCCCGTATCCAGAGTNNNNNNNNNNCAACCTCCAGTTACGGTTCAGAGAGGAGGGAAAGTCCAGGTGGTAATGCCCCGTAAGAAGGGTGGCAAACCGATCATTGTCGAGTACCCACATGAGGATTGCTTAGAAATCTTGGAGTCCTACGACACTGAACCAGAACAGGGGAAGGCAATTCTGAGAGCACATAAGGCCTACAATCAATTCGTTTTGACATGCCGTTACCGAGAGATAACGATCAAAATCACAAAACAGTAACACAGATCGAGAAAAAAACTCGGTATTGCAATCCTGAGTAACAATGTCTCTTATTAAGACTGTTAGAGTCGATGCGCTCTTAGATATAAAATTCCGTGACCCTCCTAGGACTTACAAAGAACTCATGGCGTCATTAGAGATTATCAAGGATGATTATAAAGGCTATGTCAATGACAAGGAGTATATCTTTTTGTTTTACATATTGGCTGGGTTCACTTGCAAATACATACATAAAGTTGGAGCCTTTTATCACTATCGTGGTACCGTAACCGGAGTTTTCAAAATTAAATTGTGCAACATCTCACCCATATTCCCTCTTACATATGATTATGATGGTGAGGGTATAGGATTAATCCCAAATATACTGGTGTCCTTCCATCTAACGTCTGAAGAAAGTACTATGAGTCATGCCTCTCTTTTGGATTATATTCGCATTCGTTCGCCTAATAAATATCGTAAGAGGCATACTATTAAGTTATCACTCTTTAAACCGTATGCTATAACGGTGGAATGCGATAAAGAGTGTATTTCTATAAAAAAGAATTGACTCAGGACTAAAACAGCTTATCATAAAACTATTGCAGACTAGCGTGCTTCAAAATCATCACCGTATATTTCTAGATATTTTCTATATCAAATTTTATGATTAATTCTAATAGGCTTATATTTTGTATTTATATAGGATATGGTAATAGGAGATAACGAGCTCTAGAAGCTGTTCTTGTGGTGGCGGTTGAGTCCGTTTGGGATGCCTCTTGTCTGAGTCTCAGAGTTCGAATGAGCTATTCAAGAAAAAAACTAACACAAGTCAACTATGCTATGACCCGTCATAATAATGGATTGCTCTCGCCAGAAGATCAAGAGTCTTATCTATGTGGCTGTACAAATAGTCGCCATGACGGAGGGTACATTAGGAGACTCATTACGAGGACAACAACATACAATGCTATTTCCAATATCTGACAATGTTCAGTGGCACGATGTGACACCGAGTGCGTTAACATGTCCCATTGGGGAAAGTTATGCAAAAATTGAGTCAGGCATAACAATTCCAATAAGATTACCCTTGATAGGGACAACCATTAATATTGCTGGATCCATATGCACCTTCATAAAATTGAGTACAACCTGTTCTAAGGGTTTCTTTGGAGGTATCACTTTGGAGTTACACACTGAACCATCGGAGGTTTCGGAGATTTTATGTAGAGAGGAGATCCAGAAAGTATCAGAGGGAGCATTCTCATCTTCTGAGCACCCAACGCCTTCATGCTCCTGGATGAAGAGCTCGACCACCTCTCGCATCTTCATCTCAGTGAGTTCTCACTCTGTCTCGTACAATCCGTATTCTGATGCCTTCAAATCCTCGATCTTCTTGGACGGAACCTGTTCGTCCTCTGTTTGTATCACACATTTTTACAATCGTCTTTGGATATCGGATCACTCGATAAAAGATTCATGTGATGATTCTCATATGTTGGATGGATCCTTAGTGATATATTACACTCCCAATCATACAGTAGCTAGTTGGAGTCCTGATATATACATATCCGAGTATGACAAACCATGTACAATGAATTTCTGCGGCCACCGAGGGTTGAGATATCCCTCTGGGGACTGGATGGCACTGGACCGAAAAAACATCCCGCACCAGGACTGGATAGGGGAGTACTTTTACCAAGTTCCGGACTGCCCGATTGGGTCACAAGTCAACATAATTAATGACAAAGAATTTATACAGAACGCTGTCCTCAGCTTGTTGGATGAATTTTTAGATCATGAGTGTGAGATAGTTGTCGATAAAATCATAGCAGGAGAAATGGTTAGTCGAGCGGAGTTGCAAACATTGACACCTCGTTATCCCGGATTTCATCCAGTCTATCGCTATAGTCCAGGGAAATTTGAGATGGGACTAAGCTTATACCAACAAGTGACTATCGAACCGAGCCTGAGTTTTCCGTATATTAGTATAAGAAGATCAGACAACCAACTTTGGACATGGCCGTACTGGGTTACATCAAATAAAACAGATGTGATCGATGGACCCAATGGGTTGTATGTCAAGAACAAGACACTGATCCTTGGCATCCAAGACATAGAAAACTATAAAAGAATCGCACGACTAAGCTCTCACTACAGAGTACCGATGGCAAAGCGGTCTCCCGAATACCGGAGAAGACAGCTACACCCCGGTACAGAGACAGATTATAATTTCGTGGATGATAGCTCTTTGACAGATTTGCACTGGACACCTATGTTATATACATTAACAGCATGCGCTGTAACAGTTCTAATCGGCGTTTTGATTCTAATCTTTTGGTCAAAACTCCTCCACTGTTTTAAGAATAAAAGGTATCAGTTAATAAGATTGTTTACAAAGACTTCAAGCAATCAAGAAACAGAATTTTACCAGCCTTGACCGTGAAAAAAACATTAACAC

>Manitoba_rhabdovirus_1_27-7554

GCCACATAGATAAGACTCTTGATCTTCTGGCGAGAGCAATCCATTATTATGACGGGTCATAGCATAGTTGACTTGTGTTAGTTTTTTTCTTGAATAGCTCATTCGAACTCTGAGACTCAGACAAGAGGCATCCCAAACGGACTCAACCGCCACCACAAGAACAGCTTCTAGAGCTCGTTATCTCCTATTACCATATCCTATATAAATACAAAATATAAGCCTATTAGAATTAATCATAAAATTTGATATAGAAAATATCTAGAAATATACGGTGATGATTTTGAAGCACGCTAGTCTGCAATAGTTTTATGATAAGCTGTTTTAGTCCTGAGTCAATTCTTCTTTATAGAAATACACTCTTTATCGCATTCCACCGTTATAGCATACGGTTTAAAGAGTGATAACTTAATAGTATGCCTCTTACGATATTTATTAGGCGAACGAATGCGAATATAATCCAAAAGAGAGGCATGACTCATAGTACTTTCTTCAGACGTTAGATGGAAGGACACCAGTATATTTGGGATTAATCCTATACCCTCACCATCATAATCATATGTAAGAGGGAATATGGGTGAGATGTTGCACAATTTAATTTTGAAAACTCCGGTTANNATAATCTCTAATGACGCCATGAGTTCTTTGTAAGTCCTAGGAGGGTCACGGAATTTTATATCTAAGAGCGCATCGACTCTAACAGTCTTAATAAGAGACATTGTTACTCAGGATTGCAATACCGAGTTTTTTTCTCGATCTGTGTTACTGTTTTGTGATTTTGATCGTTATCTCTCGGTAACGGCATGTCAAAACGAATTGATTGTAGGCCTTATGTGCTCTCAGAATTGCCTTCCCCTGTTCTGGTTCAGTGTCGTAGGACTCCAAGATTTCTAAGCAATCCTCATGTGGGTACTCGACAATGATCGGTTTGCCACCCTTCTTACGGGGCATTACCACCTGGACTTTCCCTCCTCTCTGAACCGTAACTGGAGGTTGCGGATCCGTGGGCTTCAACTCGAGTTGAGGATTTCTCTCCGGTTTTGATACCTTCTTAGCCTGAGAAAGGTTTCTTGGCTTCACGGGCGGAATGAAAGGCGGCGGAGTCTCCTGCTGAGACGTTTGTGTCCTCCACTGAGGATTACTCTGGATACGGGAAGCTTTATTCCTGAGTTCACCAATTAAGAATTTACCCTCTGCACGGAACGCAACGATCTCCTTATCTGCCAATTCGATTGCCTGAGATAGCATTAAGTTAACTCCGATCAACAATTCATCAGCAGTGATTATTCCGTTTCGGTCTGGGGTCATTTCTACAGCCAGTTCCTTGGACTTTTCCGTGATCTCACCTGCCAGTAGACCTAATTCGTGACATTCTGCCAGCGTCTGGAGATCGATATCAGATGGTTCCGATTCTCTCCCGAGAAGGGTTGCTGTAGAGGAATCTGATCCGCCGCTAGCGAGATCGGTTGGATCGAAATCGGTCCCACTTCCGCTTTCCTGAGAGTAGTGCTGAGTCCGTTGCAACTGTAGGTTTCTCTGCGAGCTAGTAATGATACCGCTGGTTTGAACTGATACTGTAGCAGTCTCATTACCTTCTAACTTCGATTCCCGTGACGGCCCGAGCTGGACGATCTTACGATTGTTTTGATCTGGAAACGTCTTAGGAATCACTCCAGTCGCAATTGGTGCCTCTTCATCTAGTCCAGGATCCAACCCGCCATCTCCAAGTGGATCCAACACCTCTCCCAATTTCTCCTCATTTAGAATTTTCTGAGGATCCAGGAGGCGACCCGGACGTGCATTTGGCTTTACTTTCAAGCTTGTACTAGCCGGTTTGTTCTTCTTGCTTTCCGTCTGTTTTCCTTTACCACGATTTCCCATGGCGAGTTCAGATTTTCCCTCTTTCTTGCCTGAAGTGTCTGGTACACTGGCAAACACATTCACATCTGAAAAGATTGCAGACTTATTCTCCATTTTGATATGTGTTAATGTTTTTTTCACGGACATTACGGATCTAGGAGCAGCGGTCCGGCCTTCTCATACAGCA

>Manitoba_rhabdovirus_1_1-2684

GTCATGGCGACTATTTGTACAGCCACATAGATAAGACTCTTGATCTTCTGGCGATAGCAATCCATTATTATGACGGGTCATAGCATAGTTGACTTGTGTTAGTTTTTTTCTTGAATAGCTCATTCGAACTCTGAGACTCAGACAAGAGGCATCCCAAACGGACTCAACCGCCACCACAAGAACAGCTTCTAGAGCACGTTATCTCCTATTACCATATCCTATATAAATACAAAATATAAGCCTATTAGAATTAATCATAAAATTTGATATAGAAAATATCTAGAAATATACGGTGATGATTTTGAAGCACGCTAGTCTGCAATAGTTTTATGATAAGCTGTTTTAGTCCTGAGTCAATTCTTTTTTATAGAAATACACTCTTTATCGCATTCCACCGTTATAGCATACGGTTTAAAGAGTGATAACTTAATAGTATGCCTCTTACGATATTTATTAGGCGAACGAATGCGAATATAATCCAAAAGAGAGGCATGACTCATAGTACTTTCTTCAGACGTTAGATGGAAGGACACCAGTATATTTGGGATTAATCCTATACCCTCACCATCATAATCATATGTAAGAGGGAATATGGGTGAGATGTTGCACAATTTAATTTTGAAAACTCCGGTTATGGCACCACGATAGTGATAAAAGGCTCCAACTTTATGTATGTATTTGCAAGTGAACCCAGCCAATATGTAAAACAAAAAGATATACTCCTTGTCATTGACATAGCCTTTATAATCATCCTTGATAATCTCTAATGACGCCATGAGTTCTTTGTAAGTCCTAGGAGGGTCACGGAATTTTATATCTAAGAGCGCATCGACTCTAACAGTCTTAATAAGAGACATTGTTACTCAAGATTGCAATACCGAGTTTTTTTCTCGATCTGTGTTACTGTTTTGTGATTTTGATCGTTATCTCTCGGTAACGGCACGTCAAAACGAATTGATTGTAGGCCTTATGTGCTCTCAGAATTGCCTTCCCCTGTTCTGGTTCAGTGTCGTAGGACTCCAAGATTTCTAAGCAATCCTCATGTGGGTACTCGACAATGATCGGTTTGCCACCCTTCTTACGGGGCATTACCACCTGGACTTTCCCTCCTCTCTGAACCGTAACTGGAGGTTGCGGATCCGTGGGCTTCAACTCGAGTTGAGGATTTCTCTCCGGTTTTGATACCTTCTTAGCCTGAGAAAGGTTTCTTGGCTTCACGGGCGGAATGCAAGGCGGCGGAGTCTCCTGCTGAGACGTTCGTGTCCTCCACTGAGGATCACTCTGGATACGGGAAGCTTTATTCCTGAGTTCACCAATTAAGAATTTACCCTCTGCACGGAACGCAACGATCTCCTTATCTGCCAATTCGATTGCCTGAGATAGCATTAAGTTAACTCCGATCAACAATTCATCAGCAGTGATTATTCCGTTTCGATCTGGGGTCATTTCTACAGCCAGTTCCTTGGACTTTTCCGTGATCTCACCTGCAAGTAGACCTAATTCGTGACATTCTGCCAGCGTCTGGAGATCGATATCAGATGGTTCCGATTCTCTCCCGAGAAGGGTTGCTGTCGAGGAATCTGATCCACCGCTAGCGAGATCGGTTGGATCGAAATCGGTCCCACTTCCGCTTTCCTGAGAGTAGTGCTGAGTCCGTTGCAACTGTAGGTTTCTCTGCGAGCTAGTAATGATACCGCTGGTTTGAACTGATACTGTAGCAGTCTCATTACCTTCTAACTTCGATTCCCGTGACGGCCCGAGCTGGACGATCTTACGATTGTTTTGATCTGGAAACGTCTTTGGAATCACTCCAGTCGCAATTGGTGCCTCTTCATCTAGTCCAGGATCCAACCCGCCATCTCCGAGTGGATCCAACACCTCTCCCAATTTCTCCTCATTTAGAATTTTCTGAGGATCCAGGAGGCGACCCGGACGTGCATTTGGCTTTACTTTCAAGCTTGTGCTAGCCGGTTTGTTCTTCTTGCTTTCCGTCTGTTTTCCTTTTCCACGATTTCCCATGGCGAGTTCAGATTTTCCCTCTTTCTTGCCTGAAGTGTCTGGTACACTGGCAAACACATTCACATCTGAAAAGATCGCAGACTTATTCTCCATTTTGATATGTGTTAATGTTTTTTTCACGGACATTACGGATCTAGGAGCAGCGGTCCGGCCTTCTCATACAGCAGTTTTCCGATCGAATCATCCCGACAAGCAGGATTCTGCTCCCACTGCGTAGCCACCTTCTTGATGATGTGATCAGGTACCTCTCCCTGATTTTTCAGAATGTGTGCCAACCACAAGGAGGACTCCTTCTCCTGTGGCTCTCCTTCTGGCGGAGGTTGCGGATCATCCTCTTCTTCCAATTCGTCTTTCTCGCCATCCACTCCGAATTGCTGAGAGAGAGTCGCAAACTTGCCCATCACATAATGCATTACGACTGCATTGGAGATGATGTTGTTCACTTCTGGATCCCCAACTACTCGGGCTCGTTTGGATCTTTCGAGATTGGAGCTTACTCCCAACGTGTGATAGAAGAAATGAAGATCCAGGTTCACCGAGGCCGAATAGGGGGATTTAGAAGACAATCCCAAGTCCATGAAGTACATGGCATAAGATCTGGGATTGTCCATTTCCTCCCCTCCTTTGACAATTCTCTCGAATTGGTCGGCACATCTGTCGGTCCAGATCCACTCCGCGAATGCAGGGAACGTCAATCCCAGACACTTGGTAATCATTCGGGTGGCAATGAGGGCTGCGCAATCCTTGAATCTGGTGACAATAGTTCCGATCCGAGCTTGAGAGTAGGGATGAAATGGAAACTCATTCAGGAACATATCAATTGCGGCCATGATCTGCACATATGGTTGATAGGCAATCCACTGTTTGCAGGCATCAATTATATCTCCCAATTCTGCGGTTTGATTCAACCCCAGCGGTACCAGAAGGTTCTTCACATTCTTCAGGACTTGTTCCTTGTACTCTACCCGTGCAATCTGTCCAACTCGGTAAACGCTGCAGATCAGTATGATGCCCCGAAGAATCTCCTGCTCGGTCAAGAGGACGTTTCCCTCGACTACAGCTGGCACCGATTCCCTTTCGTCCGGAATGATGATCGCACTTAATCCAACGCGGTCTGTTCGCTTTCCGATGTCGCGATTGAATGAAGACCAATCCTGATTGAGGAGTCCCTTGACCAACTCGAACTCGTCGTATAGGAAGCGTACCACGAATCGATGGTCGACTTTGCCTTTGGTTAGTCCTGATCTTATCGCGGCATGGATCTGTTCTCTCGTCGCACCTGTCCCGGGTATCATCATTATAGTCGGCTTAGTATTACCCGGCTTATTAAACCAGGCCGACGGATACTCAATGGATCTCTCAGTTCCTATGTCAAAATGAACGATAGGAGTAGCGGTCGCAGGATTGTCAGTGATACGTTTGATTACACGTGTCTCAGCCATTGAATACAAACAAAATCGATCTGATACTTTCCTCAGAGGTACTTACACAATTCTGACTTGAAGATGTGTTGACGGTACTAGTCGTTTAATTGTTTTGTTTAGAT

>Manitoba_rhabdovirus_1_1-11625

CTTAAAACAGTGGAGGAGTTTCGACCAAAAGATTAGAATCAAAACGCCGATTAGAACTGTTACAGCGCATGCTGTTAATGTATATAACATAGGTGTCCAGTGCAAATCTGTCAAAGAGCTATCATCCACGAAATTATAATCTGTCTCTGTACCGGGGTGTAGCTGTCTTCTCCGGTATTCGGGAGACCGCTTTGCCATCGGTACTCTGTAGTGAGAGCTTAGTCGTGCGATTCTTTTATAGTTTTCTATGTCTTGGATGCCAAGGATCAGTGTCTTGTTCTTGACATACAACCCATTGGGTCCATCGATCACATCTGTTTTATTTGATGTAACCCAGTACGGCCATGTCCAAAGTTGGTTGTCTGATCTTCTTATACTAATATACGGAAAACTCAGGCTCGGTTCGATAGTCACTTGTTGGTATAAGCTTAGTCCCATCTCAAATTTCCCTGGACTATAGCGATAGACTGGATGAAATCCGGGATAACGAGGTGTCAATGTTTGCAACTCCGCTCGACTAACCATTTCTCCTGCTATGATTTTATCGACAACTATCTCACACTCATGATCTAAAAATTCATCCAACAAACTGAGGACAGCGTTCTGTATAAATTCTTTGTCATTAATTATGTTGACTTGTGACCCAATCGGGCAGTCCGGAACTTGGTAAAAGTACTCCCCTATCCAGTCCTGGTGCGGGATGTTTTTCCGGTCCAGTGCCATCCAGTCCCCAGAGGGATATCTCAACCCTCGGTGGCCGCAGAAATTCATTGTACATGGTTTGTCATACTCGGATATGTATATATCAGGACTCCAACTAGCCACTGTATGATTGGGAGTGTAATATATCACTAAGGATCCATCCAACATATGAGAATCATCACATGAATCTTTTATCGAGTGATCCGATATCCAAAGACGATTGTAAAAATGTGTGATACAAACAGAGGACGAACAGGTTCCGTCCAAGAAGATCGAGGATTTGAAGGCATCAGAATACGGATTGTACGAGACAGAGTGAGAACTCACTGAGATGAAGATGCGAGAGGTGGTCGAGCTCTTCATCCAGGAGCATGAAGGCGTTGGGTGCTCAGAAGATGAGAATGCTCCCTCTGATACTTTCTGGATCTCCTCTCTACATAAAATCTCCGAAACCTCCGATGGTTCAGTGTGTAACTCCAAAGTGATACCTCCAAAGAAACCCTTAGAACAGGTTGTACTCAATTTTATGAAGGTGCATATGGATCCAGCAATATTAATGGTTGTCCCTATCAAGGGTAATCTTATTGGAATTGTTATGCCTGACTCAATTTTTGCATAACTTTCCCCAATGGGACATGTTAACGCACTCGGTGTCACATCGTGCCACTGAACATTGTCAGATATTGGAAATAGCATTGTATGTTGTTGTCCTCGTAATGAGTCTCCTAATGTACCCTCCGTCATGGCGACTATTTGTACAGCCACATAGATAAGACTCTTGATCTTCTGGCGAT

>Manitoba_rhabdovirus_1_42-2118

CTGTGTTACTGTTTTGTGATTTTGATCGTTATCTCTCGGTAACGGCATGTCAAAACGAATTGATTGTAGGCCTTATGTGCCCTCAGAATTGCCTTCCCCTGTTCTGGTTCAGTGTCGTAGGACTCCAAGATTTCTAAGCAATCCTCATGTGGGTACTCGACAATGATCGGTTTGCCACCCTTCTTACGGGGCATTACCACCTGGACTTTCCCTCCTCTCTGAACCGTAACTGGAGGTTGCGGATCCGTGGGCTTCAACTCGAGTTGAGGATTTCTCTCCGGTTTTGATACCTTCTTAGCCTGAGAAAGGTTTCTTGGCTTCACGGGCGGAATGAAAGGCGGCGGAGTCTCCTGCTGAGACGTTTGTGTCCTCCACTGAGGATTACTCTGGATACGGGAAGCTTTATTCCTGAGTTCACCAATTAAGAATTTACCCTCTGCACGGAACGCAACGATCTCCTTATCTGCCAATTCGATTGCCTGAGATAGCATTAAGTTAACTCCGATCAACAATTCATCAGCAGTGATTATTCCGTTTCGGTCTGGGGTCATTTCTACAGCCAGTTCTTTGGACTTTTCCGTGATCTCACCTGCCAGTAGACCTAATTCGTGACATTCTGCCAGCGTCTGGAGATCGATATCAGATGGTTCCGATTCTCTCCCGAGAAGGGTTGCTGTAGAGGAATCTGATCCGCCGCTAGCGAGATCGGTTGGATCGAAATCGGTCCCACTTCCGCTTTCCTGAGAGTAGTGCTGAGTCCGTTGCAACTGTAGGTTTCTCTGCGAGCTAGTAATGATACCGCTGGTTTGAACTGATACTGTAGCAGTCTCATTACCTTCTAACTTCGATTCCCGTGACGGCCCGAGCTGGACGATCTTACGATTGTTTTGATCTGGAAACGTCTTAGGAATCACTCCAGTTGCAATTGGTGCCTCTTCATCTAGTCCAGGATCCAACCCGCCATCTCCAAGTGGATCCAACACCTCTCCCAATTTCTCCTCATTTAGAATTTTCTGAGGATCCAGGAGGCGACCCGGACGTGCATTTGGCTTTACTTTCAAGCTTGCACTAGCCGGTTTGTTCTTCTTGCTTTCCGTCTGTTTCCCTTTACCACGATTTCCCATGGCGAGTTCAGATTTTCCCTCTTTCTTGCCTGAAGTGTCTGGTACACTGGCAAACACATTCACATCTGAAAAGATTGCAGACTTATTCTCCATTTTGATATGTGTTAATGTTTT

>Manitoba_rhabdovirus_1_42-3728

ACGGACATTACGGATCTAGGAGCAGCGGTCCGGCCTTCTCATACAGCAGTTTTCCGATCGAATCATCCCGACAAGCAGGATTTTGCTCCCACTGCGTAGCCACCTTCTTGATGATGTGATCAGGTACCTCTCCCTGATTTTTCAGAATGTGTGCCAACCACAAGGAGGACTCCTTCTCCTGTGGCTCTCCTTCTGGCGGAGGTTGCGGATCATCCTCTTCTTCCAATTCGTCTTTCTCGCCATCCACTCCGAATTGCTGAGAGAGAGTCGCAAACTTGCCCATCACATAATGCATTACGACTGCATTGGAGATGATGTTGTTCACTTCTGGATCCCCAACTACTCGAGCTCGTTTGGATCTTTCGAGATTGGAGCTTACTCCCAACGTGTGATAGAAGAAATGAAGATCCAGGTTCACCGAGGCCGAATAGGGTGATTTAGAAGACAATCCCAAGTCCATGAAGTACATGGCATAAGATCTGGGATTGTCCATCTCCTCTCCTCCTTTGACAATTCTCTCGAATTGGTCGGCACATCTGTCGGTCCAGATCCACTCCGCGAATGCAGGGAACGTCAATCCCAGACACTTGGTAATCATTCGGGTGGCAATGAGAGCTGCGCAATCCTTGAATCTGGTGACAATAGTTCCGATCCGAGCTTGAGAGTAGGGATGAAATGGAAACTCATTCAGGAACATATCAATTGCGGCCATGATCTGCACATATGGTTGATAGGCAATCCACTGCTTGCAGGCATCAATTATGTCTCCCAATTCTGCGGTTTGATTCAACCCCAGCGGTACCAGAAGGTTCTTCACATTCTTCAGGACTTGTTCCTTGTACTCTACCCGTGCAATCTGTCCAACTCGGTAAACGCTGCAGATCAGTATGATGCCCCGAAGAATCTCCTGCTCGGTCAAGAGGACGTTTCCCTCGACTACAGCTGGCACCGATTCCCTTTCGTCCGGAATGATGATCGCACTTAATCCAACGCGGTCTGTTCGCTTTCCGATGTCGCGATTGAATGAAGACCAATCCTGATTGAGGAGTCCCTTGACCAACTCGAACTCGTCGTATAGGAAGCGTACCACGAATCGATGGTCGACTTTGCCTTTGGTTAGTCCTGATCTTATCGCGGCATGAATCTGTTCTCTTGTCGCACCTGTCCCGGGTATCATCATTATAGTCGGCTTAGTATTACCCGGCTTATTAAACCAGGCCGACGGATACTCAATGGATCTCTCAGTTCCTATGTCAAAATGAACGATAGGAGTAGCGGTCGCAGGATTGTCAGTGATGCGTTTGATTACACGTGTCTCAGCCATTGAATACAAACAAAATCGATCTGATACTTTCCTCAGAGGTACTTACACAATTCTGACTTGAAGATGTGTTGACGGTACTAGTCG

>Manitoba_rhabdovirus_1_20-9066

GAAGCACGCTAGTCTGCAATAGTTTTATGATAAGCTGTTTTAGTCCTGAGTCAATTCTTTTTTATAGAAATACACTCTTTATCGCATTCCACCGTTATAGCATACGGTTTAAAGAGTGATAACTTAATAGTATGCCTCTTACGATATTTATTAGGCGAACGAATGCGAATATAATCCAAAAGAGAGGCATGACTCATAGTACTTTCTTCAGACGTTAGATGGAAGGACACCAGTATATTTGGGATTAATCCTATACCCTCACCATCATAATCATATGTAAGAGGGAATATGGGTGAGATGTTGCACAATTTAATTTTGAAAACTCCGGTTACGGTACCACGATAGTGATAAAAGGCTCCAACTTTATGTATGTATTTGCAAGTGAACCCAGCCAATATGTAAAACAAAAAGATATACTCCTTGTCATTGACATAGCCTTTATAATCATCCTTGATAATCTCTAATGACGCCATGAGTTCTTTGTAAGTCCTAGGAGGGTCACGGAATTTTATATCTAAGAGCGCATCGACTCTAACAGTCTTAATAAGAGACATTGTTACTCAAGATTGCAATACCGAGTTTTTTTCTCGATCTGTGTTACTGTTTTGTGATTTTGATCGTTATCTCTCGGTAACGGCATGTCAAAACGAATTGATTGTAGGCCTTATGTGCTCTCAGAATTGCCTTCCCCTGTTCTGGTTCAGTGTCGTAGGACTCCAAGATTTCTAAGCAATCCTCATGTGGGTACTCGACAATGATCGGTTTGCCACCCTTCTTACGGGGCATTACCACCTGGACTTTCCCTCCTCTCTGAACCGTAACTGGAGGTTGCGGATCCGTGGGCTTCAACTCGAGTTGAGGATTTCTCTCCGGTTTTGATACCTTCTTAGCCTGAGAAAGGTTTCTTGGCTT

>Manitoba_rhabdovirus_1_37-8088

ATCACACCCGGATCAGCCATCAGAGACTCCGACTCCGCCCACAAATCTGAGGTCCAGGTCTCATGCTCCATCTGATGCACCTCTACTTCAAATGCATCCAACTCGTTTCTCTCAACATCCGCGAACCGATGACCGTACATGTTCGACATTTTGAATGACAACTATTCGACCGTGATCACTTTCGACACTAGCAGTGGAATTGTGTTAATGTTTTTTTCACGGTCAAGGCTGGTAAAATTCTGTTTCTTGATTGCTTGAAGTCTTTGTAAACAATCTTATTAACTGATACCTTTTATTCTTAAAACAGTGGAGGAGTTTTGACCAAAAGATTAGAATCAAAACGCCGATTAGAACTGTTACAGCGCATGCTGTTAATGTATATAACATAGGTGTCCAGTGCAAATCTGTCAAAGAGCTATCATCCACGAAATTATAATCTGTCTCTGTACCGGGGTGTAGCTGTCTTCTCCGGTATTCGGGAGACCGCTTTGCCATCGGTACTCTGTAGTGAGAGCTTAGTCGTGCGATTCTTTTATAGTTTTCTATGTCTTGGATGCCAAGGATCAGTGTCTTGTTCTTGACATACAACCCATTGGGTCCATCGATCACATCTGTTTTATTTGATGTAACCCAGTACGGCCATGTCCAAAGTTGGTTGTCTGATCTTCTTATACTAATATACGGAAAACTCAGGCTCGGTTCGATAGTCACTTGTTGGTATAAGCTTAGTCCCATCTCAAATTTTCCTGGACTATAGCGATAGACTGGATGAAATCCGGGATAACGAGGTGTCAATGTTTGCAACTCCGCTCGACTAACCATTTCTCCTGCTATGATTTTATCGACAACTATCTCACACTCATGATCTAAAAATTCATCCAACAAACTGAGGACAGCGTTCTGTATAAATTCTTTGTCATTAATTATGTTGACTTGTGACCCAATCGGGCAGTCCGGAACTTGGTAAAAGTACTCCCCTATCCAGTCCTGGTGCGGGATGTTTTTCCGGTCCAGTGCCATCCAGTCCCCAGAGGGATATCTCAACCCTCGGTGGCCGCAGAAATTCATTGTACATGGTTTGTCATACTCGGATATGTATATATCAGGACTCCAACTAGCTACTGTATGATTGGGAGTGTAATATATCACTAAGGATCCATCCAACATATGAGAATCATCACATGAATCTTTTATCGAGTGATCCGATATCCAAAGACGATTGTAAAAATGTGTGATACAGACAGAGGACGAACAGGTTCCGTCCAAGAAGATCGAGGATTTGAAGGCATCAGAATACGGATTG

>Manitoba_rhabdovirus_3_12-867

TGTCTTATTTTGTCAGTTCAGAAAGATTCATTTTGATCACCCTGTACATTCTCGGTCGATGACAATTTATGCATAGTTGGCGATTAGAAGTTATGTTTGTGGTTAATAATAGTCAAGGAAATCAGAGCAGGAAGTGAAATTAGTCATAAATCTCTGAGTCTGTAACCTCTTTTATATCCTCTTTAGACGAAGCTTGAGAAGAGCCGTCCGAAACAGGAGGTTTGAATGTTAGTAGTCTAATAGCTTCTGCACAGCTATTGAGAGCTGCAGTTTGCTTTTCCAATTGATTTACAATTTTGGTCAGCAACAGTTGATTCTTCAATGATTGAGTCACCATATGGTCGAGATATAGTTCGTTCTTGACACATTTAGCTGGGTGATTAAATGGGCAATCTATCCCCGACTTGCAGAACCCTGATGTTTTGATGTTAACACAGGTAATGATCTTGTTGTCAGGGTTTAGGCTCTCGTTTGGAGAGAGTGCTTTCTTGGGACTCATGACTCGGCGGATTATGATCTCGGATACCCCTCGGAGTGATCAGAAAATATAGATAGCTATCGAAAGAAGAGTGTTCTTTATGTTAATA

>Manitoba_rhabdovirus_3_43-14258

GTCTTATTTTGTTAGTTCAGAAAGATTCATTTTGATCACCCTGTACATTCTCGGTCGATGACAATTTATGCATAGTTGGCGATTAGAAGTTATGTTTGTGGTTAATAATAGTCAAGGAAATCAGAGCAGGAAGTGAAATTAGTCATAAATCTCTGAGTCTGTAACCTCTTTTATATCCTCTTTAGATGAAGCTTGAGAAGAGCCGTCCGAAACAGGAGGTTTGAATGTTAGTAGTCTAATAGCTTCTGCACAGCTATTGAGAGCTGCAGTTTGCTTTTCCAATTGATTTACAATTTTGGTCAGCAACAGTTGATTCTTCAATGATTGAGTCACCATGTGGTCGAGATATAGTTCGTTCTTGACACATTTAGCTGGATGATTAAATGGGCAATCTATCCCCGACTTGCAGAACCCTGATGTTTTGATGTTAACACAGGTAATGATCTTGTTGTCAGGGTTTAGGCTCTCGTTTGGAGAGAGTGCTTTCTTGGGACTCATGACTCGGCGGATTATGATCTCGGATACCCCTCGGAGTGATCAGAAAATATAGATAGCTATCGAAGGAAGAGTGTTCTTTATGTTAATATCT

>Manitoba_rhabdovirus_3_30-6143

GTCTTATTTTGTTAGTTCAGAAAGATTCATTTTGATCACCCTGTACATTCTCGGTCGATGACAATTTATGCATAGTTGGCGATTAGAAGTTATGTTTGTGGTTAATAATAGTCAAGGAAATCAGAGCAGGAAGTGAAATTAGTCATAAATCTCTGAGTCTGTAACCTCTTTTATATCCTCTTTAGACGAAGCTTGAGAAAAGCCGTCCGAAACAGGAGGTTTGAATGTTAGTAGTCTAATAGCTTCTGCACAGCTATTGAGAGCTGCAGTTTGCTTTTCCAATTGATTTACAATTTTGGTCAGCAACAGTTGATTCTTCAATGATTGAGTCACCATGTGGTCGAGATATAGTTCGTTCTTGACACATTTAGCTGGGTGATTAAATGGGCAATCTATCCCCGACTTGCAGAACCCTGATGTTTTGATGTTAACACAGGTAATGATCTTGTTGTCAGGGTTTAGGCTCTCGTTTGGAGAGAGTGCTTTCTTGGGACTCATGACTCGGCGGATTATGATCTCGGATACCCCTCGGAGTGATCAGAAAATATAGATAGCTATCGAAAGAAGAGTGTTCTTTATGTTAATATCTTAATGACTT

>Manitoba_rhabdovirus_3_26-13138

CTTAATTTATTTGTTAGTCTTATTTTGTTAGTTCAGAAAGATTCATTTTGATCACCCTGTACATTCTTGGTCGATGACAATTTATGCATAGTTGGCAATTAGAGGTTATGTTTGTGGTTAATAATAGTCAAGGAAATCAGAGCAGGAAGTGAAATTAGTCATAAATCTCTGAGTCTGTAACCTCTTTTATATCCTCTTTAGACGAAGCTTGAGAAGAGCCGTCCGAAACAGGAGGTTTGAATGTTAGTAGTCTAATAGCTTCTGCACAGCTATTGAGAGCTGCAGTTTGCTTTTCCAATTGATTTACAATTTTGGTCAGCAACAGTTGATTCTTCAATGCTTGAGTCACCATGTGGTCGAGATATAGTTCGTTCTTGACACATTTAGCTGGGTGATTAAATGGGCAATCTATCCCCGACTTGCAGAACCCTGATGTTTTGATGTTAACACAGGTAATGATCTTGTTATCAGGGTTTAGGCTCTCGTTTGGAGAGAGTGCTTTCTTGGGACTCATGACTCTGCGGATTATGATCTCGGATACCCCTCGGAGTGATCAGAAAATATAGATAGCTATCGAAAGAAGAGTGTTCTTTATGTTAATATCT

>Manitoba_tombus-like_virus_1_19-1219

TTTCCCCCGGTGATCAGCCGGTTCGTTAATGGTTTACTAGTGTGAGGATGAATCCGTGTATTATAGGAACAGTCACTACCGTTGCCAACACAGCATACAGGTTACCGCGCGTGTAGGCTGTGTAGACAACGGCAGCGGCCGTCCAGGGGGTGAAGACGTGTAGCAAATTTAGGAGTGTCACTACCGGGCGAATAAGGTCCCAAAGTCGGTGCAACACCCATTCAAGAACGGTCTCTAGAACATGTGTTATCGTGCCAAATATAGCAGATAAGCCAAGTGGAAGAGTGTTCATTGTGGCCAAATAATAGTCACAACGAAGAGCCTCAAAATCGGTCTGCAATTCATAGAATACAGACGGATAATGGTAACGTAGGAATTGTGTTGGGTCAAGAGATGGTTGTGGCAATTTGATGTCCGATTGCCTGGCCAACTGTGAGAACCACGTTACATTAGAGTACAACTCTTCCATGTTGTAGGGACCGTTTAAGTATGGGTGTAGTTTATTACCTGACCAGTACGGAATACAATAACGACCATCGGATGAGGTACCCAAAGTTGGTAAGATTAGGGTATCATTGAATCCAATGAGCGTTGCTGTATATTGCGCACTGGCAAAGTAATACACGTTCGGTAATCCAGGAAGCGCTTGCTGGCAAGAGTATACATCGATGTCAGAATTCATGGCTGAAAGAGGCATTGCGGGTATCTCTTTCAAATACATGACCTGGCACCATTCAGGAGCTATGACTTTGGACGCAGTCCTATATCTGTCCTTTATCGTTATCTTTGCCAGCGGTTTCCTGGAGCCGATACATACGCCGACTTTGGAAAACCAACCATTCACTAGCTCACGGTACACAACGTCCCCACAGTATGTTGGCAGTGAACAGTCAGCCCTAGAAAAGGTTGCCGTCACAATCTGCAAACACGCCGTGGGATCGTTCCCGGGCAATTGCACCCACAATCCATTCGGTGTTAAACTATGGATGAAATCTTCCAGAGTGTAGACCGAACTAAAGTGTGTATGCCTTCCGGTGGGAGACTGAACGAAGTTGGTTCGATCGTACTCGACCACCAGTTGTGCCATTGTGTATACCGCGAGTGAGAGAATGATTGTGATCCTCATGCTATGCTTTCGGCAGTGTCAAAACGTCGTAAGAAATTAGGATGATAGCTAGGATTGCCACTACAGCACGTACCTGCAGTGACCTAGCTTTGAAGAAAATGAGAGCTAGAGCCGCAACGCCGATAAAGTCTCGAAGGGCAGTTGTCTCCTTGGCTATCTCAAAGCAATAGAAGCCGGAGATTGCAACGCCAGCCAACTGATAGATCGACGGGAGAGAGAGTGCGGCGCCACAAAACGCGATGAACCCGTTTGTGCGGTTGGGTTTGTCTTTGAGGAAGTCACCCAGAGGCTTGAGGAAGGGTGCCGATTGAAGTTTGGCAATGAACTTAGTAAGAAGTGTTGGGTCGTGGTGGGAGTACAATGCCAGCCCAACTAGCAGAATAAACGGGGCGCTTGAAAGCGATCGCGAAATCACGTTGAAAGTCGTGTTGATTAGGTTAGATCCGGTACGTTGAACTCTACCACGTGCCATATTGGATCACCAAGATTGAAGTTAAACGATTCGTTATTTAATTGGGAAAACTTGGAGGACGCTTGCTACGTAGAAACCCTGAAGAATTG

>Manitoba_tombus-like_virus_1_10-512

GGGGCCCCTAAATGAACCCCGTGAGGGGTTTTAGGTGCAATTTTTTAGTGTCGGCTTGAGGCCCTCCTTGCGGATCCCCTTGCGGGGTTTCCCGACACTTGCCAGTGGTTGTGCAGCAACCATAATTCTGGACTTTTCCATTTCCGTTTGCTGGTGTCCTATACCGTTGTTGTATCCGTCGAGCTTGTCAGTTTTCGTTAGCGATTAATGGGTGATTAATGATGATGTCCTTAAAATTACAGCATATCCTTACCTAACTTACCTAGTTGTTTTTTCGACCACATTATGCATGGGTTTATTTGGGTTTTTGTATTTTTTTTTTTTTTTCCCCCGGTGATCAGCCGGTTCGTTAATGTTTTATTAGTGTGAGGATGAATCCGTGTATTATAGGAACAGTCACTACCGTTGCCAACACAGCATACAGGTTACCACGCGTGTAGGCTGTGTAGACAACGGCAGCGGCCGTCCAAGGGGTGAAGACGTGTAGCAAATTTAGAAGTGTCACTACCGGGCGAATAAGGTCCCAAGTTCGGTGTAACACCCACTCAAGAACGGTCTCTAGAACATGTGTTATCGTGCCGAATACAGCAGATAAGCCAAGTGGAAGAGTGTTCATTGTGGCCAGGTAGTAGTCACAACGAAGAGCCTCAAAATCGGTCTGCAATTCATAGAATACAGACGGATAATGGTAACGTAGAAACTGTGTTGGGTCAATAGATGGTTGTGGCAATTTGATATCTGATTGCCTGGCCAACTGTGAAAACCACGTTACGTTAGAGTACAACTCTTCCATGTTATAGGGACCGTTCAAGTATGGGTGTAGTTTGTTACCTGACCAGTGCGGAATACAATAACGACCATCGGATGAGGTACCCAAAGTTGGTAAGATTAGGGTATCATTGAATCCAATGAGCGTTGCTGTATATTGCGCACTGGCGAAGTAATAAACAGTCGGTAGTCCAGGAAGCGCTTGCTGGCAAGAGTATACATCGATGTCAGAATTCATGGCTGAAAGAGGCATTGCAGGTATCTCTTTCAAATACATGACCTGGCACCATTCAGGAGCTATAACTTTGGACGCAGTCCTATATCTGTCCTTTATCGTTATCTTTGCCAGCGGTTTCCTGGAGCCGATACATACACCGACCTTGGAAAACCAACCATTCACCAACTCGCGGTATACAACGTCCCCACAGTATGTTGGCAGTGAACAGTCAGCCCTAGAAAAGGTTGCCGTCACAATCTGCAAACACGCCGTGGGATCGTTCCCGGGCAGTTGCACCCACAATCCATTCGGTGTTAAACTATGGATGAAATCTTCCAGAGTGTAAACCGAACTAAAGTGTGTGTGCCGTCCGGTGGGAGACTGAACGAAGTTGGTTCGATCGTACTCGACCACCAGTTGTGCCGTTGTGTACACCGCGAGTAAGAGAATGATTGTAATCCTCATGCTATGCTTTCGGTAGTGTCAAAACGTCGTAGGAAATAAGAATAACAGCTAGGATGGCCACCACAGCACGTACCTGCAGTGACCTAGCTTTAAAGAAAATAAGAGCTAGAGCCGCAACGCCGATAAAGTCTCGCAGGGCAGTTGCCTCCTTGGCTATCTCAAAGCAATAGAAGCCCGAGATTGCAACGCCGGCCAATTGATAGATCGACGGTAGAGAAAGTGCGGCGCCACAAAACGCGATGAAGCCGTTTGTGCGGTTGGGTTTGTCTTTGAGGAAGTCACCCAGAGGCTTGAGGAAGGGTGCCGATTGAAGTTTGGCAATGAACTTAGTAAGAAGTGTTGGGTCGTGATGGGAGTACAATGCCAGCCCAACTAGCAGAATAAACGGGGCGCTTGAAAGCGATCGCGAAATCACGTTGAATGTCGTGTTGATTAGGTTAGATCCGGTACGTTGAACTCTACCACGTGCCATATTGGATCACCAAGATTGAAGTTAAACGATTCGTTATACAATTGGGAAAACTTGGAGGACGCTTGCTACGTTGAAACCCTGAAGAATTG

>Manitoba_tombus-like_virus_1_17-1214

GGGCCCCTAAATGAACCCCGTGAGGGGTTTTAGGTGCAATTTTTTTAGTGTCGGCTTGAGGCCCTCCTTGCGGATCCCCTTGCGGGGTTTCCCGACACTTGCCAGTGGTTGTGCAGCAACCATAATTCTGGACTTTTCCATTTCCGTTTGCTGGTGTCCTATACCGTTATTATATCCATCGGGTTTGTCAGTTTTCGTTGGCGATTAATGGGTGATTGATGATGATGTCCTTAAAATTACAGCATATCCTTACCTAACTTACCTAGTTGTTTTTTCGACCACATTATGCATGGGTTTATTTGGGTTTTGTATATTTTTTTTTTTTCCCCCGGTGATCAGCCGGTTCGTTAATTGTTTATTAGTGTGAGGATGAATCCGTGTATTATAGGAACAGTCACTACCGTGGCCAACACAGCGTACAGGTTACCACGCGTGTAGGCTGTGTAGACAACGGCAGCGGCCGTCCAGGGGGTGAAGACGTGTAGCAAATTTAGAAGTGTCACTACCGGGCGAATAAGGTCCCAAGCTCGGTGCAACACCCATTCAAGAACGGTCTCTAGAACATGTGTTATCGTGCCGAATACAGCAGATAAGCCAAGTGGAAGAGTGTTCATTGTGGCCAAGTAATAGTCACAACGAAGAGCCTCAAAATCGGTCTGCAATTCATAGAATACAGACGGATAATGGTAACGTAGAAATTGTGTTGGGTCAATAGATGGTTGTGGCAATTTGATGTCCGATTGCCTGGCCAACTGTGAAAACCACGTTACATTAGAGTACAACTCTTCCATGTTGTAGGGGCCGTTCAAGTATGGGTGTAGCTTATTACCTGACCAGTACGGTATACAATAACGACCATCGGATGAGGTACCCAAAGTTGGTAAGATTAGGGTATCATTGAATCCAATGAGCGTTGCTGTATATTGCGCACTGGCGAAGTAATACACGTTCGGCAATCCAGGAAGCGCTTGCTGGCATGAGTATACATCGATGTCAGAATTCATGGCTGAAAGAGGCATTGCAGGAATCTCTTTCAGATACATGACCTGACACCATTCAGGAGCTATGACTTTGGAGGCAGTCCTATATCTGTCCTTTATCGTTATCTTTGCCAGCGGTTTCCTGGAGCCGATACATACACCGACCTTGGAAAACCAACCATTCACCAGCTCGCGGTATACAACGTCCCCGCAGTATGTTGGCAGTGAACAGTCAGCCCTAGAAAAGGTTGCTGTCACAATCTGCAAACACGCCGTGGGATCGTTCCCGGGCAGTTGCACCCACAATCCATTCGGTGTTAAACTATGGATGAAATCTTCCAGAGTGTAAACCGAACTAAAGTGTGTGTGCCGTCCGGTGGGAGACTGAACGAAGTTGGTTCGATCGTACTCGACCACCAGTTGTGCCGTTGTGTATGCCGCGAGCAAGAGAATGATTGGAATCCTCATGCTATGCTTTCGGTAGTGTCAAAACGTCGTAAGAAATGAGGATGACAGCTAGGATGGCCACTACAGCACGTACCTGCAGTGACCTAGCTTTGAAGAAAATGAGAGCTAGAGCCGCAACGCCGATGAAGTCCCGTAGGGCAGTTGTCTCCTTGGCTATCTCAAAGCAATAGAAGCCGGAGATTGCAACGCCAGCCAATTGATAGATCGACGGGAGAGAGAGTGCGGCGCCACAAAACGCGATGAACCCGTTTGTGCGGTTGGGTTTGTCTTTGAGGAAGTCACCCAGGGGCTTGAGGAAGGGTGCCGATTGAAGTTTGGCAATGAACTTAGTAAGAAGTGTTGGGTCGTGATGGGAGTACAATGCCAGCCCAACTAGCAGAATAAACGGGGCGCTTGAAAGCGATCGCGAAATCACGTTGAAAGTCGTGTTGATTAGGTTAGATCCGGTACGTTGAACTCTACCACGTGCCATATTGGATCACCAAGATTGAAGTTAAAACGATTCGTTATACAATTGGGAAAACTTGGAGGACGCTTGCTACGTTGAAACCCTGAAGAATTGGATC

>Manitoba_tombus-like_virus_1_26-5

GGGGCCCCTAAATGAACCCCGTGAGGGGTTTTAGGTGCAATTTTTTAGTGTCGGCTTGAGGCCCTCCTTGCGGATCCCCTTGCGGGGTTTCCCGACACTTGCCAGTGGTTGTGCAGCAACCATAATTCTGGACTTTTCCATTTCCGTTTGCTGGTGTCCTATACCGTTGTTATATCCGTCGAGCTTGTCAGTTTTCGTTAGCGATTAATGGGTGATTGATGATGATGTCCTTAAAATTACAGCATATCCTTACCTAACTTACCTAGTTGTTTTTTCGACCACATTATGCATGGGTTTATTTGGGTTTTGTATTTTTTTTTTTTTTTCCCCCGGTGATCAGCCGGTTCGTTAATGTTTTATTAGTGTGAGGATGAATCCGTGTATTATAGGAACGGTCACTACCGTTGCCAACACAGCATACAGGTTACCACGCGTGTAGGCTGTGTAGACAACGGCAGCGGCCGTCCAAGGGGTGAAGACGTGTAGCAAATTTAGGAGTGTCACTACCGGGCGAATAAGGTCCCAAGTTCGGTGTAACACCCACTCAAGAACGGTCTCTAGAACATGTGTTATCGTGCCGAATACAGCAGATAAGCCAAGTGGAAGAGTGTTCATTGTGGCCAGATAGTAGTCACAACGAAGAGCCTCAAAATCGGTCTGCAATTCATAGAATACAGACGGATAATGGTAACGTAGAAACTGTGTTGGGTCAATAGATGGTTGTGGCAATTTGATGTCCGATTGCCTGGCCAACTGTGAAAACCACGTTACGTTAGAGTACAACTCTTCCATGTTATAGGGACCGTTCAAGTATGGGTGTAGTTTGTTACCTGACCAGTACGGAATACAATAACGACCATCGGATGAGGTACCCAAAGTTGGTAAGATTAGGGTATCATTGAATCCAATGAGCGTTGCTGTATATTGCGCACTGGCGAAGTAATAAACAGTCGGTAATCCGGGAAGCGCTTGCTGGCAAGAGTATACATCGATGTCAGAATTCATGGCTGAAAGAGGCATTGCAGGTATCTCCTTCAAATACATGACCTGGCACCATTCAGGAGCTATGACTTTGGACGCAGTCCTATATCTGTCCTTTATCGTTATCTTTGCCAGCGGTTTCCTGGAGCCGATACATACACCGACCTTGGAAAACCAACCATTCACCAACTCGCGGTATACAACGTCCCCACAGTATGTTGGCAGTGAACAGTCAGCCCTAGAAAAGGTTGCCGTCACAATCTGCAAACACGCCGTGGGATCGTTCCCGGGCAGTTGCACCCACAATCCATTCGGTGTTAAACTATGGATGAAATCTTCCAGAGTGTAAACCGAACTAAAGTGTGTGTGCCGTCCGGTGGGAGACTGAACGAAGTTGGTTCGATCGTACTCGACCACCAGTTGTGCCGTTGTGTACACCGCGAGTAAGAGAATGATTGTAATCCTCATGCTATGCTTTCGGTAGTGTCAAAACGTCGTAAGAAATAAGAATAACAGCTAGGATGGCCACAACAGCACGTACCTGCAGTGACCTAGCTTTAAAGAAAATAAGAGCTAGAGCCGCAACGCCGATAAAGTCTCGAAGGGCAGTTGCCTCCTTGGCTATCTCAAAGCAATAGAAGCCCGAGATTGCAACGCCGGCCAACTGATAGATCGACGGTAGAGAAAGTGCGGCGCCACAAAACGCGATGAAGCCGTTTGTGCGGTTGGGTTTATCTTTGAGGAAGTCACCCAGAGGCTTGAGGAAGGGTGCCGATTGAAGTTTGGCAATGAACTTAGTAAGAAGTGTTGGGTCGTGATGGGAGTACAATGCCAGCCCAACTAGCAGAATAAACGGGGCGCTTGAAAGCGATCGCGAAATCACGTTGAATGTCGTGTTGATTAGGTTAGATCCGGTACGTTGAACTCTACCACGTGCCATATTGGATCACCAAGATTGAAGTTAAACGATTCGTTATACAATTG

>Manitoba_toti-like_virus_1_23-3712

AATATCTTCAATAATTGTTGGATTCTCTGTCCTTTAACTATATAATATATTTTATAAGCACTTATTCTTCAAAGACGACGCTAGTAGAGCTGTCTAGGTTCAGCTCTTTGCCGATGTCTACACCATGGCCATAGAAGACATCGTCTATATCGGCCCACTGCCTAGTAATGGGTGTTATCTTTGCCTCGACGAGGGCACGATTCACAACGTCCTTGAACCTTTTGTATTTATCAGGTCCGTGCCCATGGGCCTGCAACAGAGCAGCTTCACAGTTGACTCTAGTGGAATCTTTAAGAGATGCCCCTTTCCACACCCACTGTGTTGTGCTAACAATTGATTCCCAAGCAAGGGGGGATAAAAACTCTCCCTTACGCGTTGGGTGCAGCACGAATTTTCTTTTAAGAAACGATGCTTCTGACATATGCACCCATGCTGTTATGGCGGCGGTTTTCTCAGCAGACGTCGAAACTATGTTGTGCGACGCGAAGTATTGGTGTATAGTTATACCATTAAAATCGCCTATATAGTCATCGCTAACCGTCATTATGGCATCATCCCCATATACTATGAGGGCGACATTAGCCTTAAATGTCTCCACCCAATCGTTGCCGCCGACTATGGACTTCCATGCTAACATAATGTACATCTGGTTAACCATACTATTGACTATGGTTGTGAAGAAAGCCCCACTTGGCGAACCACCTAGTTGGCTATAAACAGTATTGTGGCATATATGTTTGCTTTGAATGCACTCGTAAACTAAAGACCACATCTCTGTCTCATCAACACCTTGAACATAGTTCTTAGTCCAATCTATCATGAGAGAGTAGGCGGCTTTAGCTACTCCAGCATTGTATCCAGGCCCGAAATTCGAGTAGTCTATTGTGATGAACTTTGTATTGATCCTAGCCAATCGCTTAATCATATCAGACCACTCTTCACTTAGCGGATTGCATCCAACGCCGTGCATCAAATTGCGGCGGGCGCTCATAAAAGAGGCAACAAAGTGTGAGAACGTCTGGCGACATGCTATAGTATAATCGGCACCCGGATTACATATTACACGTGTACCCCCCCTCTTCTTAATCTTTTCCAATGACCTTTTCTCGTCCTTCAAAGTATCGATGTAGAGAGTCTCAGGTATTATTCCATTTCTACGCAAAAGCTCTTTACGAGCTATCTCGTCAGAAAGAGCGGGATCAATCCATTCCACACCTCTAACACTACCGTCACCATTACGTGCGTATGTAATGTAATCTTCTTTAGCTGTCTTCCGAGTGAGCACATAGGGAAAACCCATGCTCGTGGACAAATCCATCGGCTTGTAGTAATCAATACCCAATCCAGTGATTGCTTTATTCACTGTCAGGCGTTCTGGATTCATCACCAATGGTTTCATTGCGCAATACCACCCATCCCACAATGCCTCCTTAACACAAGTAAGCTCTTCACTAGTGAAGTCTGTGGTCAATAAGCCGTGTTTTCTCACTCCCTCAAACAACGGAGTGGTGTCAAAGTCGTACCGCGCATCCTTCTTGTGTAAAACAGCAGGTTGCATGTCGGTGCTCATGCCACAACTGTTATAAACCAAAGACTTAACTAGCCGTGTTGTCTGAGGTAAGTAGGGTACTTTCTCCTTAGGAAGTGAACCCCCATAATCTAGACGAACTTCCATATCGTATATTATGGAAGCGTCCATGATAGTACCGTGCTCAACTTCTTCCTTGGTGAGCACAATATCGGACTGTGTAGTGGGCAAGAGCGCCTCTTTTGTAAGGACTATGCCATAACCTGTCCCCTTATGCTCTTGACCAAGCCCAGCAACATGCATAGACATAATGGGGCGATTACTACCCTCCCTAAGTAGCAATGATCCACACGCTCCGGGTCTAGAGTAGGTGTACTCTAAACAATCGCGTATCTCAAAGCATTTGTTCTCCTCATCAGCTACAACGTAAGTATCGAGGAAACCTATAAGATCTACCTCGATAAGTACCATGTAGTCTTCACCACGCCTAGGTACTGAGAACAAAGTGGCGTTGGCTGGTATAGATGCCTGCATTTCGTCCTCCGTAGCAAGGAACTTGCGCAAGTCCTTGAATAGGGGTTGCGATGGTTGCAACTTGAGCATAGCTATATCGGTATGCGTGGACTCCGTGATATCTTTCTCACATATCACTAGGAGCGACCTCAACTGGGGTTTGTGGAGGGGGTTACCGTATATGTTGTAACCCATTTCCATTGCTTTCCTGATTTCGTGTATGTAGTGCCTAGGTAGAAGCAAATTGTGATTGTACAACCCAACACCGTACAATACGCGTTCCTTGTTACTACCTCTATCTATAGAGATACGGAATGTGTTATTGGCTATGTACGACTTAGCCACGGCGTACGGAGTAGGAGTTTCCTCGCCTCCTTGGAAGTACCTGACTTGTCTCTGCCTATCATAGCGTCGGCGCATTACTCGGTGCGGTTGATTCGAATCATATTCATCAGATTGGGGTAGAGCCACTACGCTCTGTGCTGTGCAACAAATCTTATATATGATTGTCGCTATTGCTGTAGCACCAACAAGACCGGCCAGTACGTAGCTGTACTTGTGGAGGGTCCCTGTTAGGTATTCCCACCACGCTTTTGGTAGTGCCACATTACTCGGCTCCACCACCCAAGCAGGTCTAAATAGTGCCGGTACAGCCCTGCACGTTTCTCCCGAAGGATTGTTAATGTAGTTAACTATAGCGGTGCGCAATACACGCGCATTAGCCTCCATGTACCTAGTAACAAAAGTAGCATACTGGGTACTGCGTGCGTAACAGCTTTCGGAGTTGCACGACGCCATCGGTACGTCAACCATAAGGTTGGTTTGCATATTGGGCACACGCCACATATTGCCGTATATCAAAGGCGACACACGCGGGTTAGACAACAACCCATGAGTACAAACAGGCAAAGGAGCACAAGCTTGCATCAAATCCTCTAGTAACTGCACGTTTACAACAGGCTCCATCAAGTCCTGGGTGGTTTGTGCGTCCACATAATCAGACTGGAATACAGCTTGCGCAGCAAATCCCATAAAATGGCCCGTAGCATAGCTCTCGTAGGCGTCCAACCTCCCTGTAATAAGGGACGCTAGATAGTACAAGACTGCCGTCAGTGTAGGCAACGATGCCCCCTCATATATGGTGGAAAGGTGCTTACCCAACCATGCTATCGGCATGCACCCGCTAGAAATTAGTCGAGCTGCCGCATATATAGAGTAGCCTTGCTCAGGTGTAACGCAAGGTATGACCACACCGTCACGACAAGTGGGGCAGCCAGAAACGCGTTCTGCCATCAAATTATTGTAACATGCAGTACATAAGTAGTGTGGGGATTCCAAGTCCACAGAGTTAGAGCACGCATATGCCACTACTCTGGTGTCTAAGCACACGTTACACCTGCTAACGGTATTGGAGGTCTTATTGACGTAGTAATCCGAGATGGAGCGC

>Manitoba_toti-like_virus_1_36-7

TAGTCCAATCTATCATGAGAGAGTAGGCGGCTTTAGCTACTCCAGCATTGTATCCAGGCCCGAAATTCGAGTAGTCTATTGTGATGAACTTTGTATTGATCCTAGCCAATCGCTTAATCATATCAGACCACTCTTCACTTAGCGGATTGCATCCAACGCCGTGCATCAAATTGCGGCGGGCGCTCATAAAAGAGGCAACAAAGTGTGAGAACGTCTGGCGACATGCTATAGTATAATCGGCACCCGGATTACATATTACACGTGTACCCCCCCTCTTCTTAATCTTTTCCAATGACCTTTTCTCGTCCTTCAAAGTATCGATGTAGAGAGTCTCAGGTATTATTCCATTTCTACGCAAAAGCTCTTTACGAGCTATCTCGTCAGAAAGAGCGGGATCAATCCATTCCACACCTCTAACACTACCGTCACCATTACGTGCGTATGTAATGTAATCTTCTTTAGCTGTCTTCCGAGTGAGCACATAGGGAAAACCCATGCTCGTGGACAAATCCATCGGCTTGTAGTAATCAATACCCAATCCAGTGATTGCTTTATTCACTGTCAGGCGTTCTGGATTCATCACCAATGGTTTCATTGCGCAATACCACCCATCCCACAATGCCTCCTTAACACAAGTAAGCTCTTCACTAGTGAAGTCTGTGGTCAATAAGCCGTGTTTTCTCACTCCCTCAAACAACGGAGTGGTGTCAAAGTCGTACCGCGCATCCTTCTTGTGTAAAACAGCAGGTTGCATGTCGGTGCTCATGCCACAACTGTTATAAACCAAAGACTTAACTAGCCGTGTTGTCTGAGGTAAGTAGGGTACTTTCTCCTTAGGAAGTGAACCCCCATAATCTAGACGAACTTCCATATCGTATATTATGGAAGCGTCCATGATAGTACCGTGCTCAACTTCTTCCTTGGTGAGCACAATATCGGACTGTGTAGTGGGCAAGAGCGCCTCTTTTGTAAGGACTATGCCATAACCTGTCCCCTTATGCTCTTGACCAAGCCCAGCAACATGCATAGACATAATGGGGCGATTACTACCCTCCCTAAGTAGCAATGATCCACACGCTCCGGGTCTAGAGTAGGTGTACTCTAAACAATCGCGTATCTCAAAGCATTTGTTCTCCTCATCAGCTACAACGTAAGTATCGAGGAAACCTATAAGATCTACCTCGATAAGTACCATGTAGTCTTCACCACGCCTAGGTACTGAGAACAAAGTGGCGTTGGCTGGTATAGATGCCTGCATTTCGTCCTCCGTAGCAAGGAACTTGCGCAAGTCCTTGAATAGGGGTTGCGATGGTTGCAACTTGAGCATAGCTATATCGGTATGCGTGGACTCCGTGATATCTTTCTCACATATCACTAGGAGCGACCTCAACTGGGGTTTGTGGAGGGGGTTACCGTATATGTTGTAACCCATTTCCATTGCTTTCCTGATTTCGTGTATGTAGTGCCTAGGTAGAAGCAAATTGTGATTGTACAACCCAACACCGTACAATACGCGTTCCTTGTTACTACCTCTATCTATAGAGATACGGAATGTGTTATTGGCTATGTACGACTTAGCCACGGCGTACGGAGTAGGAGTTTCCTCGCCTCCTTGGAAGTACCTGACTTGTCTCTGCCTATCATAGCGTCGGCGCATTACTCGGTGCGGTTGATTCGAATCATATTCATCNNGTATTCCCACCACGCTTTTGGTAGTGCCACATTACTCGGCTCCACCACCCAAGCAGGTCTAAATAGTGCCGGTACAGCCCTGCACGTTTCTCCCGAAGGATTGTTAATGTAGTTAACTATAGCGGTGCGCAATACACGCGCATTAGCCTCCATGTACCTAGTAACAAAAGTAGCATACTGGGTACTGCGTGCGTAACAGCTTTCGGAGTTGCACGACGCCATCGGTACGTCAACCATAAGGTTGGTTTGCATATTGGGCACACGCCACATATTGCCGTATATCAAAGGCGACACACGCGGGTTAGACAACAACCCATGAGTACAAACAGGCAAAGGAGCACAAGCTTGCATCAAATCCTCTAATAACTGCACGTTTACAACAGGCTCCATCAAGTCCTGGGTGGTTTGTGCGTCCACATAATCAGACTGGAATACAGCTTGCGCAGCAAATCCCATAAAATGGCCCGTAGCATAGCTCTCGTAGGCGTCCAACCTCCCTGTAATAAGGGACGCTAGATAGTACAAGACTGCCGTCAGTGTAGGCAACGATGCCCCCTCATATATGGTGGAAAGGTGCTTACCCAACCATGCTATCGGCATGCACCCGCTAGAAATCAGTCGAGCTGCCGCATATATAGAGTAGCCTTGCTCAGGTGTAACGCAAGGTATGACCACACCGTCACGACAAGTGGGGCAGCCAGAAACGCGTTCTGCCATCAAATTATTGTAACATGCAGTACATAAGTAGTGTGGGGATTCCAAGTCCACAGAGTTAGAGCACGCATATGCCACTACTCTGGTGTCTAAGCACACGTTACACCTGCTAACGGTATTGGAGGTCTTATTGACGTAGTAATCCGAGATGGAGCGCAATGCTTTCATCGAGTACTTCGCCACAGTTTTCCACATTCCGCTTCCAATAGCAAGCCCTAGTATAGTGTGACCCAAATTCTGCTGCCTAGGCAATTCCCAAGGCATAGCTTCAGGTTGAGATGGATTCTCAGGAGCTGGGGCAACAGAAACCTGGTTAACAATATTGCTAACCGCTATTTCTAGCTCTTCGTATGGAGTAAACGCATTTTGAGCGAGCGACTCATCCTCACGTATAGCAGTATTCAATTGGTAAAACAGGGTAAAGGGGTCATTTAACTGTATCTCCCTAACGTCCTGGGAACTCAAGGACGATAAAAGTCTATCCATCCGCTTTTGAACGTTGGCAACTTCCGTAGCATGATAGCGTTGGAAAGTATTCTGGAGGTATGACACAGTCTGTGAGAAGGATCTAATAGCTCCATCTAGCACTGTGCCGTCTTTCACGTTCGGGTACCTCCTAAACATGAGGTGGGGGAAGTCGTTTAGGGATCCTATAGGGAGGTCCCTCAAATTGACATTCCGATACTCCTCAGCAAGCTCAGCCCGCAAAACTGTATCTCGACGGCGATAAATAGCCTCAGGATACCTAGCGTAATTCGCTAAGTTTGGAAAAGCCCCGTTACACAATATTATAACGACCAAGGGATTACCTCGGCGCTTCTTCTCCTCCAGGTGTGCCATCTCAGGGATGAACAGTGACGTGGATTTCAATTTATTCAATTCCACTATCATATCGTTACATCTCTGGGAATCTTGGGAATTAAGAAACTCGTCATAGACTATCACTGGTTGATTACTATAACCCGACCAGAATTTCTCCCCAGCAGTCCTGTAATATATACTTGCTGACGAGGGACACTGATAACCGATAGACTTGAGGAGTTGTACTACAATTTCCTCACAGGCTGAACTCTTACCAATTCCAGGAGCCCCTTCTATACACAACACGTATGGTTCGTATCGCACCGGGGACGCTGATAAATCTGCAAACCTCTCATTTGCCACTTTTATAACGTCCGAGCACAACTTGGCTAACTGTGCGTTTCCCACGCCTGATGGAACAGAACATAAGAGTCGCTGGTATTGGTAAGCACTAAGTACAGTCTTCCAACACCGCAAGCGAAAGTTACCATTCGATAAGAGCGAAGTAGATGCTTCACTGGTGACTATCTGGGCTTCGCGTATGAAATGAGCTATCTGTTCACTCCCATCGGCTAGCATCTTGAGAGCACAAGCCTCTGGTGAGACATAACCAAGAGCATGCATGACATACTCCTTAATAATTTCGAATGTGGATTGCACATAACGCAATATAGACAAAAGATACGATATACCGGAGGTATTTGTGATTCGCTCCAATAGAGCTCCCGGCACACTACGCACACGCTTTGGGTCTATATAAACACCCATTAGCGTACCCACAATGCCGGCCAGGATACCAGTGAGCGTAGACTCATTAGAAGGGGGCCCTTGTGTGGTAGGAGTGTCCTGATTTAATTGTGCGTAAAACGCACCCAACTGTGGGCCAAAAGCCGATAAGGCGAGAGCTTGTGACGTGTCAAAGAAATGGGATATGAAACGCACCACGGAAACGCCTACTACAACCCACGATTTTGAGACCCATGCACTGACAATGTCTAAAAATAAGTCAAAAAATATCCTCGCGTACGAGGCCACGGAAGACGTAGTTGAAATAACTTTGTCTACAGCTGCTTTTATAATGCCCTTAAATTCGTCAACAGACGAAGACATACCGGACGCTAATCTCTCTAAGCTGGTAGCAGCTTGATCGATTTTCGGTACACTCTGGCCGAGGGCGTTACTTAAATTGCTACACGAGCGCATGGCGTTCCTCACTTCAGGTGCCAGCTCGTTCACAGCACTCGCGGCGACTAAGCCACCTCCAACGATTGGTATAGCGCCTAGTGCCATACGTGCCACTGAAGTTGGCGAAGTAGATTGTATAACATTATTTATAGCCCCTGTGATGGTATCCATCTGAGTTTTAGGGATACTGCGAACCGAAGTTCCAGGATCTTTATCCAGAAAAACCGCCGGTGACGCAGGTTTCTTAGTAACTCCATTTTCATCAGTATACACTGTGTACTGACCATAATCGATACAGTAGGGTATACCATGGAAATTGGCTACTTGGAAGTCATCCCCTGCTGACCACCACACTGTAACCTTAGCCTCAACTTCGGATGACAGTATTAAATGACCAGAGTTGTAGTTACCCTTATCACGCCAGGAGTAAGACAAGCCAGGGTTGTCCTCCTGCATAAGAGTCCAGACGTTTTCGGTGTCATACGGGACCTCAAAGGTTAACATGGGATTAACAGCCGGAACTATGACCTCGGTGACCATACCTACGCCCTCCAAAGGTGACCAGGAGATGTGGTTGTTATCACCGCTAGTCGCCTTCTGTATATAGGCGTGATTACCTATAATGCGAGCGCCTGTGTGTGGCATGTATGACACATAAACTGTGGCTCCGTTCCGCACTAAGATTGTGTAACGCATACTACCTCTCCACATACGAAACATATTGCATATATTGTACTGGGCAGTGTGTTGTACCCCAGGGTTAAATATAGCACTGGCTATGTGCGTCATAGGGTCATTAATCATAGACCTATTCGGAGGCATCAACGGTATAAAGAAAGCCGTAGCAGTATCACTATCGGGTCTCAAATCGATAGTGGCCCTAGTTATAAGGCAGGGCATCCTCAATCTGTCCTTGAAATCCATATGACAGTCCAATGTTTGGAGCTGCGTATTTACGCGATTGACGCGGAAATTGGGCGTGTCGTCTTGCGCTACAGCATCCATCTGTGTCCTAGGCATTTCATTCCACGCAGATCCTGTGGCCCTTGCAATAGCAACGAGCTCTCTCGAGGTCATCTCTATAGCAGGAGGATTGCGCTTGACAATCCTATCTCTGCTATAACCACCCCCAGGTAGCAGCGTGGGGTAATCACCCGGGAAATTATCGATGGGCTTAATAGTATTATATTCCCAGAAATGCCGCATGGACGCTGGCTTTATGCCATGCATAGCAAAATCCTCACCAGCTCTGATATATATAATACACTTAACCTCTTTGGCTACGACAGGGCTACAACGCAAAGGTGTGCTAACTCGTATTTTAAGCATGGCTTTGGTTTGTGCTGCAATCCCGATAGCAGACGCACAGACCCTTGGATCTGGCTTAAGCCCGTAAAAGAGCGGTAGAGGATCAACCGTTGACCTACGCATGATGGTATCATAAACATAGGGTATGGTGTACTCGCACACGTTACCGCTACTGAGGGGGTAGTACTTGGAGTAAGAACATGACGATGCCGACATATTAGACTCCTCACTAACACGGTTGAACTCCGTGCCTATCATGACTTCTCCACTATGGCCCTTGTTAGCAATAAAGCGGAGTTTGACCTCTATGGTACCAGACCAGAATTGAAAGTTGGTCATTGCGTACTCCAGAGGGGTAGGCTCGCCACGGTATGTTGTAGGCCGTAAATTACCACCATTATTGCGGCATGATGGATCTACAAAGAATTCGGCCAAAACAGCACTTACGGCATCGGTGGTTCTCCACTCAAACTGCTTATAGTAACCCCACTTCCGGGCTAGTTCCCAATAATTGCTTGGGTCGTTCTGCGGTATCTCGATCTTGTCGTAGTTAGTTTGGGTATAAGGGTTCAAACGCAGTGAAACTACGTTGATAGGTCCTTTCCCTGTTGCGAAATTCATAGCAGGGTGGGGGATAGCTATGGTAGACTGGGTGTCACTAGGCTTATCCCGGTTATGGGCTCTCCCTCGCACGTCGTAAGCATTTTCATACTTGCCGAGTATGCTCCTTAAGGCACCATTAGCAGCACCTGCAATAACATCAGTTATCCCCAACATCTGGACCACAGCTCTGGCAGTAAGGCCAGCAAAGTGAGCCCTTGGCATTCGTATGTATAGAGTGCCAGTGATGGAGGATGGACCATTCTCGCCAACGGCAAGTGGCGCAACACACATGACGGTTATAGAGCAAGACTCTGCTGGTCTAACACCTTTTGACACGCCAGGTGCATTTAGTAGCCTAATGAAGGGTGTGTGGTATATGTACGGGCAGTACAGAACTGCATCGTTGTCCTCCGATAGATCAACCTCGACGTGACGTCTAGCTTGCTCAGATTGATGTCCCGCCATGGCACGTTTACACTGGTACGAATCGTGCTTCACTCCCATGGTGTAAAATCCAGCAAACGCATCGTTACCAGGCACTACTAGGCGTACTTCGAAATCCATATTAGCGTAAGCATGGGTCTGGAAGGGAAGCAGAGCAGGGGAATTTGGCATCTGCGTGTACAAAAATTCCGGTAGCACACACGTAAAGATAACTTTACCTATGGCATCACTGGTAGAGAAACTAACCTGTTCCAGGTAAACCCACCTATTAACAAGATCGTCGTAAGTGGATATCTTCTCCGATGTTACAAATTCATCCAACGAAATATCGGGTTGCTCGGGTTGAGCAGTAATATCGGGCCCGGTGTGCACCATTATGGTGTTGCCAGAAACATCTGCCTCCGAATCCATCTGAACGCGAGGGATACACGCGCACTTCCTGATAGCAGGACAACCGTATCGACCTTTGTACGTGTTACGAAAATTGTACATCAAACGCCAGGACGCCTCGTTGGTGGCTGCCTTGACCGTAGTAGTAATGATATCAGCTTCGATTACTACAGGTCCCTGCAGCGTTGACCAAGTGAGGGATCTTCTACATCTTGCTGTAGCGGGGTCTCTTCCACGTCGCGTGGTGTTGGTGATTTGGCAACTTGCCGACAAATAAAAAGTAGCATTATACCAACTATTAAGGACAAGGGTATAGCTAGTGTTGCTATGATACGCCTCGTCAAAGGAGGCAGCGTATCTATCCAACGATTTAGTGAGCGCATCCATGCACTCACTCTTACTAGAGTAGACTCTAGCCTCAGCTCTCTCGGAACTGCTCTTCTCTCCGGATACAAATGGTCCCATGATGTAAAAGTTGAAACGTCGTTAGACGGGTAAACCAAGTTGTTATATGGCGTACCTTGAACTAAAGCTGAAGAACAATCTTCACTATTAGCTGGGACAGTGATGGTGCAAGTATTTGTCATACAATGACACGATATAGCACTGAAAATCCGGTTGTTCGCAAGAATAAAATAGCAAATTACACACGCAAATAACGAATGCGCCCACGGGTGTAGGACACAAAGGTAAATGGGAGGTGCAAAGGCTAAAATAATTGCTGCACGTTTTGAATGTTTATCATTCGGGGTTTTGTTGGTTGTTTGGCGTTAATAAACGCGTTGTTGTATGGTTCTTTCTCTGAAGAATCAAATCACGGTAATAAGGCACTTTCTCTGAAGTGTGGGCGGCGTACAATATGAAGAGTTGTTCTCTGACAAATCTTCGTATTT

>Manitoba_toti-like_virus_1_38-436

GTTGGATTCTCTGTCCTTTAACTATATAATATATTTTATAAGCACTTACTCTTCAAAGACGACGCTAGTAGAGCTGTCTAGGTTCAGCTCTTTGCCGATGTCTACACCATGGCCATAGAAGACATCGTCTATATCGGCCCACTGCCTAGTAATGGGTGTTATCTTTGCCTCGACGAGGGCACGATTCACAACGTCCTTGAACCTTTTGTATTTATCAGGCCCGTGCCCATGGGCCTGCAACAGAGCAGCTTCACAGTTGACTCTAGTGGAATCTTTAAGAGATGCCCCTTTCCACACCCACTGGGTTGTGCTAACAATTGATTCCCAAGCAAGGGGGGATAAAAACTCTCCCTTACGCGTTGGGTGCAGCACGAATTTTCTTTTAAGAAACGATGCTTCTGACATATGCACCCATGCTGTTATGGCGGCGGTTTTCTCAGCAGATGTCGAAACTATGTTGTGCGACGCGAAGTATTGGTGTATGGTTATACCATTAAAATCACCTATATAGTCATCGCTAACCGTCATTATGGCATCATCCCCATATACTATGAGGGCGACATTAGCCTTAAATGTCTCCACCCAATCGTTGCCGCCGACTATGGACTTCCATGCTAACATAATGTACATCTGGTTAACCATACTGTTGACTATGGTTGTGAAGAAAGCCCCACTTGGCGAACCACCTAGTTGGCTATAAACAGTATTGTGGCATATATGTTTACTTTGAATGCACTCGTAAACTAGAGACCACATCTCTGTCTCATCAACACCTTGAACATAGTTCTTGGTCCAATCTATCATGAGAGAGTAGGCGGCTTTAGCTACTCCAGCATTGTATCCAGGCCCGAAGTTCGAGTAGTCTATTGTGATGAACTTCGTATTGATCCTAGCCAATCGCTTAATCATATCAGACCACTCTTCACTTAGCGGATTGCATCCAACGCCGTGCATCAAATTGCGGCGGGCGCTCATAAAAGAGGCAACAAAGTGTGAGAACGTCTGGCGACATGCTATAGTGTAATCGGCACTCGGATTACATATTACACGTGTACCCCCCCTCTTCTTAATCTTTTCCAATGACCTTTTCTCATCCTTCAAAGTATCGATGTAGAGGGTCTCAGGTATTACTCCATTTCTACGCAAAAGCTCTTTACGAGCTATCTCGTCAGAAAGAGCGGGATCAATCCATTCCACACCTCTAACACTACCGTCACCATTACGTGCGTATGTAATGTAATCTTCTTTAGCTGTCTTCCGAGTGAGCACATAGGGAAAACCCATGCTCGTGGACAAATCCATTGGCTTGTAGTAATCAATACCCAATCCAGTGATTGCTTTATTCACTGTCAGGCGTTCTGGATTCATCACCAATGGTTTCATTGCGCAATACCACCCATCCCACAGTGCCTCCTTAACACAAGTAAGCTCTTCACTCGTGAAGTCTGTGGTCAATAAGCCGTGTTTTCTCACTCCCTCAAACAACGGAGTGGTGTCAAAGTCGTACCGCGCATCCTTCTTGTGCAAGACAGCAGGTTGCATGTCGGTGCTCATACCACAACTGTTATAAACCAAAGACTTAACTAGCCGTGTTGTCTGGGGTAAGTAGGGTACTTTCTCCTTAGGAAGTGAACCCCCGTAATCTAGACGAACTTCCATATCGTATATTATGGAAGCGTCCATGATAGTACCGTGCTCAACTTCTTCCTTGGTGAGCACAATATCGGACTGTGTAGTGGGCAAGAGCGCCTCTTTTGTAAGGACTATGCCATAACCTGTCCCCTTATGCTCTTGACCAAGCCCAGCAACATGCATAGACATAATGGGGCGATTACTACCCTCCCTAAGTAGCAATGATCCACACGCTCCGGGTCTAGAGTAGGTGTACTCTAAACAATCACGTATCTCAAAGCATTTGTTCTCCTCATCAGCTACAACGTAAGTATCGAGGAAACCTATGAGATCTACCTCGATAAGTACCATGTAGTCTTCACCACGCCTAGGTACTGAGAACAAAGTGGCGTTGGCTGGTATAGATGCCTGCATTTCGTCCTCCGTAGCAAGGAACTTGCGCAAGTCCTTGAATAGGGGTTGCGATGGTTGCAACTTGAGCATAGCTATATCGGTATGCGTGGACTCCGTGATATCTTTCTCACATATCACTAGGAGCGACCTCAACTGGGGTTTGTGGAGGGGGTTACCATATATGTTGTAACCCACCTCCATTGCTTTCCTGATTTCGTGTATGTAGTGCCTAGGTAGAAGCAAATTGTGATTGTACAACCCAACACCGTACAATACGCGTTCCTTGTTACTACCTCTATCTATAGAGATACGGAATGTGTTATTGGCTATGTACGACTTAGCCACGGCGTACGGAGTAGGAGTTTCCTCGCCTCCTTGGAAGTACCTGACTTGTCTCTGCCTATCGTAGCGTCGACGCATTACTCGGTGCGGTTGATTCGAATCATATTCATCAGATTGGGGTAGCGCCACAACGCTCTGTGCTGTGCAACAAATCTTATATATGATTGTCGCTATTGCTGTAGCACCAACAAGACCGGCCAGTACGTAGCTGTACTTATGGAGGGTCCCTGTTAAGTATTCCCACCACGCTTTTGGTAGTGCAACATTACTTGGCTCCACCACCCAAGCAGGTCTAAATAGTGCCGGTACAGCCCTGCACGTTTCTCCCGAAGGATCGTTAATGTAGTTAACTATAGCGGTGCGCAATACACGCACATTAGCCTCCATGTACCTAGTAACAAAAGTGGCATACTGGGTACTGCGCGCGTAACAGCTTTCGGAGTTGCACGACGCCATGGGTACGTCAACCATAAGATTGGTTTGCATATTGGGCACACGCCACATATTGCCGTATATCAAAGGCGACACACGCGGGTTAGACAACAACCCATGAGTACAAACAGGCAAAGGAGCACAAGCTTGCATCAATTCCTCTAGTAACTGCACGTTTACAACAGGCTCCATCAAGTCCTGGGTGGTTTGTGCGTCCACATAGTCAGACTGGAATACAGCTTGCACAGCAAATCCCATAAAATGGCCCGTAGCATAGCTCTCGTAGGCGTCCAACCTCCCTGTAATAAGGGACGCTAGATAGTACAAGACTGCCGTCAGTGTAGGCAACGATGCCCCCTCATATATGGTGGAAAGGTGCTTACCCAACCATGCTATCGGCATGCACCCGCTAGAAACTAGTCGAGCTGCCGCATATATAGAGTAGCCTTGCTCAGGTGTAACGCAAGGTATGATCACACCGTCACGACAAGTGGGGCAGCCAGCAACGCGTTCTGCCATCAAATTATTGTAACATGCAGTACATAAGTAGTGTGGGGATTCCAAGTCCACAGAGTTAGAGCACGCATATGCCACTACTCTGGTGTCTAAGCACACGTTACACCTGCTAACGGTATTGGAGGTCTTATTGACGTAGTAATCCGAGATGGAGCGCAATGCTTTCATCGAGTACTTCGCCACAGTTTTCCACATTCCGCTTCCAATAGCAAGCCCTAATATAGTGTGACCCAAATTCTGCTGCCTAGGCAATTCCCAAGGCATAGCTTCAGGTTGAGATGGATTCTCAGGAGCTGGGGCAACAGAAACCTGGTTGACAATATTGCTAACCGCTATTTCTAGCTCTTCGTATGGAGTAAACGCATTTTGAGCGAGCGACTCATCCTCACGTATAGCAGTATTCAATTGGTAAAACAGGGTAAAGGGGTCATTTAACTGTATCTCCCTAACGTCCTGGGAACTCAAGGACGATAAAAGTCTATCCATCCGCTTTTGAACATTGGCAACTTCCGTAGCATGATAGCGTTGGAAAGTATTCTGGAGGTATGACACAGTCTGTGAAAAGGATCTAATAGCTCCATCTAGCACTGTGCCGTCTTTCACGTTCGGGTACCTCCTAAACATGAGGTGGGGGAAGTCGTTTAGGGATCCTATAGGGAGGTCCCTCAAATTGACATTCCGATACTCCTCAGCAAGCTCAGCCCGCAAAACTGTATCCCGACGGCGATAAATGGCCTCGGGATACCTAGCGTAATTCGCTAAGTTTGGAAAAGCCCCATTACACAATATTATAACGACCAAGGGATTACCTCGGCGCTTCTTCTCCTCCAGGTGTGCCATCTCAGGGATGAACAGTGACGTGGATTTCAATTTATTCAATTCCACTATCATATCGTTACATCTCTGGGAATCTTGGGAATTAAGAAACTCGTCATAGACTATCACTGGTTGATTACTATAACCCGACCAGAATTTCTCCCCAGCAGTCCTGTAATATATACTTGCTGACGAGGGACACTGATAACCGATAGACTTGAGGAGTTGTACTACAATTTCCTCACAGGCTGAACTCTTACCAATTCCAGGAGCCCCTTCTATACACAACACGTATGGTTCGTATCGCACCGGGGACGCTGATAAATCTGCAAACCTCTCATTTGCCACTTTTATAACGTCCGAGCATAACTTGGCTAGCTGTGCGTTTCCCACGCCTGATGGAACAGAACATAAGAGTCGCTGGTATTGGTAAGCACTAAGTACAGTCTTCCAACACCGCAAGCGAAAGTTACCATTCGATAAGAGCGAAGTAGATGCTTCACTGGTGACTATCTGGGCTTCGCGTATGAAATGAGCTATCTGTTCACTCCCATCGGCTAGCATCTTGAGAGCACAAGCCTCTGGTGAGACATAACCAAGAGCATGCATGACATACTCCTTAATAATTTCGAATGTGGATTGCACATAACGCAATATAGACAAAAGATACGATATACCGGAGGTATTTGTGATTCGCTCCAATAGAGCTCCCGGCACACTACGCACACGCTTCGGGTCTATATAAACACCCATTAGCGTACCCACAATGCCGGCCAGGATACCAGTGAGCGTAGACTCATTAGAAGGGGGCCCTTGTGTGGTAGGAGTGTCCTGATTTAATTGTGCGTAAAACGCACCCAACTGTGGGCCAAAAGCCGATAAGGCGAGAGCTTGTGACGTGTCAAAGAAATGGGATATGAAACGCACCACGGAAACGCCTACTACGACCCACGATTTTGAGACCCATGCACTGACAATGTCTAAAAATAAGTCGAAAAATATCCTCGCGTACGAGGCCACGGAAGACGTAGTTGAAATAACTTTGTCTACGGCTGCTTTTATAATGCCCTTAAATTCGTCAACAGACGAAGACATACCGGACGCTAATCTCTCTAAGCTGGTAGCAGCTTGATCGATTTTCGGTACGCTCTGGCCGAGGGCGTTACTTAAATTGCTACACGAGCGCATGGCGTTCCTCACTTCAGGTGCCAGCTCGTTCACAGCACTCGCGGCGACTAAGCCACCACCAACGATTGGTATAGCGCCTAGCGCCATACGTGCCACCGAAGTTGGCGAAGTAGATTGTATAACATTATTTATAGCCCCTGTGATGGTATCCATCTGAGTTTTAGGGATACTGCGAACCGAAGTTCCAGGATCTTTATCCAGAAAAACCGCCGGTGACGCAGGTTTCTTAGTAACTCCATTTTCATCAGTATACACTGTGTACTGACCATAATCGATACAGTAGGGTATACCATGGAAATTGGCTACTTGGAAGTCATCCCCTGCTGACCACCACACTGTAACCTTAGCCTCAACTTCGGATGACAGTATTAAATGGCCAGAGTTGTAGTTACCCTTGTCACGCCAGGAATAAGACAAGCCAGGGTTGTCCTCCTGCATAAGAGTCCAGACGTTTTCGGTGTCATACGGGACCTCAAAGGTTAACATGGGATTAACAGCCGGAACTATTACCTCAGTGACCATACCTACGCCCTCCAAAGGTGACCAGGAGATGTGGTTGTTATCACCGCTAGTCGCCTTCTGTATATAGGCGTGATTACCTATAATGCGAGCGCCTGTGTGTGGCATGTATGACACATAAACTGTGGCTCCGTTCCGCACTAAGATTGTATAACGCATACTACCTCTCCACATACGGAACATATTGCATATATTATACTGGGCAGTGTGTTGTACCCCAGGGTTAAATATAGCACTGGCTATGTGCGTCATAGGGTCATTAATCATGGACCTATTCGGAGGCATCAACGGTATAAAGAAAGCCGTGGCAGTATCACTATCGGGTCTCAAATCGATAGTGGCCCTGGTTATAAGGCAGGGCATCCTCAATCTGTCCTTGAAATCCATATGACAGTCCAATGTTTGGAGCTGCGTATTTACGCGATTGACGCGGAAATTAGGCGTATCGTCTTGCGCTACAGCATCCATCTGTGTCCTAGGCATTTCATTCCACGCAGATCCTGTGGCCCTTGCAATAGCAACGAGCTCTCTCGAGGTCATCTCTATAGCAGGAGGGTTGCGCTTGACAATCCTATCTCTGCTATAACCACCCCCGGGTAGCAGCGTGGGGTAATCACCCGGGAAATTATCGATGGGCTTAATAGTATTATATTCCCAGAAGTGCCGCATGGACGCTGGCTTTATGCCGTGCATAGCAAAATCCTCACCAGCTCTGATATATATAATACACTTAACCTCCTTGGCTACGACAGGGCTACAACGCAAAGGTGTGCTAACTCGTATTTTAAGCATGGCTTTGGTTTGTGCTGCAATTCCGATAGCAGACGCACAGACCCTTGGATCTGGCTTAAGCCCATAAAAGAGCGGTAGAGGATCAACCGTTGACCTACGCATGATGGTATCATAAACATAGGGTATGGTGTACTCGCACACGTTACCGCTACTGAGGGGGTAGTACTTGGAGTAAGAACATGACGATGCCGACATATTAGACTCCTCACTAACACGGTTGAACTCCGTGCCTATCATGACTTCTCCACTGTGGCCCTTGTTAGCAATAAAGCGGAGTTTGACCTCTATGGTACCAGACCAGAATTGAAAGTTGGTCATAGCGTACTCCAGAGGGGTAGGCTCGCCACGGTATGTTGTAGGCCGTAAATTTCCACCATTATTGCGGCATGATGGATCTACAAAGAATTCGGCCAAAACAGCACTTACGGCATCGGTGGTTCTCCACTCAAACTGCTTATAGTAACCCCATTTCCGGGCTAGCTCCCAATAATTGCTTGGGTCGTTCTGCGGTATCTCGATCTTGTCGTAGTTAGTTTGGGTATAAGGGTTCAAACGCAGTGAAACTACGTTGATAGGTCCTTTCCCTGTTGCGAAATTCATAGCAGGGTGGGGGATAGCTATGGTAGACTGGGTGTCACTAGGCTTATCCCGGTTACGGGCTCTCCCTCGCACGTCGTAAGCATTTTCATACTTGCCAAGTATGCTCCTTAAGGCACCATTAGCAGCACCTGCAATAACATCAGTTATCCCCAACATCTGGACCACAGCTCTGGCAGTAAGACCAGCAAAGTGAGCCCTTGGCATTCGTATGTATAGAGTGCCAGTGATGGAGGATGGACCATTCTCGCCAACGGCAAGTGGCGCAACACACATGACGGTTATAGAGCAAGACTCTGCTGGTCTAACACCTTTTGACACGCCAGGTGCATTTAGTAGCCTAATGAAGGGTGTGTGGTATATGTACGGGCAGTACAGAACTGCATCGTTGTCCTCCGATAGATCAACCTCGACGTGACGTCTAGCTTGTTCAGATTGATGTCCCGCCATGGCACGTTTACACTGGTACGAATCATGCTTCACTCCCATGGTGTAAAATCCAGCAAACGCATCATTACCAGGCACTACTAGGCGTACTTCGAAATCCATGTTAGCGTAAGCATGGGTCTGGAAGGGAAGCAGAGCAGGGGAATTTGGCATCTGCGTGTACAAAAATTCCGGTAGCACACACGTAAAGATAACTTTACCTATGGCATCACTGGTAGAGAAACTAACCTGTTCCAGGTAAACCCACCTATTAACAAGATCGTCGTAAGTGGATATCTTCTCCGATGTTACAAATTCATCCAACGAAATATCGGGTTGCTCGGGTTGGGCAGTAATATCGGGCCCGGTGTGCACCATTATGGTGTTGCCAGAAACATCTGCCTCCGAGTCCATCTGAACGCGAGGGATACACGCGCACTTCCTAATAGCAGGACAACCATATCGACCTTTGTACGTGTTACGAAAATTGTACATCAAACGCCAGGATGCCTCATTGGTGGCTGCCTTGACCGTAGTAGTAATGATATCAGCTTCGATTACTACAGGTCCCTGCAACGTTGACCAAGTGAGGGATCTTCTACATCTTGCTGTAGCGGGGTCTCTTCCACGTCGCGTGGTGTTGGTGATTTGGCAACTTGCCGACAAATAAAAAGTAGCATTATACCAACTATTAAGGACAAGGGTATAGCTAGTGTTGCTATGATACGCCTCGTCAAAGGAGGCAGCGTATCTATCCAACGATTTAGTGAGCGCATCCATGCACTCACTCTTACTAGAGTAGACTCTAGCCTCAGCTCTCTCGGAACTGCTCTTCTCTCCGGATACAAATGGTCCCATGATGTAAAAGTTGAAACGTCGTTAGACGGGTAAACCAAGTTGTTATATGGCGTACCTTGGACTAAAGCTGAAGAACAATCTTCACTATTAGCTGGGACAGTGATGGTGCAAGTATTTGTCATACAATGACACAATATAGCACTGAAAATCCGGTTGTTCGCAAGAATAAAATAGCAAATTACACACGCAAATAACAAATGCGCCCACGGGTGTAGGACACAAAGGTAAATGAGAGGTGCAAAGGCTAAAATAATTGCTGCACGTTTTGAATGTTTATCATTCGGGGTTTTGTTGGTTGTTTGGCGTTAATAAACGCGTTGTTGTGTGGTTCTTTCTCTGAAGAATCAAATCACGGTAATAAGGCACTTTCTCTGAAGTGTGGGCGGCGTACAATATGAAGAGTTGTTCTCTGACAAATC

>Manitoba_toti-like_virus_1_22-1695

ATAGAATATCTTCAATAATTGTTGGATTCTCTGTCCTTTAACTATATATTATATTTTATAAGCACTTATTCTTCAAAGACGACGCTAGTAGAGCTGTCTAGGTTCAGCTCTTTGCCGATGTCTACACCATGACCATAGAAGACATCGTCTATATCGGCCCACTGCCTAGTAATGGGTGTTATCTTTGCCTCGACGAGGGCACGATTCACAACGTCCTTGAACCTTTTGTATTTATCAGGCCCGTGCCCATGGGCCTGCAACAGAGCAGCTTCACAGTTGACTCTAGTGGAATCTTTAAGAGATGCCCCTTTCCACACCCACTGTGTTGTGCTAACAATTGATTCCCAAGCAAGGGGGGATAAAAACTCTCCCTTACGCGTTGGGTGCAGCACGAATTTTCTTTTAAGAAACGATGCTTCTGACATATGCACCCATGCTGTTATGGCAGCGGTTTTCTCAGCAGATGTCGAAACTATGTTGTGCGACGCGAAGTATTGGTGTATAGTTATACCATTAAAATCGCCTATATAGTCATCGCTAACCGTCATTATGGCATCATCCCCATATACTATGAGGGCGACATTAGCCTTAAATGTCTCCACCCAATCGTTGCCGCCAACTATGGACTTCCATGCTAACATAATGTACATCTGGTTAACCATACTATTGACTATGGTTGTGAAGAAAGCCCCACTTGGCGAACCACCTAGTTGGCTATAAACAGTATTGTGGCATATATGTTTACTTTGAATGCACTCGTAAACTAGAGACCACATCTCTGTCTCATCAACACCTTGAACATAGTTCTTGGTCCAATCTATCATGAGAGAGTAGGCGGCTTTAGCTACTCCAGCATTGTATCCAGGCCCGAAGTTCGAGTAGTCTATTGTGATGAACTTTGTATTGACCCTAGCCAATCGCTTAATCATATCAGACCACTCTTCACTTAGCGGATTGCATCCAACGCCGTGCATCAAATTGCGGCGGGCGCTCATAAAAGAGGCAACAAAGTGTGAGAACGTCTGGCGACATGCTATAGTATAATCGGCACCCGGATTACATATTACACGTGTACCCCCCCTCTTCTTAATCTTCTCCAATGACCTTTTCTCATCCTTCAAAGTATCGATGTAGAGGGTCTCAGGTATTATTCCATTTCTACGCAAAAGCTCTTTACGAGCTATCTCGTCAGAAAGAGCGGGATCAATCCATTCCACACCCCTAACACTACCGTCACCATTACGTGCGTATGTAATGTAATCTTCTTTAGCTGTCTTCCGAGTGAGCACATAGGGAAAACCCATGCTCGTGGACAAATCCATTGGCTTGTAGTAATCAACACCCAATCCAGTGATTGCTTTATTCACTGTCAGGCGTTCTGGATTCATCACCAATGGTTTCATTGCGCAATACCACCCATCCCACAGTGCCTCCTTAACACAAGTAAGCTCTTCACTCGTGAAGTCTGTGGTCAATAAGCCGTGTTTTCTCACTCCCTCAAACAACGGAGTGGTGTCAAAGTCGTACCGCGCGTCCTTCTTGTGTAAAACAGCAGGTTGCATGTCGGTGCTCATACCACAACTGTTGTAAACCAAAGACTTAACTAGCCGTGTTGTCTGGGGTAAGTAGGGTACTTTCTCCTTAGGAAGTGAACCCCCGTAATCTAGACGAACTTCCATATCATATATTATGGAAGCGTCCATGATGGTACCGTGCTCAACTTCTTCCTTGGTGAGCACAATATCGGACTGTGTAGTGGGTAAGAGCGCCTCTTTTGTAAGGACTATGCCATAACCTGTCCCCTTATGCTCTTGACCAAGCCCAGCAACATGCATAGACATAATGGGGCGATTACTACCCTCCCTAAGTAGCAATGATCCACACGCTCCGGGTCTAGAGTAGGTGTACTCTAAACAATCACGTATCTCAAAGCATTTATTCTCCTCATCAGCTACAACGTAAGTATCGAGGAAACCTATGAGATCTACCTCGATAAGTACCATATAGTCTTCACCACGCCTAGGTACTGAGAACAAAGTGGCGTTGGCTGGTATAGATGCCTGCATTTCGTCCTCCGTAGCAAGGAACTTGCGCAAGTCCTTGAATAGGGGTTGCGATGGTTGCAACTTGAGCATAGCTATATCGGTATGCGTGGACTCCGTGATATCTTTCTCACATATCACTAGGAGCGACCTCAACTGGGGTTTGTGGAGGGGGTTACCGTATATGTTGTAACCCATCTCCATTGCTTTCCTGATTTCGTGTATGTAGTGTCTAGGTAGAAGCAAATTGTGATTGTACAACCCAACACCGTACAATACGCGTTCCTTGTTACTGCCTCTATCTATAGAGATACGGAATGTGTTATTGGCTATGTACGACTTAGCCACGGCGTACGGAGTAGGAGTTTCCTCGCCTCCTTGGAAGTACCTGACTTGTCTCTGCCTATCGTAGCGTCGGCGCATTACTCGGTGCGGTTGATTCGAATCATATTCATCAGATTGGGGTAGCGCCACTACGCTCTGTGCTGTGCAACAAATCTTATATATGATTGTCGCTATTGCTGTAGCACCAACAAGACCAGCCAGTACGTAGCTGTACTTATGGAGGGTCCCTGTTAAGTATTCCCACCACGCTTTTGGTAGTGCCACATTACTTGGCTCCACTACCCAAGCAGGTCTAAATAGTGCCGGTACAGCCCTGCACGTTTCTCCCGAAGGATCGTTAATGTAGTTAACTATAGCGGTGCGCAATACACGCGCATTAGCCTCCATGTACCTAGTAACAAAAGTGGCATACTGGGTACTGCGTGCGTAACAGCTTTCGGAGTTGCACGACGCCATGGGTACGTCAACCATAAGGTTGGTTTGCATATTGGGCACACGCCACATATTGCCGTATATCAAAGGCGACACACGCGGGTTAGACAACAACCCATGAGTACAAACAGGCAAAGGAGCACAAGCTTGCATCAAATCCTCTAATAACTGCACGTTTACAACAGGCTCCATCAAGTCCTGGGTGGTTTGTGCGTCCACATAATCAGACTGGAATACAGCTTGCACAGCAAATCCCATAAAATGGCCTGTAGCATAGCTTTCGTAGGCGTCCAACCTCCCTGTAATAAGGGACGCTAGATAGTACAAGACTGCCGTCAATGTAGGCAACGATGCCCCCTCGTATATGGTGGAAAGGTGCTTACCCAACCATGCTATCGGCATGCACCCGCTAGAAATTAGTCGAGCTGCTGCATATATAGAGTAGCCTTGCTCAGGTGTAACGCAAGGTATGACCACACCATCACGACAAGTGGGGCAGCCAGCAACGCGTTCTACCATCAAATTATTGTAACATGCAGTACATAAGTAGTGTGGGGATTCCAAGTCCACAGAGTTAGAGCACGCATATGCCACTACTCTGGTATCTAAGCACACGTTACACCTGCTAACGGTATTGGAGGTCTTATTGACGTAGTAATCCGAGATGGAGCGCAATGCTTTCATCGAGTACTTCGCTACAGTTTTCCACATTCCGCTTCCAATAGCAAGCCCTAATATAGTGTGACCCAAATTCTGCTGCCTAGGCAATTCCCAAGGCATAGCTTCAGGTTGAGATGGATTCTCAGGAGCTGGGGCAACAGAAACCTGGTTGACAATATTGCTAACCGCTATTTCTAGCTCTTCGTATGGAGTAAACGCGTTTTGAGCGAGCGACTCGTCCTCACGTATAGCAGTATTCAATTGGTAAAACAGGGTAAAGGGGTCATTTAACTGTATCTCCCTAACGTCCTGGGAACTCAAGGACGATAAAAGTCTATCCATCCGCTTCTGAACATTGGCAACTTCCGTAGCATGATAGCGTTGGAAAGTATTCTGGAGGTATGACACAGTCTGTGAAAAGGATCTAATAG

>Manitoba_toti-like_virus_3_15-14549

TTTGATTTTTTATTTTTTATTATTTACATTTTTACGGGAAATTCACGTCAATTATTGAATTTCCCAGCCCATCATGAGCCTTGATTTACAGGATCCTTAGATCCTTAACACGATAGTTGTCATAATTCGGTATACATTGTTTGTCAAATCCGGTTTACGTTTGGACACCCAATCATCGGGGGTTCCATGGTCATCATTTGTGTTGTAATCAACTGTTGTTGATTGTCCGATAAACCAGCTGTTAATAATGCGATATGTCTATCAACACTGGTCTGTTCCTAGCTGATCGAATGTATCGATCAATTGTCAATCGTATGAACTTGAGAACTTCATTGACATCTCTAACTTGCTTCAGCAAGATCGTTCTCACTGCATAGTCCATAGCATAATCTTTTATCCAAGCCAATATAGTGGGGTGCAGTTCCCCCTGGTACGTGGTACTTCCCGTTCTAATCTCGTCGATCAAAACCGATGCGACGCCTTTCCCACAGTTATCGTTGAGCTGCTGCAAGGCCAAACTAGCACTCCTCCTCAAATTTGAGTTGAAATGGGTAGAGATAGCAACCTCCGCAGCCTCAAACCTGTCCAAACCTGTAGCAACCATAGCGTTCGTCAAGCCTTTAAAAGGACAAGAATTCAAAGCCCTAAAAATAGTAGAGAGAGGCCCTACTACTTTTCTACTCATCTTAGCCAAACTACCATTCTCCAACGCGTCAAGCACTTTATTGAAATCATCAGGGGCTGGGTCGCCAGTGAGAGACCTAGCATATAATTCGCGATTCCGAGTCACCTTAGGCAGATCAAGGCTGGCCTTCCATTTGACACAAGACCTTTCCAGGCTCTTCAAACCAGCCATTCTGTCCTCCATTCGTAATGAATCCATCACGTTGGCTCTATGTAGCACATCAACCCAATTGTCAGCAAGAATAGGAGCCCTAATCTGAGTACTGGCATAGTTGACCCAGTCTTGAGACATTTTAGATTCTATGCTCCTTTCCAATTCAAGGCTCCTAAATTCCATAGACGGTATTGGCGCTACTGAGCTTGTGTGGGAGGCCATTGTCCCTGGTGGACCTAGGTCTAGTCCATTACAATCAAACCGCATTTTTAGAAACGTGACTGGAATACTGAAAGTTCCTGAGGGAAACGTTGATGTGGCAGCAAAAGGTACTACTGCTGACCACAACAAGTCTACCCCTTTCTCAGTCATCCCACGCCGATACAGTATTTGTATTTGATCATTTAACGCAGTGGCCCTCTCAGCAGGGCTAACCACCTCCACTGATTGTATCGGTTTGATTATTAGCGTGTTAACCGCTCGTGCCAGGTACCCGTTACACCCTTGACTGGTGTAGACAACTCGCAGGAATTCACCCACCTGCGTCCCAAACAATTGCTTCTCTTCCTGGAAAATCAATCCTGTCGCTTGCATAACGTTGAATAGTGCCATAGACCATAACCTCGAATCGTTACTGATCCACACATCATCTCCTTGGTGTACATTTAATAGATTGTCAGGTTTTAAACGTAGATATCTGAAGACCCAATCTCTAGCATGTCTATAATATCCAACATTCAACGTTGTATTTAAAAAGTGAGTAGCCCTACAACCTGAGAACATCCCTTGTGTGATCTTAAGTTCCTTGTTATTATTCGTGGGAAACTTGCACCATTGATTGAGTAGACCAGCTGCTACCCAGTCTGTTGCCCTGACCTTATCTGAGTGATAGCCTAGTTGTGATAGCCTTTTGGAAAGTGCTTGAAAAACAGCCGACTGTGCGGGTAGTGTGTGTTGGTAGTTAAAATCAGCATAATCTACCATAGTACACTCCTTTCTATCCTCCTTTACAGCCGAATATTTTTTGAAGACGGCATTGGTTACATCTAGCCCCAAAAGACCACTTTCAATTCCTTCAACTCTAAACATATGCTCTTCCACCTCAGATATGCTGTAGCTGGTGATGACGTAGTCGGGCGGCTTGGTCCCGTAGATGGCTCTACCTTTGCCCATTTCGAACTTCTCACTTCCTACGGCGCATATTTTCGGTTCTTCATCTAGCCAGCCAAGCATCTCCTTGGTGCTGACTCCTTCGAAATAAGCATGTTTATTAATTCTGATCGTACTACCATCACTCAAATTGGCCTTAGCACCTCCGGTACTCCCTGAAGACATCCACGACTGTCTGGTCTTACAAAAATCAAGGAACGACTTGTTTAACGGAAACGGCTTTACCATTTCCACCATGATAGTGTCAAGAACTTCATCTAGTTGTTTCAAATACAGAGCATTAGTATATTCAGATTTGTCCACCAACTCAGGCGGGCTCAAAGGCACAAAACATTCTGTACGCTTATGCTGTTCC

>Manitoba_toti-like_virus_3_13-10025

CAAATTTTATAATTTTTATTTGATTTTTATTTTTTATTATTTACATTTTTACGGGAAATTCACGTCAATTATTGAATTTCCCAGCCCATCATGAGCCTTGATTTACAGGATCCTTAGATCCTTAACACGATAGTTGTCATAATTCGGTATACATTGTTTGTCAAATCCGGTTTACGTTTGGACACCCAATCATCGGGGGTTCCATGGTCATCATTTGTGTTGTAATCAACTGTTGTTGATTGTCTGATAAACCAGCTGTTAATAATGCGATATGTCTATCAACACTGGTCTGTTCCTAGCTGATCGAATGTATCGATCAATTGTCAATCGTATGAACTTGAGAACTTCATTGACATCTCTAACTTGCTTCAGCAAGATCGTTCTCACTGCATAGTCCATAGCATAATCTTTTATCCAAGCCAATATAGTGGGGTGCAGTTCCCCCTGGTACGTGGTACTTCCCGTTCTAATCTCGTCGATCAAAACCGATGCGACGCCTTTCCCACAGTTATCGTTGAGCTGCTGCAAGGCCAAACTAGCACTCCTCCTCAAATTTGAGTTGAAATGGGTAGAGATAGCAACCTCCGCAGCCTCAAACCTGTCCAAACCTGTAGCAACCATAGCGTTCGTCAAGCCTTTAAAAGGGCAAGAATTCAAAGCCCTAAAAATAGTAGAGAGAGGCCCTACTACTTTTCTACTCATCTTAGCCAAACTACCATTCTCCAACGCGTCAAGCACTTTATTGAAATCATCAGGGGCTGGGTCGCCAGTGAGAGACCTAGCATATAATTCGCGATTCCGAGTCACCTTAGGCAGATCAAGGCTGGCCTTCCATTTGACACAAGACCTTTCCAGGCTCTTCAAACCAGCCATTCTGTCCTCCATTCGTAATGAATCCATCACGTTGGCTCTATGTAGCACATCAACCCAATTGTCAGCAAGAATAGGAGCCCTAATCTGAGTACTGGCATAGTTGACCCAGTCTTGAGACATTTTGGATTCTATGCTCCTTTCCAATTCAAGGCTCCTAAATTCCATAGACGGTATTGGCGCTACTGAGCTTGTGTGGGAGGCCATTGTCCCTGGTGGACCTAGGTCTAGTCCATTACAATCAAACCGCATTTTTAGAAACGTGACTGGAATACTGAAAGTTCCTGAGGGAAAAGTTGATGTGGCAGCAAAAGGTACTACTGCTGACCACAACAAGTCTACCCCTTTCTCAGTCATCCCACGCCGATACAGTATTTGTATTTGATCATTTAACGCAGTGGCCCTCTCAGCAGGGCTAACCACCTCCACTGATTGTATCGGTTTGATTATTAGCGTGTTAACCGCTCGTGCCAGGTACCCGTTACACCCTTGACTGGTGTAGACAACTCGCAGGAATTCACCCACCTGCGTCCCAAACAATTGCTTCTCTTCCTGGAAAATCAATCCTGTCGCTTGCATAACGTTGAATAGTGCCATAGACCATAACCTCGAATCGTTACTGATCCACACATCATCTCCTTGGTGTACATTTAATAGATTGTCAGGTTTTAAACGTAGATATCTGAAGACCCAATCTCTAGCATGTCTATAATATCCAACATTCAACGTTGTATTTAAAAAGTGAGTAGCCCTACAACCTGAGAACATCCCTTGTGTGATCTTTAGTTCCCTGTTATTATTCGTGGGAAACTTGCACCATTGATTGAGTAGACCAGCTGCCACCCAGTCTGTTGCCCTGACCTTATCTGAGTGATAGCCTAGTTGTGATAGCCTTTTGGAAAGTGCTTGAAAAACAGCCGACTGTGCAGGTAGTGTGTGTTGGTAGTTAAA

>Manitoba_toti-like_virus_3_14-20694

CCCAGCCCATCATGAGCCTTGATTTACAGGACCCTTAGATCCTTAACACGATAGTTGTCATAATTCGGTATACATTGTTTGTCAAATCCGGTTTACGTTTGGACACCCAATCATCGGGGGTTCCATGGTCATCATTTGTGTTGTAATCAACTGTTGTTGATTGTCTGATAAACCAGCTGTTAATAATGCGATATGTCTATCAACACTGGTCTGTTCCTAGCTGATCGAATGTATCGATCAATTGTCAATCGTATGAACTTGAGAACTTCATTGACATCTCTAACTTGCTTCAGCAAGATCGTTCTCACTGCATAGTCCATAGCATAATCTTTTATCCAAGCCAATATAGTGGGGTGCAGTTCCCCCTGGTACGTGGTACTTCCCGTTCTAATCTCGTCGATCAAAACCGATGCGACGCCTTTCCCACAGTTATCGTTGAGCTGCTGCAAGGCCAAACTAGCACTCCTCCTCAAATTTGAGTTGAAATGGGTAGAGATAGCAACCTCCGCAGCCTCAAACCTGTCCAAACCTGTAGCAACCATAGCGTTCGTCAAGCCTTTAAAAGGACAAGAATTCAAAGCCCTAAAAATAGTAGAGAGAGGCCCTACTACTTTTCTACTCATCTTAGCCAAACTACCATTCTCCAACGCGTCAAGCACTGTATTGAAATCATCAGGGGCTGGGTCGCCAGTGAGAGACCTAGCATATAATTCGCGATTCCGAGTCACCTTAGGCAGATCAAGGCTGGCCTTCCATTTGACACAAGACCTTTCCAGGCTCTTCAAACCAGCCATTCTGTCCTCCATTCGTAATGAATCCATCACGTTGGCTCTATGTAGCACATCAACCCAATTGTCAGCAAGAATAGGAGCCCTAATCTGAGTACTGGCATAGTTGACCCAGTCTTGAGACATTTTAGATTCTATGCTCCTTTCCAATTCAAGGCTCCTAAATTCCATAGACGGTATTGGCGCTACTGAGCTTGTGTGGGAGGCCATTGTCCCTGGTGGACCTAGGTCTAGTCCATTACAATCAAACCGCATTTTTAGAAACGTGACTGGGATACTGAAAGTTCCTGAAGGAAAAGTTGATGTGGCAGCAAAAGGTACTACTGCTGACCACAACAAGTCTACCCCTTTCTCAGTCATCCCACGCCGATACAGTATTTGTATTTGATCATTTAACGCAGTGGCCCTCTCAGCAGGGCTAACCACCTCCACTGATTGTATCGGTTTGATTATTAGCGTGTTAACCGCTCGTGCCAGGTACCCGTTACACCCTTGACTGGTGTAGACAACTCGCAAGAACTCACCCACCTGCGTCCCAAACAATTGCTTCTCTTCCTGGAAAATCAATCCTGTCGCTTGCATAACGTTGAATAGTGCCATAGACCATAACCTCGAATCGTTACTGATCCACACATCATCTCCTTGGTGTACATTTAATAGATTGTCAGGTTTTAAACGTAGATATCTGAAGACCCAATCTCTAGCATGTCTATAATATCCAACATTCAACGTTGTATTTAAAAAGTGAGTAGCCCTACAACCTGAGAACATCCCTTGTGTGATCTTTAGTTCCCTGCTATTATTCGTGGGAAACTTGCACCATTGATTGAGTAGACCAGCTGCTACCCAGTCTGTTGCCCTGACCTTATCTGAGTGATAGCCTAGTTGTGATAGCCTTTTGGAAAGTGCTTGAAAAACAGCCGACTGTGCAGGTAGTGTGTGTTGGTAGTTAAAATCAGCATAATCTACCATAGTACACTCCTTTCTATCCTCCTTTACAGCCGAATATTTTTTGAAGACGGCATTGGTTACATCTAGCCCCAAAAGACCACTTTCAATTCCTTCAACTCTAAACATATGCTCTTCCACCTCAGATATGCTGTAGCTGGTGATGACGTAGTCGGGCGGCTTGGTCCCGTAGATGGCTCTACCTTTGCCCATTTCGAACTTCTCACTTCCTACGGCGCATATTTTCGGTTCTTCATCTAGCCAGCCAAGCATCTCCTTGGTGCTGACTCCTTCGAAATAAGCATGTTTATTAATTCTGATCGTACTACCATCACTCAAATTGGCCTTAGCACCTCCGGTACTCCCTGAAGACATCCACGACTGTCTGGTCTTACAAAAATCAAGGAACGACTTGTTTAACGGAAACGGCTTTACCATTTCCACCATGATAGTGTCAAGAACTTCATCTAGTTGTTTCAAATACAGAGCATTAGTATATTCAGA
