## Supplementary Figures: S1, S2, S3 for "Metatranscriptomic analysis of common mosquito vector species in the Canadian Prairies"

**Supplementary Figure S1**. Total number of viral reads from previously reported viruses identified in each mosquito species. Viruses are sorted by family and colour coded based on their genome configuration. Also displayed is the number of sequencing libraries for each species.

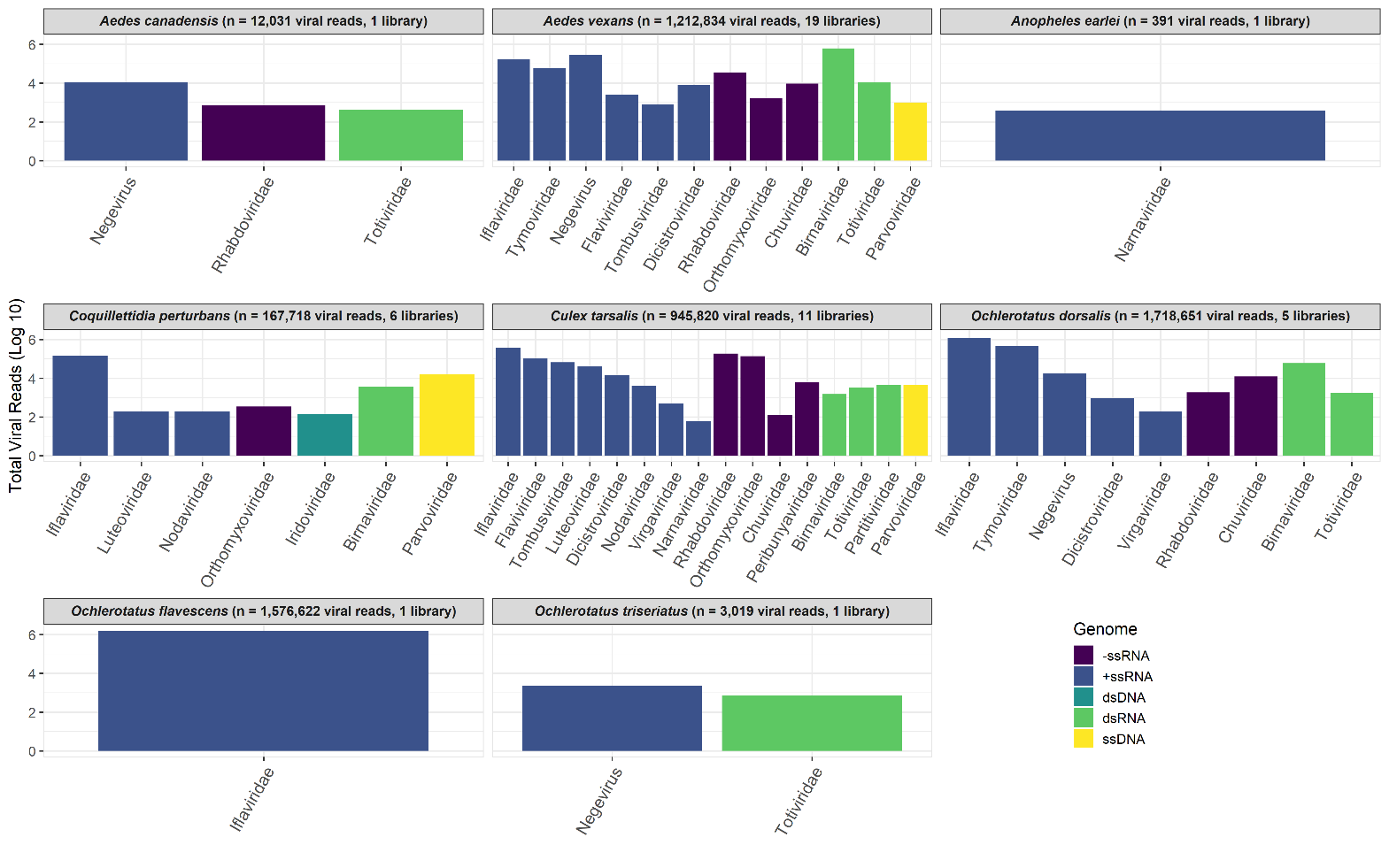

**Supplementary Figure S2**. Number of novel viruses identified for each mosquito species (A). Viruses are sorted by family and colour coded based on their genome configuration. Also displayed is the total numbers of viruses detected and sequencing libraries for each species. Also displayed is a Venn diagram showing the partitioning of novel viruses shared among mosquito genera.

**(A)**

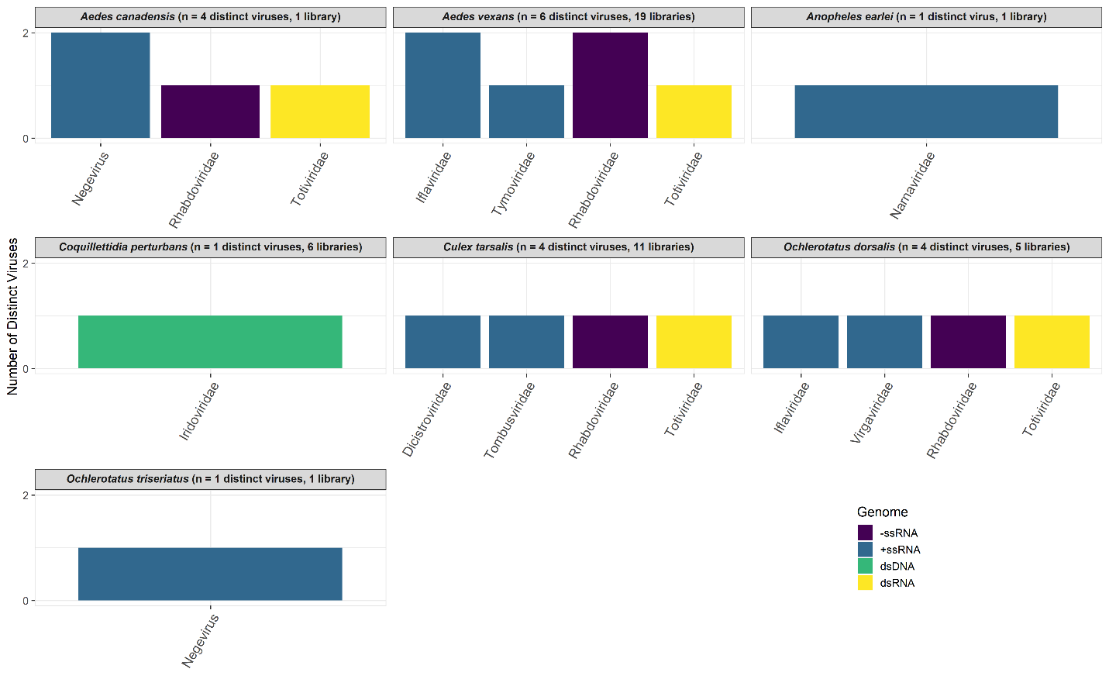

**(B)**

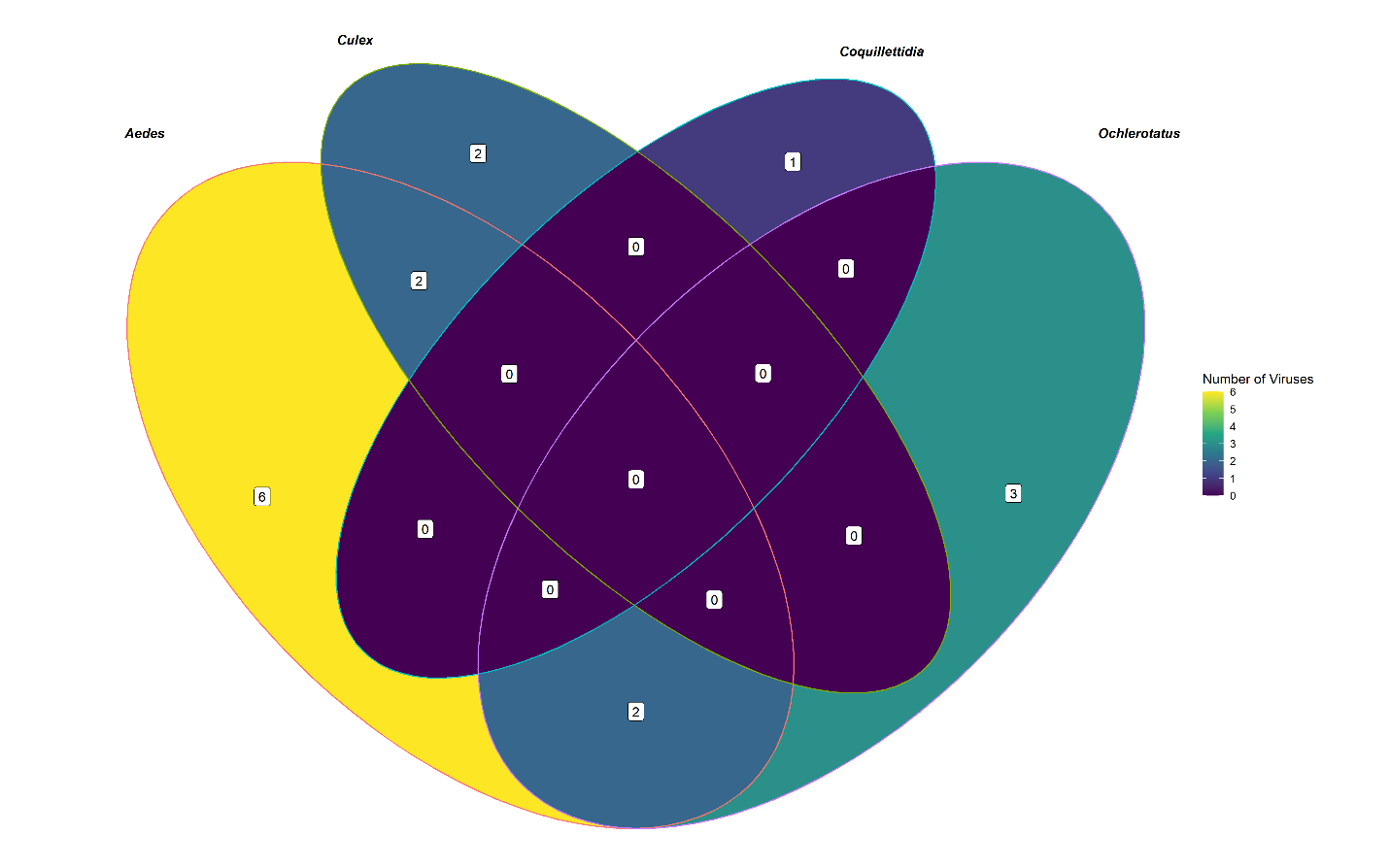

**Supplementary Figure S3**. Non viral, non-host organisms identified in Canadian Prairie mosquitoes. Displayed is the number of total sequencing reads (A) and number of organisms detected (B) for the various groups of fungi, invertebrate parasites/protozoa, plants, and vertebrates. The bars are colour-coded based on the mosquito species each organism was identified in.

**A)**
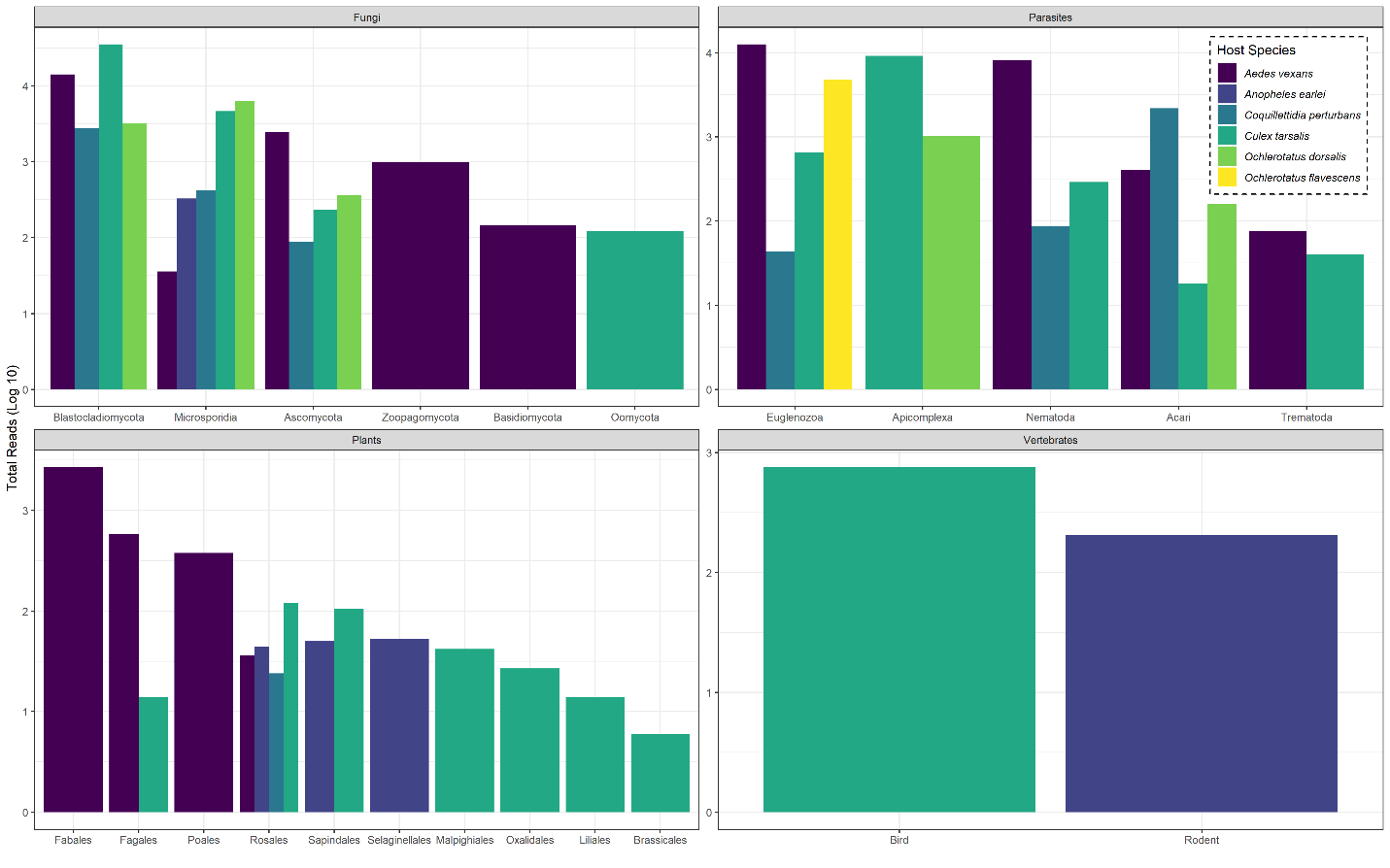

**B)**
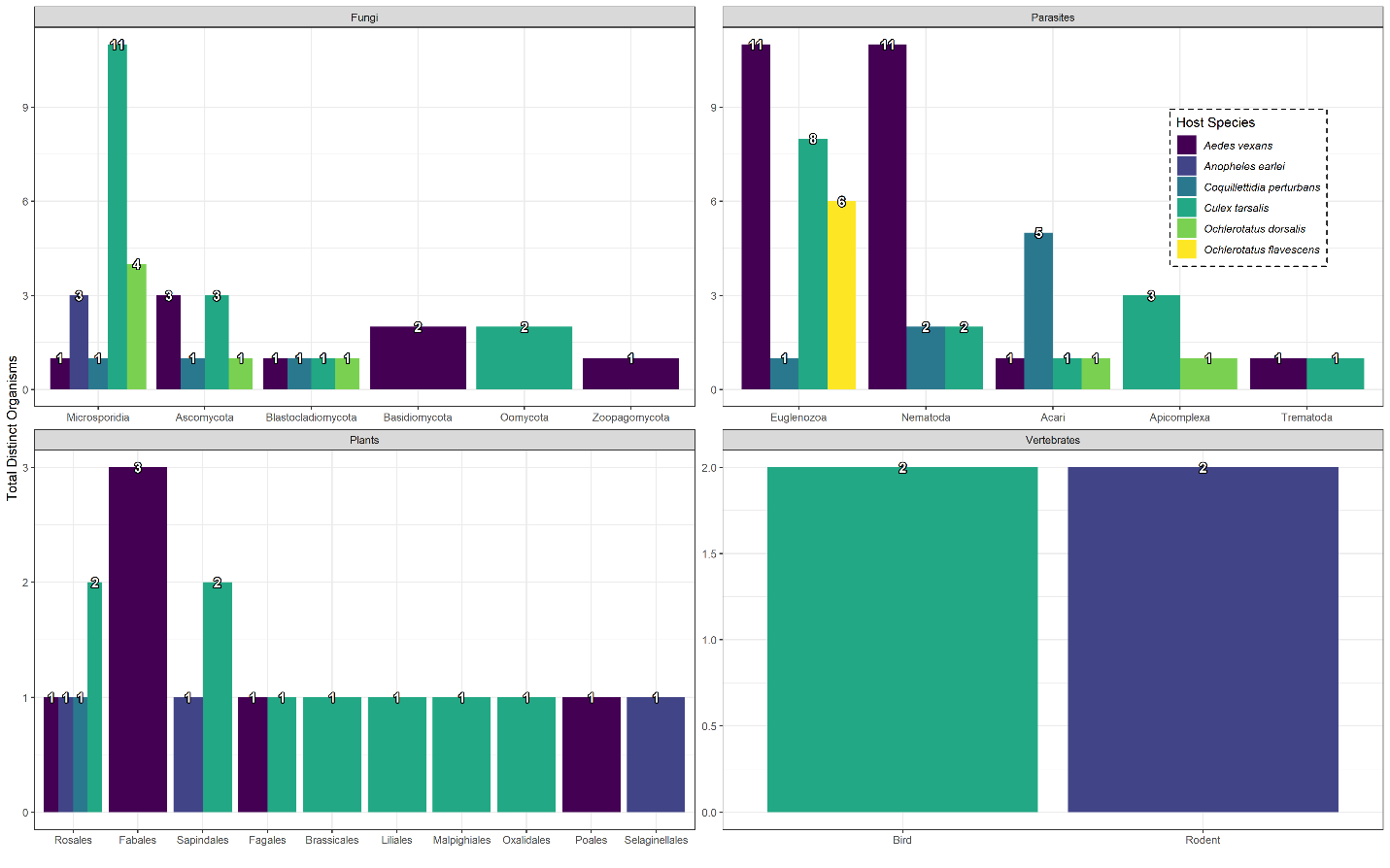
